# Integrative omics approach to identify the molecular architecture of inflammatory protein levels in healthy older adults

**DOI:** 10.1101/2020.02.17.952135

**Authors:** Robert F. Hillary, Daniel Trejo-Banos, Athanasios Kousathanas, Daniel L. McCartney, Sarah E. Harris, Anna J. Stevenson, Marion Patxot, Sven Erik Ojavee, Qian Zhang, David C. Liewald, Craig W. Ritchie, Kathryn L. Evans, Elliot M. Tucker-Drob, Naomi R. Wray, Allan F. McRae, Peter M. Visscher, Ian J. Deary, Matthew R. Robinson, Riccardo E. Marioni

## Abstract

The molecular factors which control circulating levels of inflammatory proteins are not well understood. Furthermore, association studies between molecular probes and human traits are often performed by linear model-based methods which may fail to account for complex structure and interrelationships within molecular datasets. Therefore, in this study, we perform genome- and epigenome-wide association studies (GWAS/EWAS) on the levels of 70 plasma-derived inflammatory protein biomarkers in healthy older adults (Lothian Birth Cohort 1936; n = 876; Olink^®^ inflammation panel). We employ a Bayesian framework (BayesR+) which can account for issues pertaining to data structure and unknown confounding variables (with sensitivity analyses using ordinary least squares- (OLS) and mixed model-based approaches). We identified 13 SNPs associated with 13 proteins (n = 1 SNP each) concordant across OLS and Bayesian methods. We identified three CpG sites spread across three proteins (n = 1 CpG each) that were concordant across OLS, mixed-model and Bayesian analyses. Tagged genetic variants accounted for up to 45% of variance in protein levels (for MCP2, 36% of variance alone attributable to one polymorphism). Methylation data accounted for up to 46% of variation in protein levels (for CXCL10). Up to 66% of variation in protein levels (for VEGFA) was explained using genetic and epigenetic data combined. We demonstrated putative causal relationships between CD6 and IL18R1 with inflammatory bowel disease, and between IL12B and Crohn’s disease. Our data may aid understanding of the molecular regulation of the circulating inflammatory proteome as well as causal relationships between inflammatory mediators and disease.

## Introduction

Inflammation represents a concerted cascade of molecular and cellular events to combat infectious pathogens and endogenous insults. Inflammatory proteins are key mediators of defence and repair responses, and tight spatiotemporal regulation of their plasma concentrations permits effective immune activation and resolution (1). Whereas acute inflammatory states may prompt severe illness and death, absence of resolution precipitates transition from acute to deleterious chronic inflammatory states (2). Chronic inflammation facilitates the pathogenesis of various disease states, including diabetes, heart disease, stroke and allergic conditions (3). Furthermore, inflammatory lesions in brain tissue are often associated with, and may contribute to, neurodegeneration and cognitive decline (4). Globally, 60% of individuals will die as a consequence of a chronic inflammation-associated disease state (5). Therefore, identifying biological factors which govern inter-individual variation in circulating inflammatory protein levels may allow for better prediction of individual disease risk and prognosis, and inform disease biology.

To date, few studies have characterised genetic factors which are associated with circulating inflammatory protein levels (also known as protein quantitative trait loci or pQTLs) (n = 9-83 proteins, 65-6,861 individuals) (6-8). Zaghlool *et al.* performed an epigenome-wide association study (EWAS) of 1123 proteins, which pointed towards networks of chronic low-grade inflammatory biomarkers (n = 944 individuals) (9). In an integrative approach, Ahsan *et al.* identified epigenetic, as well as genetic, markers associated with protein biomarkers including inflammatory mediators (n ≤ 1,033 individuals) (10). Notably, most studies examining the molecular architecture of human traits have relied on linear model-based methods which examine marker or probe effects marginally (11, 12). This fails to account for issues including correlation structure within molecular datasets, data structure (i.e. cellular heterogeneity, batch effects) and omitted variable bias which together may result in overfitting as well as poor and biased estimates of effect sizes. A number of approaches have been proposed to address these issues (13-19) and these encompass strategies which permit the joint and conditional estimation of effect sizes while accounting for correlations among markers and confounding variables. Here, we consider a Bayesian penalised regression framework termed BayesR+ which was developed to assess genetic and epigenetic architectures of complex traits (20). In BayesR+, marker effects (polymorphism or CpG site) can be estimated jointly while controlling for data structure and correlations among molecular markers of different types. Indeed, this method permits the estimation of variance explained in the trait by all methylation probes or genetic markers, either separately or together.

In the present study, we use the BayesR+ method (and sensitivity analyses using ordinary least squares (OLS) (21, 22) and mixed model methods (23)) to examine both the genetic and epigenetic architectures of 70 blood inflammatory proteins in 876 relatively healthy older adults from the Lothian Birth Cohort 1936 study (mean age: 69.8 ± 0.8 years; levels adjusted for age, sex, population structure and array plate). Hereinafter we refer to the adjusted inflammatory protein levels as protein levels. These proteins are present on the Olink^®^ inflammation panel and comprise a mixture of proteins with defined functions pertinent to human inflammatory pathways as well as putative roles in inflammation-related disease states. Applying a stringent approach, we consider only markers or probes concordantly identified across methods and integrate multiple levels of ‘omics’ data to investigate mechanisms by which pQTLs may influence protein levels. Finally, we use our GWAS summary data to test for putatively causal relationships between inflammatory protein biomarkers and neurological or inflammatory disease states.

## Results

### Genome wide studies of inflammatory protein levels

In a Bayesian penalised regression model (BayesR+), 16 pQTLs were identified for 14 proteins (Supplementary Table 1). Thirteen of these 16 pQTLs (n = 13 proteins) directly, or through variants in high linkage disequilibrium (LD): *r*^2^ > 0.75, replicated conditionally significant pQTLs from the OLS model (Supplementary Note 1; Supplementary Tables 2-4). The correlation structure among these 13 proteins is shown in Supplementary Figure 1.

Twelve (92.30%) of the concordant SNPs were *cis* pQTLs (SNP within 10 Mb of the transcription start site (TSS) of a given gene (24, 25)) and 1 pQTL (7.70%) was a *trans*-associated variant (Figure 1A; Supplementary Table 5). There was an inverse relationship between the minor allele frequency of variants and their effect size (Figure 1B). The majority of SNPs were located in exonic regions (38.5%), as identified by FUMA (**FU**nctional **M**apping and **A**nnotation analysis) (Figure 1C). Four of the five SNPs annotated to exonic regions result in missense mutations. From the Bayesian model, pQTLs explained between 5.28% (rs10005565; CXCL6) and 35.80% (rs3138036; MCP2) of inter-individual variation in protein levels (Figure 1D). The estimates for variance accounted for in protein levels by single SNPs were correlated 99% between the BayesR+ and OLS regression models (Figure 2A; Supplementary Table 5). The BayesR+ common (minor allele frequency > 1%) SNP-based heritability estimates ranged from 11.4% (CXCL9; 95% credible interval: [0%, 43.5%]) to 45.3% (MCP2; 95% credible interval: [23.5%, 70.6%]), with a mean estimate of 20.2% across the 70 proteins (Supplementary Table 6). Figure 2B shows heritability estimates for the 13 proteins exhibiting concordantly identified pQTLs across OLS and Bayesian approaches. Figure 3 demonstrates the effect of genetic variation at the most significant *cis* pQTL (rs3138036; MCP2) and the sole *trans* pQTL (rs12075; MCP4) on protein levels.

**Figure 1.**
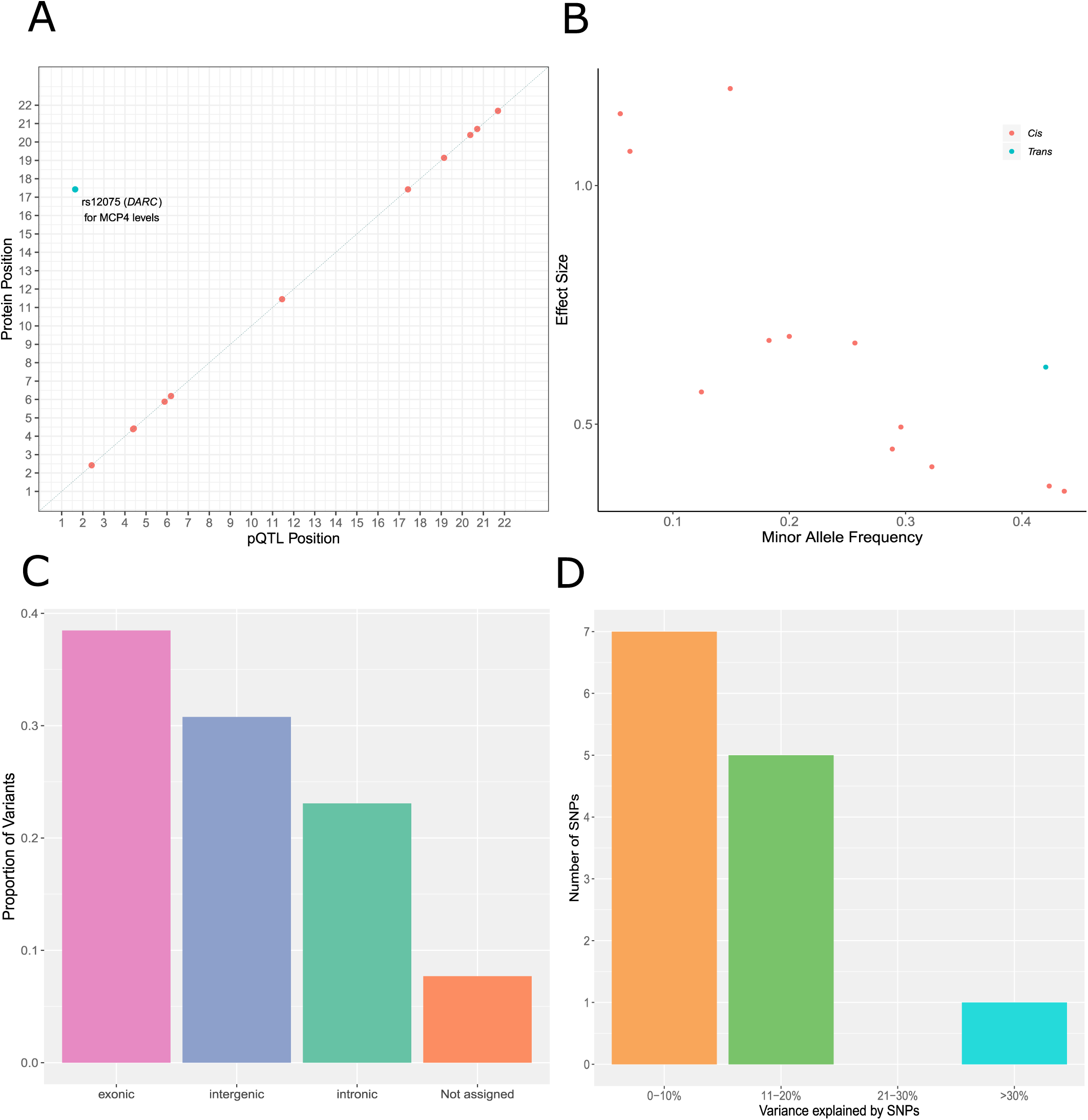
Genetic architecture of inflammatory protein biomarkers in the Lothian Birth Cohort 1936. (A) Chromosomal locations of pQTLs concordant between Bayesian penalised and ordinary least squares regression models for genome-wide association studies (n = 13 pQTLs). The x-axis represents the chromosomal location of concordantly identified *cis* and *trans* SNPs associated with the levels of Olink^®^ inflammatory proteins. The y-axis represents the position of the gene encoding the associated protein. The sole conditionally significant concordant *trans* association is annotated. *Cis* (red circles); *trans* (blue circles). (B) Absolute effect size (per standard deviation of difference in protein level per effect allele) of pQTLs versus minor allele frequency. *Cis* (red circles); *trans* (blue circles). (C) Classification of 13 pQTLs by function as defined by functional enrichment analysis in FUMA. (D) Variance in protein levels explained by pQTLs (estimates from Bayesian penalised regression are displayed).

**Figure 2.**
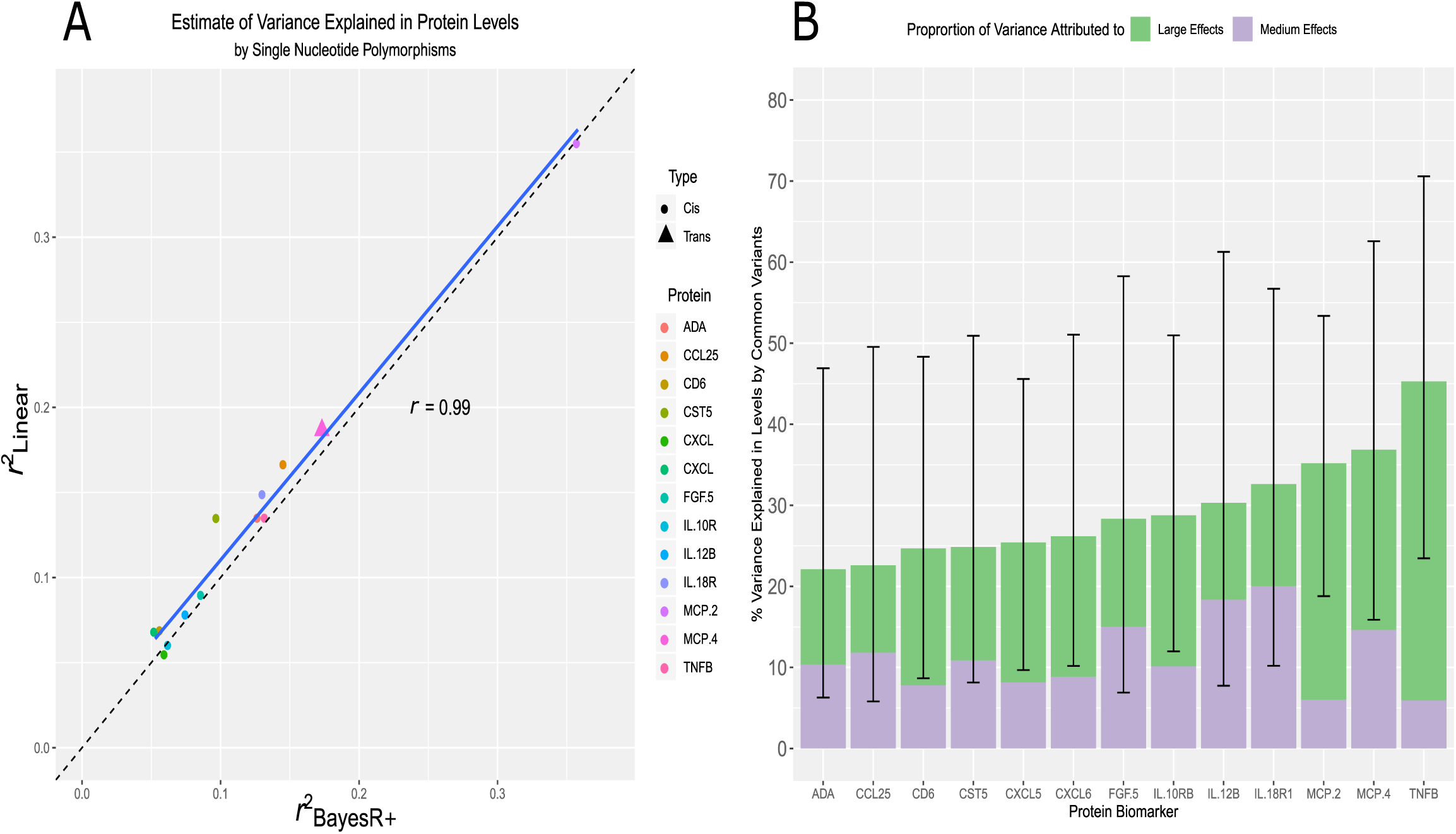
(A) In this panel, the variance explained (*r*^2^) by consensus SNPs (n = 13 SNP, 1 per protein) in the ordinary least squares regression model was compared against the variance explained by the same SNP set identified in the Bayesian penalised regression approach. (B) The proportion of variance explained in Olink^®^ inflammatory protein levels by common genetic variants genotyped in the LBC1936 participants is shown. Only those proteins which had significant pQTL associations in both the ordinary least squares and Bayesian methods are presented (n = 13). Additionally, the proportion of variance explained attributable to medium effects (prior: variance of 1% explained) and large effects (prior: variance of 10% explained) are demonstrated in purple and green, respectively. Error bars represent 95% credible intervals.

**Figure 3.**
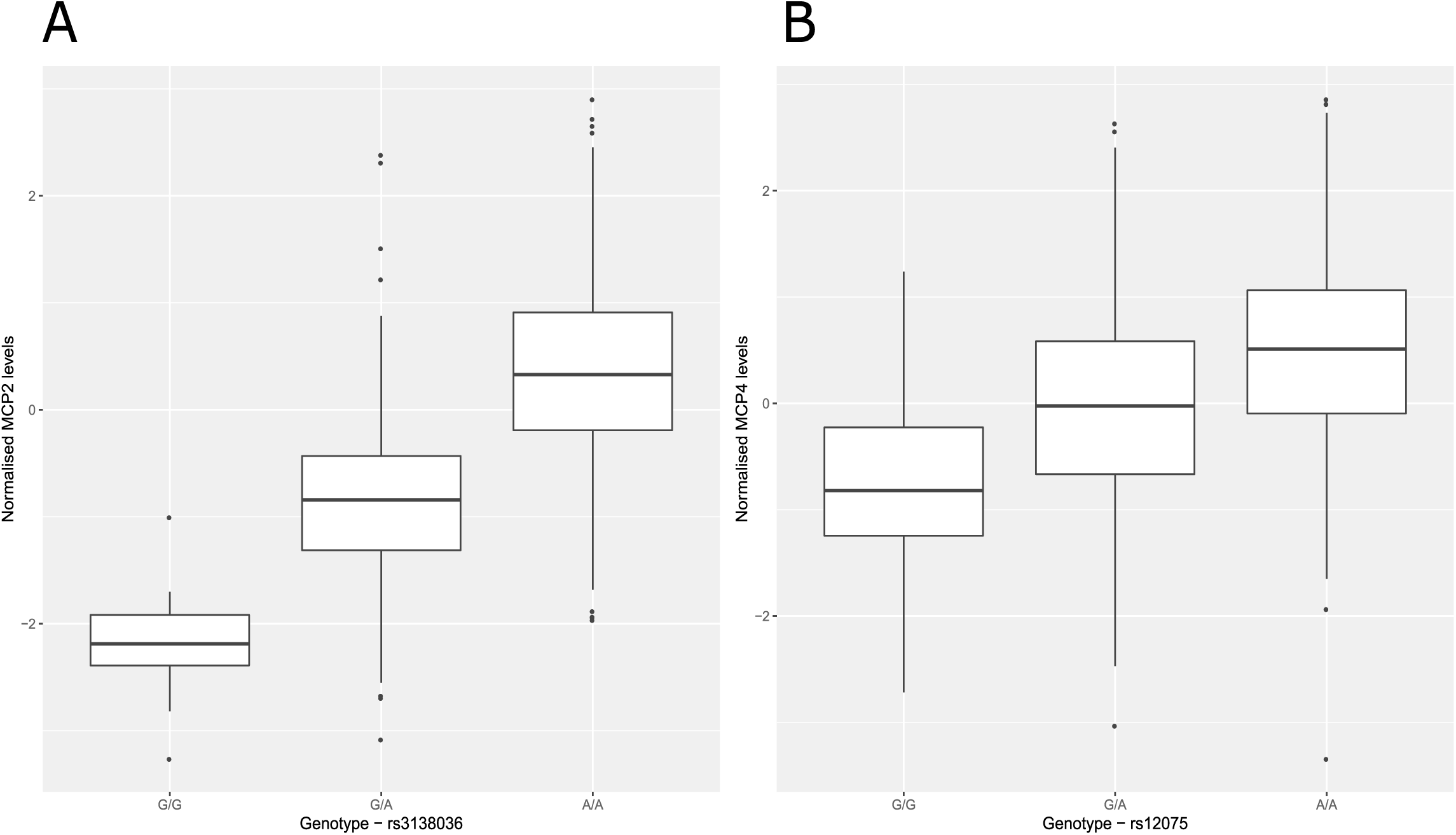
Effect of genetic variation on inflammatory protein levels. (A) Box plot of MCP2 levels as a function of genotype (rs3138036, effect allele: G, other allele: A, beta = −1.20, se = 0.06). (B) Box plot of MCP4 levels as a function of genotype (rs14075, effect allele: G, other allele: A, beta = −0.62, se = 0.05). Centre line of boxplot: median, bounds of box: first and third quartiles and tips of whiskers: minimum and maximum.

There was a strong correlation between our SNP-based heritability estimates and those from a previous study of 961 individuals (10): 29 overlapping proteins, *r*: 0.71, 95% CI: [0.43, 0.84] (Supplementary Figure 2 and Supplementary Table 7).

### Molecular mechanisms underlying pQTLs: colocalisation analysis

Of the 12 *cis* pQTLs which were identified across OLS regression and BayesR+, 8 SNPs (66.67%) previously have been identified as *cis*-acting expression QTLs (eQTLs) in blood (Supplementary Table 8). Using *coloc* (26), we tested the hypothesis that one causal variant might underlie both a pQTL and eQTL for each protein. For 4/8 proteins, there was strong evidence (posterior probability (PP) > 0.95) for colocalisation of *cis* pQTLs and *cis* eQTLs (Supplementary Table 9). These proteins were CCL25, CD6, CXCL5 and CXCL6.

Mendelian Randomisation analyses (MR; see methods) indicated that altered gene expression was causally associated with changes in protein levels for each of the four aforementioned proteins (CCL25, CD6, CXCL5 and CXCL6; range of beta: [0.68, 12.25], se: [0.09, 1.12], P: [9.54 × 10^−7^, 1.05 × 10^−37^]). However, a second colocalisation approach termed Sherlock (27) suggested that, from the 13 proteins with concordantly identified pQTLs, only expression of *ADA, CXCL5* and *IL18R1* were associated with levels of their respective protein products (Supplementary Note 1; Supplementary Table 10).

### Epigenome wide studies of inflammatory protein levels

In the Bayesian model, 8 CpG-protein associations (n = 8 proteins) had a posterior inclusion probability of greater than 95% (Supplementary Table 11). Five of these associations overlapped with those identified from the OLS regression model (P < 5.14 × 10^−10^; Supplementary Table 12); three of which were also identified in the mixed model approach (P < 5.14 × 10^−10^; Supplementary Table 13). These were: the smoking-associated probe cg05575921 for CCL11 levels (*trans* association at *AHRR*; mixed model - beta: −1.97, se: 0.31, P: 4.86 × 10^−10^), cg07839457 for CXCL9 levels (*trans* association at *NLRC5*; beta: −2.91, se: 0.39, P: 8.03 × 10^−14^) and cg03938978 for IL18R1 levels (*cis* association at *IL18RAP*; beta: −1.37, se: 0.16, P: 5.86 × 10^−17^) (Supplementary Table 13). Adjustment for smoking attenuated the association between CCL11 levels and the cg05575921 probe (linear model - before adjustment: beta: −0.02, P: 2.68 × 10^−10^, after adjustment: beta: 0.003, P: 0.15; % attenuation: 88.60%). Figure 4 depicts an epigenetic map of CpG-protein associations within this study and demonstrates the degree of overlap between methodologies. The correlation among the three proteins with concordantly identified CpG associations is shown in Supplementary Figure 3. A lookup analysis of the top GWAS and EWAS findings with those reported in the literature showed moderate to strong concordance (Supplementary Note 1).

**Figure 4.**
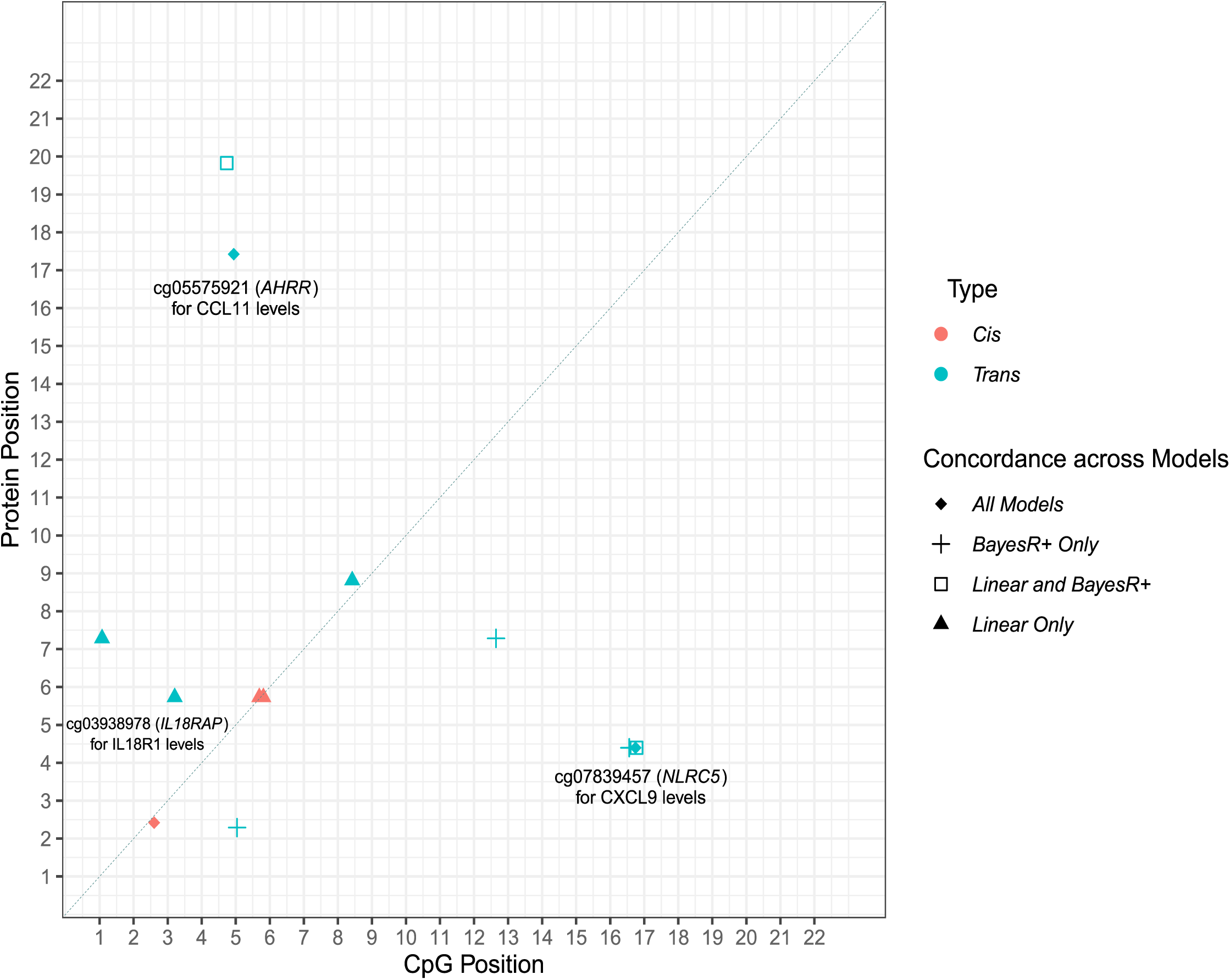
Genomic locations of CpG sites associated with differential inflammatory protein levels. The x-axis represents the chromosomal location of CpG sites associated with the levels of Olink^®^ inflammatory biomarkers. The y-axis represents the position of the gene encoding the associated protein. The level of concordance across three models used to perform epigenome-wide association studies is represented by different shape patterns. Those CpG sites (n = 3) which were identified by linear (ordinary least squares), mixed-model and Bayesian penalised regression models, and passed a Bonferroni-corrected significance threshold, are represented by diamonds and annotated. Three proteins (CXCL9, CXCL10 and CXCL11) were associated with differential methylation levels at the cg07839457 site in the *NLRC5* transcription factor locus. Additionally, two proteins (CCL11 and TGF-alpha) were associated with the smoking-associated cg05575921 site in the *AHRR* locus. *Cis* (red); *trans* (blue).

We conducted tissue specificity and pathway enrichment analyses based on genes identified by methylation for each of the 3 proteins with significant CpG associations. Tissue-specific patterns of expression were observed for 2/3 proteins (Supplementary Figures 4-6). For CCL11, differential expression was observed in breast, adipose and kidney tissue. For IL18R1, differential expression of associated genes was observed in pancreatic tissue. Furthermore, down-regulation of genes associated with IL18R1 was observed in the hippocampus and substantia nigra. There was no significant enrichment of pathways incorporating genes annotated to CXCL9, CCL11 or IL18R1 following multiple testing correction.

One protein, IL18R1, harboured both a significant *cis* pQTL and *cis* CpG site in our study (Supplementary Figure 7). This SNP (rs917997) previously has been identified as a methylation QTL (mQTL) for the single *cis* CpG site associated with IL18R1 levels identified by our epigenome-wide studies (cg03938978) (28). Using bidirectional MR analysis (Wald ratio test; see methods), we found that DNA methylation at this locus was causally associated with circulating IL18R1 levels (beta: −0.81, se: 0.17, P: 2.14 × 10^−33^). Conversely, IL18R1 levels were also causally associated with altered DNA methylation (beta: −1.22, se: 0.16, P: 3.4 × 10^−14^).

The methylation data explained an average of 18.2% of variance in protein levels using BayesR+; estimates ranged from 6.3% (IL15RA, 95% credible interval: [0.0%, 27.3%]) to 46.1% (CXCL10, 95% credible interval: [24.1%, 67.1%]) (Supplementary Table 14). There was strong concordance with estimates from the mixed model sensitivity analysis (Supplementary Table 15 and Fig 5A). Figure 5B shows the variance explained by methylation data for the 3 proteins exhibiting concordantly identified CpGs across OLS regression, mixed model and Bayesian approaches.

**Figure 5.**
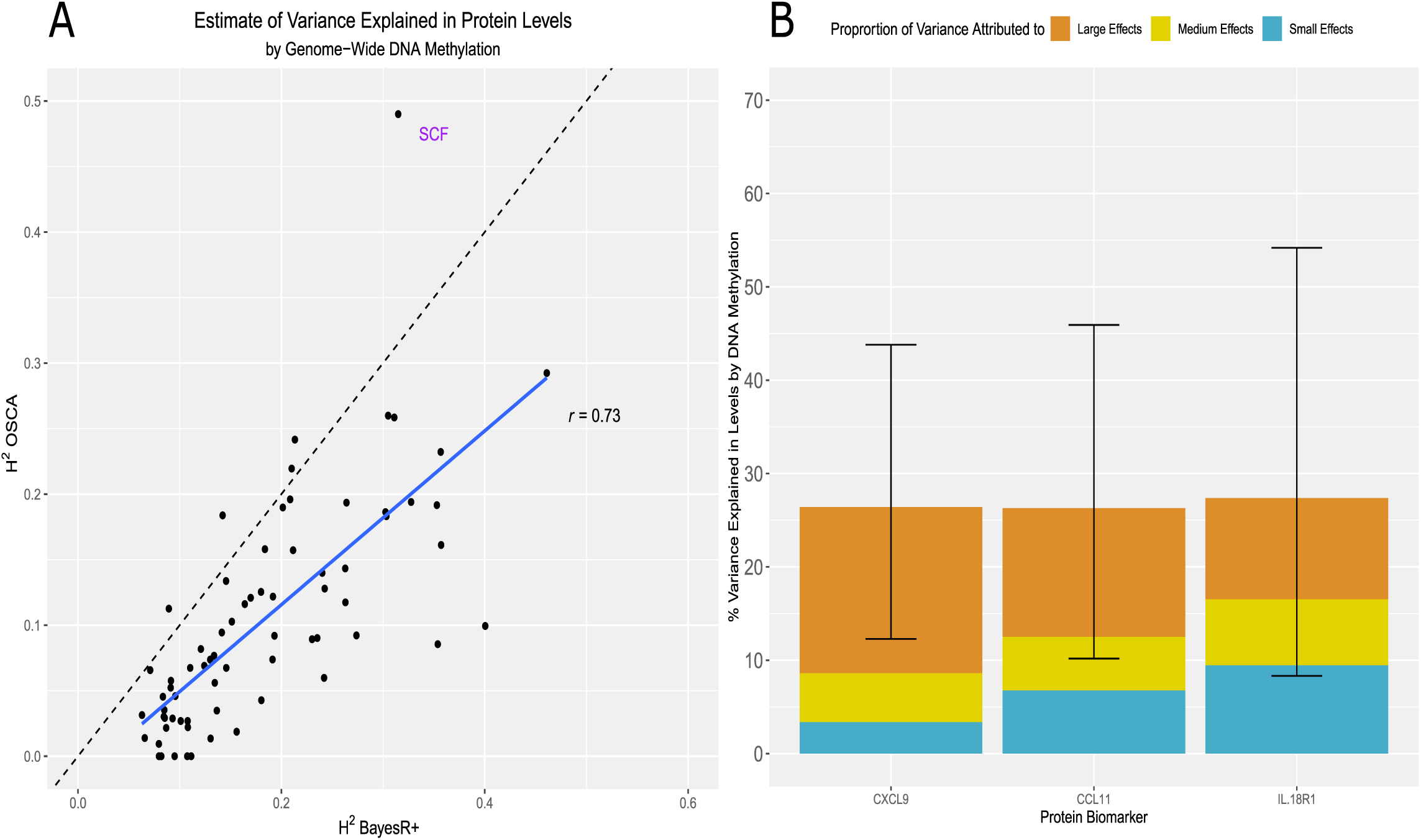
Variance in circulating inflammatory protein levels explained by DNA methylation. (A) In this panel, the variance explained in circulating protein levels by complete methylation data from sites present on the Infinium 450 K methylation array was examined. A comparison between variance explained (H^2^) by a mixed model approach (OSCA) and a Bayesian penalised regression approach (BayesR+) is shown. (B) The proportion of variance explained in Olink^®^ inflammatory protein levels by DNA methylation, as estimated by BayesR+, is shown. Only those proteins (n = 3) which had significant CpG associations in ordinary least squares, mixed model and Bayesian methods are presented. Additionally, the proportion of variance explained attributable to small effects (prior: variance of 0.1% explained), medium effects (prior: variance of 1.0% explained) and large effects (prior: variance of 10% explained) are demonstrated in blue, gold and dark orange, respectively. Error bars represent 95% credible intervals.

### Variation in inflammatory protein levels explained by genetics and DNA methylation

When accounting for genetic data, the estimates for variance explained by methylation data were largely unchanged for most proteins (Supplementary Table 16; n = 9 proteins with change > 5%, 1 with change < −5% (VEGFA)). The mean absolute change was 2.6% (minimum: 0.01% for TNFRSF9 and maximum: 15.0% for IL18R1). Similarly, estimates from genetic data were largely unchanged in the combined analysis (n = 2 proteins with change > 5%). The mean absolute change was 1.8% (minimum: 0.02% for CD244 and maximum: 6.7% for CCL28). For 22 proteins, the variance explained by methylation data was greater than that explained by genetic data (Supplementary File 1).

The combined estimate for variance explained by genetic and methylation data ranged from 23.4% for CXCL1 to 66.4% for VEGFA. The mean and median estimates were 37.7% and 36.0%, respectively.

### Evaluating causal associations between inflammatory biomarkers and human traits

The 13 independent pQTL associations were queried against GWAS Catalog to identify existing associations between these pQTLs and phenotypes (29). We investigated whether these associations represented causal relationships. Using two-sample MR, we showed that CD6 levels were causally associated with inflammatory bowel disease (IBD) (beta: 0.20, se: 0.04, P: 2.59 × 10^−6^). Furthermore, FGF-5 levels were causally associated with systolic and diastolic blood pressure (beta: 0.07 and 0.07, se: 0.01 and 0.01, P: 1.04 × 10^−34^ and 4.29 × 10^−42^, respectively). IL12B levels were associated with Crohn’s disease (beta: 0.42, se: 0.05, P: 2.76 × 10^−15^). Circulating IL18R1 levels showed a causal relationship with IBD (beta: 0.17, se: 0.03, P: 1.63 × 10^−9^).

Peripheral inflammatory processes and proteins have been linked to risk of late-onset Alzheimer’s disease (AD) (30, 31). We tested whether the 13 proteins with significant genetic correlates in our study were causally associated with AD risk (Supplementary Note 1). One protein, IL18R1, showed a nominally significant, unidirectional relationship with AD risk (beta: 0.02, se: 0.01, P: 0.04) (Supplementary Table 17).

## Discussion

Using a Bayesian framework and sensitivity analyses with ordinary least squares and mixed linear models, we robustly identified 13 independent genetic and 3 epigenetic correlates of circulating inflammatory protein levels. This is the first study to have integrated genetic and epigenetic data together using multiple methods to identify molecular correlates of, and estimate the contribution of these molecular factors towards inter-individual variability in, the circulating proteome. Using integrative causal frameworks, we identified mechanisms through which genetic variation may perturb plasma protein levels. Additionally, we demonstrated causal relationships between prioritised circulating inflammatory proteins and blood pressure as well as inflammatory bowel diseases.

By relying on careful corrections for multiple testing and cross-validation across methods, our stringent approach was well-equipped to identify true biological signal as opposed to false positives. This is particularly pertinent in relation to *trans* associations which show poor replication and often have smaller effect sizes than *cis* associations (32). We identified one *trans* pQTL (rs12075) associated with levels of the chemokine MCP4 (encoded for by *CCL13* gene on chromosome 17). This SNP represents a nonsynonymous polymorphism (Asp42Gly) annotated to the *Duffy antigen/chemokine receptor* (*DARC*) gene on chromosome 1. Previously, this SNP has been associated with lower MCP1 levels and evidence shows that the base-change results in altered chemokine-receptor binding (33-35). Additionally, this polymorphism has been shown to explain approximately 20% of variation in MCP1 levels, similar to our estimate of 18.66% in MCP4 levels (34). The Duffy antigen receptor is expressed on erythrocytes and acts as a reservoir for circulating chemokines resulting in reduced distribution of chemokines to extravascular tissue and dampened pro-inflammatory effects (36). Our findings suggest that this polymorphism may also lead to reduced MCP4 levels, possibly through augmented chemokine-receptor interaction.

In the EWAS analyses, the well-documented smoking-associated probe cg05575921 (*AHRR* locus) was associated with CCL11 levels. This association was attenuated after adjustment for smoking. Higher levels of CCL11 have been associated with tobacco smoking and cannabis use (37-39). We also found altered methylation at the *NLRC5* locus (*NOD-like receptor* family CARD domain containing 5) is associated with circulating CXCL9 levels. NRLC5 acts as a potent regulator of the inflammasome (40, 41). Zaghlool *et al.* showed that altered methylation at the *NLRC5* locus associates with several inflammatory markers, including CXCL10 and CXCL11, with pathway analyses linking it to disease states in which NLRC5 dysfunction is implicated such as cancer and cardiovascular disease (9).

Using our database of genotype-protein associations, we tested for causal relationships between inflammatory protein biomarkers and human phenotypes. However, in each case, only one variant was available to test for such associations which does not allow for the testing of pleiotropic effects. CD6 was associated with clinically diagnosed IBD. Expression of the CD6 receptor and its ligand, ALCAM, are overexpressed in the intestinal mucosa of IBD patients where it may promote CD4^+^ T cell proliferation and differentiation into pro-inflammatory Th1/Th17 cells (42). FGF-5 levels were associated with automated readings of systolic and diastolic pressure; previously, FGF-5 levels have been significantly correlated with blood pressure (43). Variation in the *IL12B* gene has been linked strongly to the pathogenesis of Crohn’s disease and an antibody targeted towards the p40 subunit of IL12 demonstrated efficacy in the treatment of moderate-to-severe Crohn’s disease (44). In our study, we showed that circulating IL12B levels may be causally linked to this disease. Lastly, IL18R1 levels may also be causally associated with IBD. A number of studies have demonstrated that increased IL18 signalling confers detrimental effects in the context of gastrointestinal inflammatory processes (45).

Our study has a number of caveats. First, proteins with high sequence homology and structural similarities to a targeted protein of interest may be inappropriately captured by assay probes resulting in quantification errors. Second, there was a strong correlation structure among the inflammatory protein panel. However, given that inflammatory proteins are often co-expressed and synergistic, overlapping loci may reveal biologically important foci or nodes of co-regulation (46). Third, our Scottish cohort contains individuals from a homogenous genetic background limiting the generalisability of our findings to individuals of other ethnic backgrounds. Fourth, ageing is closely linked to chronic low-grade inflammation. Therefore, the distributions of, and correlation structure among, inflammatory protein biomarkers may differ in our cohort of healthy older ageing when compared to other age ranges and the general older adult population. Fifth, the sample size within our study resulted in large confidence and credible intervals in the reported estimates for heritabilities in inflammatory protein levels.

To conclude, our integrative and cross-method approach has identified genetic and epigenetic loci associated with inflammatory protein biomarker levels. Furthermore, we have provided novel estimates for the contribution of common genetic and epigenetic variation towards differences in circulating inflammatory biomarker levels. Together, our data may have important implications for informing the molecular regulation of the human proteome. Furthermore, our data provides a platform upon which other researchers may investigate relationships between inflammatory biomarkers and disease, and a resource to further inform biological insights into immunological and inflammatory processes.

## Methods

### The Lothian Birth Cohort 1936

The Lothian Birth Cohort 1936 (LBC1936) study is a longitudinal study of ageing. Cohort members were all born in 1936 and most took part in the Scottish Mental Survey 1947 at age 11 years. Participants who were living mostly within the Edinburgh area were re-contacted approximately 60 years later (n = 1,091, recruited at mean age 70 years). Recruitment and testing of the LBC1936 cohort have been described previously (47, 48).

### Ethical approval

Ethical permission for the LBC1936 was obtained from the Multi-Centre Research Ethics Committee for Scotland (MREC/01/0/56) and the Lothian Research Ethics Committee (LREC/2003/2/29). Written informed consent was obtained from all participants.

### Protein measurements in the Lothian Birth Cohort 1936

Plasma was extracted from 1,047 blood samples and collected in lithium heparin tubes at mean age 69.8 ± 0.8 years. Following quality control, 1,017 samples remained. Plasma samples were analysed using a 92-plex proximity extension assay (Olink^®^ Bioscience, Uppsala Sweden). One protein from the panel, BDNF, failed quality control and was removed from the study. For a further 21 proteins, over 40% of samples fell below the lowest limit of detection. These proteins were removed from analyses leaving a final set of 70 proteins. The proteins assayed comprise the Olink^®^ inflammatory biomarker panel. Briefly, 1 µL of sample was incubated in the presence of proximity antibody pairs linked to DNA reporter molecules. Upon appropriate antigen-antibody recognition, the DNA tails form an amplicon by proximity extension which is quantified by real-time PCR. Data pre-processing was performed by Olink^®^ using NPX Manager software. Protein levels were transformed by rank-based inverse normalisation and regressed onto age, sex, four genetic principal components of ancestry and array plate. Standardised residuals from these regression models were brought forward for all genetic-protein and epigenetic-protein analyses. Pre-adjusted protein level distributions are presented in Supplementary File 2. Associations between pre-adjusted protein levels and biological as well as technical covariates are detailed in Supplementary Table 18.

### Genome-wide association studies

LBC1936 DNA samples were genotyped at the Edinburgh Clinical Research Facility using the Illumina 610-Quadv1 array (n = 1005; mean age: 69.6 ± 0.8 years; San Diego). Quality control procedures for genetic data are detailed in Supplementary Methods.

BayesR+ is a software implemented in C++ for performing Bayesian penalised regression on complex traits (20). The joint and conditional effects of typed SNPs (n = 542,095 variants) on transformed protein levels were examined. The prior distribution is specified as a mixture of Gaussian distributions, corresponding to effect sizes of different magnitude, and a discrete spike at zero which enables the omission of probes and markers with negligible effect on the phenotype. Informed by linear models and data from our previous pQTL study (49), mixture variances for genetic data were set to 0.01 and 0.1. Input data were scaled to mean zero and unit variance, and adjusted for age and sex. To obtain estimates of effect sizes, Gibbs sampling was used to sample over the posterior distribution conditional on the input data. The Gibbs algorithm consisted of 10,000 samples and 5,000 samples of burn-in after which a thinning of 5 samples was utilised to reduce autocorrelation. Genetic markers which exhibited a posterior inclusion probability of ≥95% were deemed to be significant.

Details for the OLS model approach are outlined in Supplementary Methods. In the linear method, markers which surpassed a Bonferroni-corrected conditional significance threshold of 7.14 × 10^−10^ (= genome-wide significance: 5.0 × 10^−8^/70 phenotypes) were considered. The genome-wide significance level of 5.0 × 10^−8^ was selected as per convention in GWAS studies.

### Epigenome-wide association studies

DNA from whole blood was assessed using the Infinium 450 K methylation array at the Edinburgh Clinical Research Facility (n = 876; mean age: 69.8 ± 0.8 years). Quality control procedures for methylation data are detailed in Supplementary Methods.

Using BayesR+, prior mixture variances for methylation data (n = 459,309 CpG sites) were set to 0.001, 0.01 and 0.1. Age, sex and Houseman-estimated white blood cell proportions (50) were incorporated as fixed effect covariates. The same settings as in the genetic analyses were applied. Methylation probes which had a posterior inclusion probability of ≥95% were deemed to be significant.

Details for the OLS and mixed linear model approaches are outlined in Supplementary Methods. For these methods, probes which surpassed a Bonferroni-corrected significance threshold of 5.14 × 10^−10^ (= genome-wide significance: 3.6 × 10^−8^/70 phenotypes) were deemed to be significant. The genome-wide significance level of 3.6 × 10^−8^ was selected as per the recommendations of Safarri *et al.* (51).

### Functional Annotation of Genetic and Epigenetic Loci

Genetic markers that were independently associated with protein levels were functionally annotated using ANNOVAR (52) and Ensembl genes (build 85) in FUMA (**FU**nctional **M**apping and **A**nnotation) (53). Epigenetic probes associated with protein levels were annotated using the *IlluminaHumanMethylation450kanno.ilmn12.hg19* package (54).

### Identification of overlap between *cis* pQTLs and *cis* eQTLs

To determine whether pQTL variants may affect protein levels through modulation of gene expression, we cross-referenced *cis* pQTLs with publicly available blood-based (and FDR-corrected significant) *cis* expression QTL (eQTL) data from the eQTLGen consortium (55).

### Colocalisation

To test whether a sole causal variant might underlie both an eQTL and pQTL association, we performed Bayesian tests of colocalisation using the *coloc* package in R (26). For each protein of interest, a 200 kb region (upstream and downstream – recommended default setting) surrounding the appropriate pQTL was extracted from our GWAS summary statistics (56). For each respective protein, the same region was also extracted from eQTLGen summary statistics. Default priors were applied. Summary statistics from all SNPs within these regions were used to determine the posterior probability for five distinct hypotheses: a single causal variant for both traits, no causal variant for either trait, a causal variant for one of the traits (encompassing two hypotheses), or distinct causal variants for the two traits. Posterior probabilities (PP) ≥ 0.95 provided strong evidence in favour of a given hypothesis.

### Pathway Enrichment and Tissue Specificity Analyses

Using methylation data, pathway enrichment was assessed among KEGG pathways and Gene Ontology (GO) terms through hypergeometric tests using the *phyper* function in R. All gene symbols from the 450 K array annotation (null set of sites) were converted to Entrez IDs using *biomaRt* (57, 58). GO terms and their corresponding gene sets were retrieved from the Molecular Signatures Database (MSigDB)-C5 (59). KEGG pathways were downloaded from the KEGG REST server (60). Tissue specificity analyses were performed using the GENE2FUNC function in FUMA. Differentially expressed gene sets with Bonferroni-corrected P values < 0.05 and an absolute log-fold change of ≥0.58 (default settings) were considered to be enriched in a given tissue type (GTEx v7).

### Mendelian Randomisation

Two-sample Mendelian Randomisation was used to test for putatively causal relationships between (i) the 4 proteins whose pQTLs were previously shown to be associated with human traits, as identified through GWAS Catalog, and the respective traits (29, 61) (http://www.nealelab.is/uk-biobank/), (ii) the 13 proteins which harboured significant pQTLs and Alzheimer’s disease risk (62), (iii) gene expression and inflammatory protein levels and (iv) DNA methylation and inflammatory protein levels. Pruned variants (LD *r*^2^ < 0.1) were used as instrumental variables (IV) in MR analyses. In tests where only one independent SNP remained after LD pruning, causal effect estimates were assessed using the Wald ratio test i.e. a ratio of effect per risk allele on trait to effect per risk allele on protein levels. In tests where multiple independent variants were identified, and if no evidence of directional pleiotropy was present (non-significant MR-Egger intercept), multi-SNP MR was carried out using inverse variance-weighted estimates. Analyses were conducted using MRbase (63). Further details are provided in Supplementary Methods.

## Acknowledgements

The authors thank LBC1936 study participants and research team members who have contributed, and continue to contribute, to ongoing LBC1936 studies. The LBC1936 is supported by Age UK (Disconnected Mind program, which supports S.E.H), the Medical Research Council (MR/M01311/1), and the University of Edinburgh. Genotyping was supported by the Biotechnology and Biological Sciences Research Council (BB/F019394/1). Methylation typing was supported by Centre for Cognitive Ageing and Cognitive Epidemiology (Pilot Fund award), Age UK, The Wellcome Trust Institutional Strategic Support Fund, The University of Edinburgh, and The University of Queensland. Proteomic analyses were supported for by the LBC1936 Age UK grant. This work was conducted in the Centre for Cognitive Ageing and Cognitive Epidemiology, which was supported by the Medical Research Council and Biotechnology and Biological Sciences Research Council (MR/K026992/1), and which supported I.J.D. We acknowledge NIH Grants R01AG054628 and R01AG05462802S1 for supporting this research and Grant P2CHD042849 for supporting the Population Research Center at the University of Texas. R.F.H. and A.J.S. are supported by funding from the Wellcome Trust 4-year PhD in Translational Neuroscience–training the next generation of basic neuroscientists to embrace clinical research [R.F.H: 108890/Z/15/Z; A.J.S: 203771/Z/16/Z]. D.L.Mc.C. and R.E.M. are supported by Alzheimer’s Research UK major project grant ARUK-PG2017B-10. This research was supported by Australian National Health and Medical Research Council (grants 1010374, 1046880 and 1113400) and by the Australian Research Council (DP160102400). P.M.V., N.R.W. and A.F.M. are supported by the NHMRC Fellowship Scheme (1078037, 1078901 and 1083656). P.M.V was also funded by the Australian Research Council (DP160102400 and FL180100072)..

