## Supplementary Note 1 for "Integrative omics approach to identify the molecular architecture of inflammatory protein levels in healthy older adults"

**Conditional and joint analysis from ordinary least squares GWAS on protein levels**

In an ordinary least squares (OLS) regression model, 1,531 SNPs were associated with the levels of 19/70 proteins at a Bonferroni-corrected threshold ($P < 7.14 \times 10^{-10}$; Supplementary Table 2). Manhattan and Q-Q plots for these 19 proteins are presented in Supplementary Files 3 and 4, respectively. Estimates for inflation factors across each of the 70 genome-wide association studies are listed in Supplementary Table 3. Conditional and joint analysis (GCTA-COJO) was performed to identify which of these hits were independent of one another, resulting in the identification of 27 conditionally significant pQTLs associated with the circulating levels of 17 proteins (Supplementary Table 4). Of note, whereas Bonferroni-corrected genome-wide significant SNPs were identified for an additional two proteins (CCL23 and MMP-10), the conditional P value for these pQTLs from GCTA-COJO did not fall below the Bonferroni-corrected threshold of $P < 7.14 \times 10^{-10}$.

**Sherlock: identifying genes whose expression associates with inflammatory biomarkers**

The Bayesian algorithm termed Sherlock uses *cis* and *trans* eQTLs to assign gene-based scores from GWAS data to identify genes whose expression associates with a trait of interest (here, protein levels). Putative gene expression-protein associations for all 13 proteins are outlined in Supplementary Table 10. From this gene-based colocalisation approach, only gene expression of *ADA*, *CXCL5* and *IL18R1* were associated with levels of their respective protein products. Expression of *MIF4GD* and *GRB2* were both associated with CD6 and TNFB levels (*r* between levels: 0.45). Expression of *TNRC6A* was associated with CD6 and ADA levels (*r:* 0.55). *CXCL4L1* expression was associated with CXCL5 and CXCL6 levels (*r*: 0.44).

The disparity between the employed colocalisation methods, coloc and Sherlock, may reflect differences in technical and biological variability. The former method considers *cis* regions in a continuous chromosomal region as defined by the pQTL whereas the latter takes into account all *cis* and *trans* eQTLs across the genome passing a soft significance threshold. Both transcript datasets and our pQTL estimates were generated in different samples which may have resulted in different molecular abundances and differences in the overlapping of transcript and protein distributions. The small sample sizes used to generate the datasets may have limited power to detect further *cis* gene expression-protein patterns.

**Replication of previous pQTLs and protein association-CpG sites**

Summary statistics from four major pQTL studies were extracted to determine whether the 13 concordant pQTLs identified in this study replicated those of previous findings (1-4). Of the 13 proteins that harboured a pQTL, 10 were available for look-up. Four (40.0%) pQTLs were replicated in our study, three of which had available beta statistics. There was good agreement between our beta values and those reported in the literature (rs6851997 for CXCL6: 0.37 vs. 0.52; rs917997 for IL18R1: 0.68 vs. 0.45 and rs3138036 for MCP2: -0.62 vs. -0.47, respectively). For the 10 proteins which were available for look-up analyses, we aimed to extract beta coefficients for all significant pQTLs associated with their levels in the literature. Of note, beta coefficients were reported for 8/10 proteins. Many of these pQTLs were non-significant in our study, though we observed a moderate correlation between the effect sizes (r: 0.70, 95% CI: [0.46, 0.84]; Supplementary Figure 8).

Of the 3 proteins with significant CpG sites (n = 3) identified by multiple methods, 1 was available for look-up from the EWAS on inflammatory proteins performed by Ahsan *et al*. (5). This CpG-protein association was replicated in our study (cg07839457 (*NLRC5*) for CXCL9 levels; $beta_{Ahsan}$: -2.91 vs. $beta_{LBC}$: -3.26).

**Evaluating causal associations between blood inflammatory proteins and Alzheimer's risk**

Using two-sample Mendelian randomisation, we tested whether the 13 inflammatory proteins with significant genetic correlates in our study were causally associated with Alzheimer's disease risk (Supplementary Table 17). One protein, IL18R1, showed a nominally significant association with AD risk (no. of instruments: 1, beta: 0.02, se: 0.01, P: 0.04; Wald ratio test). Conversely, AD risk was not associated with IL18R1 levels (no. of instruments: 22, beta: 0.03, se: 0.21, P: 0.85; inverse variance-weighted method). The intercept from MR Egger regression was −0.07 (P = 0.11) which does not provide evidence for directional pleiotropy.
