## Supplementary Methods for "Integrative omics approach to identify the molecular architecture of inflammatory protein levels in healthy older adults"

**Quality control of genetic data in Lothian Birth Cohort 1936**

SNPs were imputed to the 1000 G reference panel (phase 1, version 3). Individuals were excluded on the basis of sex discrepancies, relatedness, SNP call rate of less than 0.95 and evidence of non-Caucasian descent. SNPs with a call rate of greater than 0.98, minor allele frequency in excess of 0.01 and Hardy-Weinberg equilibrium test with $P \geq 0.001$ were included in analyses.

**Genome-wide association studies**

*Ordinary least squares regression model*

Genome-wide association analyses were conducted on 8,675,776 (typed and imputed) autosomal variants against protein phenotypes which were pre-corrected for age, sex, four genetic principal components of ancestry and Olink® array plate. Ordinary least squares regression was used to assess the effect of each genetic variant on transformed protein levels using mach2qtl (1, 2).

To identify independent genetic associations with Olink® inflammatory levels, we performed approximate genome-wide stepwise conditional analysis through GCTA-COJO (3). We used the 'cojo-slct' option and individual level genotype data. Default settings of the software were applied.

**Quality control of methylation data in Lothian Birth Cohort 1936**

Raw intensity data were background-corrected and normalised using internal controls. Following background correction, manual inspection permitted removal of low quality samples presenting issues relating to bisulphite conversion, staining signal, inadequate hybridisation or nucleotide extension. Quality control analyses were performed to remove probes with low detection rate <95% at $P < 0.01$. Samples with a low call rate (samples with <450,000 probes detected at $p$-values of less than 0.01) were also eliminated. Furthermore, samples were removed if they had a poor match between genotype and SNP control probes, or incorrect DNA methylation-predicted sex.

### Epigenome-wide association studies

*Ordinary least squares regression model*

In the ordinary least squares model strategy (limma; linear models for microarray data), each CpG site (n = 459,309) was regressed on transformed protein levels with adjustments for age, sex, estimated white blood cell proportions ($CD4^+$ T cells, $CD8^+$ T cells, B cells, Natural Killer Cells and granulocytes) and technical covariates (plate, position, array, hybridisation, date). Proportions of white blood cells were estimated from methylation data using the Houseman method (4).

*Mixed linear model*

In contrast to limma, CpG site (n = 459,309) was the independent variable whereas Olink® protein levels were input as dependent variables in all mixed models (performed using **O**mic**S**-data-based **C**omplex trait **A**nalysis: OSCA) (5). The same covariates were adjusted for as in the limma strategy. The MOMENT method was used to test for associations between traits of interest and methylation at individual probes. MOMENT is a mixed linear model-based method that can account for unobserved confounders and the correlation between distal probes which may be introduced by such confounders. The same Bonferroni-corrected threshold as the linear model was applied: $5.14 \times 10^{-10}$ (= genome-wide significance: $3.6 \times 10^{-8}$/70 phenotypes).

### Sherlock

A Bayesian algorithm termed Sherlock (6) was used to detect gene-protein associations by incorporating information from publicly available eQTL data and GWAS summary statistics. This was carried out to infer genes whose differential expression may contribute to alterations in circulating levels of Olink® inflammatory proteins. Sherlock identifies all *cis* and *trans* eQTLs or expression-associated SNPs (eSNPs) for a given gene in a selected data set. The algorithm evaluates the association of each eSNP with the trait of interest (i.e. protein levels) using supplied GWAS data. A score is assigned to each gene based on aligning P-values for the association of the SNP with gene expression and the studied trait. There are three possible scenarios which affect this gene-based score: (i) if the eSNP for the gene is also associated with the trait, a positive score is assigned, (ii) if the eSNP is not associated with the trait, a negative score is assigned and (iii) if the SNP is associated with the trait only (non-eSNP), the score is not affected. The total score of a gene increases in tandem with an increase in the number of SNPs with combined evidence (SNPs that are associated with trait and expression). For each SNP in the alignment, the logarithm of Bayes factor is computed and the sum of constituent SNPs in the gene constitutes the final score for the gene. SNPs that have moderate statistical significance in GWAS and eQTL data sets, that are otherwise missed by traditional GWAS thresholds, are considered. SNPs with stronger associations with the trait contribute more to the final gene-based score than moderately-associated variants. Default settings were applied. As our protein data was collected from whole blood, analyses were restricted to the eQTL GTEx (V7) Whole Blood data set (n = 369) (7). Correction for multiple testing was carried out using the Benjamini-Hochberg procedure at a threshold of $P < 1.0 \times 10^{-5}$ (8).

**Mendelian Randomisation**

(i) Pruned protein QTL variants were used as instrumental variables (IV) to determine the relationship between circulating inflammatory protein biomarkers and their respective phenotypic associations, as identified through GWAS Catalog. Four proteins were shown to have an association with five human traits. Thus, a Bonferroni-corrected significance threshold of 0.01 (0.05/5 tests) was applied.

(ii) For 11/13 proteins, only one SNP remained after linkage disequilibrium (LD) pruning. For 2 proteins (CCL25 and CST5), two independent SNPs were present after pruning. Pruned SNPs were used as IV to test for causal associations between each of the 13 inflammatory proteins and risk of late-onset Alzheimer's disease (9). A Bonferroni-corrected significance threshold of $3.85 \times 10^{-3}$ (0.05/13 tests) was applied. For one protein (IL18R1) which showed a nominally significant association with AD risk, a bidirectional analysis was performed to assess for a putatively causal association in which AD risk affected circulating IL18R1 levels. For this test, 22 independent SNPs remained after LD pruning.

(iii) Expression QTLs obtained from eQTLGen consortium were used as IV to test whether changes in gene expression were causally associated with protein levels (10).

(iv) One protein (IL18R1) harboured both genome- and epigenome-wide significant associations in this study. Therefore, we wished to determine whether methylation affected protein levels and/or whether protein levels affected methylation. We used Phenoscanner to determine whether the pQTL identified for IL18R1 levels (rs917997) has been reported as a methylation QTL for the corresponding *cis* genome-wide significant CpG site identified for IL18R1 levels in our study (cg03938978) (11). The methylation QTL was used as an instrument to test whether altered DNA methylation was causally associated with inflammatory protein levels. Conversely, the IL18R1 pQTL was used as an instrument to assess whether altered IL18R1 levels were causally linked to differential methylation.
