## Supplementary Figures for "Integrative omics approach to identify the molecular architecture of inflammatory protein levels in healthy older adults"

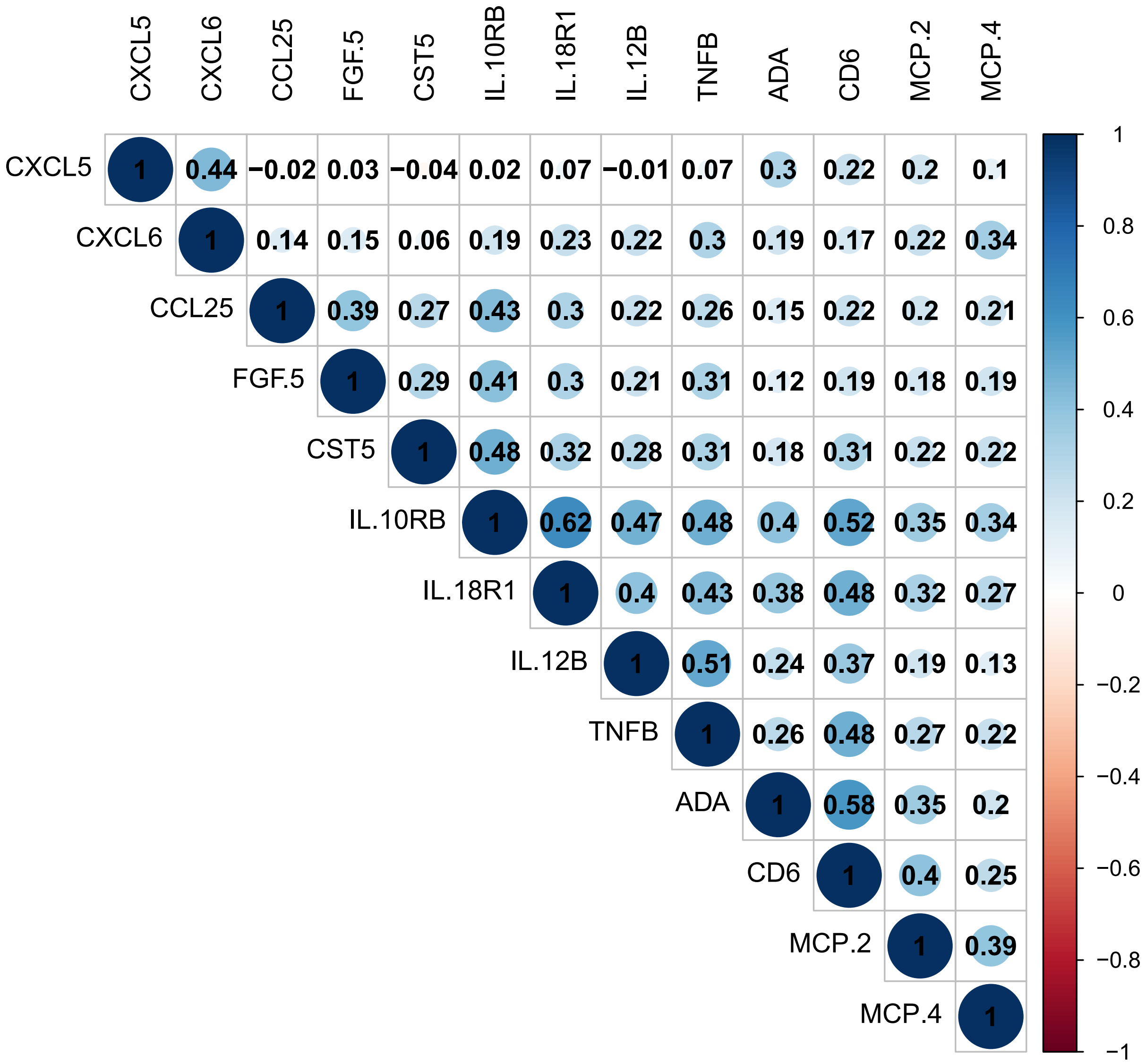


**Supplementary Figure 1.** Correlation between the 13 proteins with significant pQTLs as identified by ordinary least squares and Bayesian penalised regression.

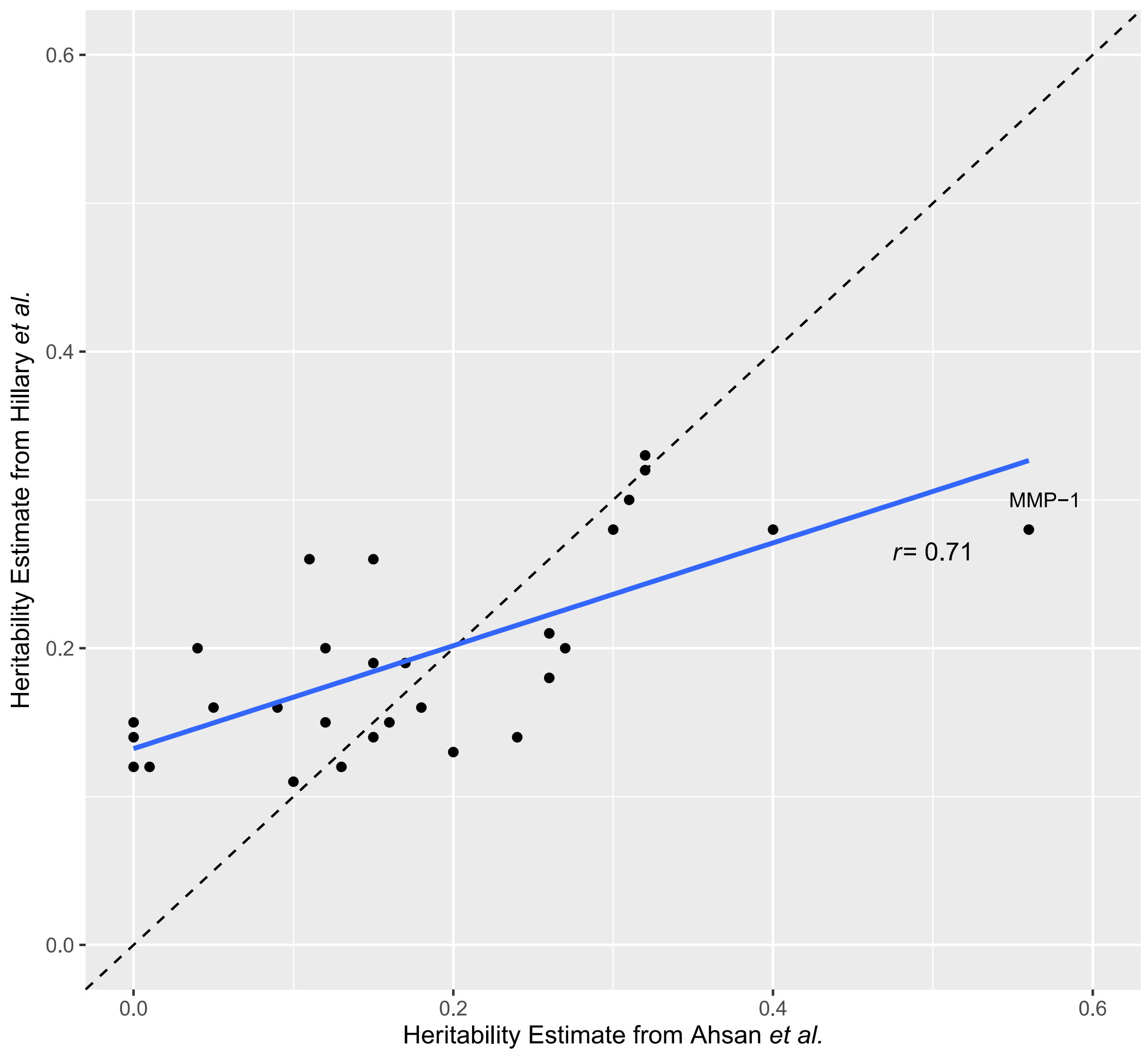


**Supplementary Figure 2.** Correlation between heritability estimates for circulating inflammatory protein biomarkers from present study and that of Ahsan *et al.* The protein with the greatest discordance between studies (MMP-1) is annotated.

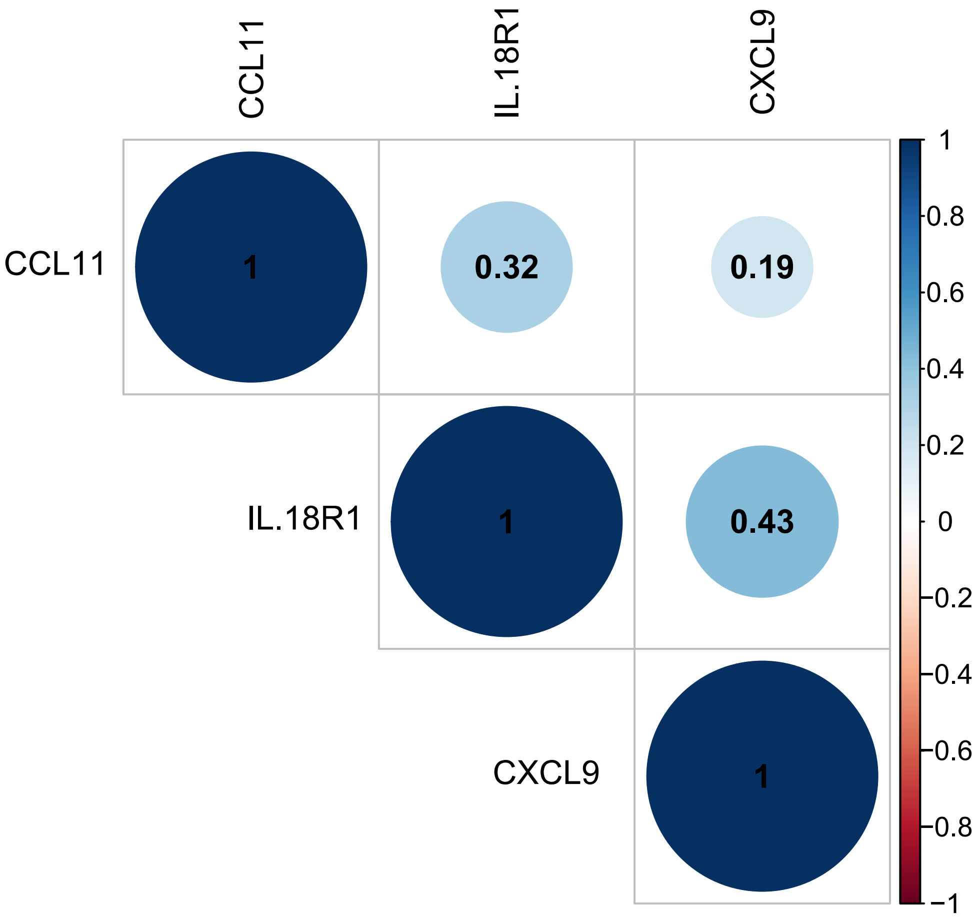


**Supplementary Figure 3.** Correlation between the 3 proteins with significant CpG associations as identified across ordinary least squares model, mixed model and Bayesian penalised regression approaches.

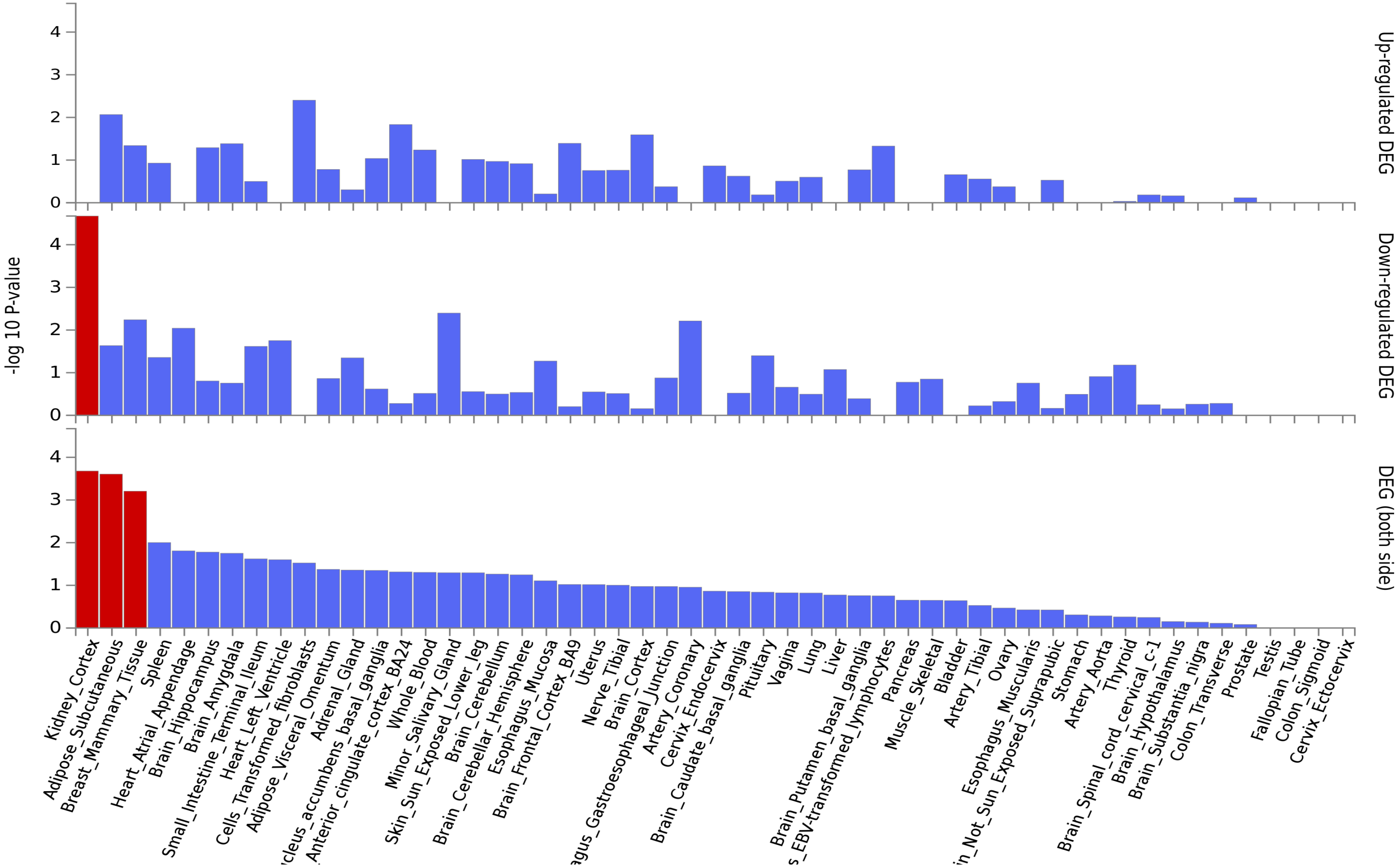


**Supplementary Figure 4.** Tissue-specific expression of genes annotated to CpGs associated with CCL11 levels at $P < 1 \times 10^{-5}$. Differential expression was observed in kidney, adipose and breast tissue.

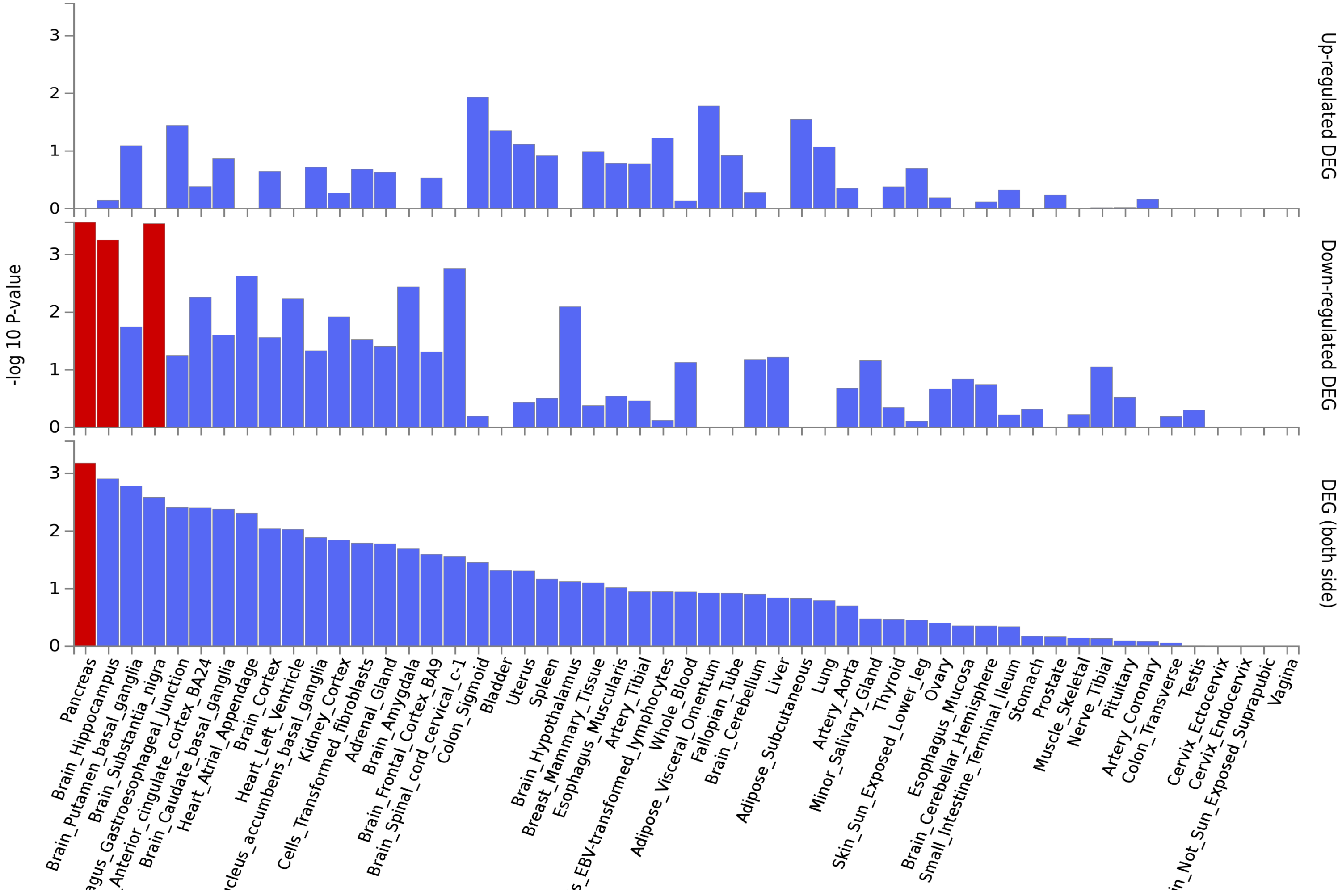


**Supplementary Figure 5.** Tissue-specific expression of genes annotated to CpGs associated with IL18R1 levels at $P < 1 \times 10^{-5}$. Differential expression was observed in pancreatic, hippocampal and substantia nigra tissue.

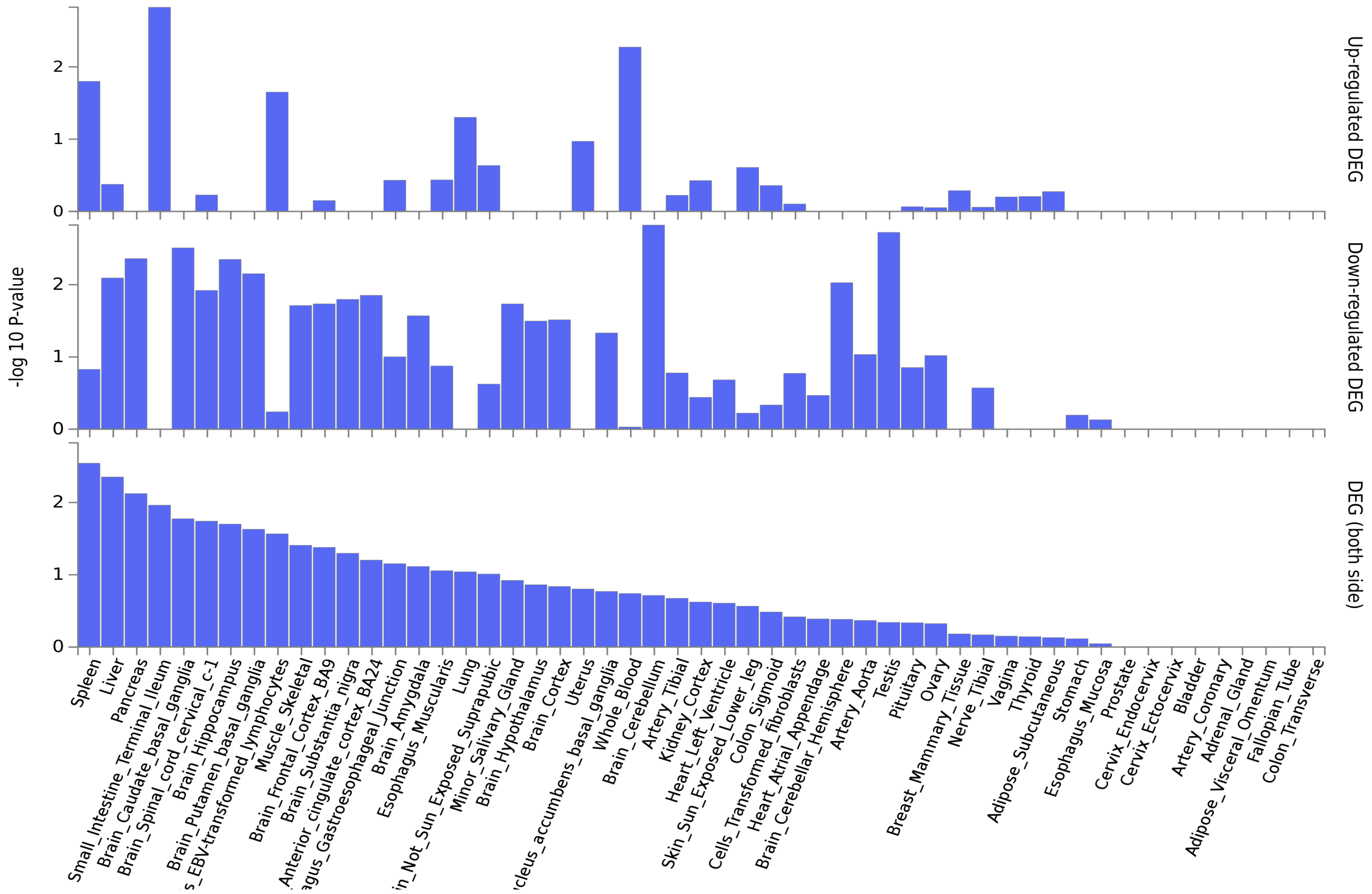


**Supplementary Figure 6.** Tissue-specific expression of genes annotated to CpGs associated with CXCL9 levels at $P < 1 \times 10^{-5}$. No tissue-specific expression was observed.

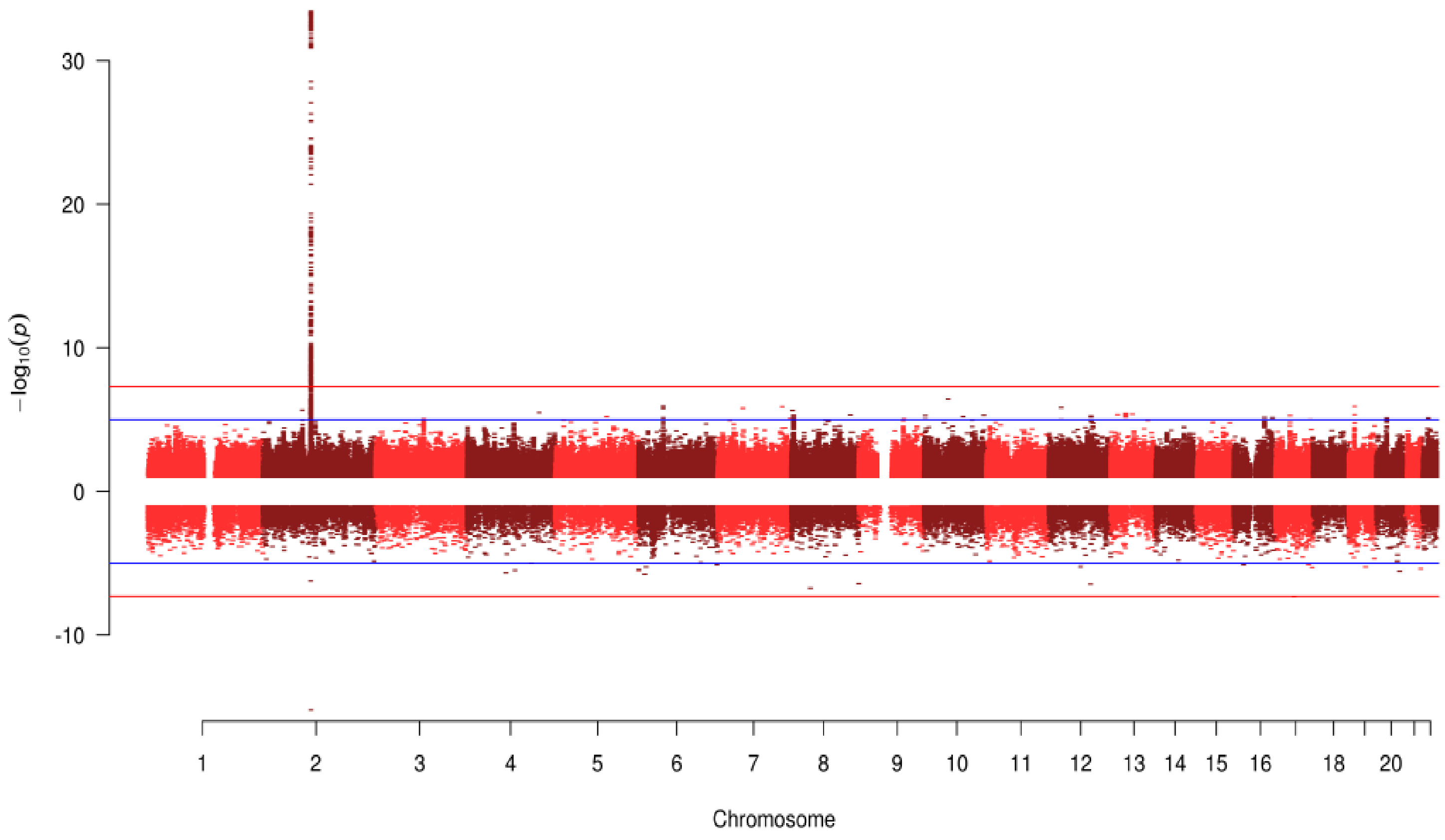


**Supplementary Figure 7.** Miami plot for IL18R1 which exhibited both genome-wide significant SNP and genome-wide significant CpG associations. The top half of the plot (skyline) shows the results from the GWAS on protein levels, whereas the bottom half (waterfront) shows the results from the EWAS. IL18R1 (chromosome 2: 102,311,529-102,398,775).

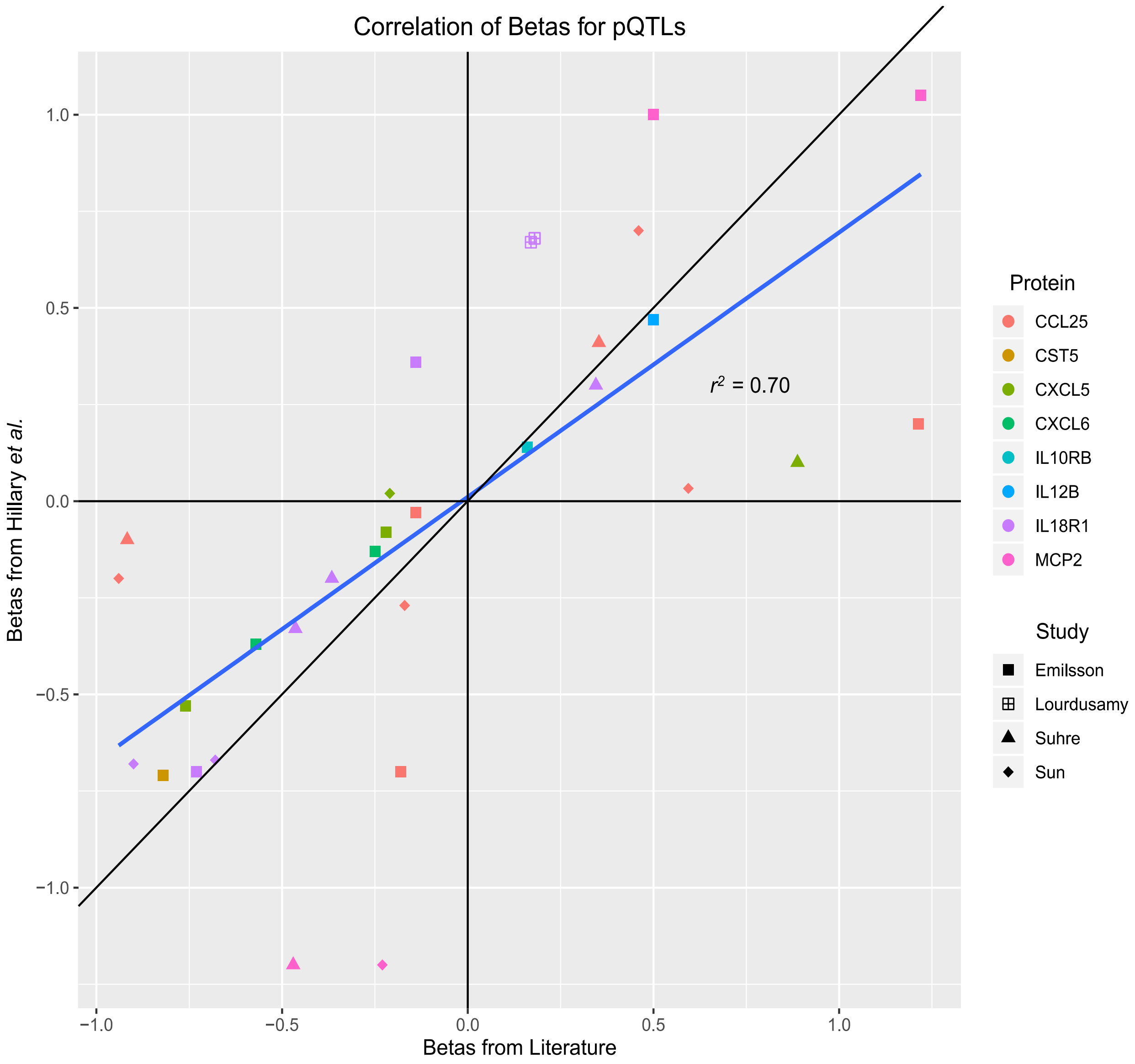


**Supplementary Figure 8.** Correlation of beta values for pQTL associations derived from literature with beta values from the present study.
