## Supplementary figures and images for "Integrative omics approach to identify the molecular architecture of inflammatory protein levels in healthy older adults"

### Supplementary File 1

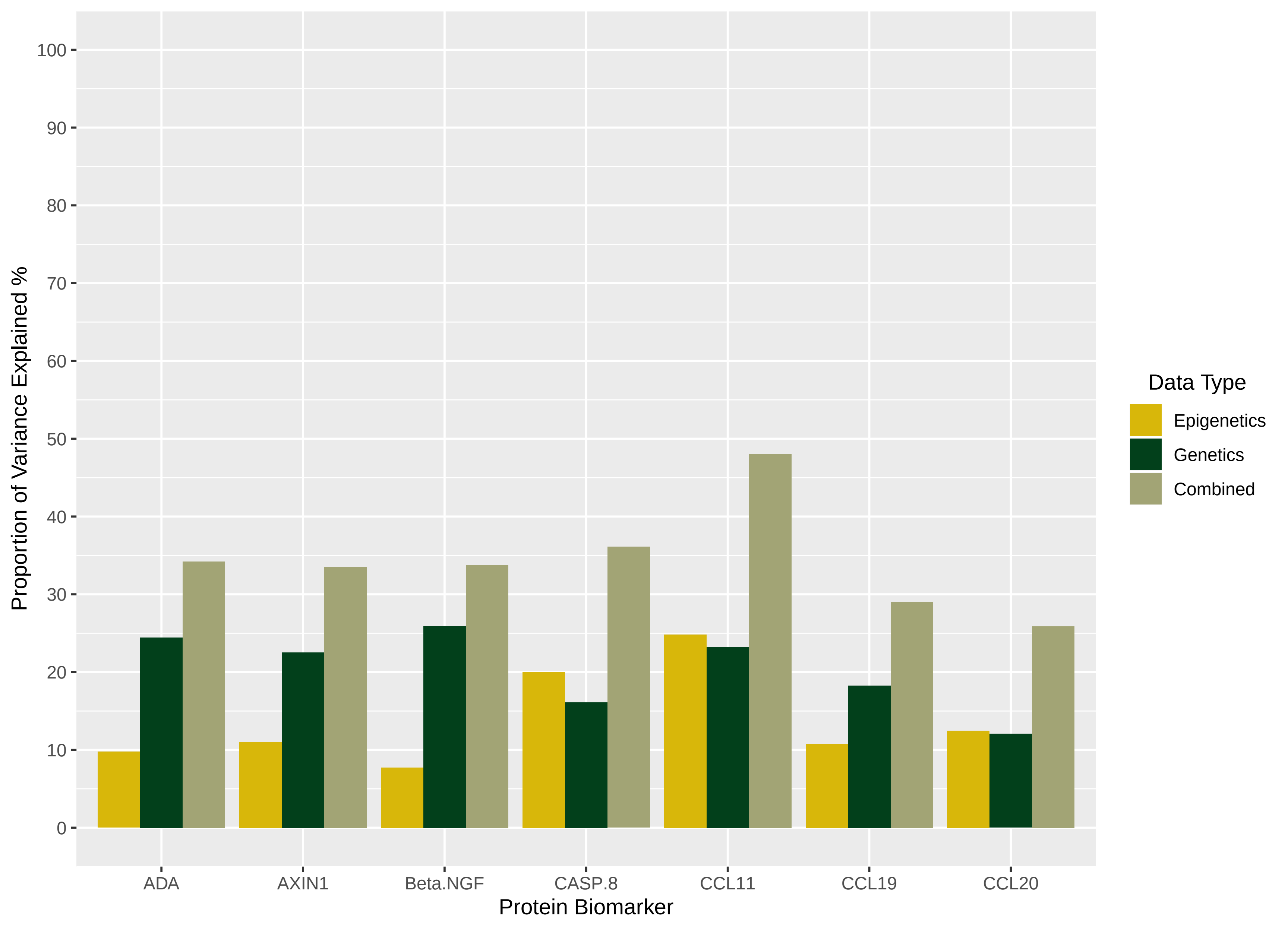

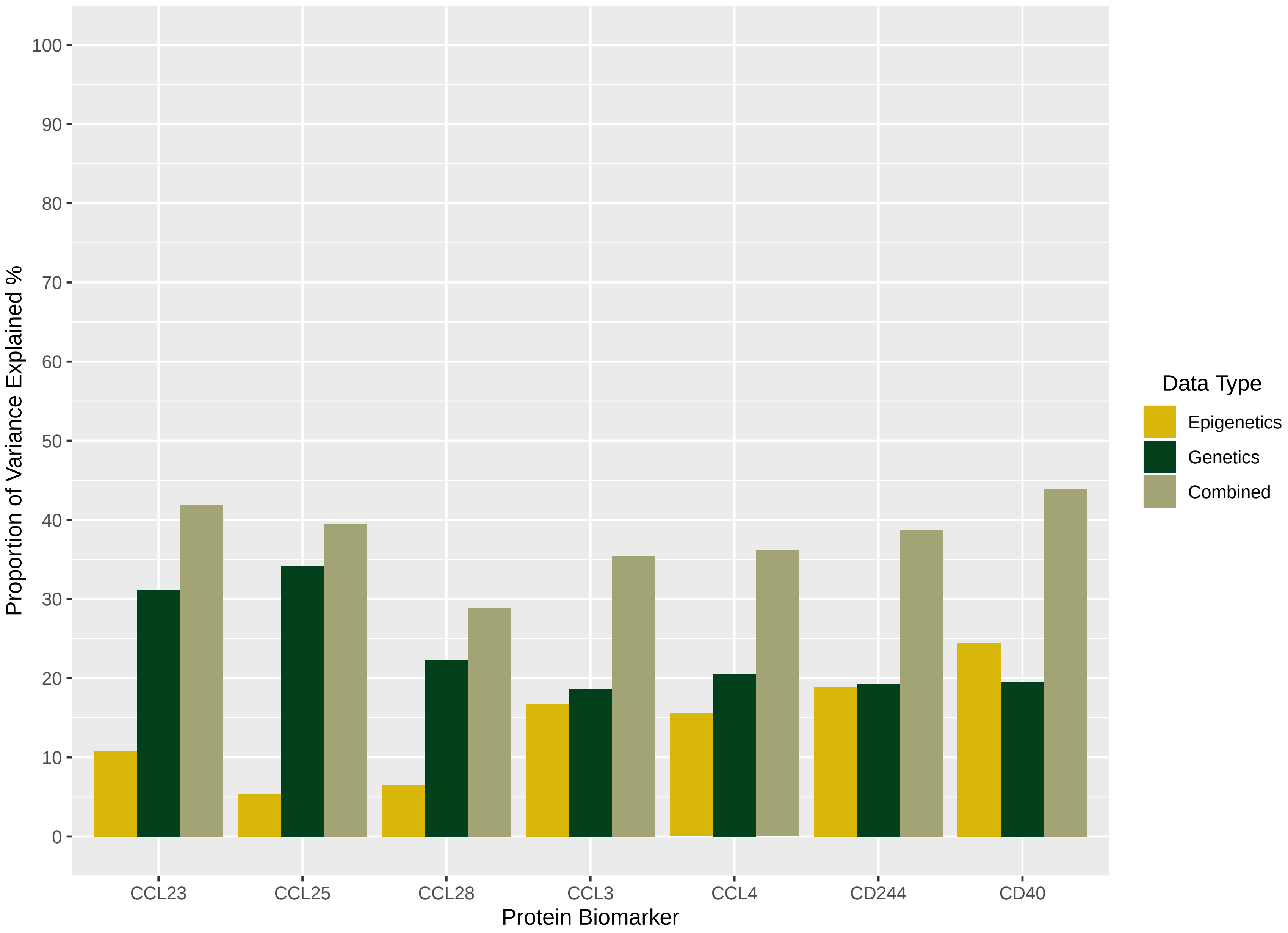

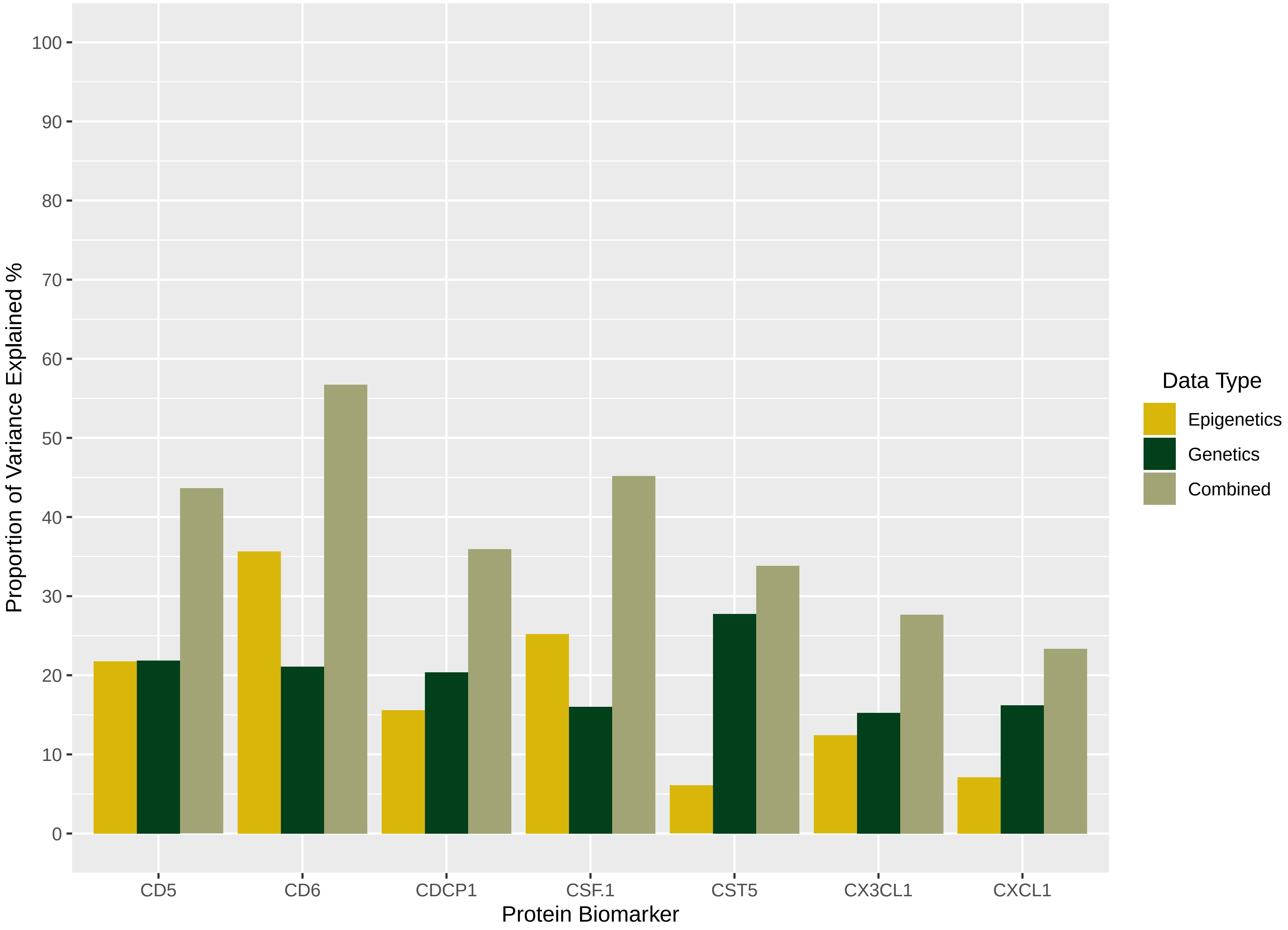

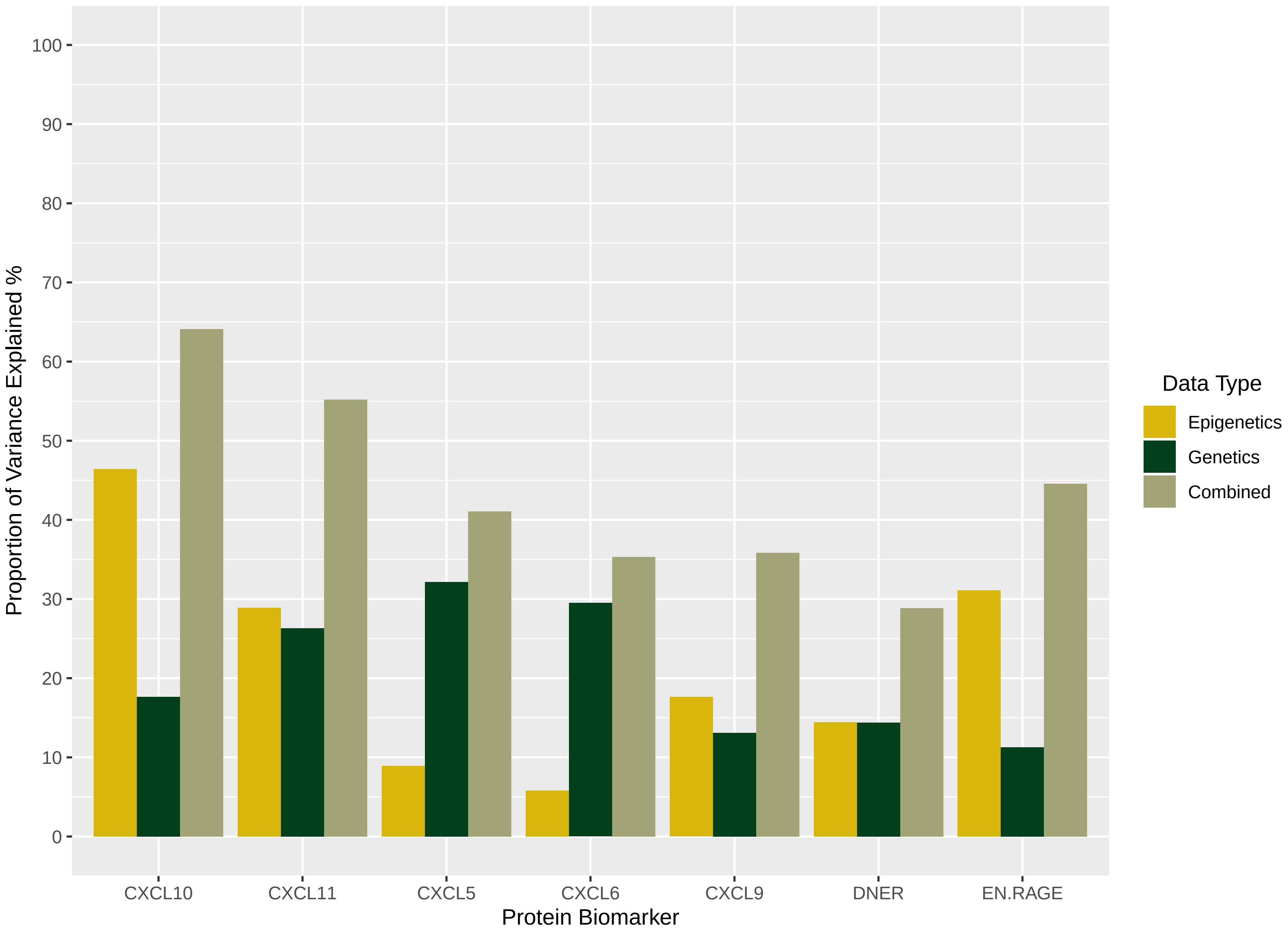

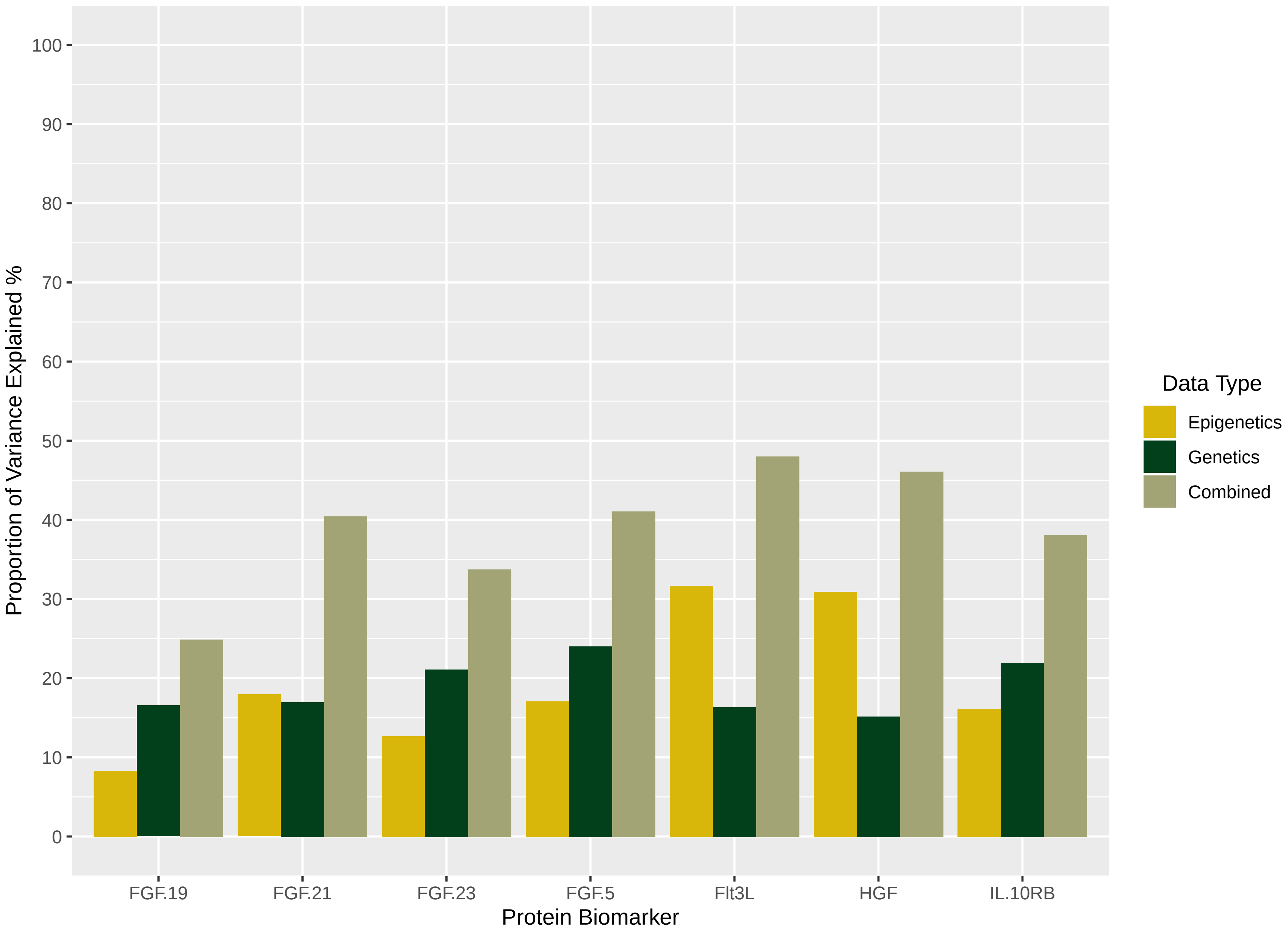

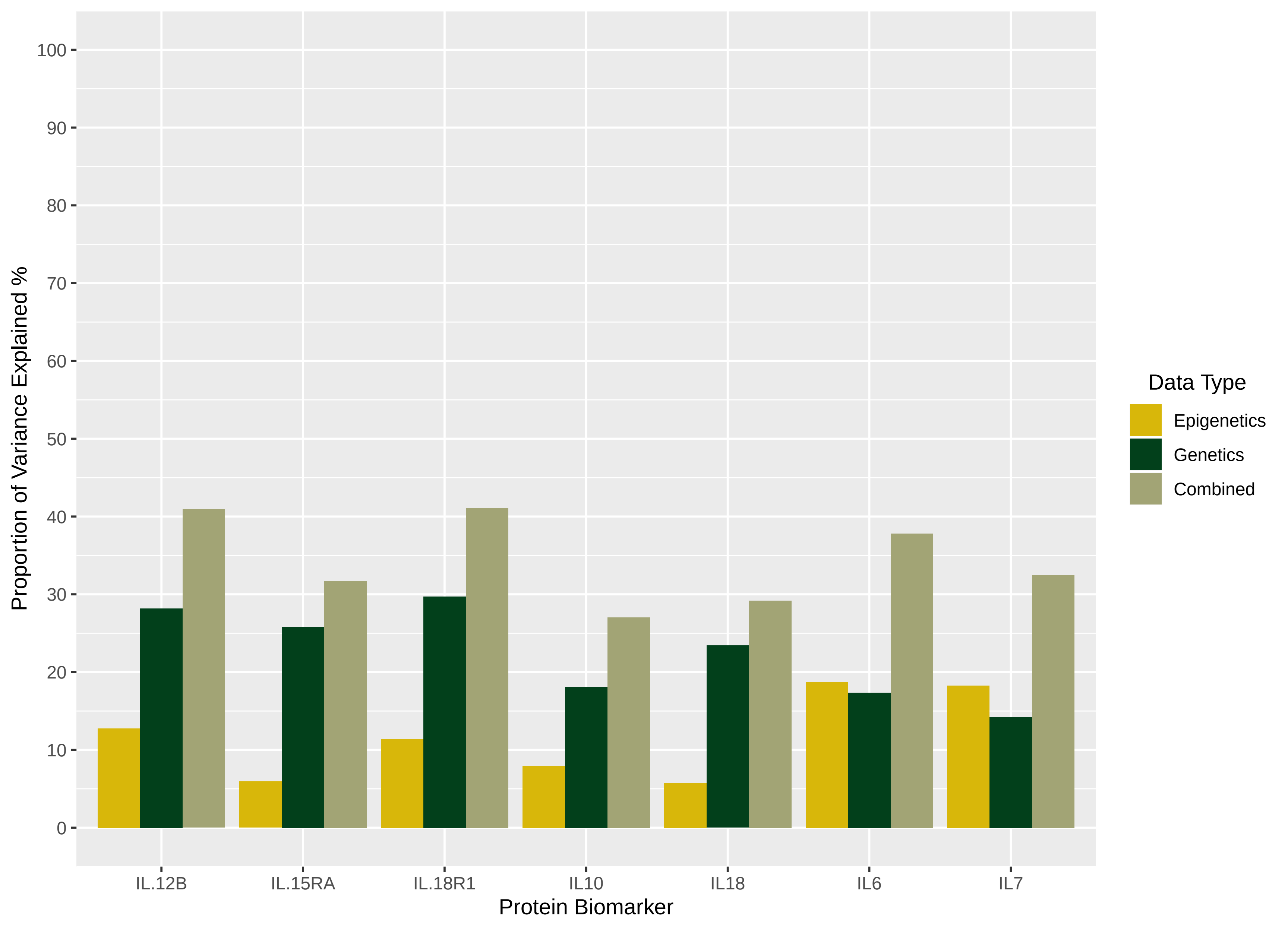

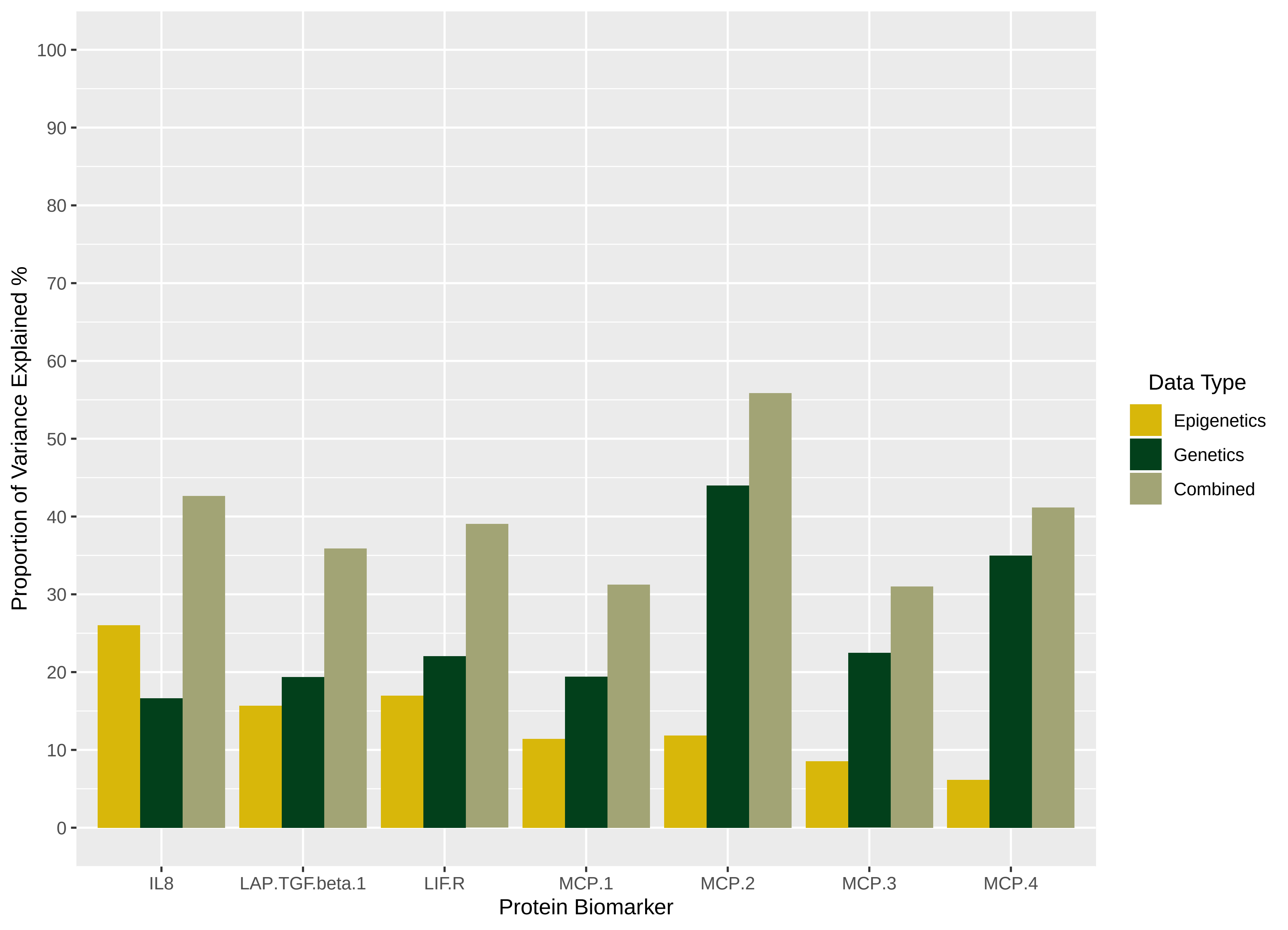

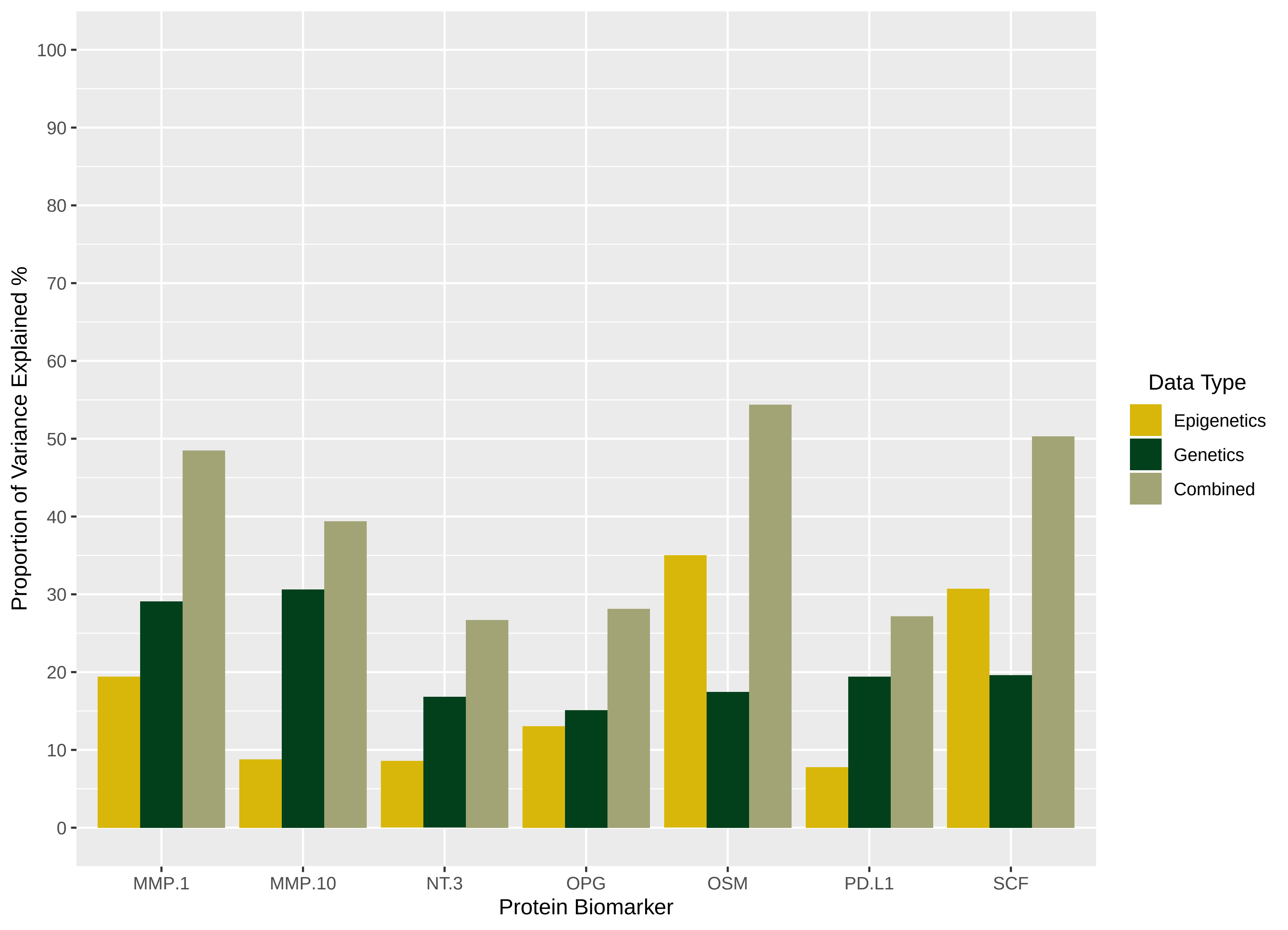

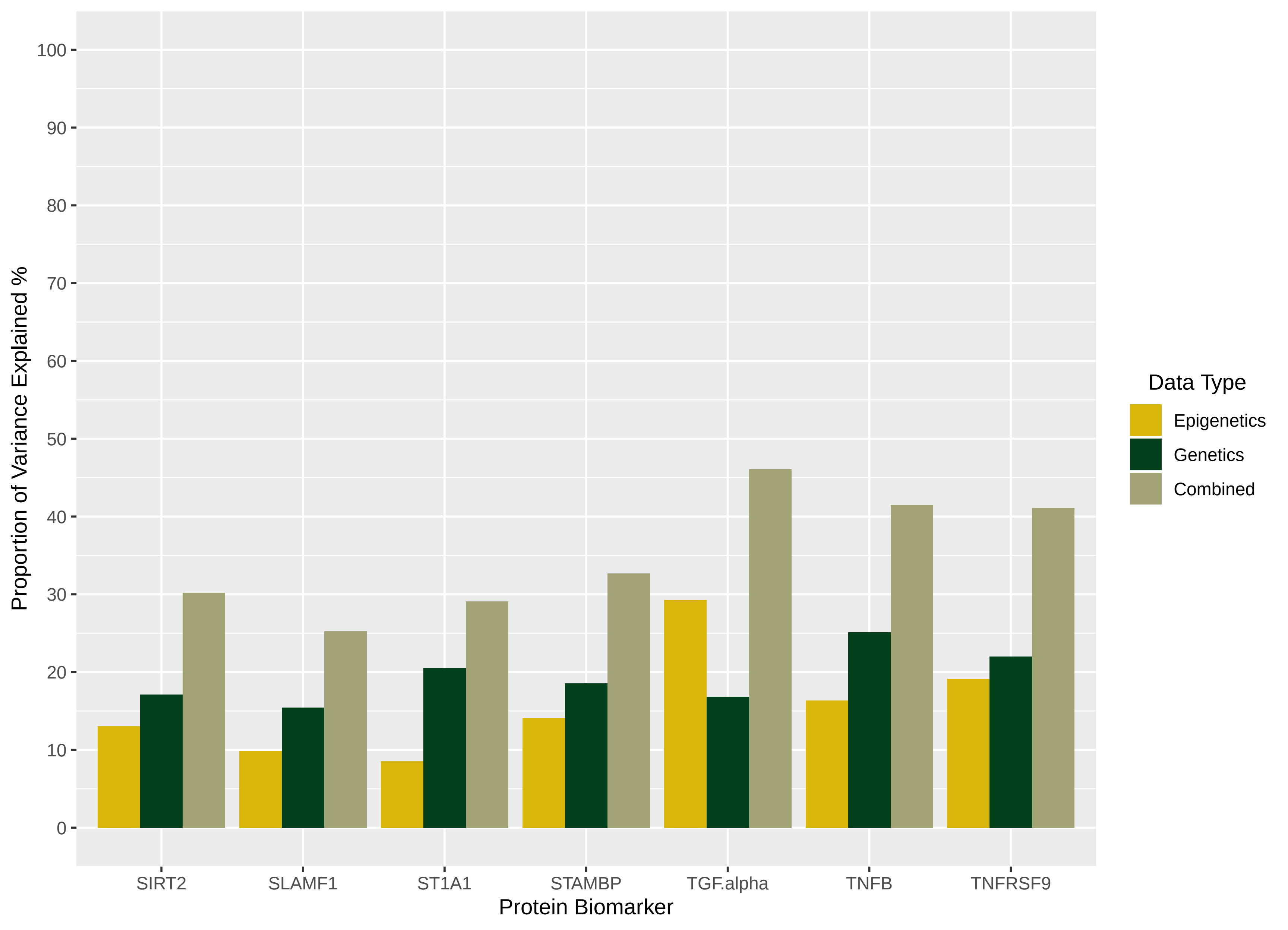

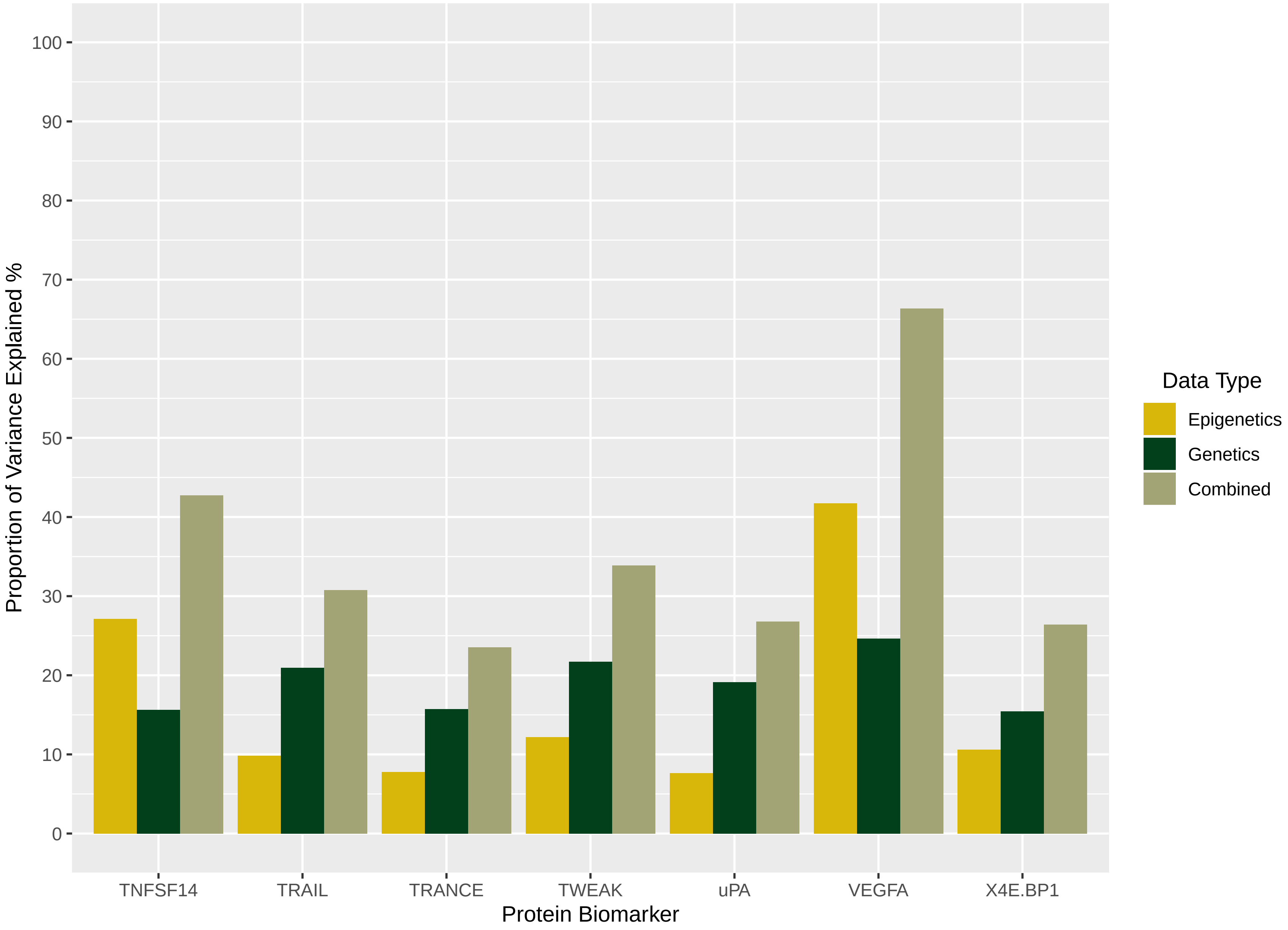
