## Supplementary File 2 for "Integrative omics approach to identify the molecular architecture of inflammatory protein levels in healthy older adults"

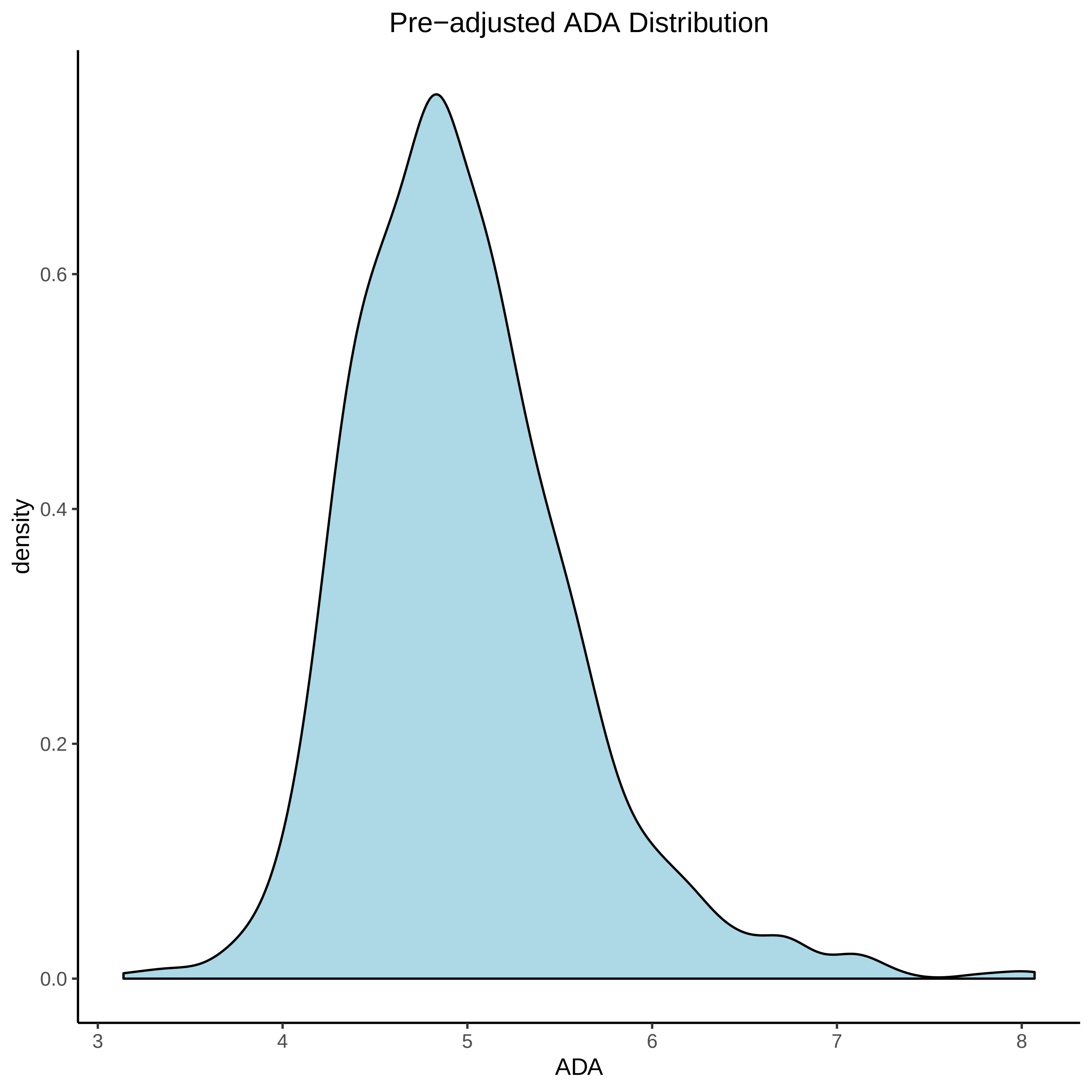

Pre−adjusted ADA Distribution
density
0.0
0.2
0.4
0.6
3
4
5
6
7
8
ADA

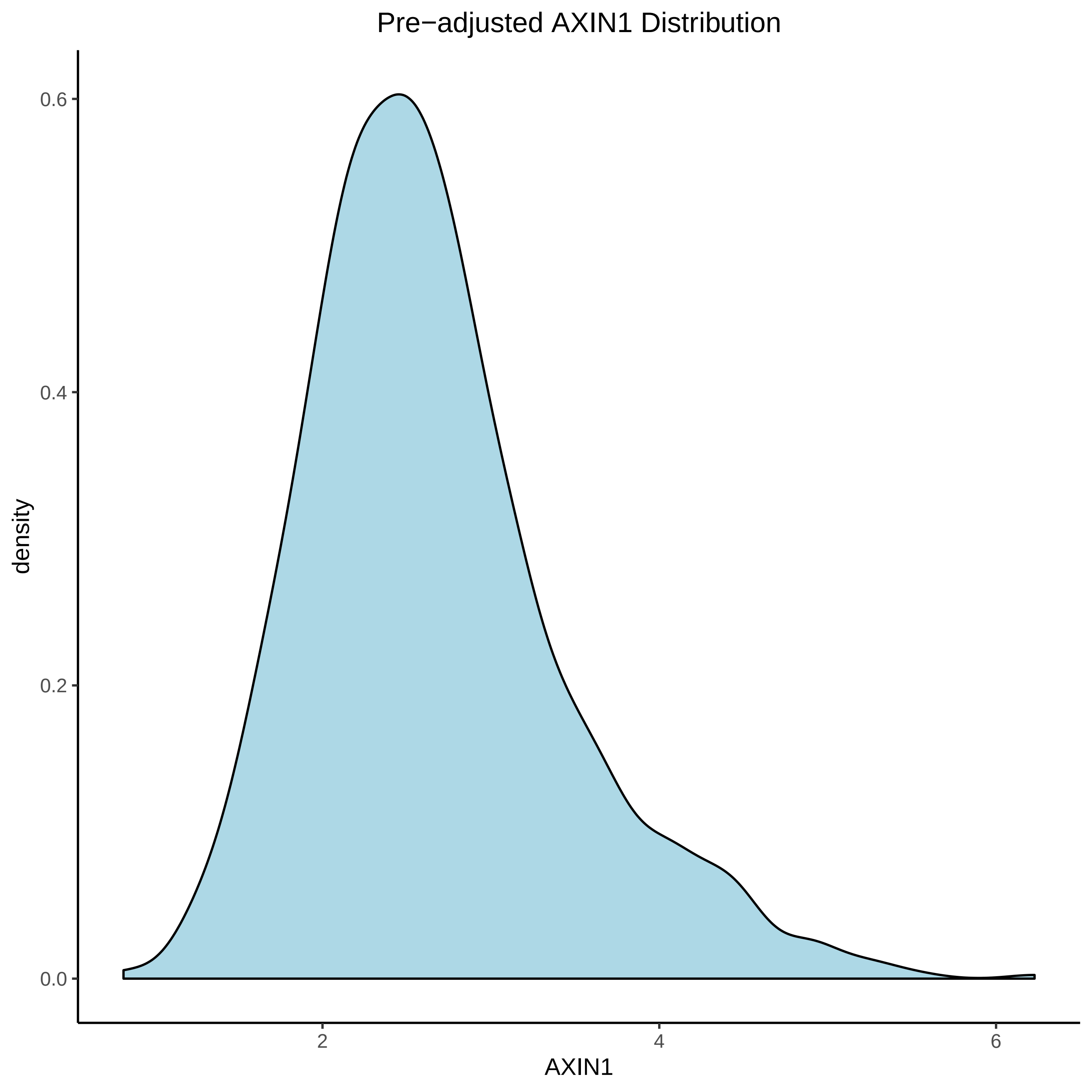

Pre−adjusted AXIN1 Distribution
density
0.6
0.4
0.2
0.0
2
4
6
AXIN1

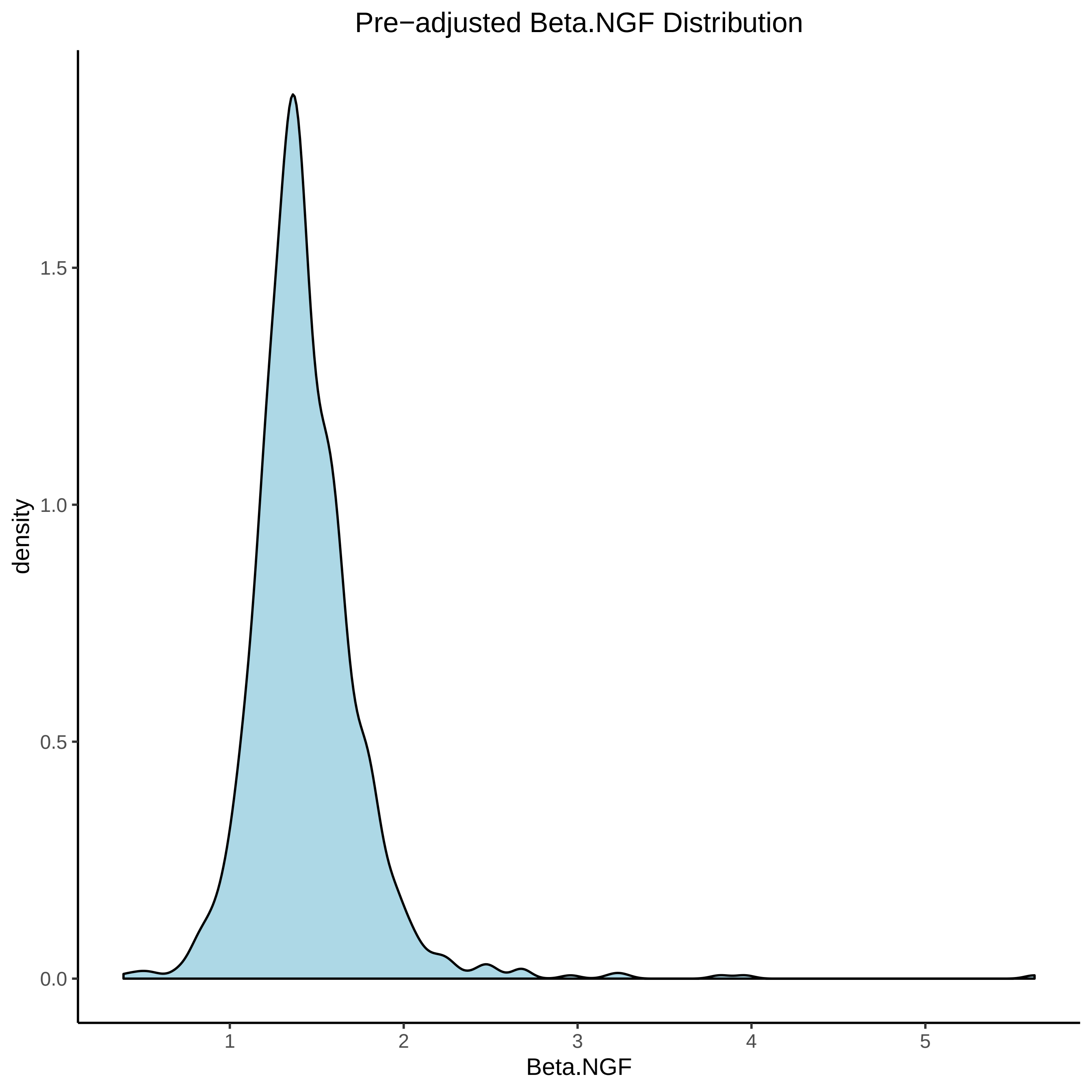
Pre−adjusted Beta.NGF Distribution
density
0.0
0.5
1.0
1.5
1
2
3
4
5
Beta.NGF

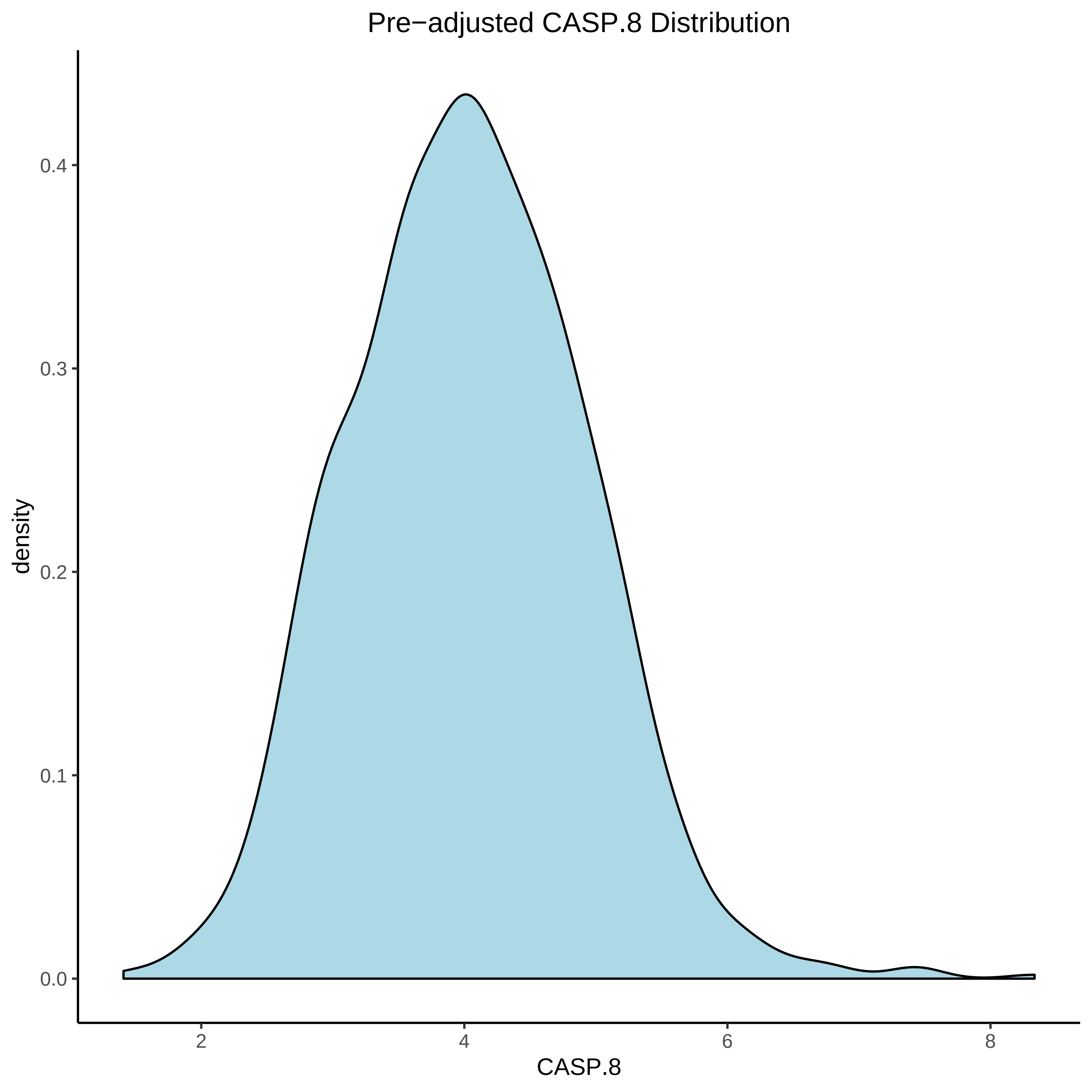

Pre−adjusted CASP.8 Distribution
density
0.0
0.1
0.2
0.3
0.4
2
4
6
8
CASP.8

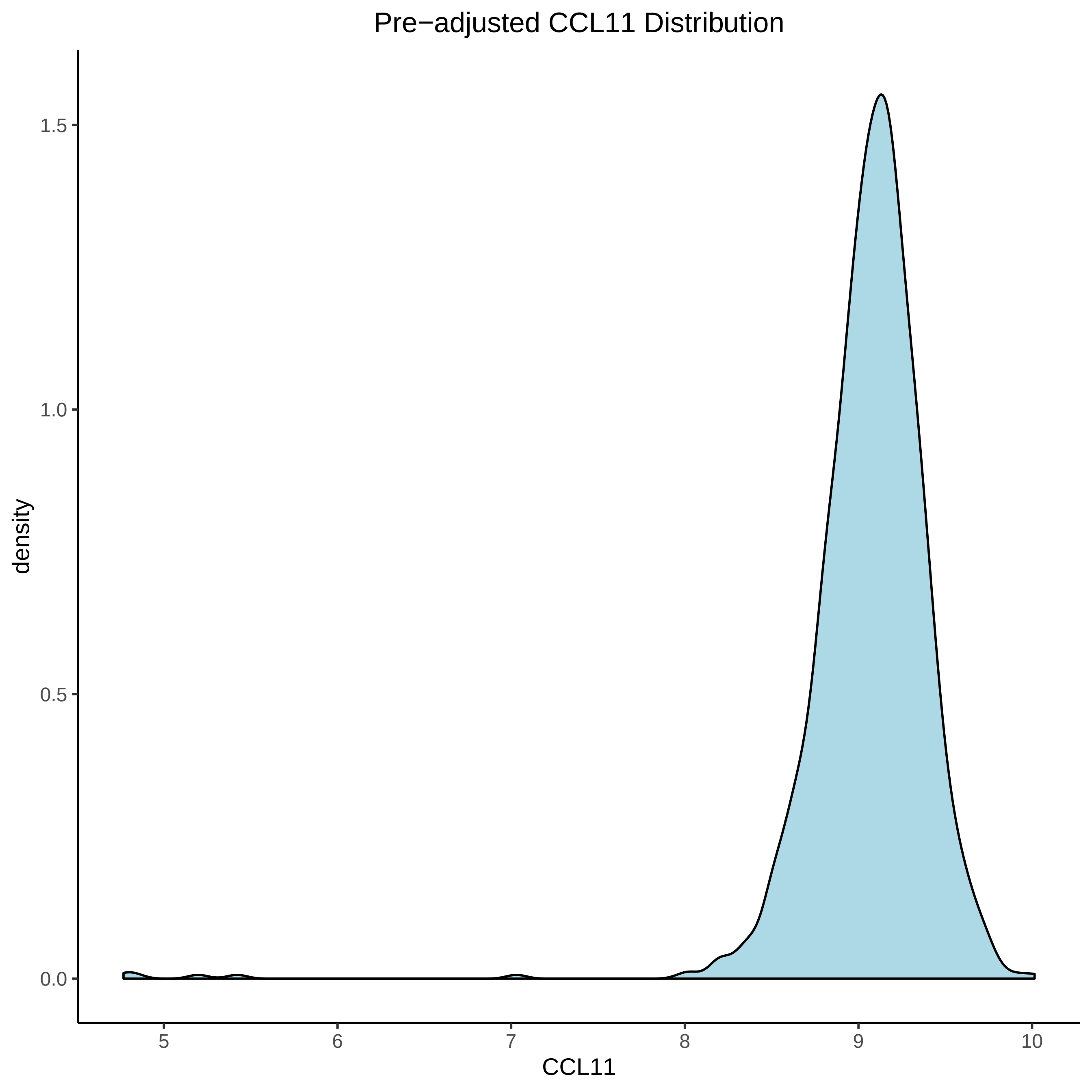

Pre−adjusted CCL11 Distribution
density
1.5
1.0
0.5
0.0
5
6
7
8
9
10
CCL11

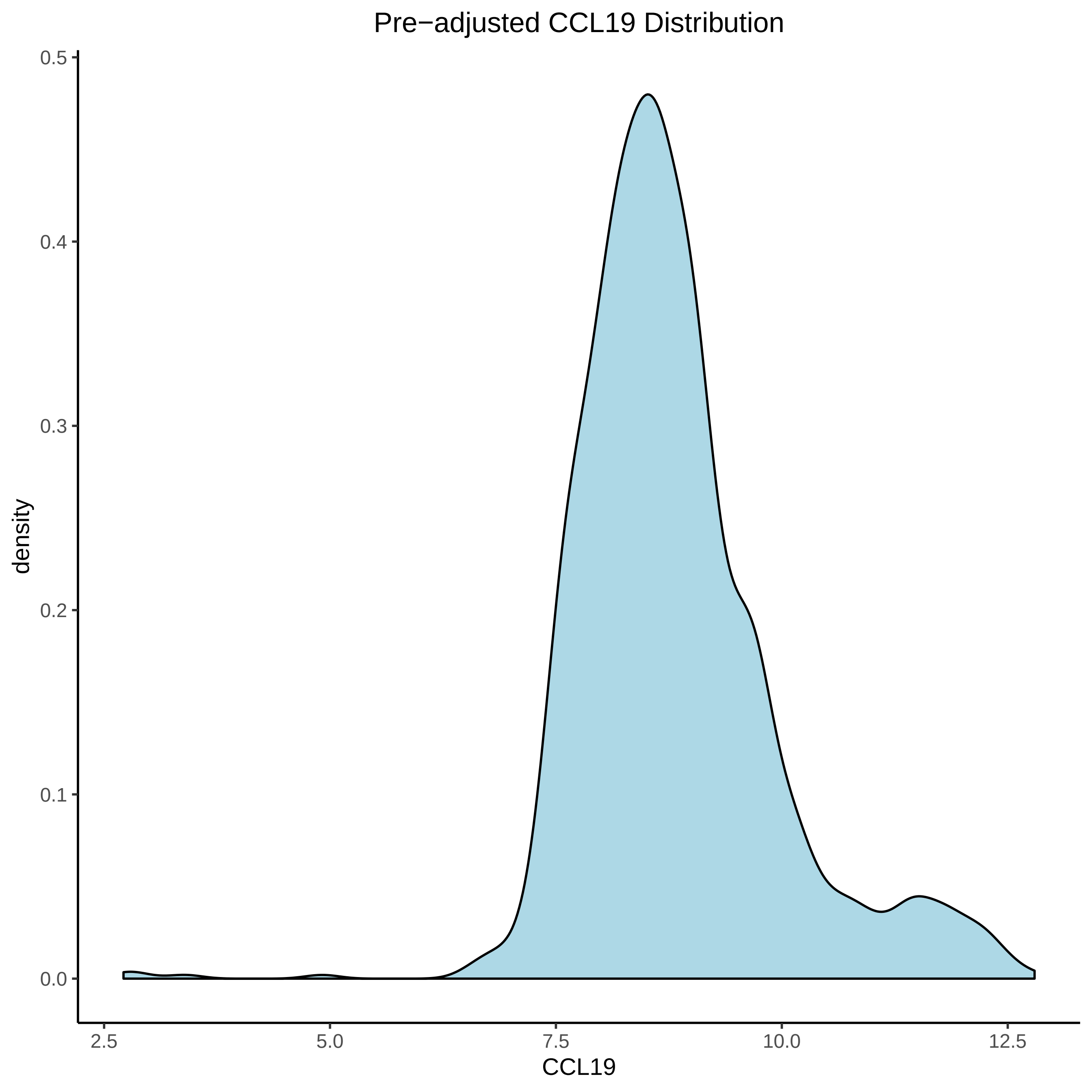

Pre−adjusted CCL19 Distribution
0.5
0.4
0.3
0.2
0.1
0.0
density
2.5
5.0
7.5
10.0
12.5
CCL19

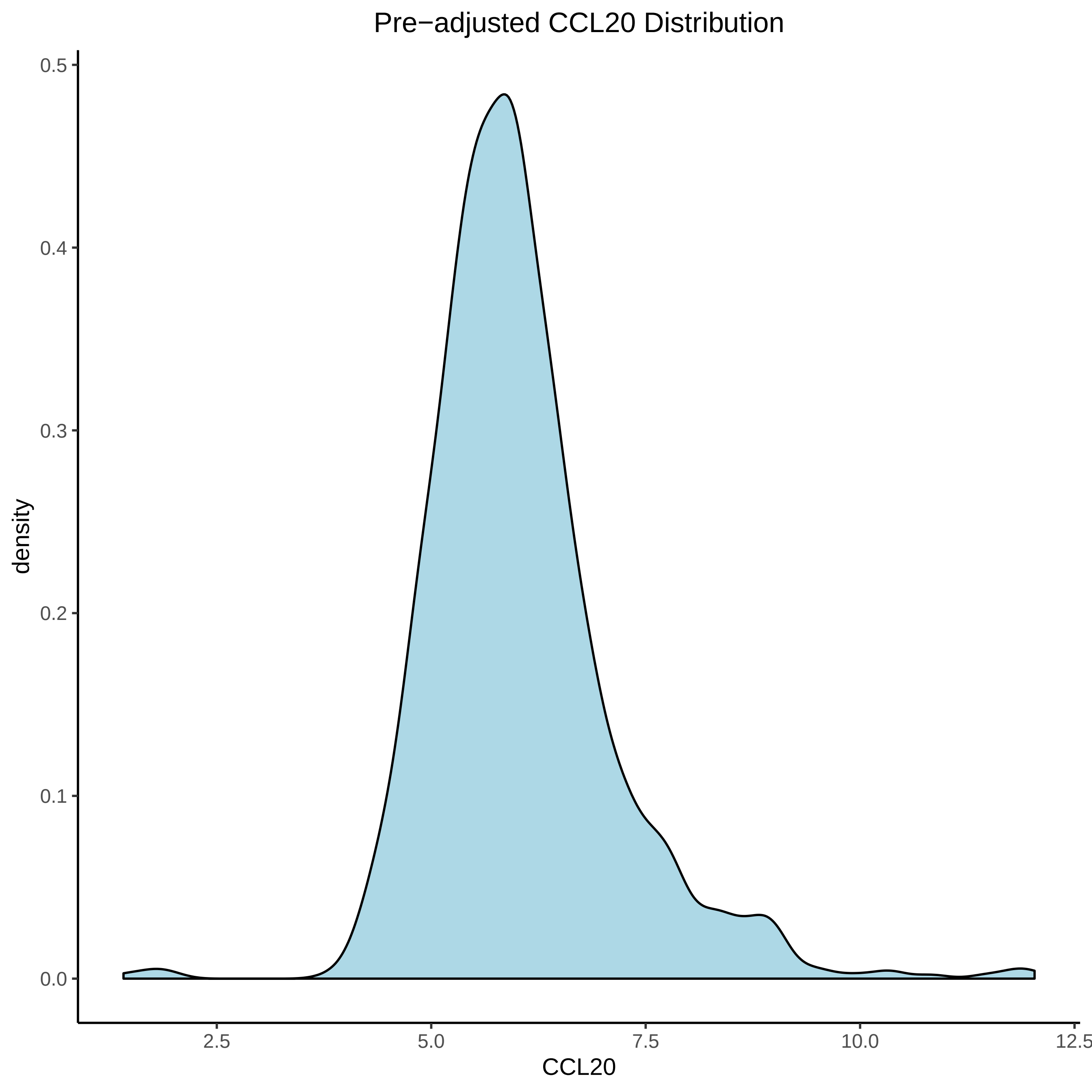
Pre−adjusted CCL20 Distribution
density
0.0
0.1
0.2
0.3
0.4
0.5
2.5
5.0
7.5
10.0
12.5
CCL20

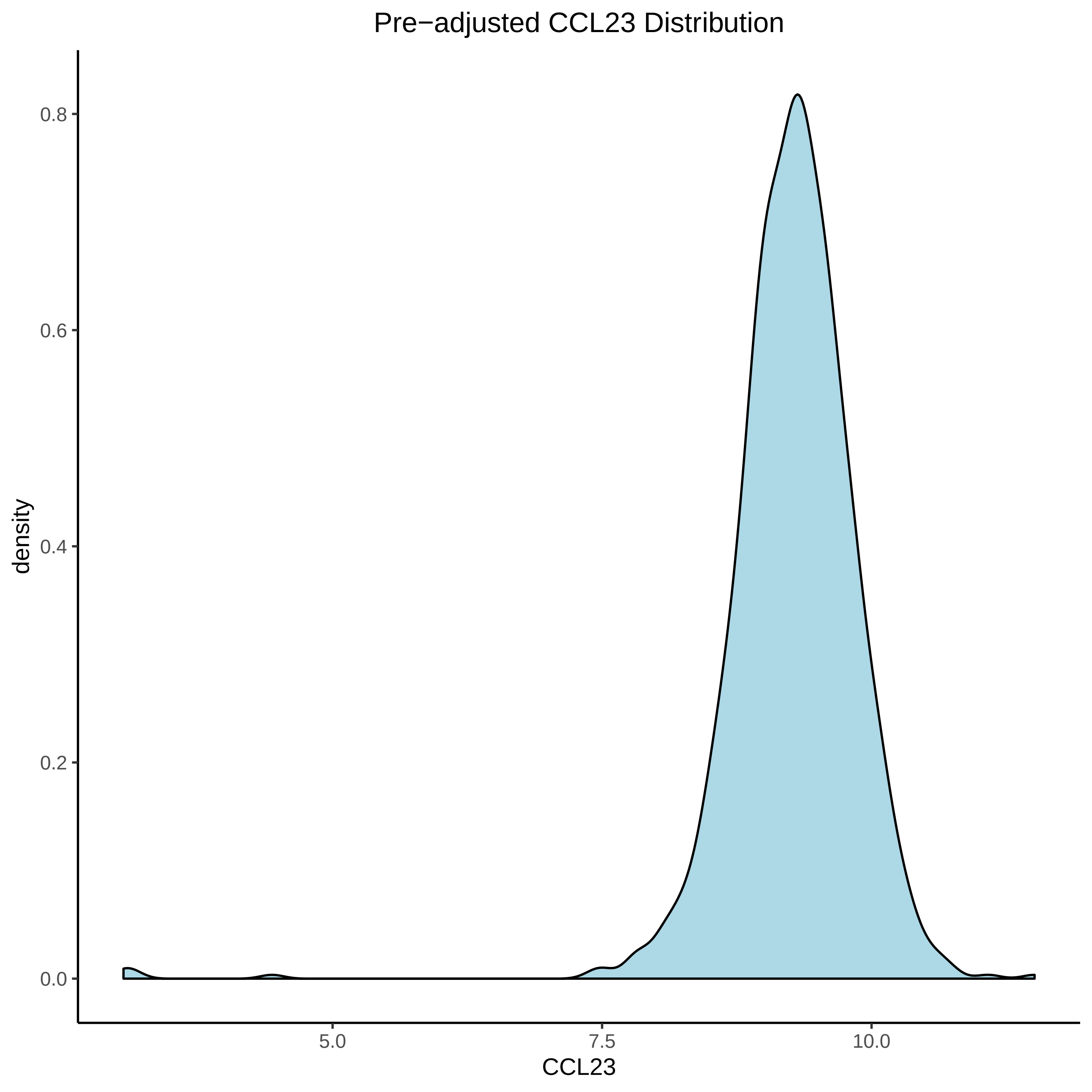

Pre−adjusted CCL23 Distribution
0.8
0.6
0.4
0.2
0.0
density
5.0
7.5
10.0
CCL23

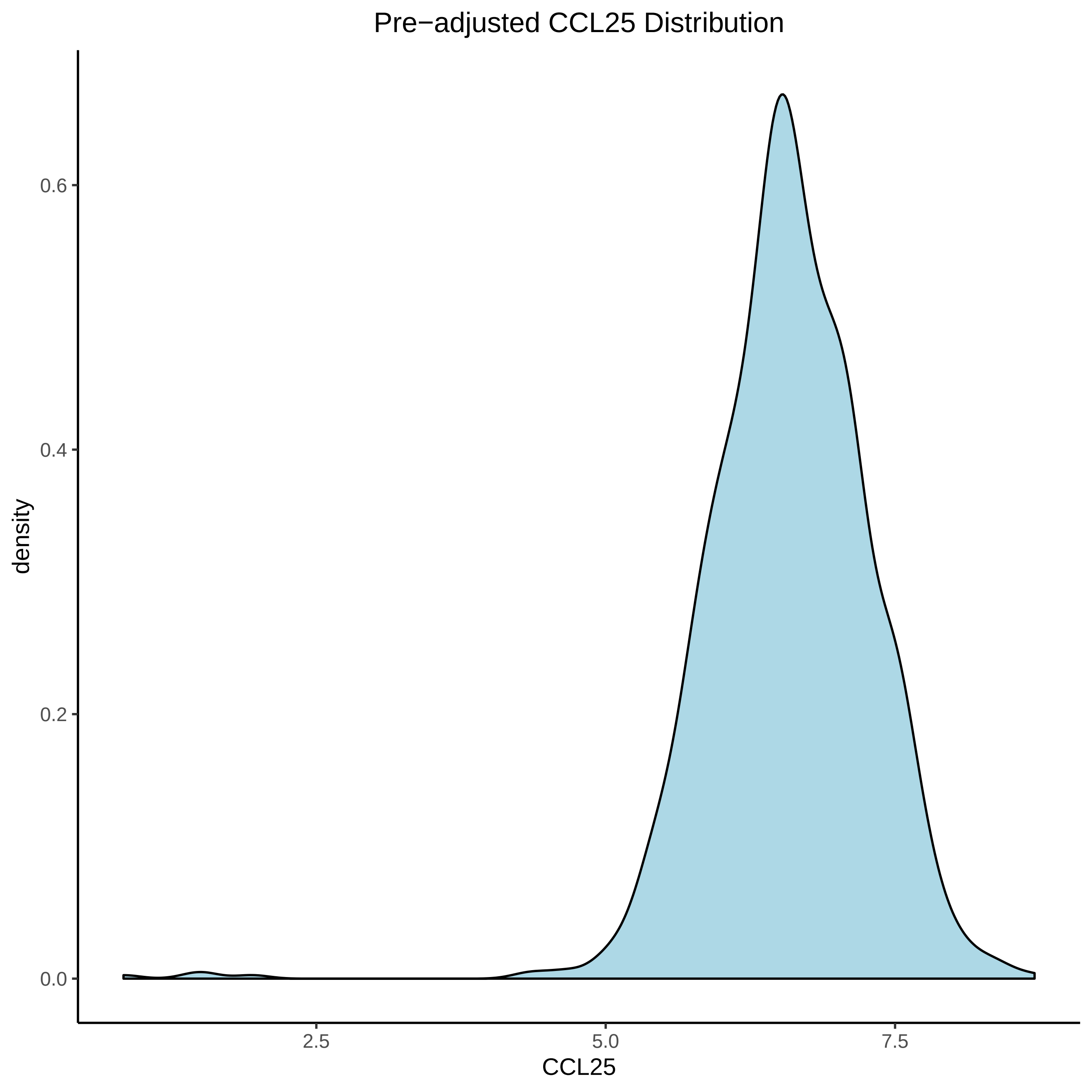

Pre−adjusted CCL25 Distribution
density
0.6
0.4
0.2
0.0
2.5
5.0
7.5
CCL25

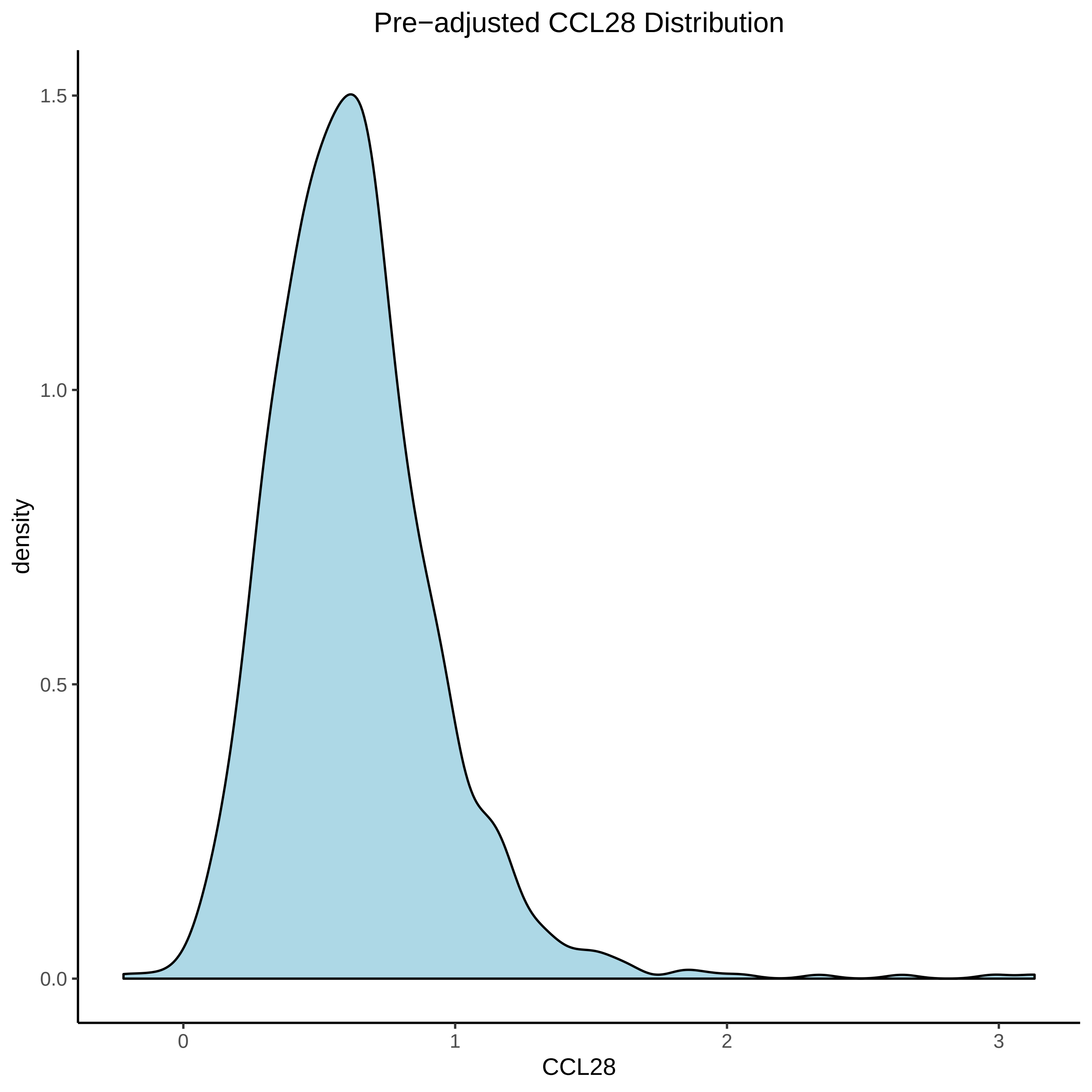

Pre−adjusted CCL28 Distribution
density
1.5
1.0
0.5
0.0
0
1
2
3
CCL28

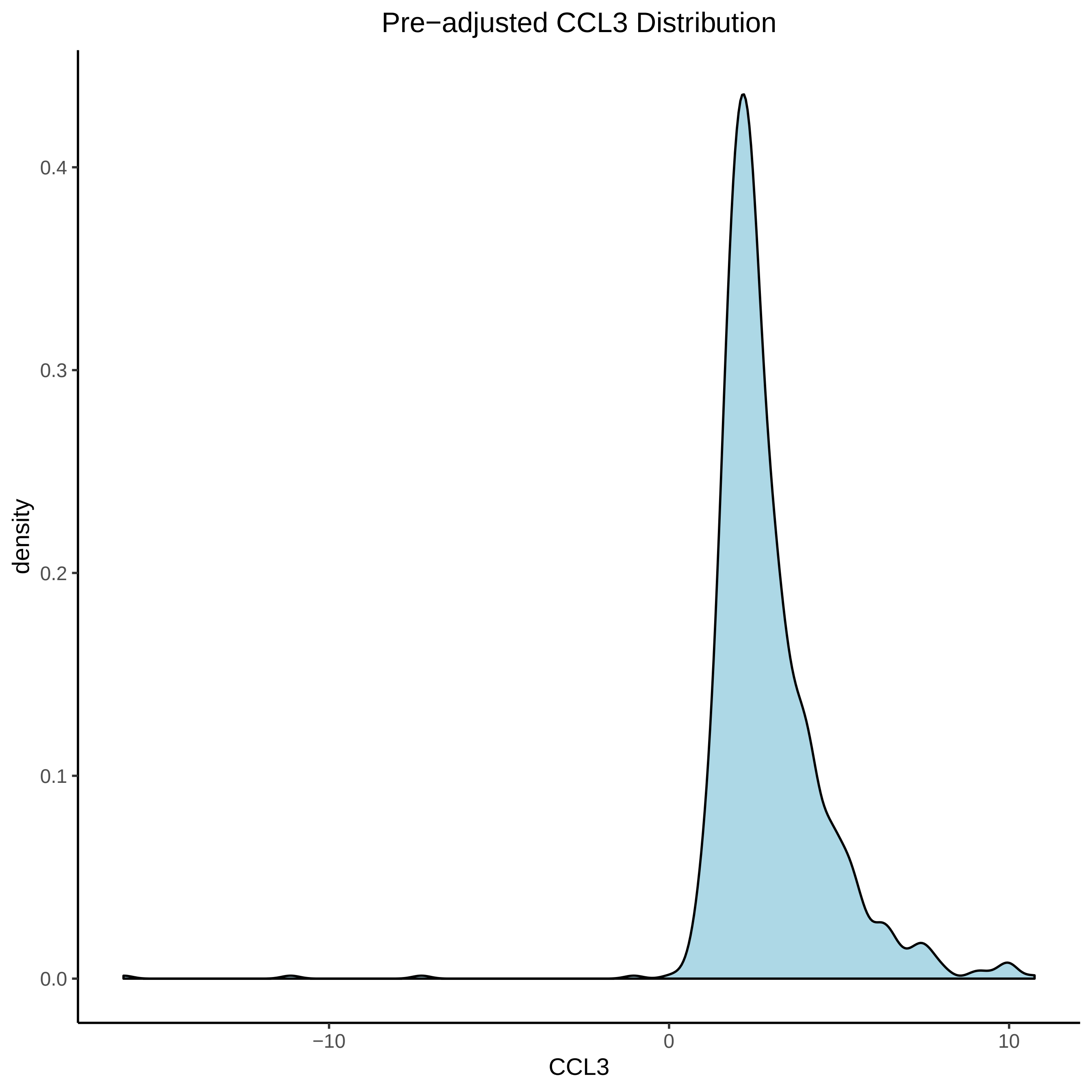
Pre−adjusted CCL3 Distribution
density
0.4
0.3
0.2
0.1
0.0
−10
0
10
CCL3

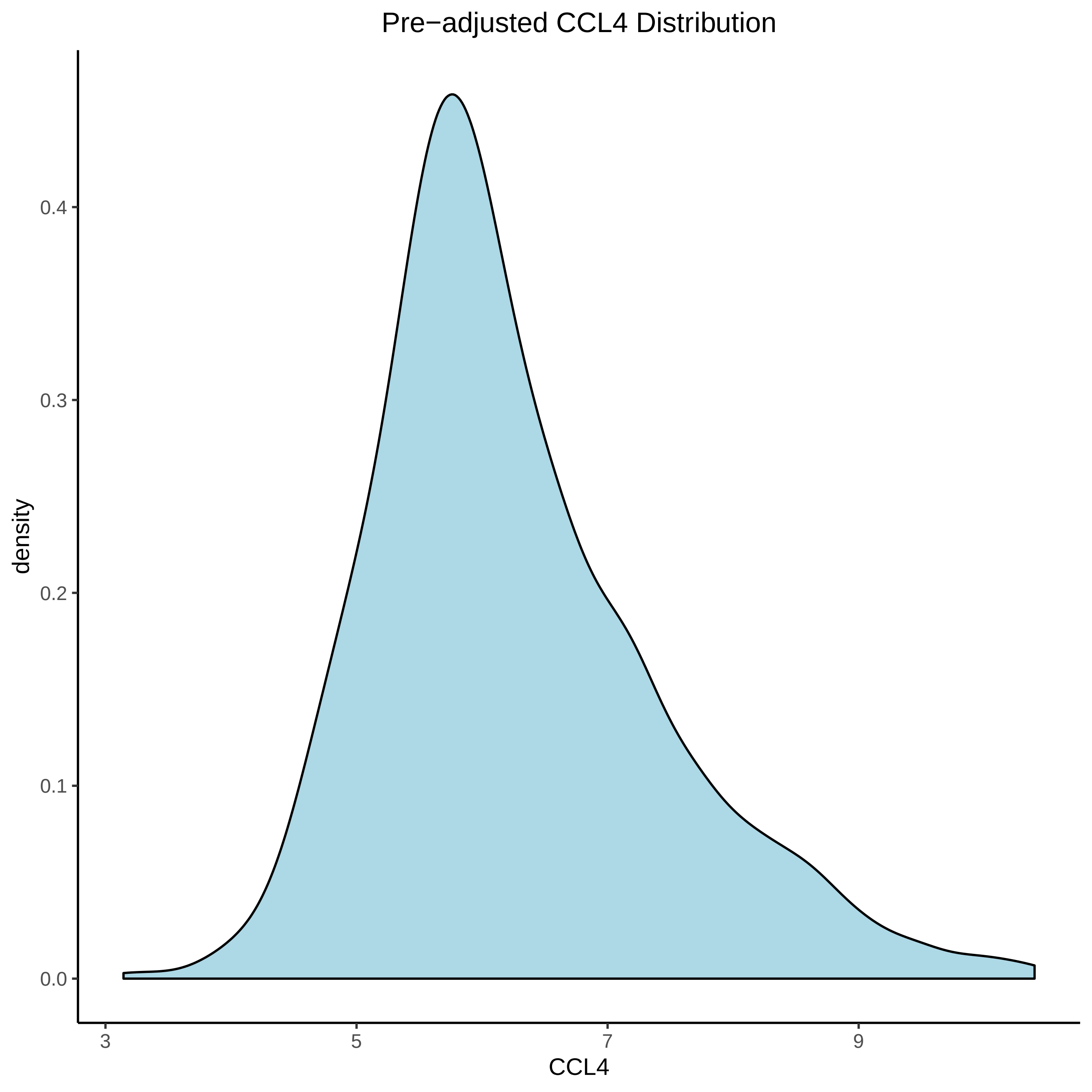

Pre−adjusted CCL4 Distribution
density
0.0
0.1
0.2
0.3
0.4
3
5
7
9
CCL4

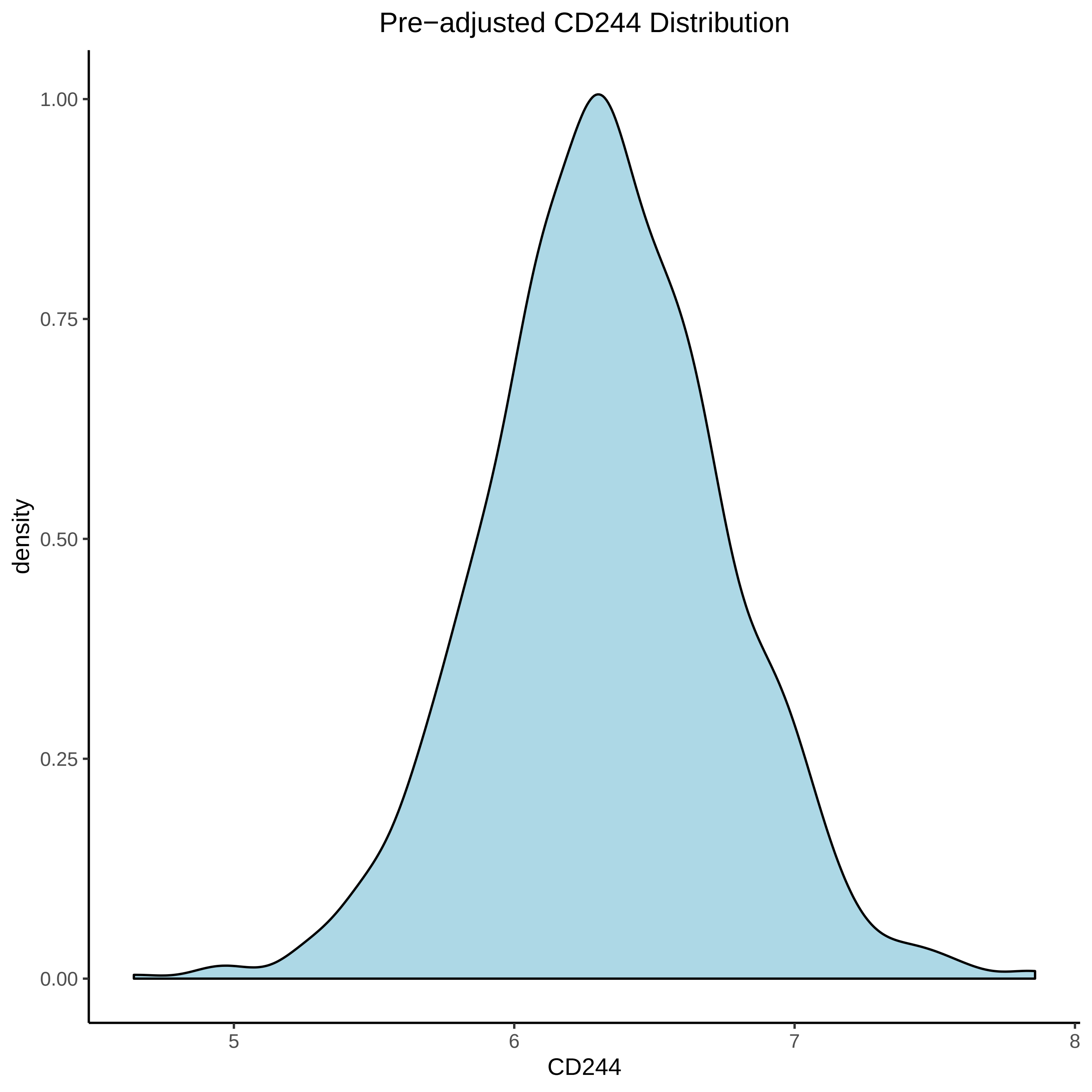
Pre−adjusted CD244 Distribution
density
1.00
0.75
0.50
0.25
0.00
5
6
7
8
CD244

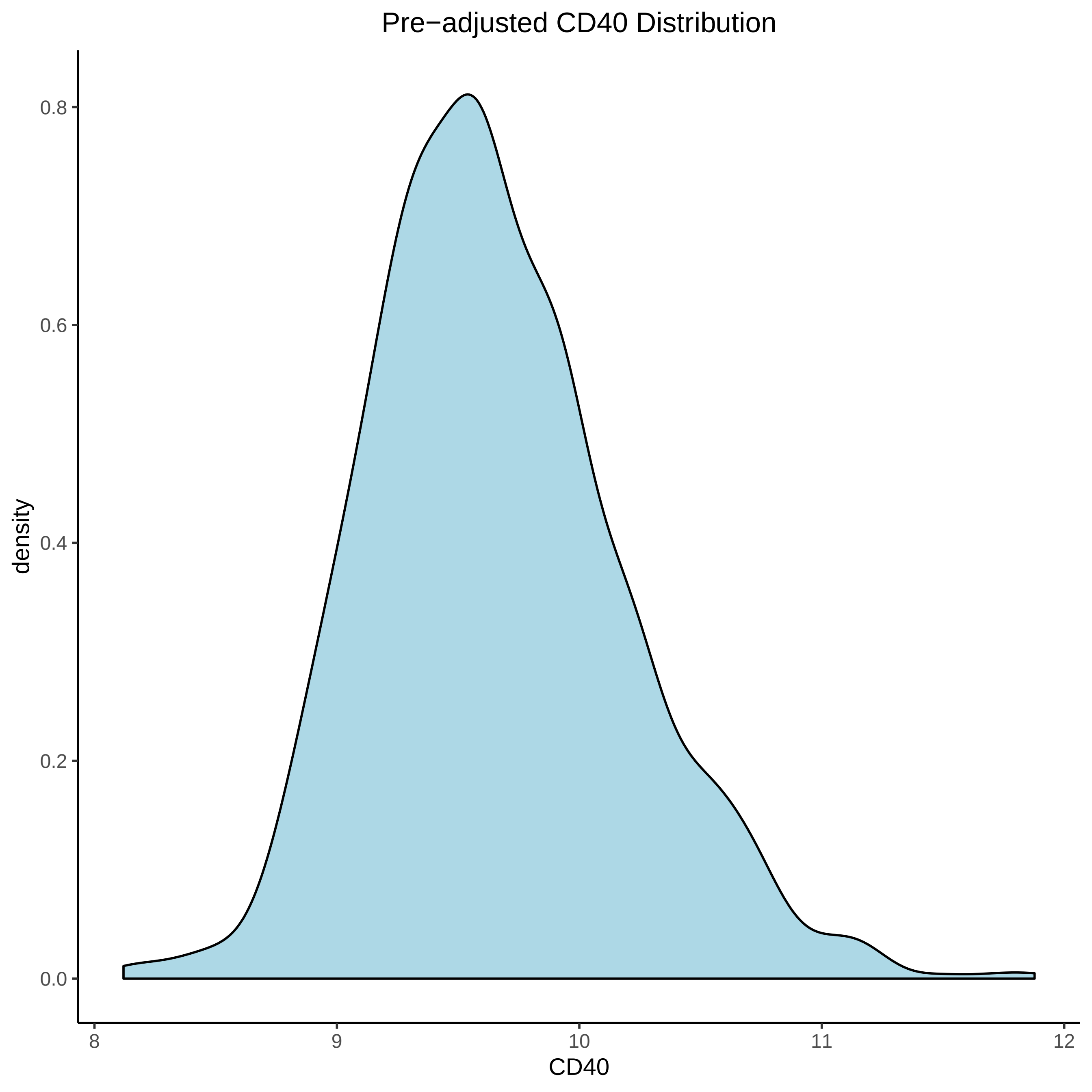

Pre−adjusted CD40 Distribution
density
0.0
0.2
0.4
0.6
0.8
8
9
10
11
12
CD40

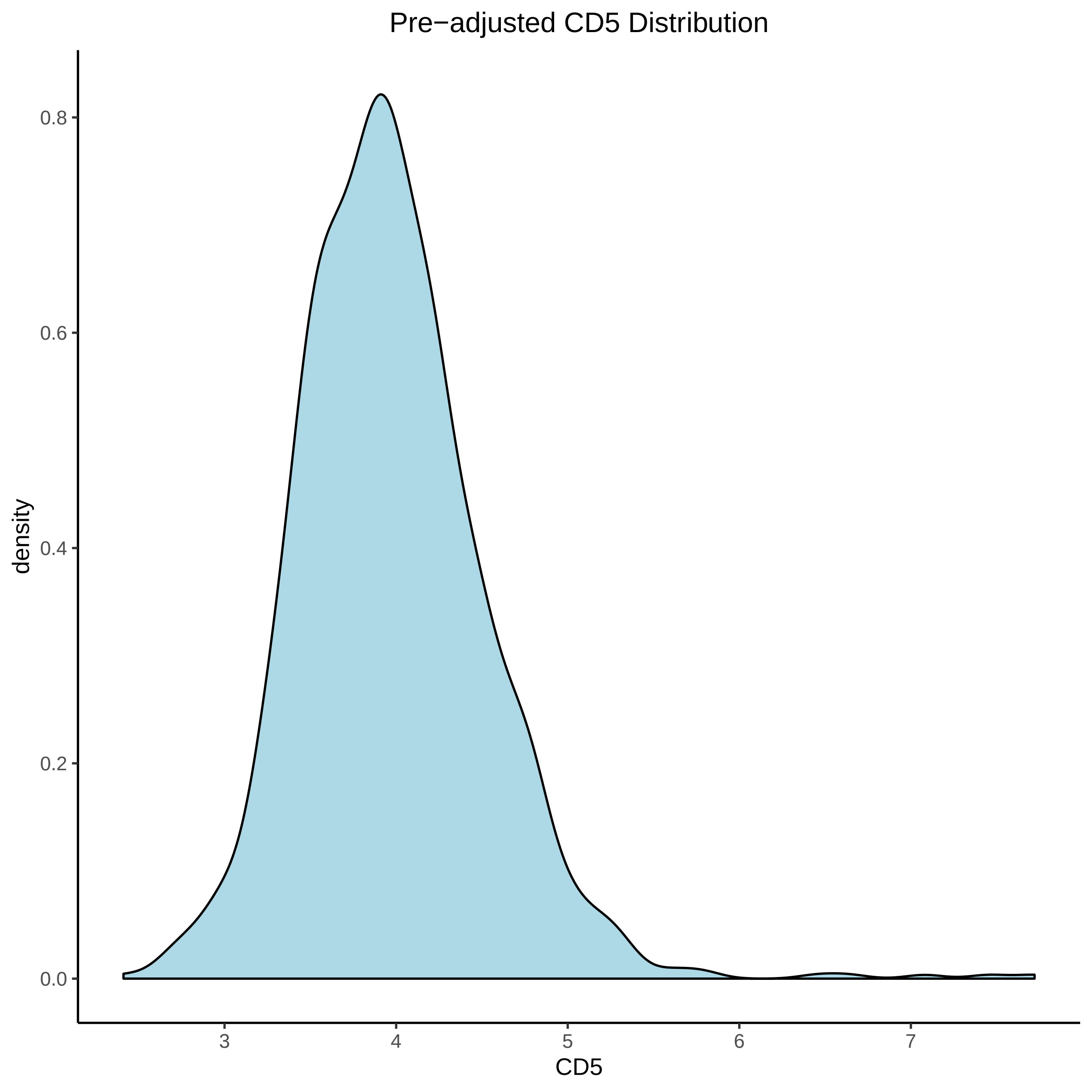
Pre−adjusted CD5 Distribution
density
0.0
0.2
0.4
0.6
0.8
3
4
5
6
7
CD5

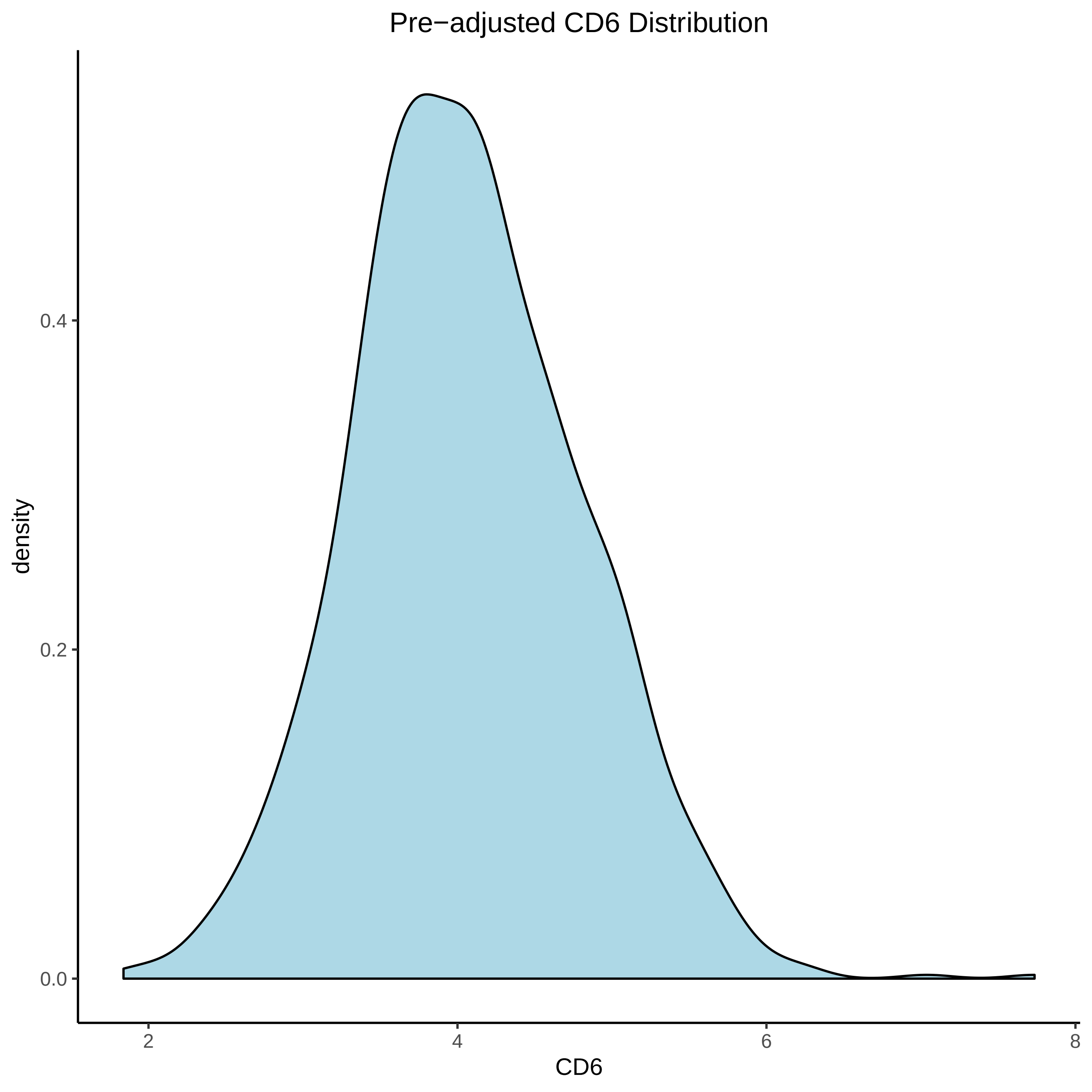

Pre−adjusted CD6 Distribution
density
0.4
0.2
0.0
2
4
6
8
CD6

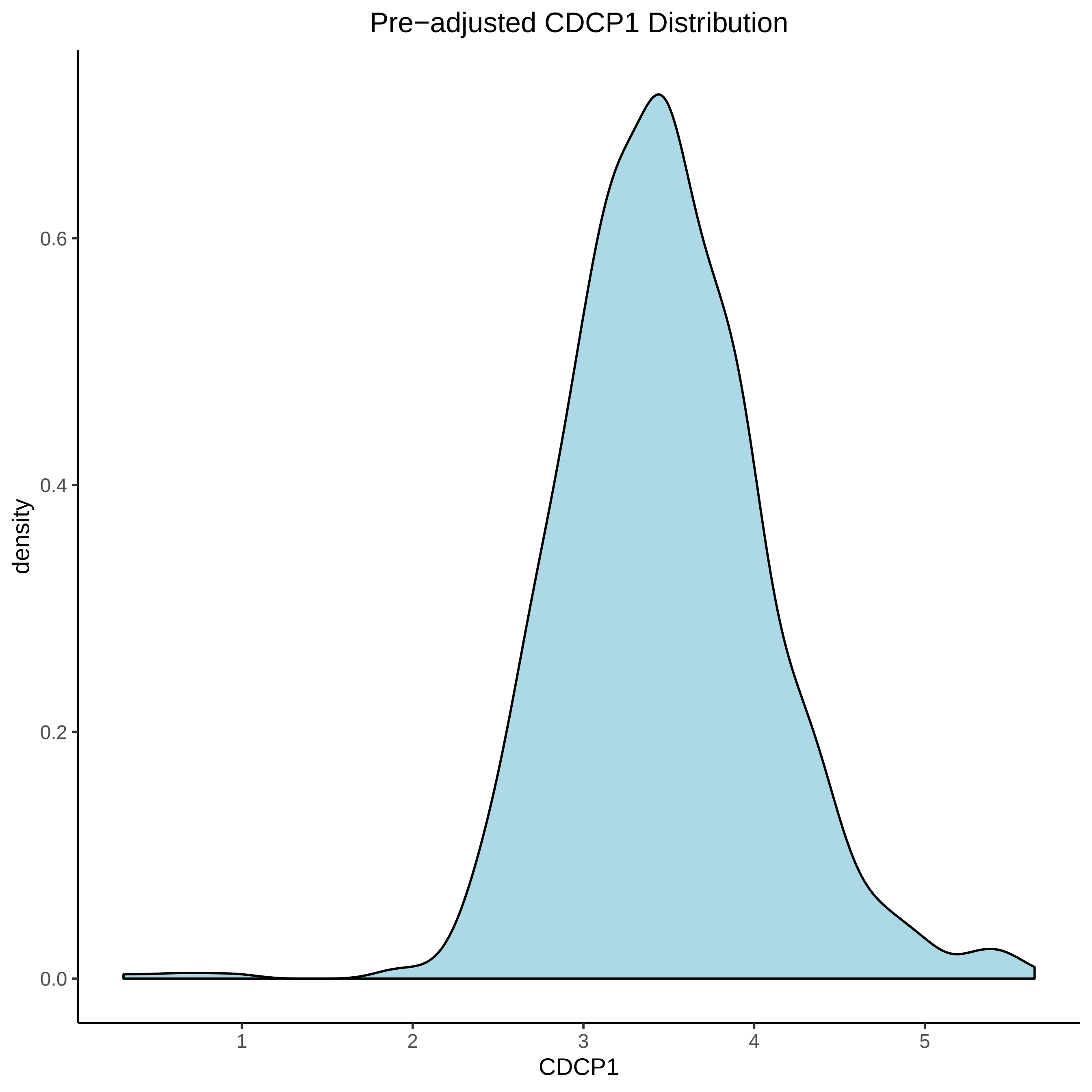

Pre−adjusted CDCP1 Distribution
density
0.6
0.4
0.2
0.0
1
2
3
4
5
CDCP1

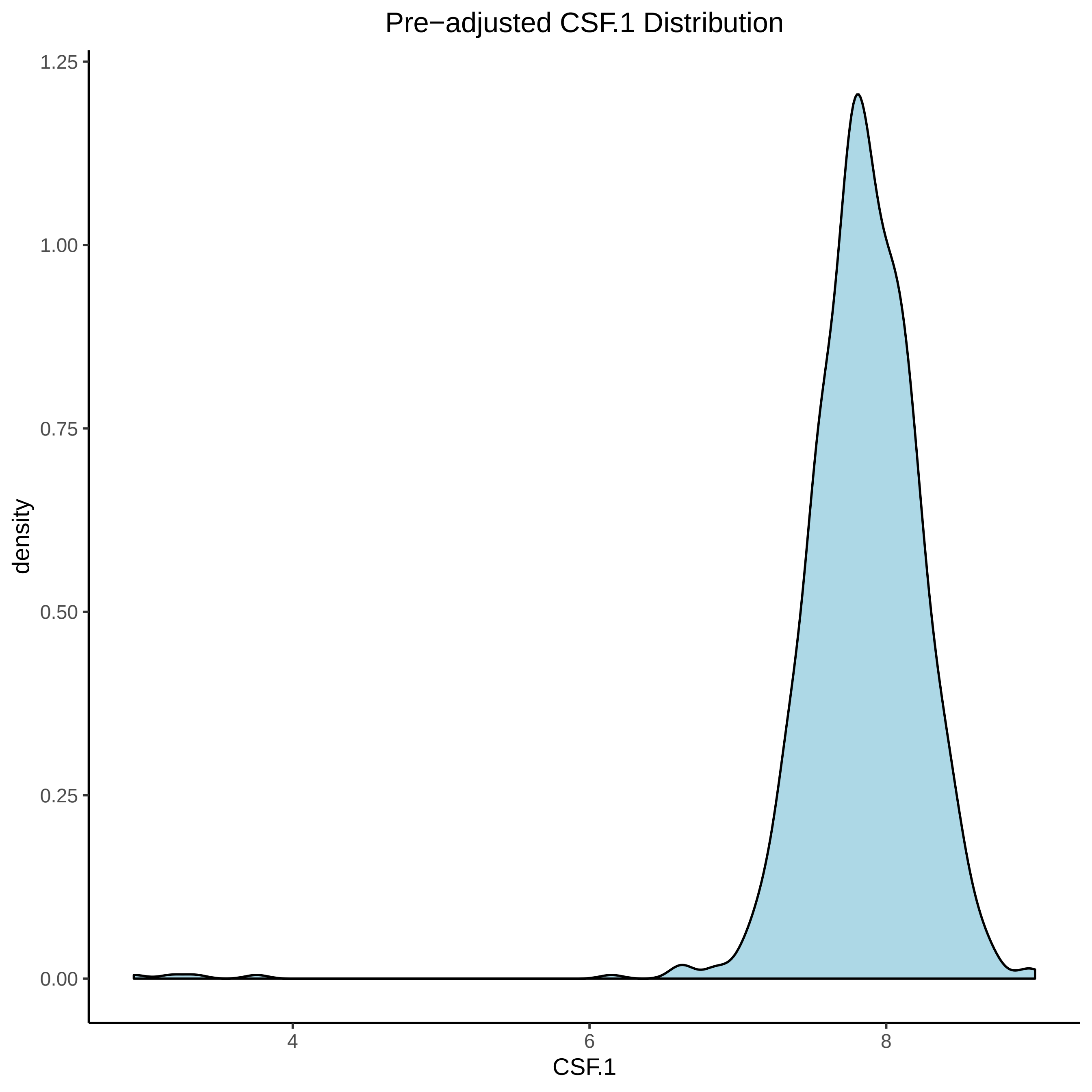

Pre−adjusted CSF.1 Distribution
1.25
1.00
0.75
0.50
0.25
0.00
density
4
6
8
CSF.1

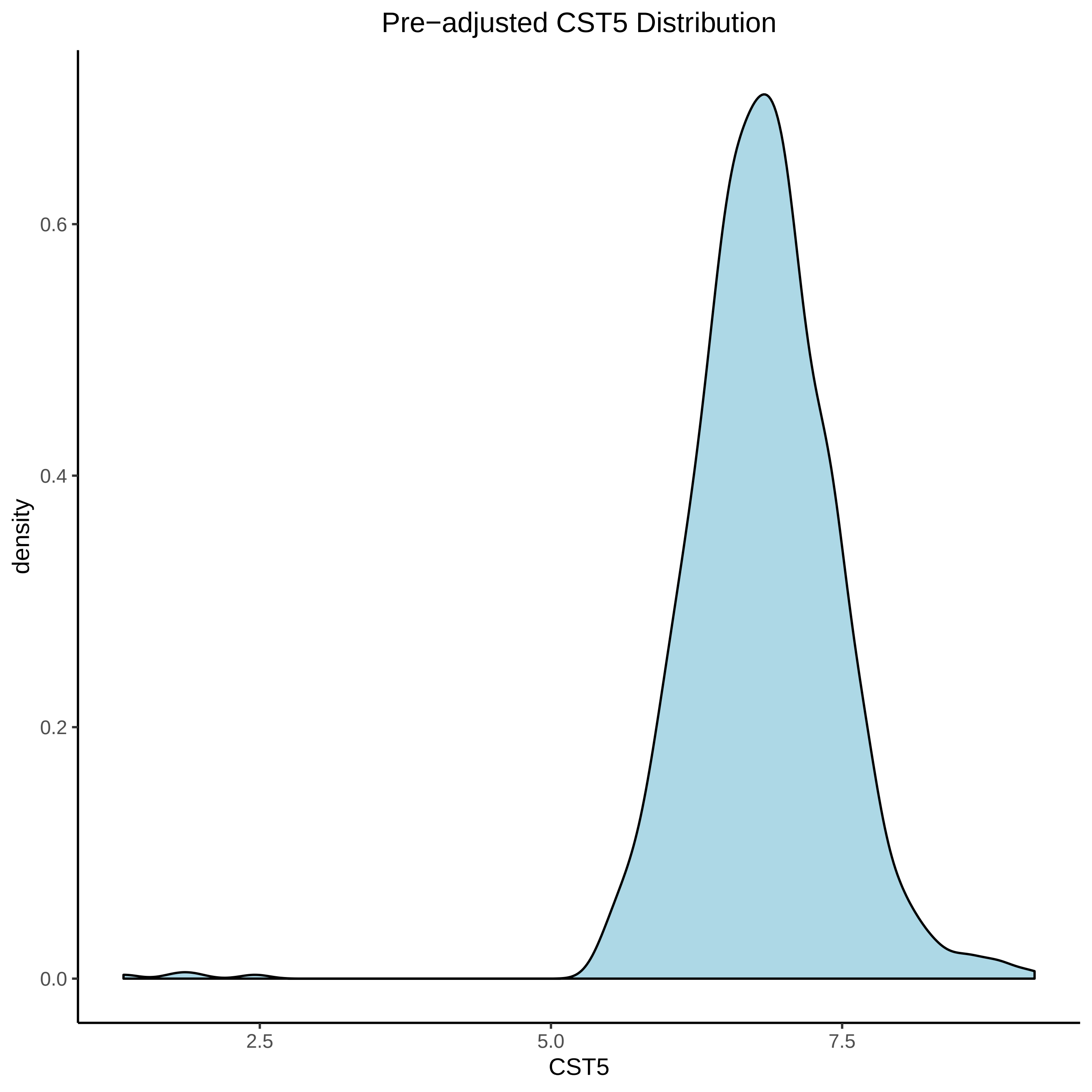

Pre−adjusted CST5 Distribution
density
0.6
0.4
0.2
0.0
2.5
5.0
7.5
CST5

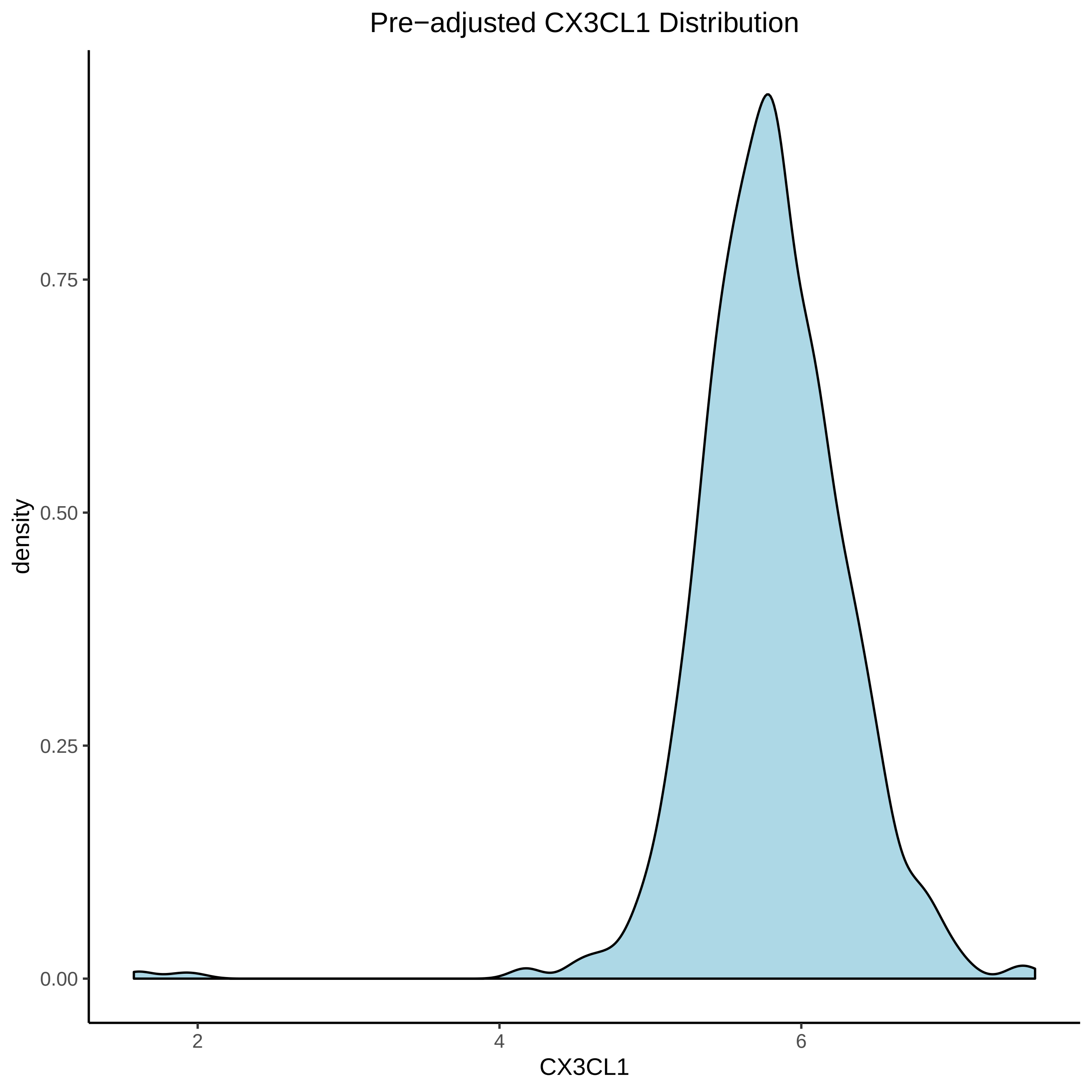

Pre−adjusted CX3CL1 Distribution
density
0.75
0.50
0.25
0.00
2
4
6
CX3CL1

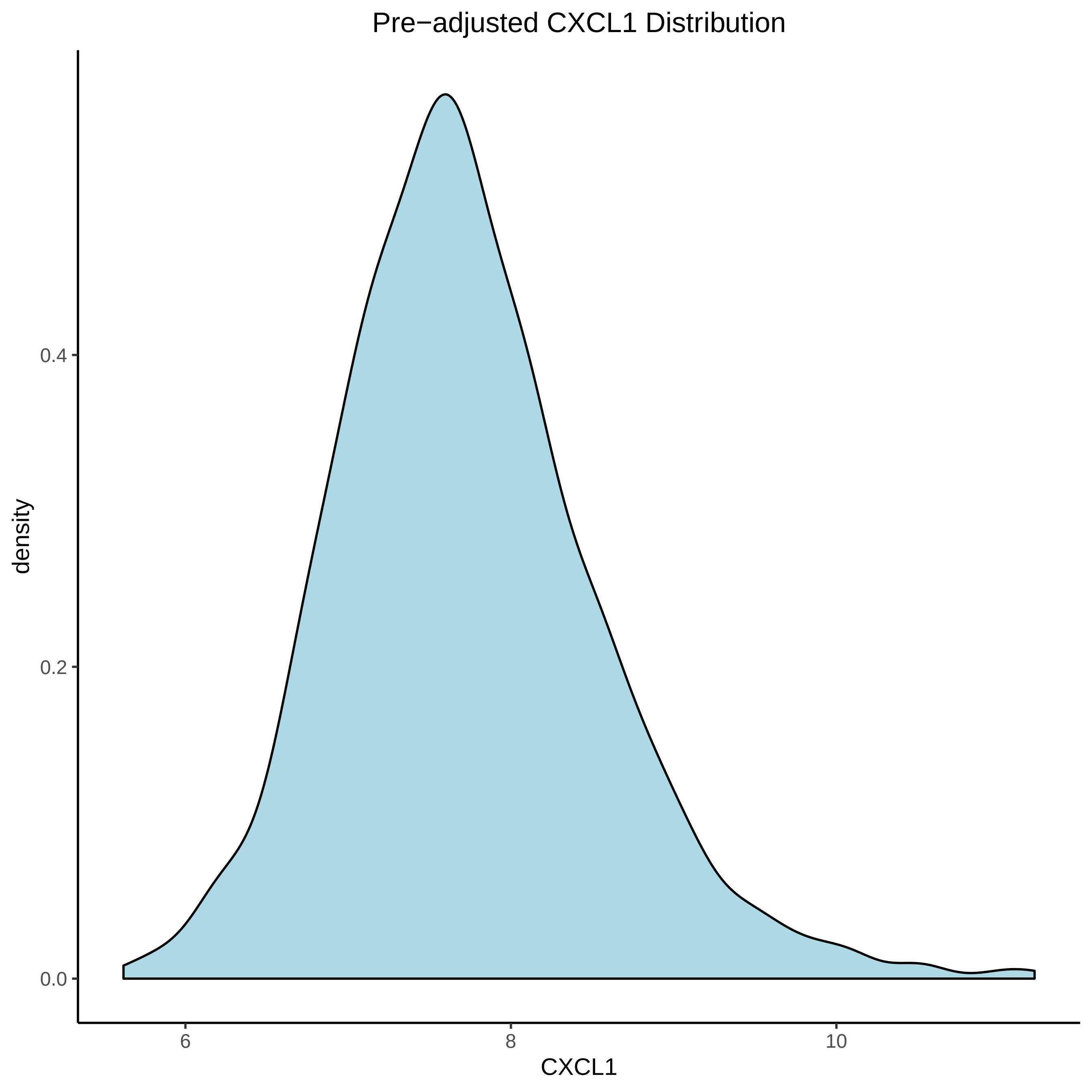

Pre−adjusted CXCL1 Distribution
density
0.0
0.2
0.4
6
8
10
CXCL1

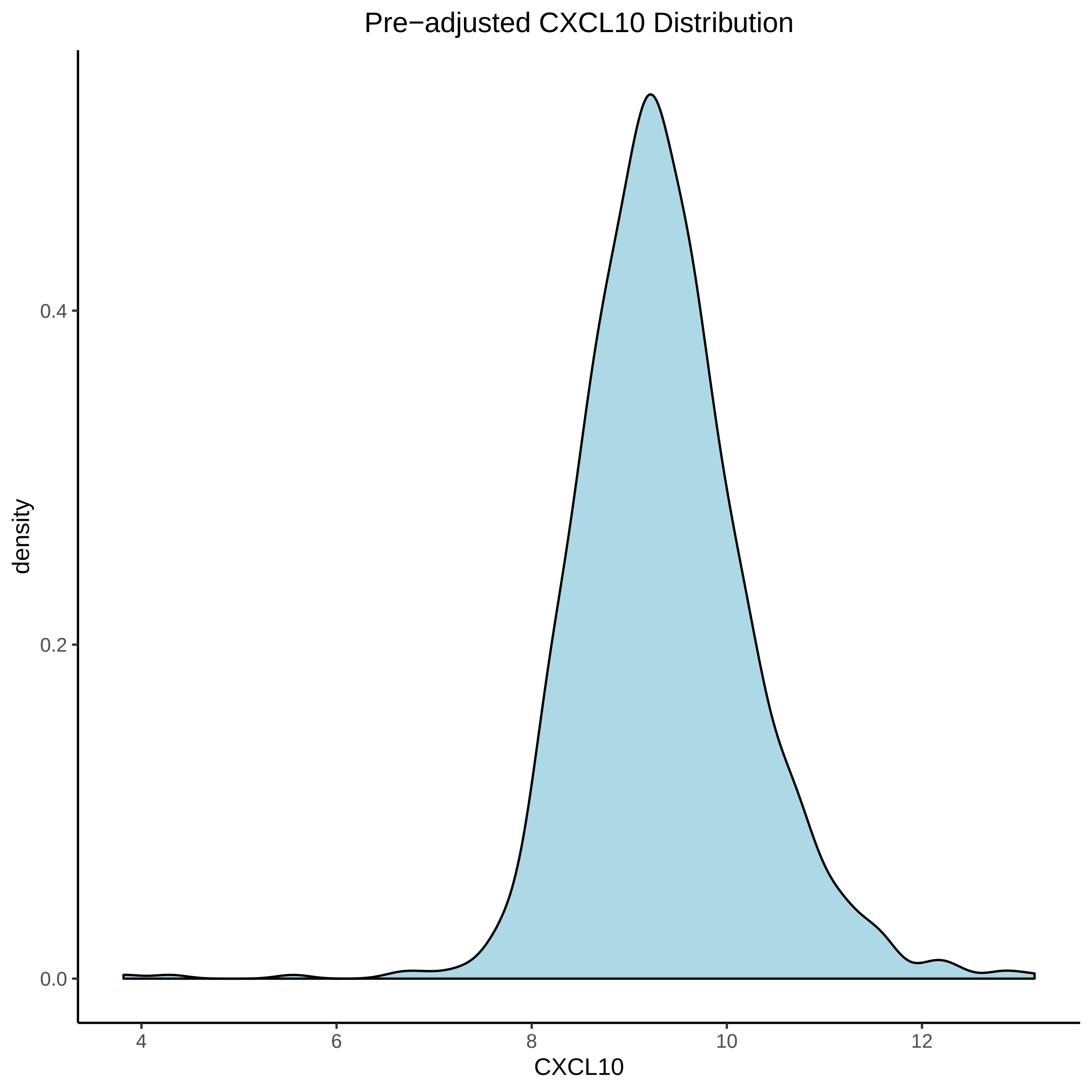

Pre−adjusted CXCL10 Distribution
density
0.0
0.2
0.4
4
6
8
10
12
CXCL10

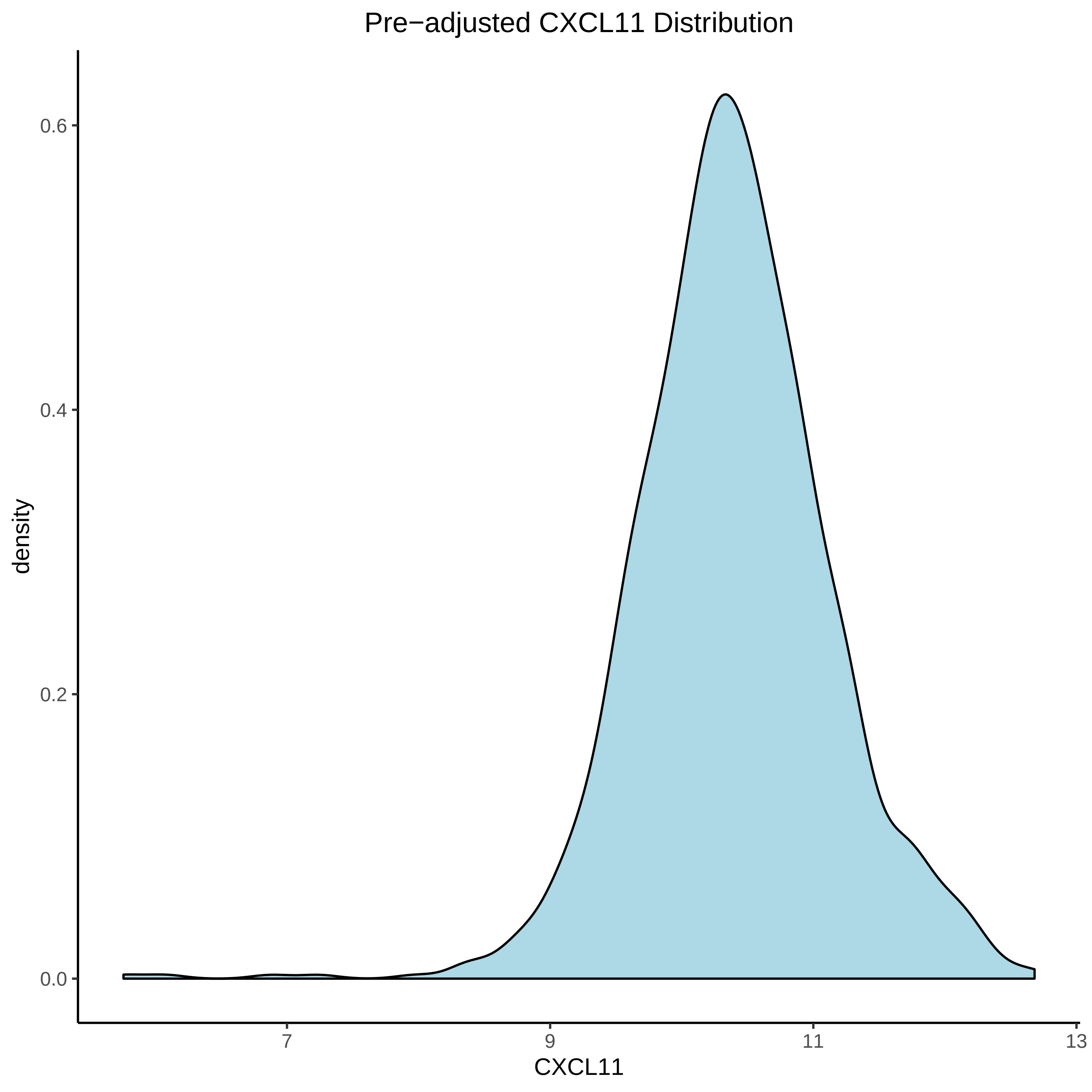

Pre−adjusted CXCL11 Distribution
density
0.6
0.4
0.2
0.0
7
9
11
13
CXCL11

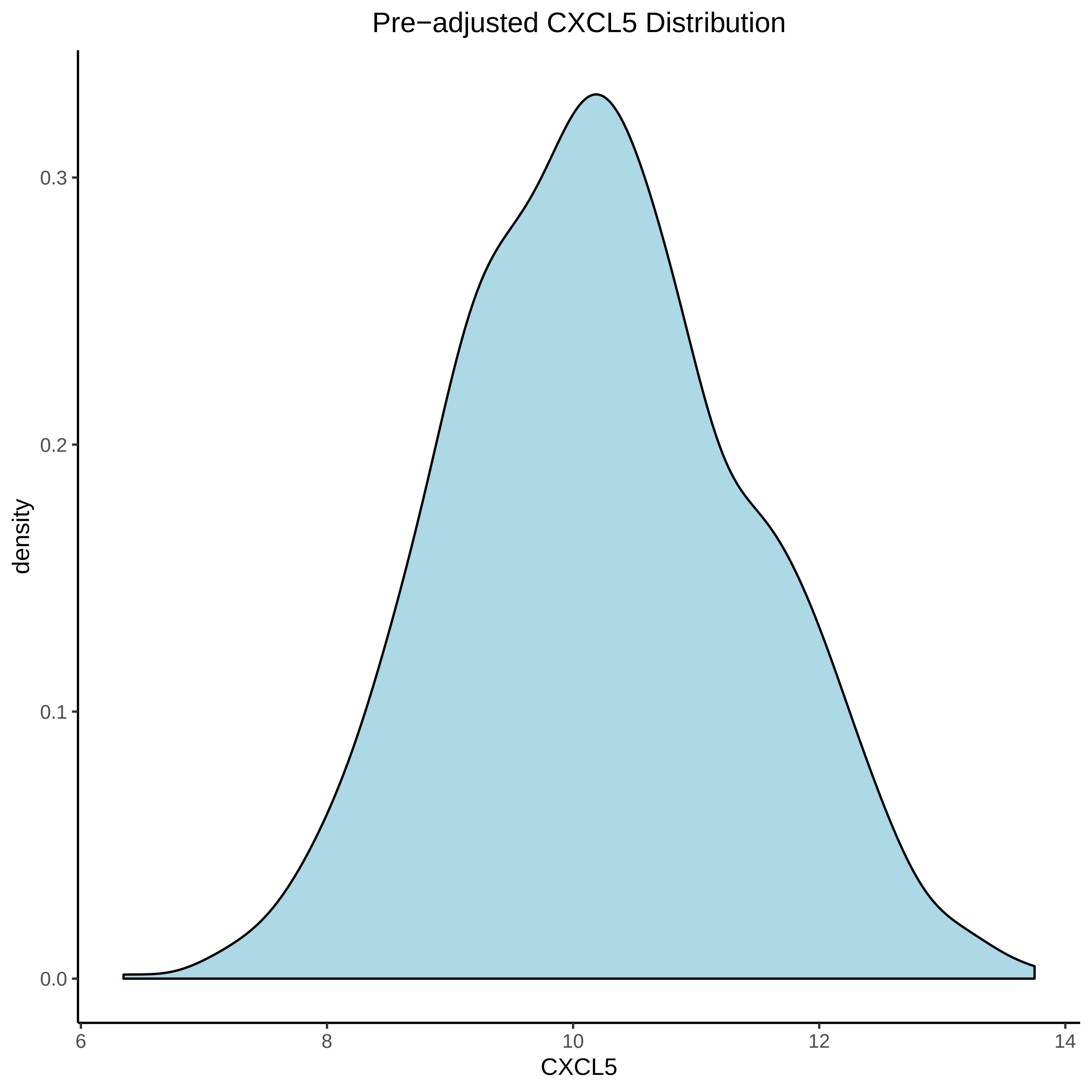
Pre−adjusted CXCL5 Distribution
0.3
0.2
0.1
0.0
density
6
8
10
12
14
CXCL5

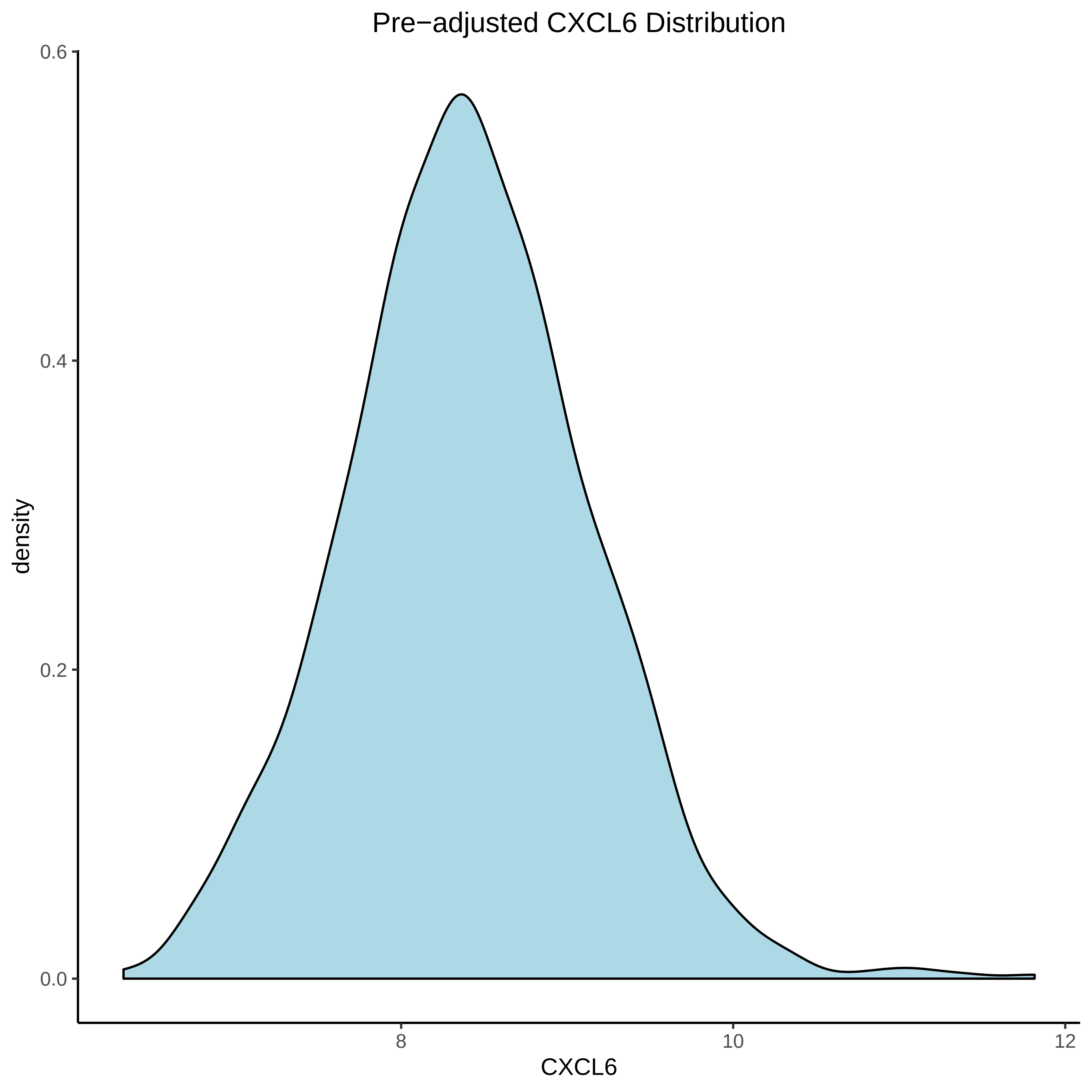
Pre−adjusted CXCL6 Distribution
density
0.6
0.4
0.2
0.0
8
10
12
CXCL6

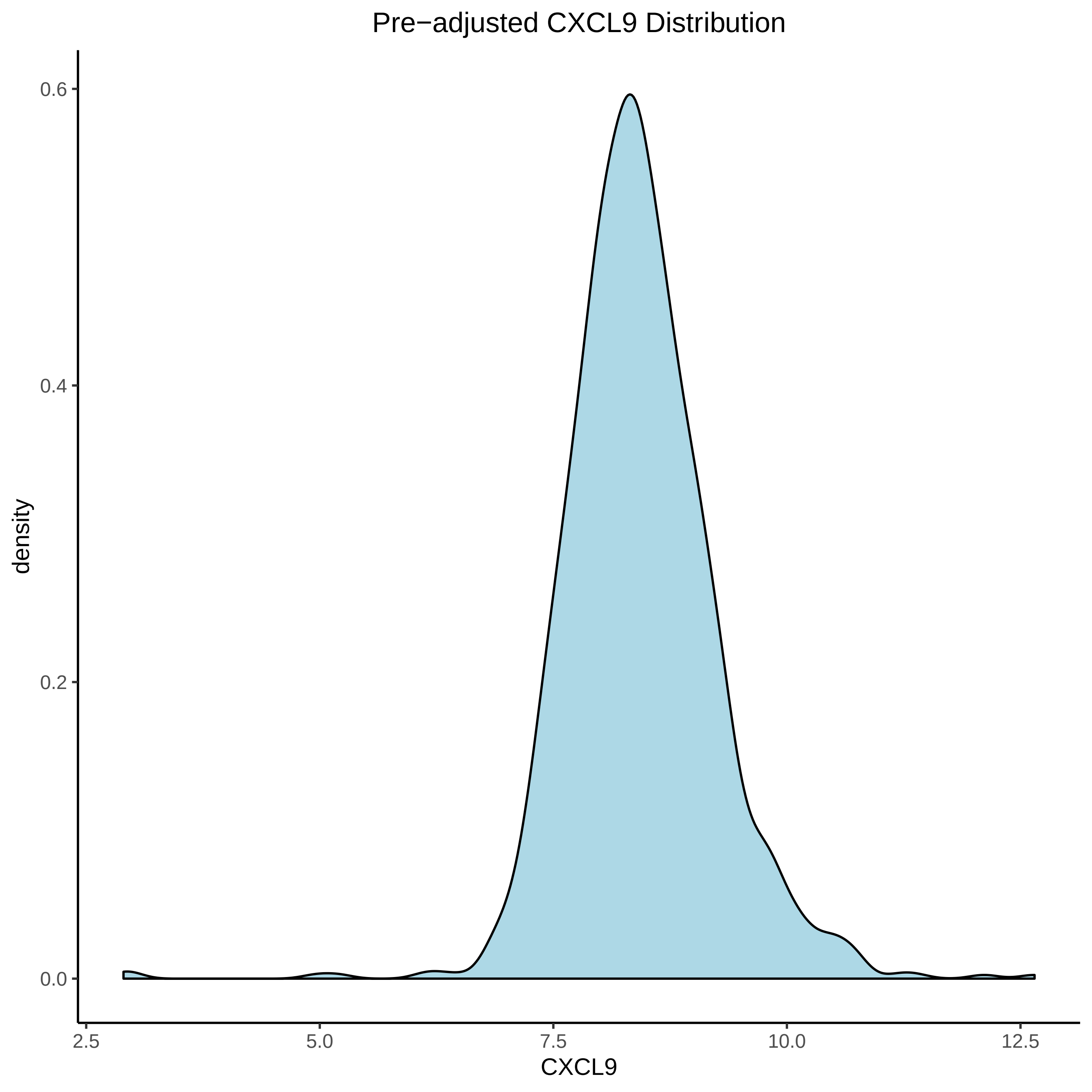

Pre−adjusted CXCL9 Distribution
density
0.6
0.4
0.2
0.0
2.5
5.0
7.5
10.0
12.5
CXCL9

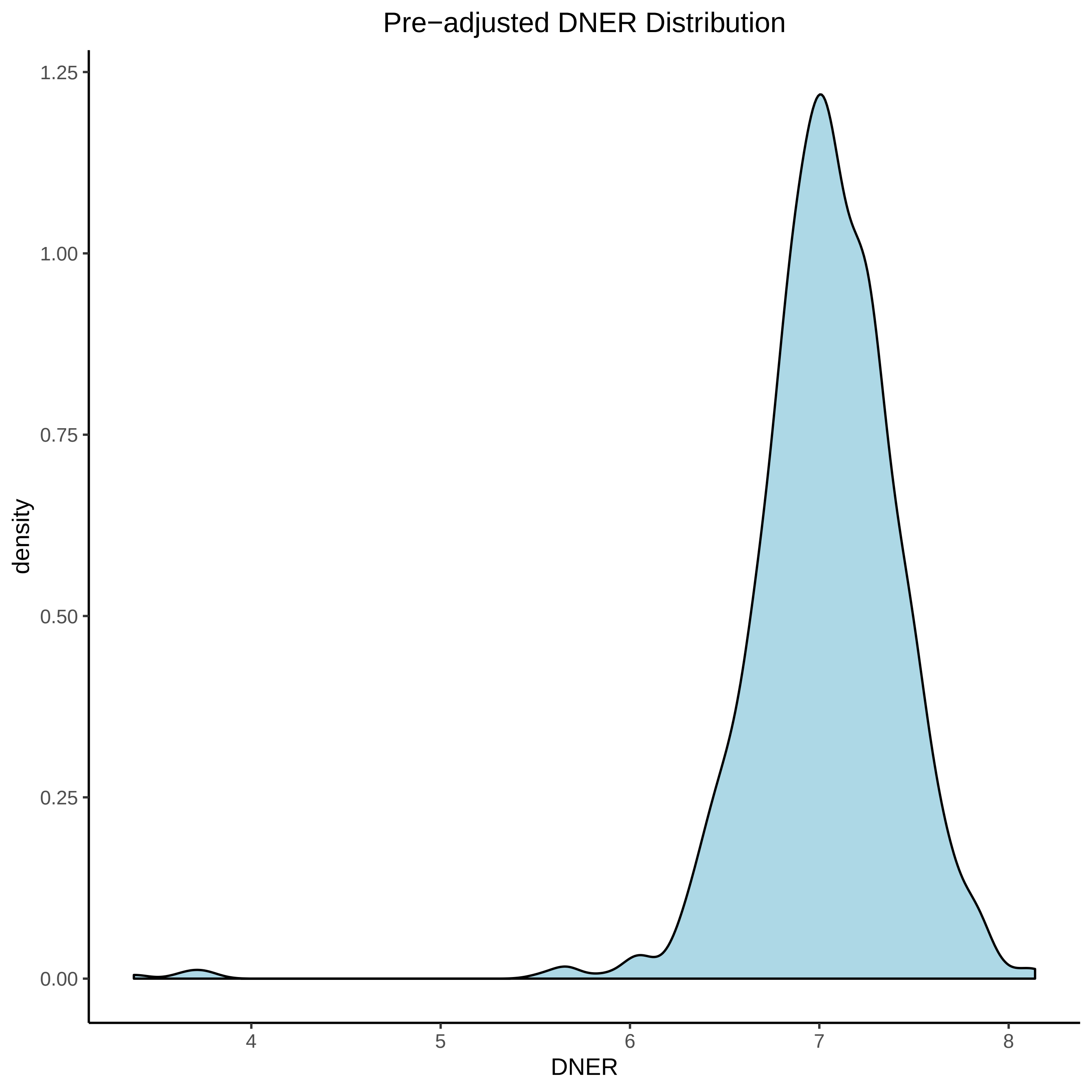

Pre−adjusted DNER Distribution
1.25
1.00
0.75
0.50
0.25
0.00
density
4
5
6
7
8
DNER

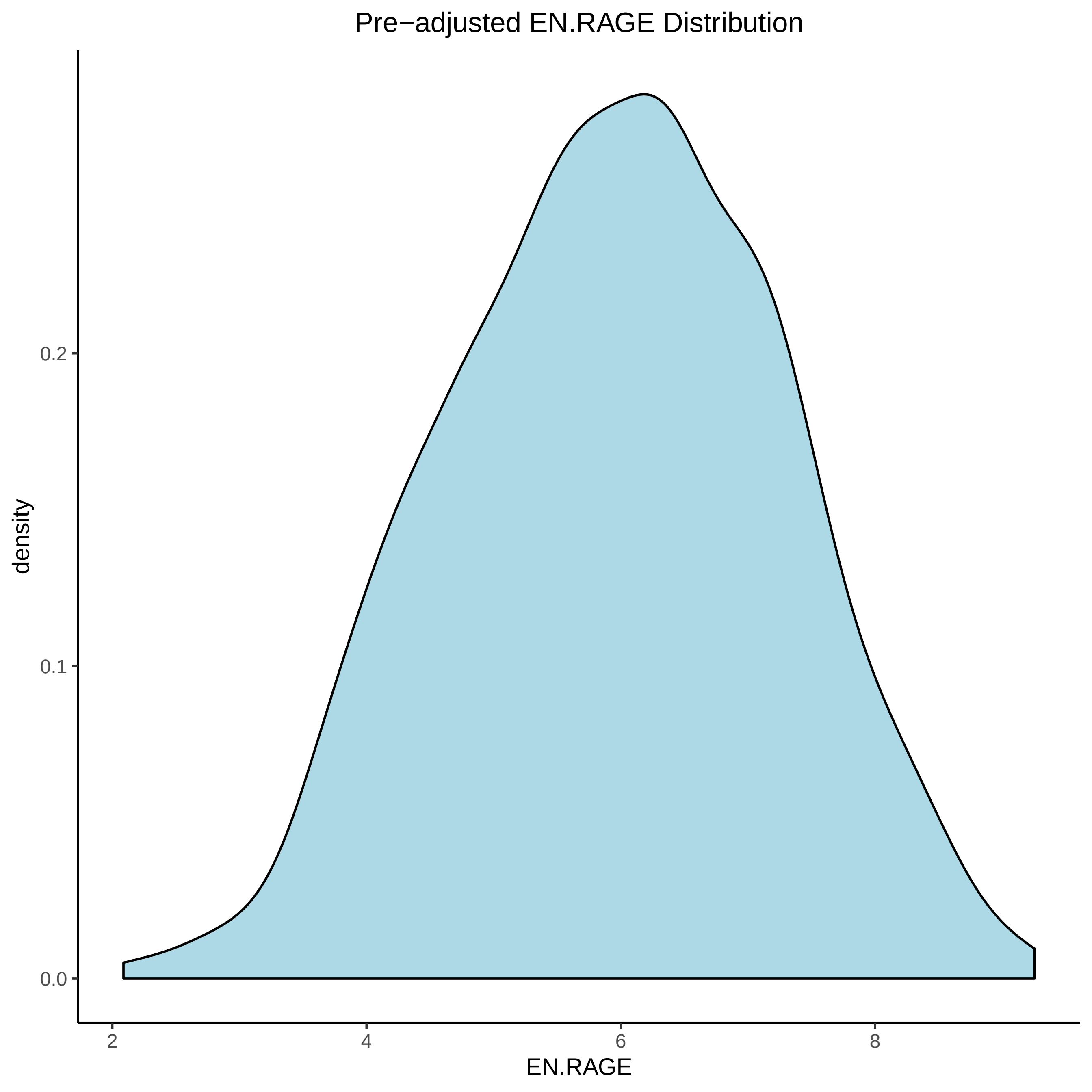

Pre−adjusted EN.RAGE Distribution
density
0.2
0.1
0.0
2
4
6
8
EN.RAGE

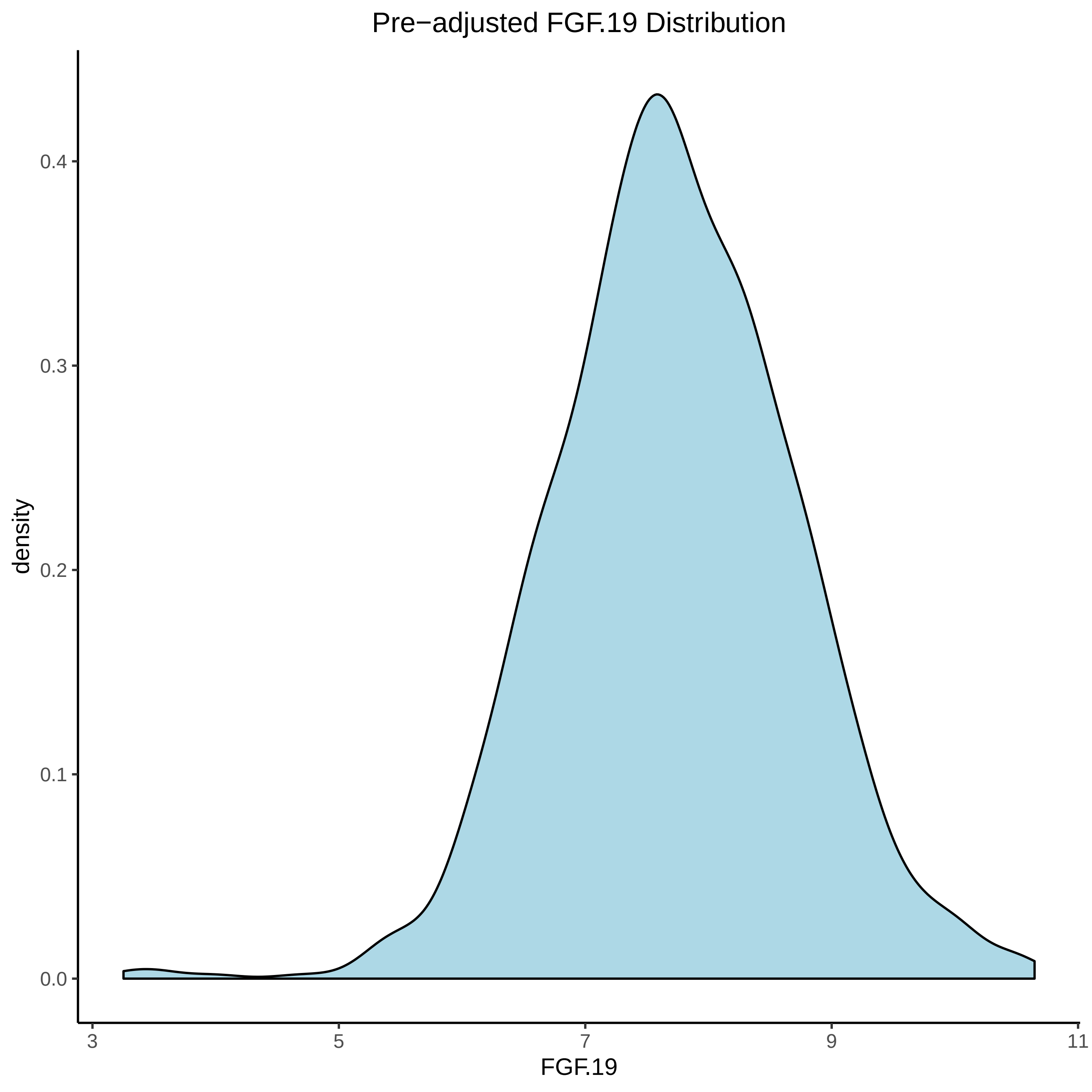

Pre−adjusted FGF.19 Distribution
density
0.4
0.3
0.2
0.1
0.0
3
5
7
9
11
FGF.19

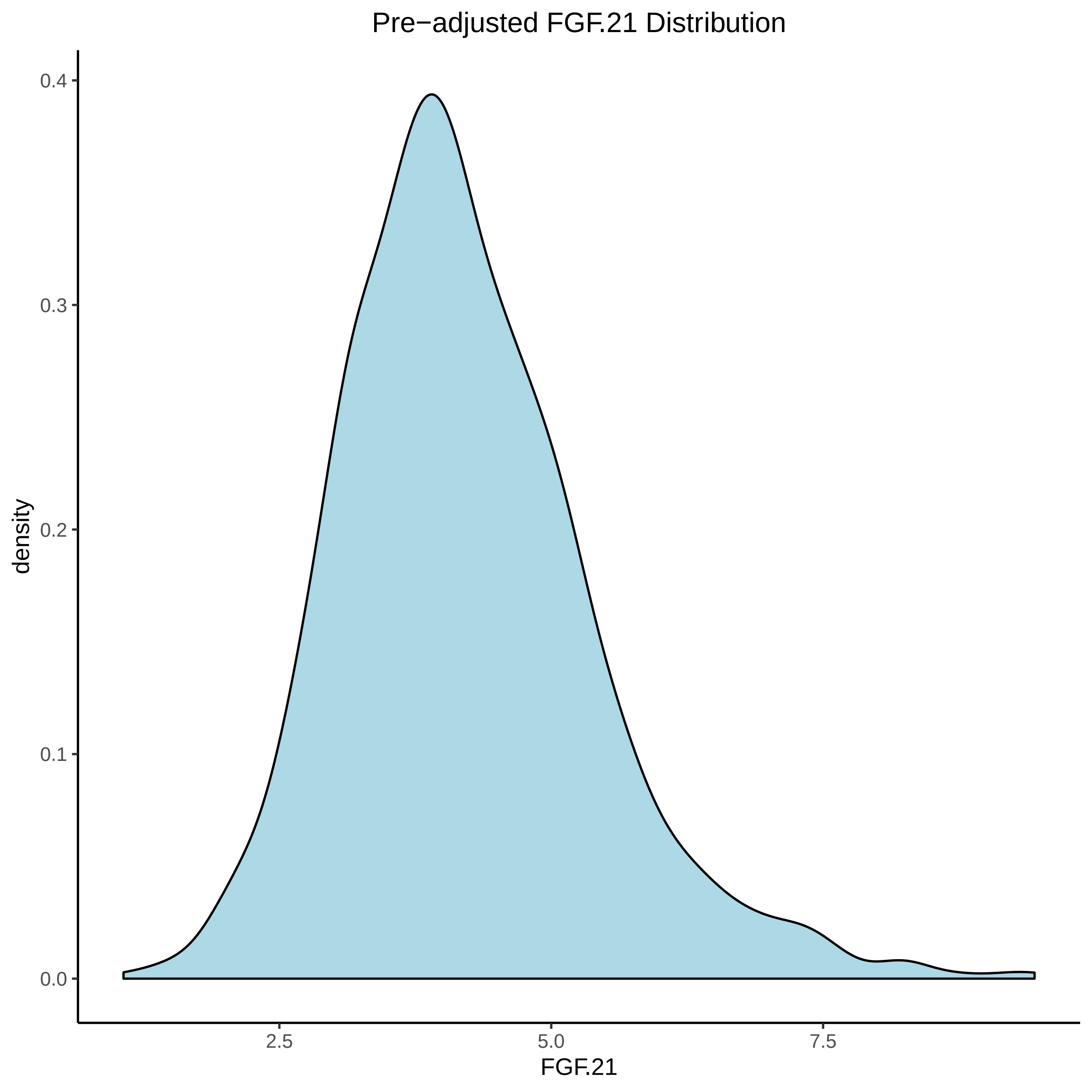
Pre−adjusted FGF.21 Distribution
density
0.4
0.3
0.2
0.1
0.0
2.5
5.0
7.5
FGF.21

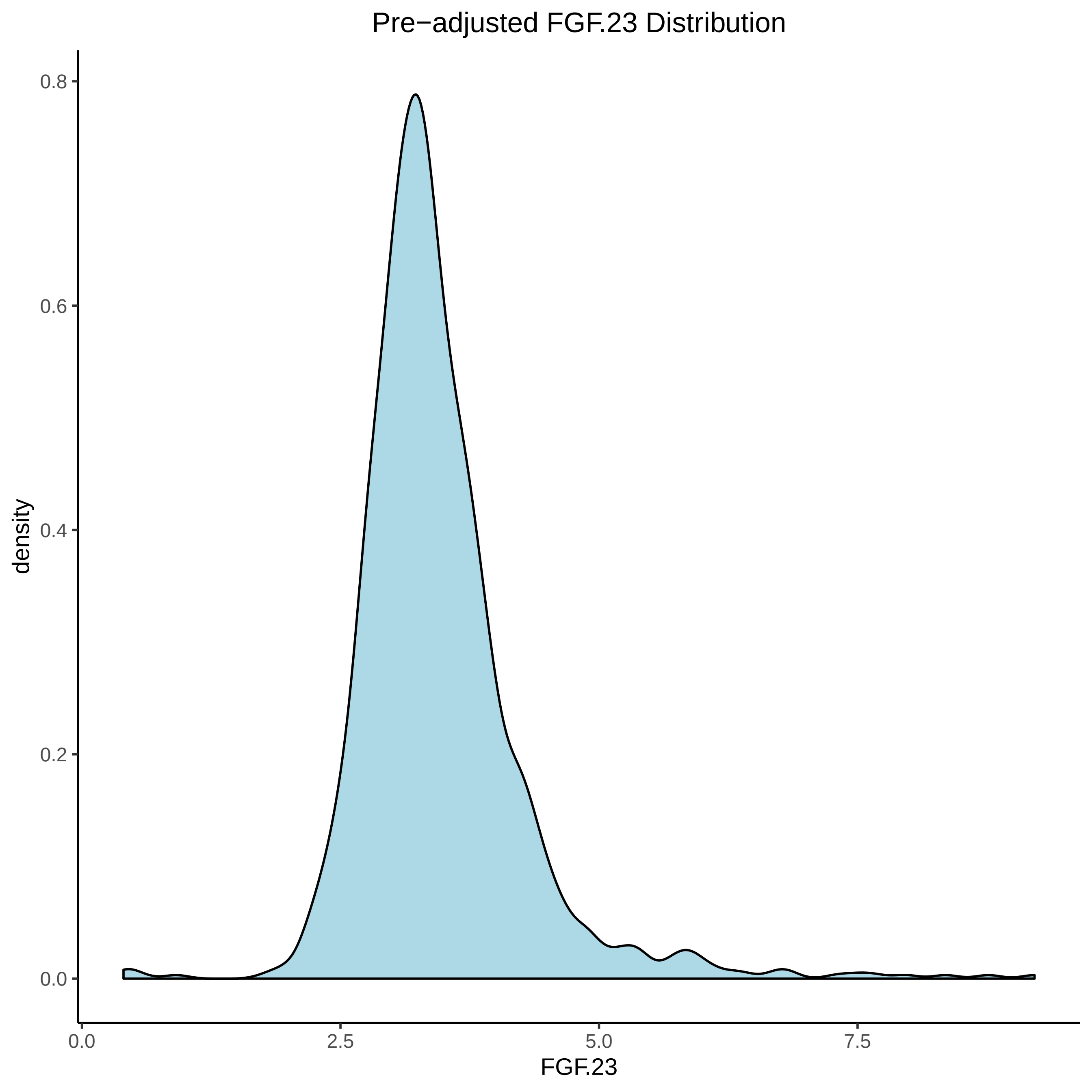

Pre−adjusted FGF.23 Distribution
0.8
0.6
0.4
0.2
0.0
density
0.0
2.5
5.0
7.5
FGF.23

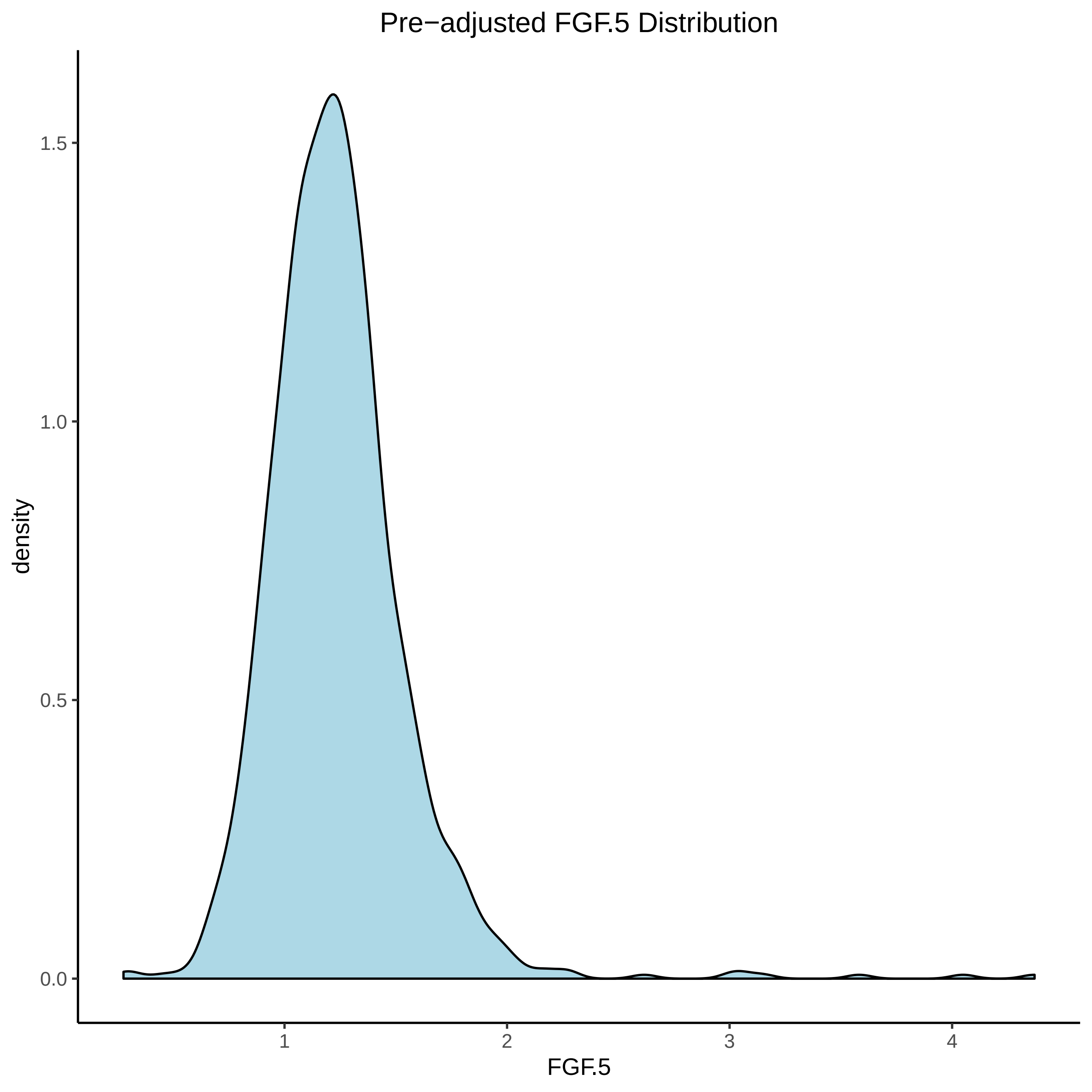
Pre−adjusted FGF.5 Distribution
density
1.5
1.0
0.5
0.0
1
2
3
4
FGF.5

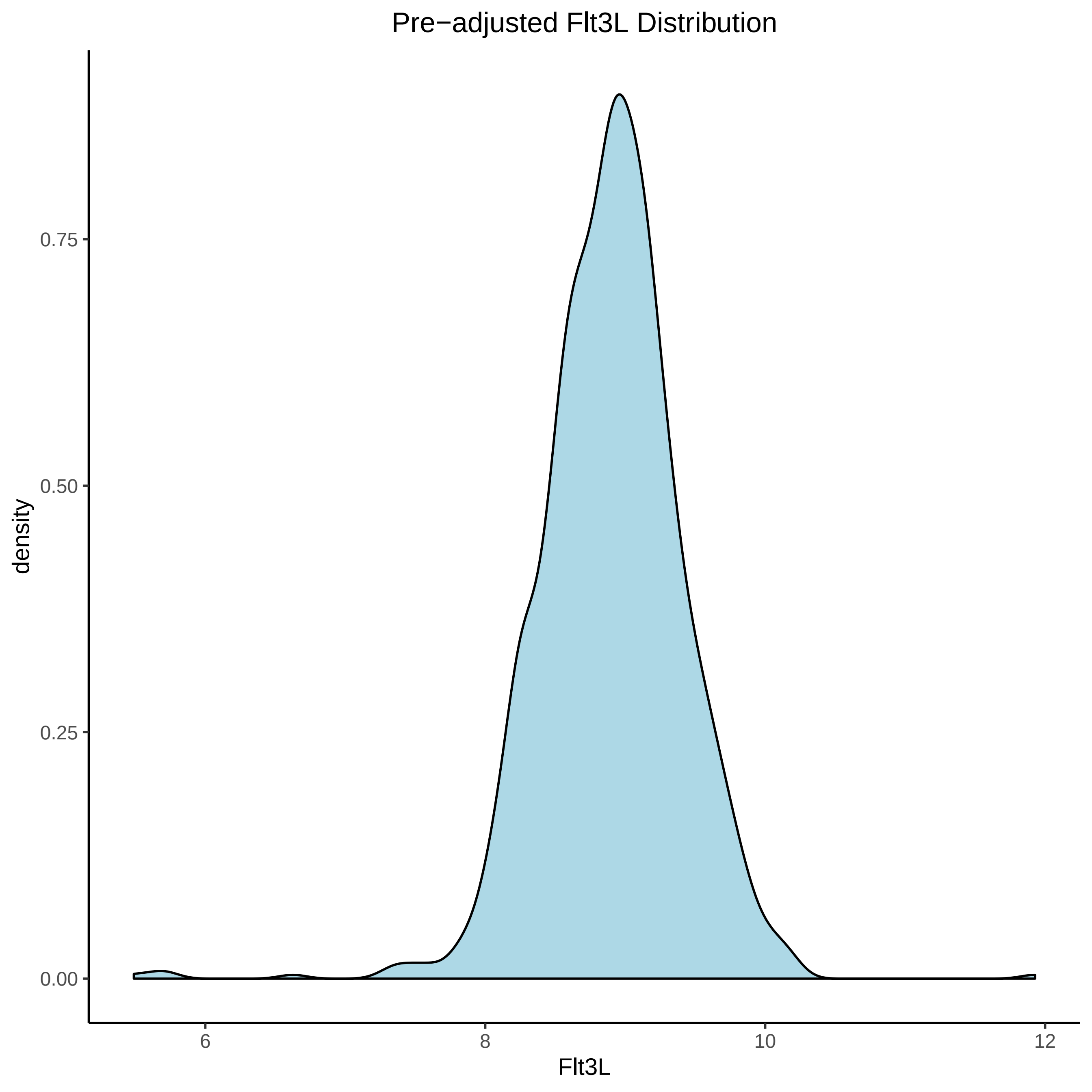

Pre−adjusted Flt3L Distribution
0.75
0.50
0.25
0.00
density
6
8
10
12
Flt3L

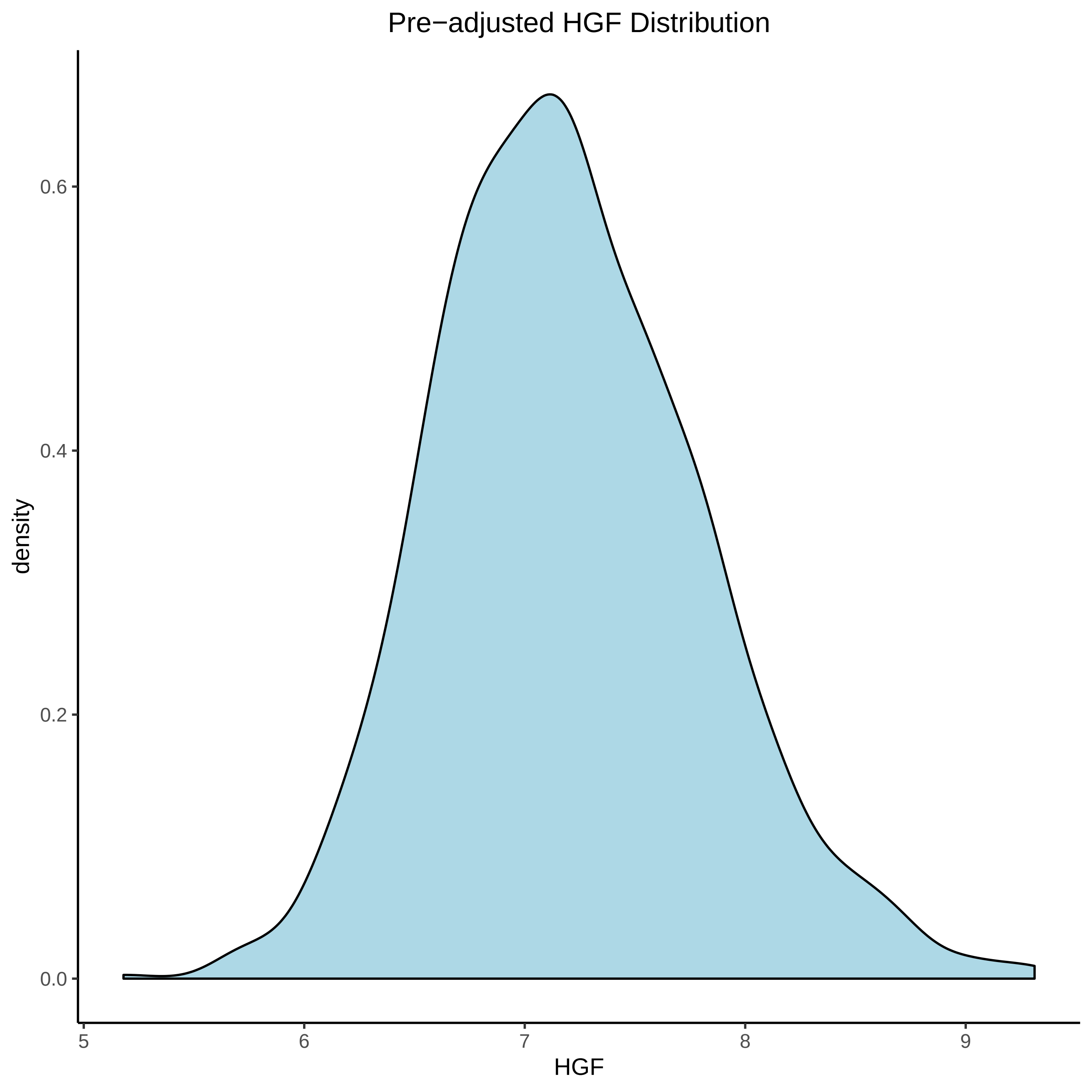

Pre−adjusted HGF Distribution
density
0.0
0.2
0.4
0.6
5
6
7
8
9
HGF

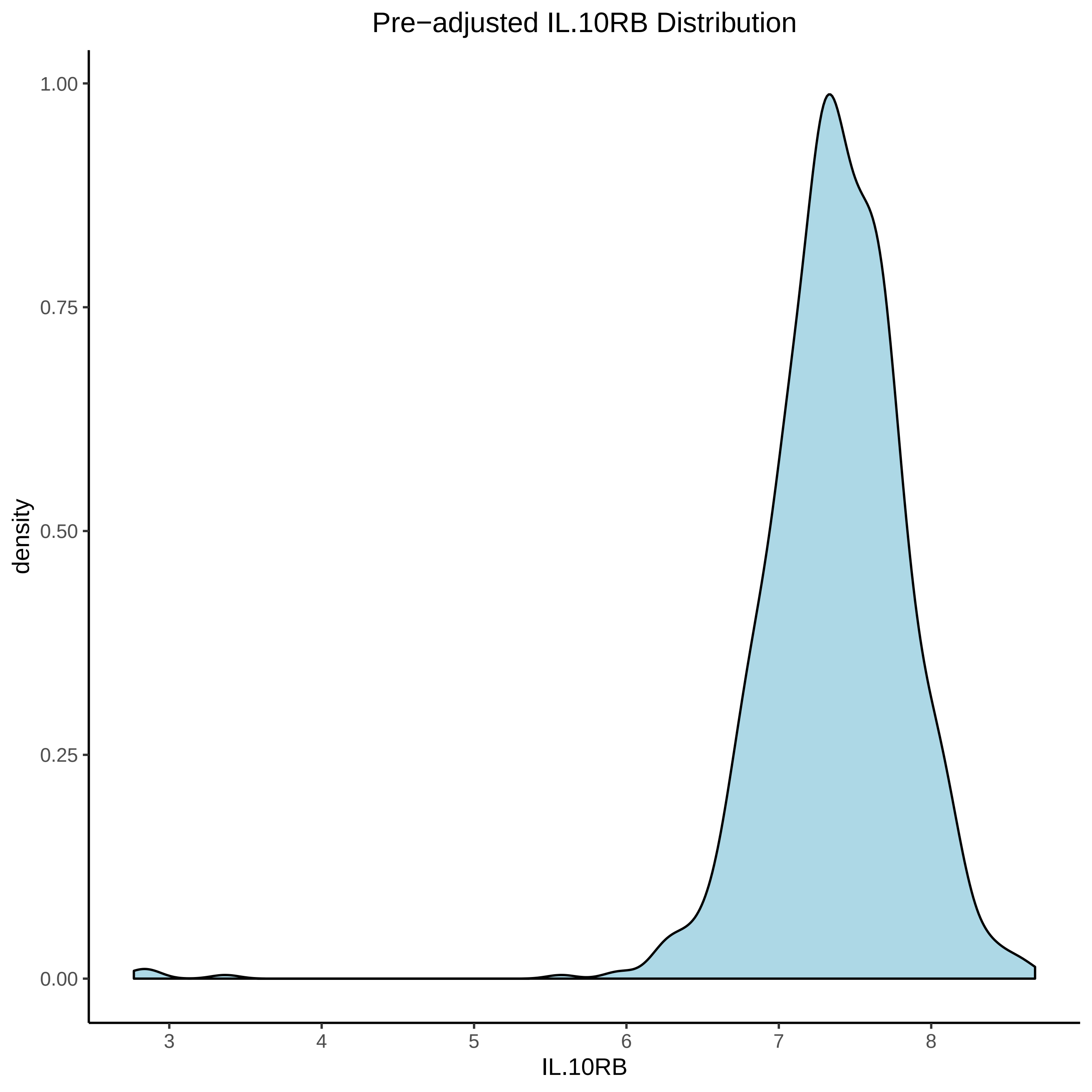

Pre−adjusted IL.10RB Distribution
1.00
0.75
0.50
0.25
0.00
density
3
4
5
6
7
8
IL.10RB

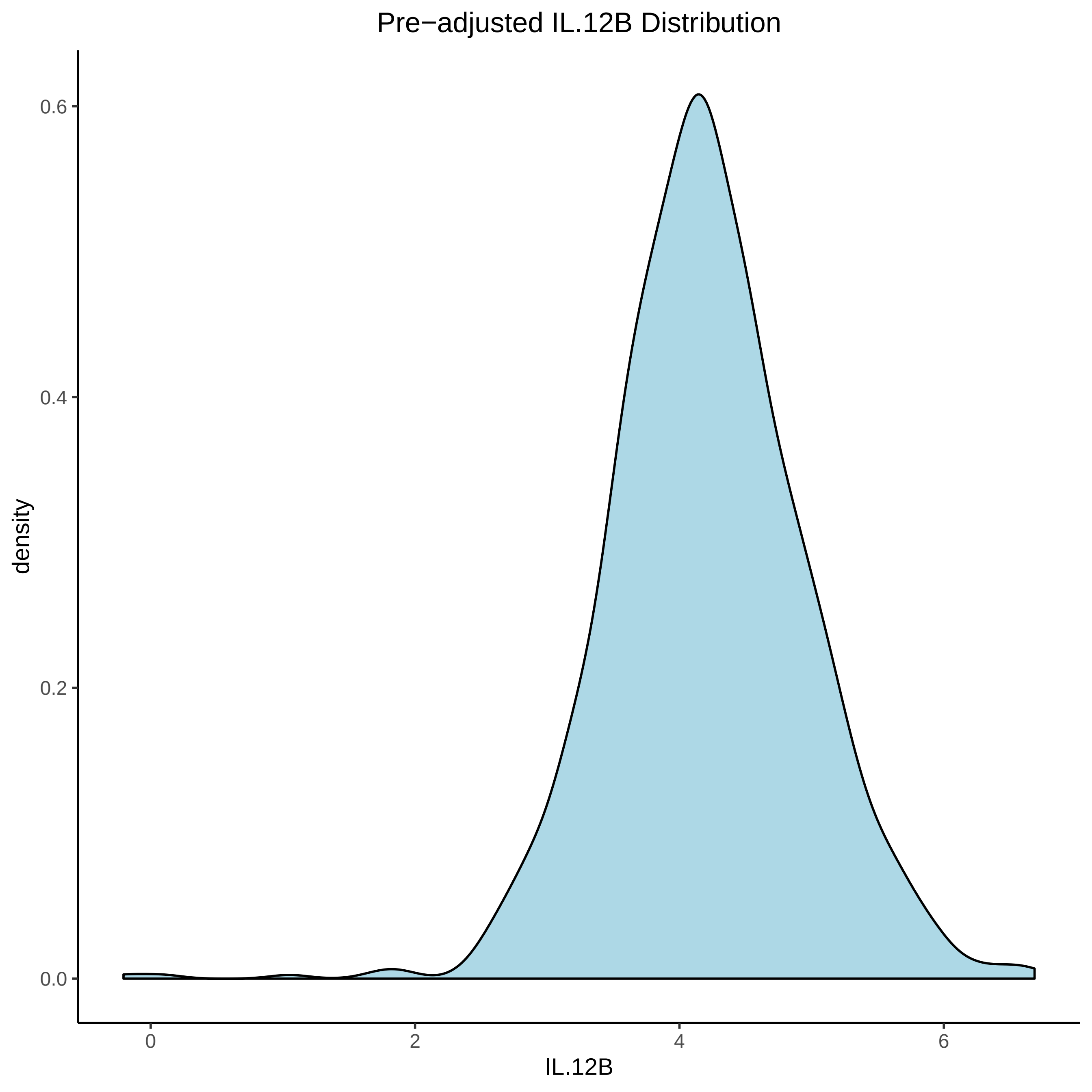
Pre−adjusted IL.12B Distribution
density
0.6
0.4
0.2
0.0
0
2
4
6
IL.12B

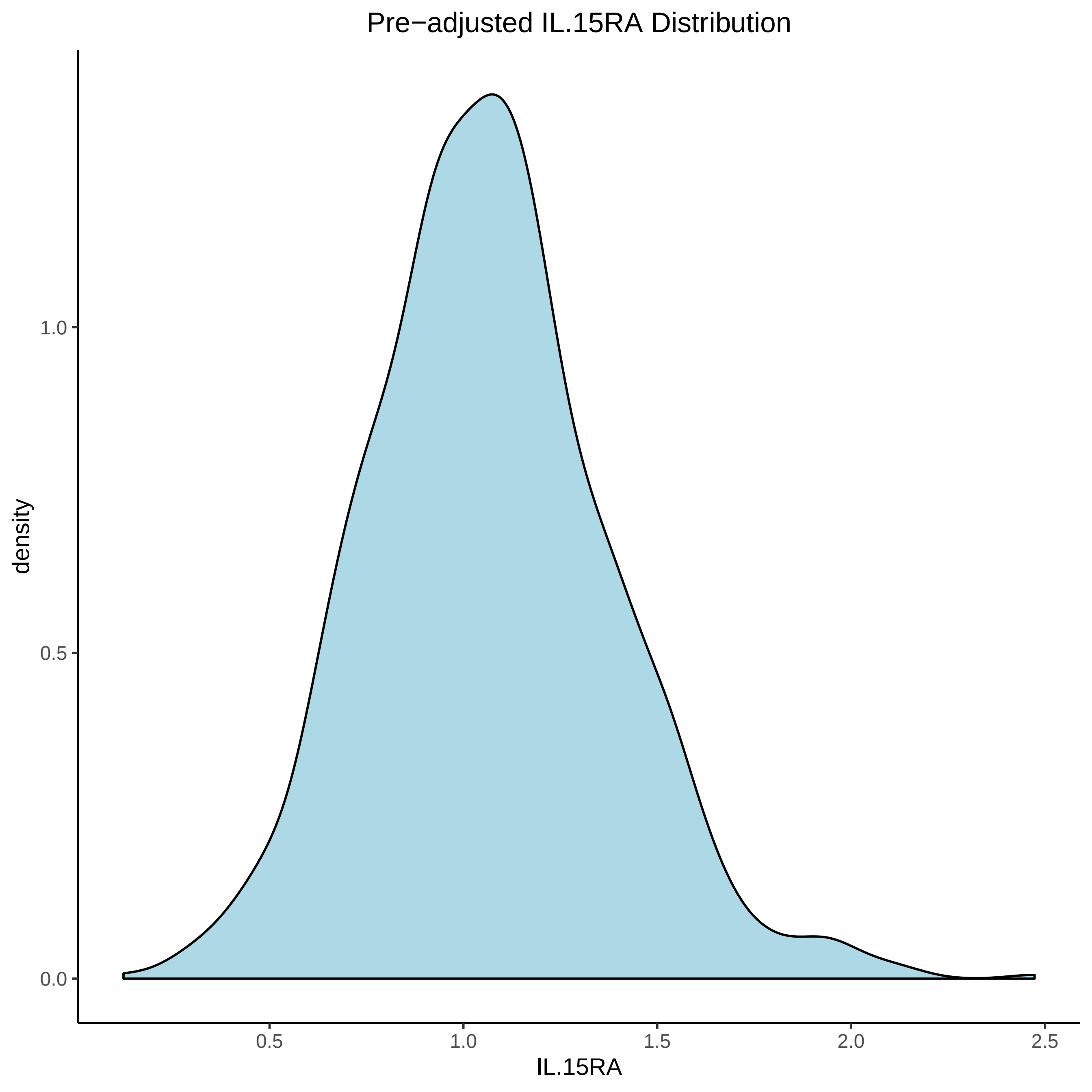

Pre−adjusted IL.15RA Distribution
density
1.0
0.5
0.0
0.5
1.0
1.5
2.0
2.5
IL.15RA

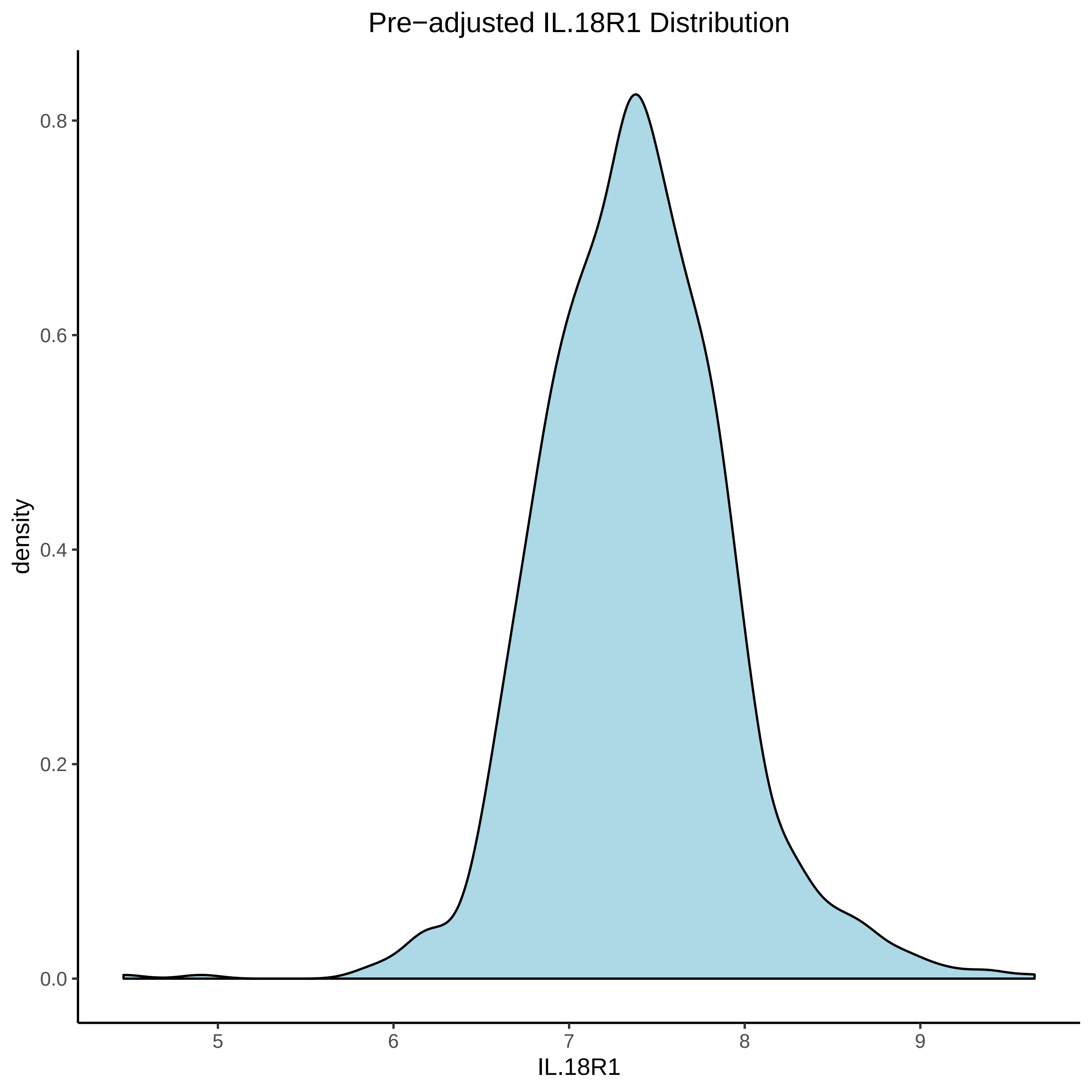

Pre−adjusted IL.18R1 Distribution
density
0.0
0.2
0.4
0.6
0.8
5
6
7
8
9
IL.18R1

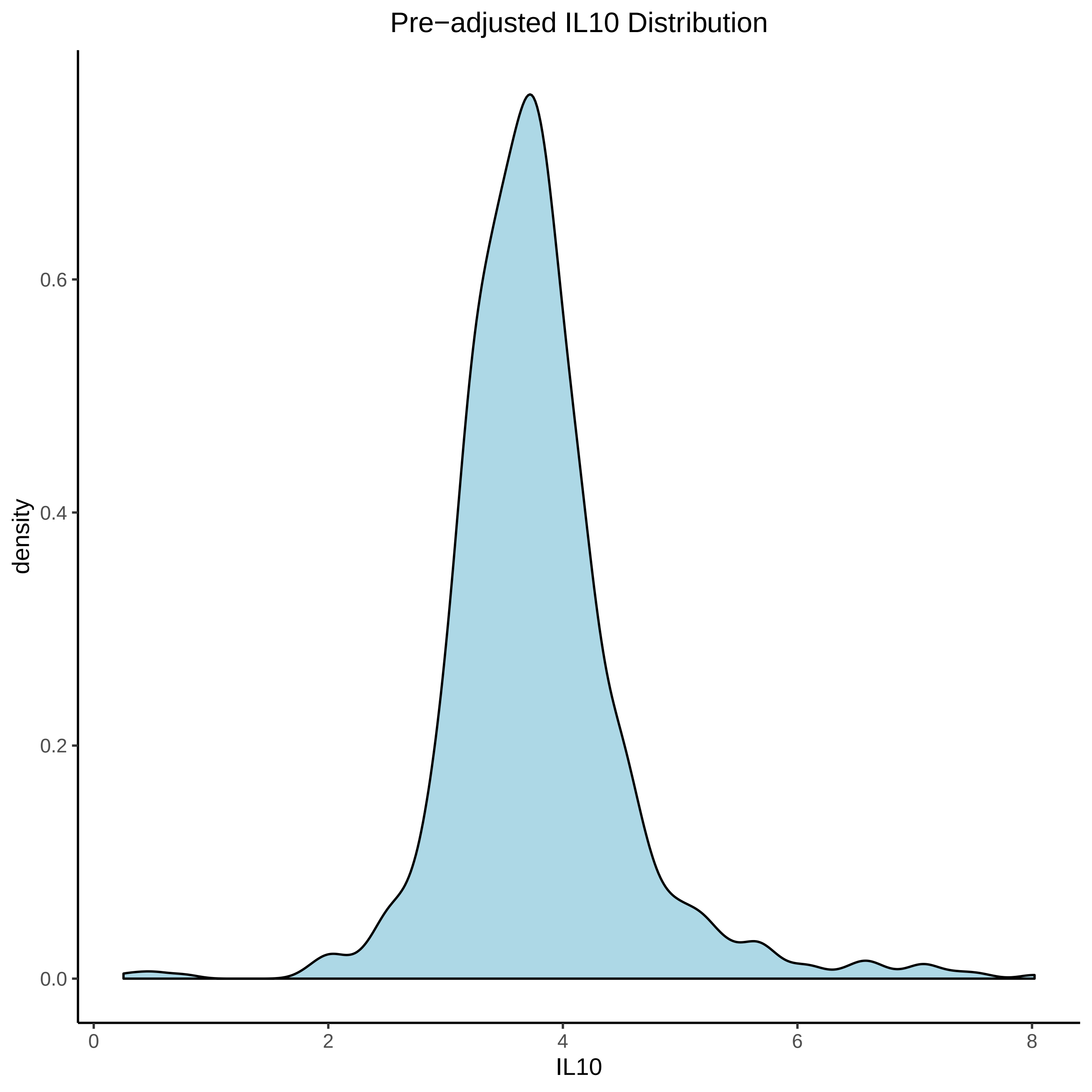
Pre−adjusted IL10 Distribution
density
0.0
0.2
0.4
0.6
0
2
4
6
8
IL10

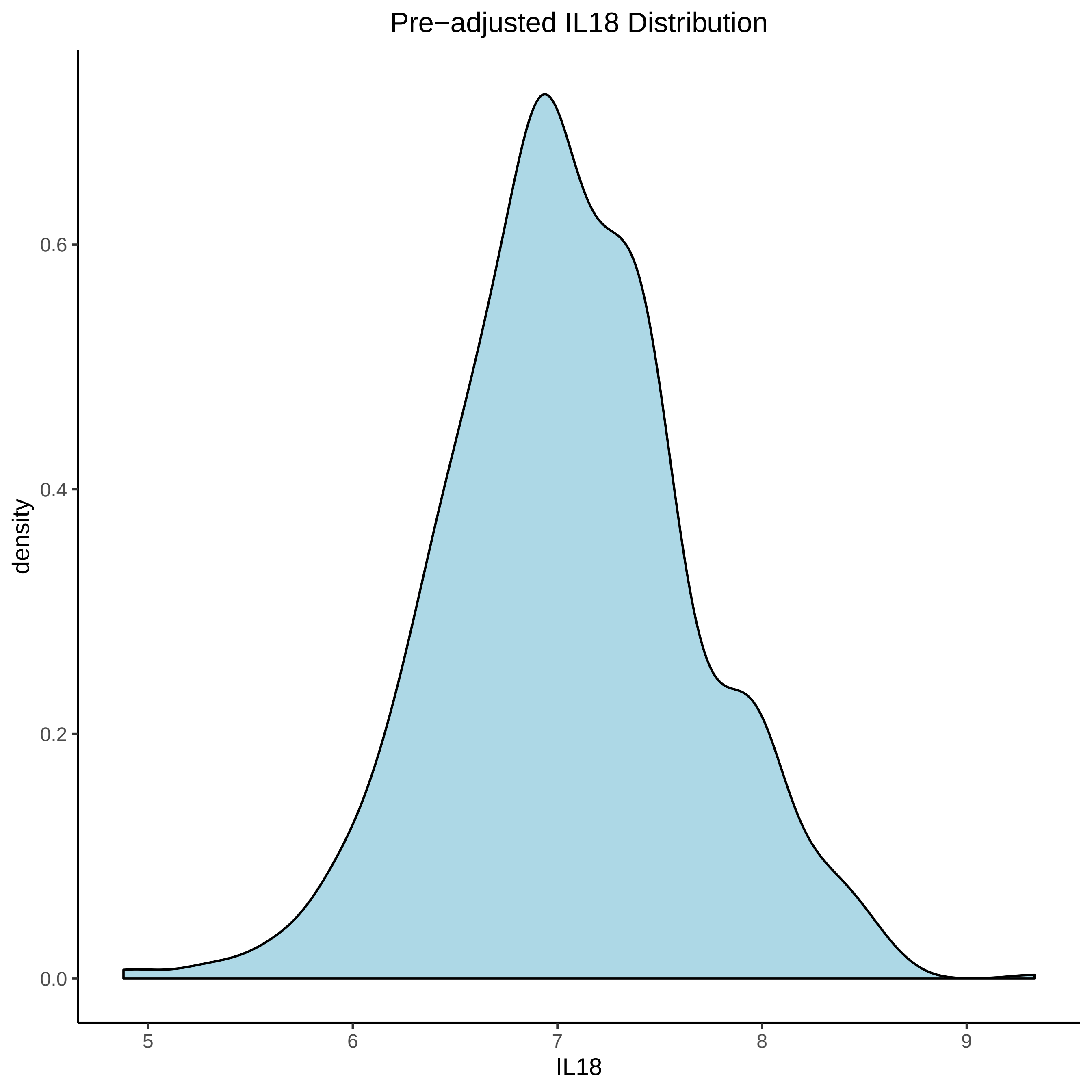

Pre−adjusted IL18 Distribution
density
0.0
0.2
0.4
0.6
5
6
7
8
9
IL18

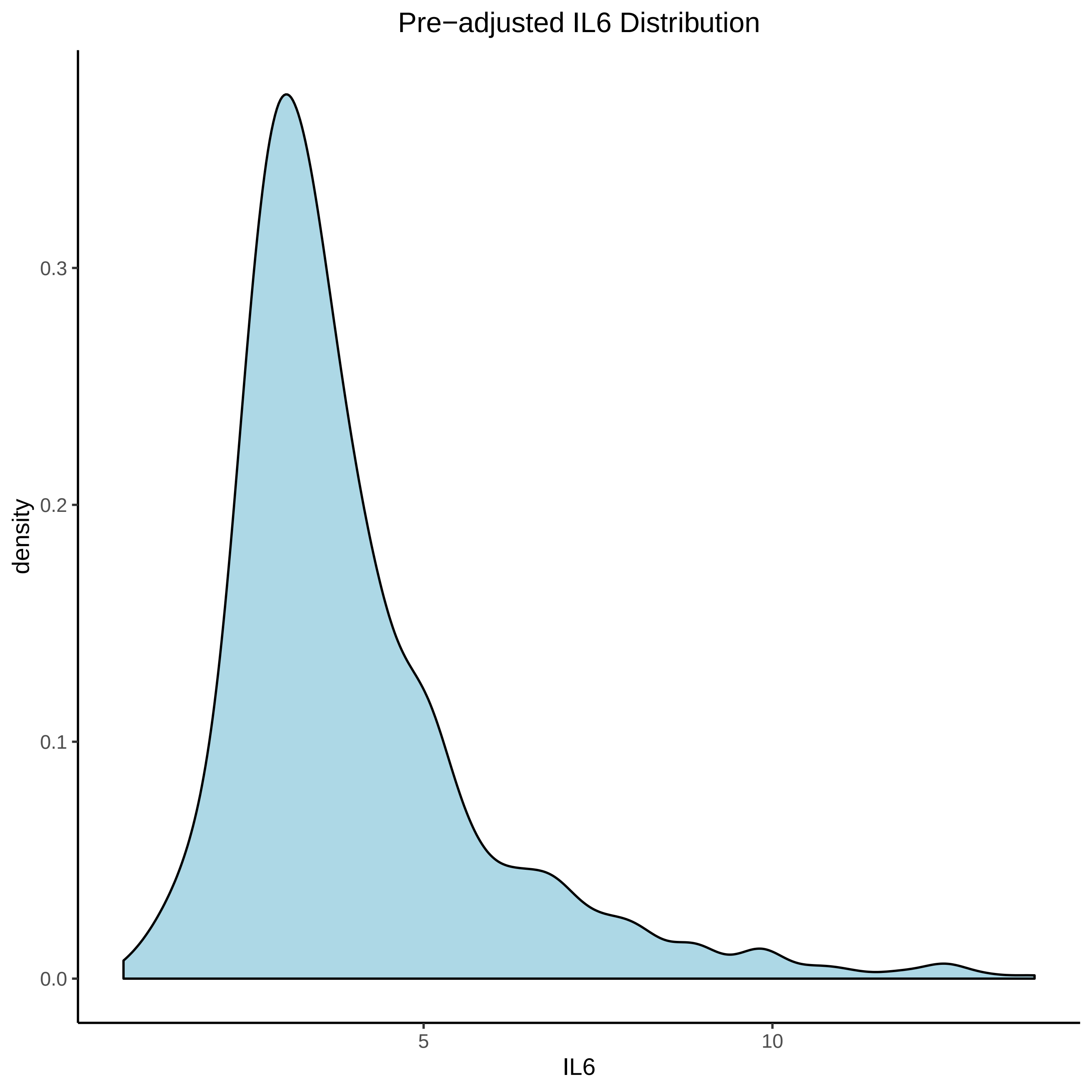

Pre−adjusted IL6 Distribution
density
0.3
0.2
0.1
0.0
5
10
IL6

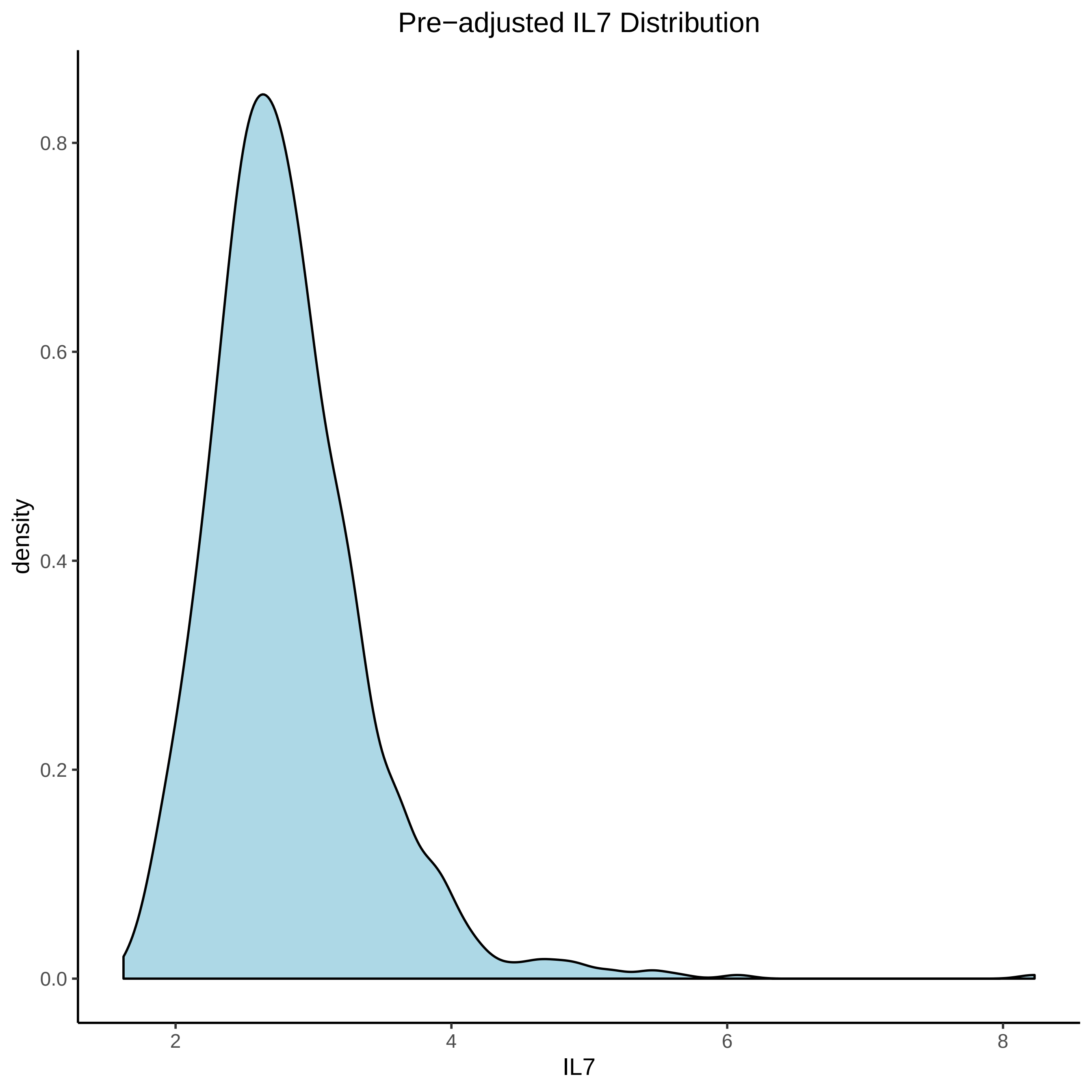
Pre−adjusted IL7 Distribution
density
0.0
0.2
0.4
0.6
0.8
2
4
6
8
IL7

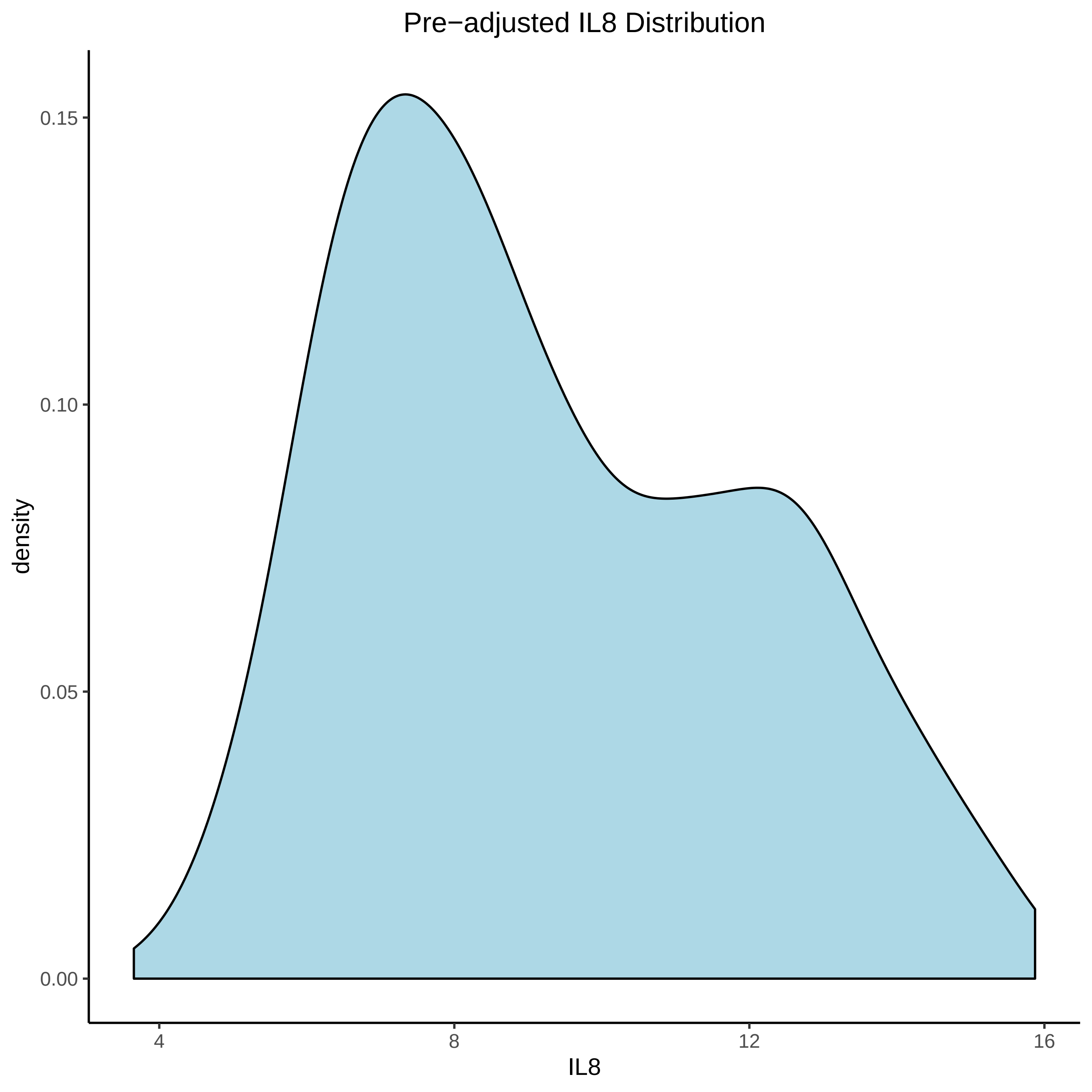

Pre−adjusted IL8 Distribution
density
0.15
0.10
0.05
0.00
4
8
12
16
IL8

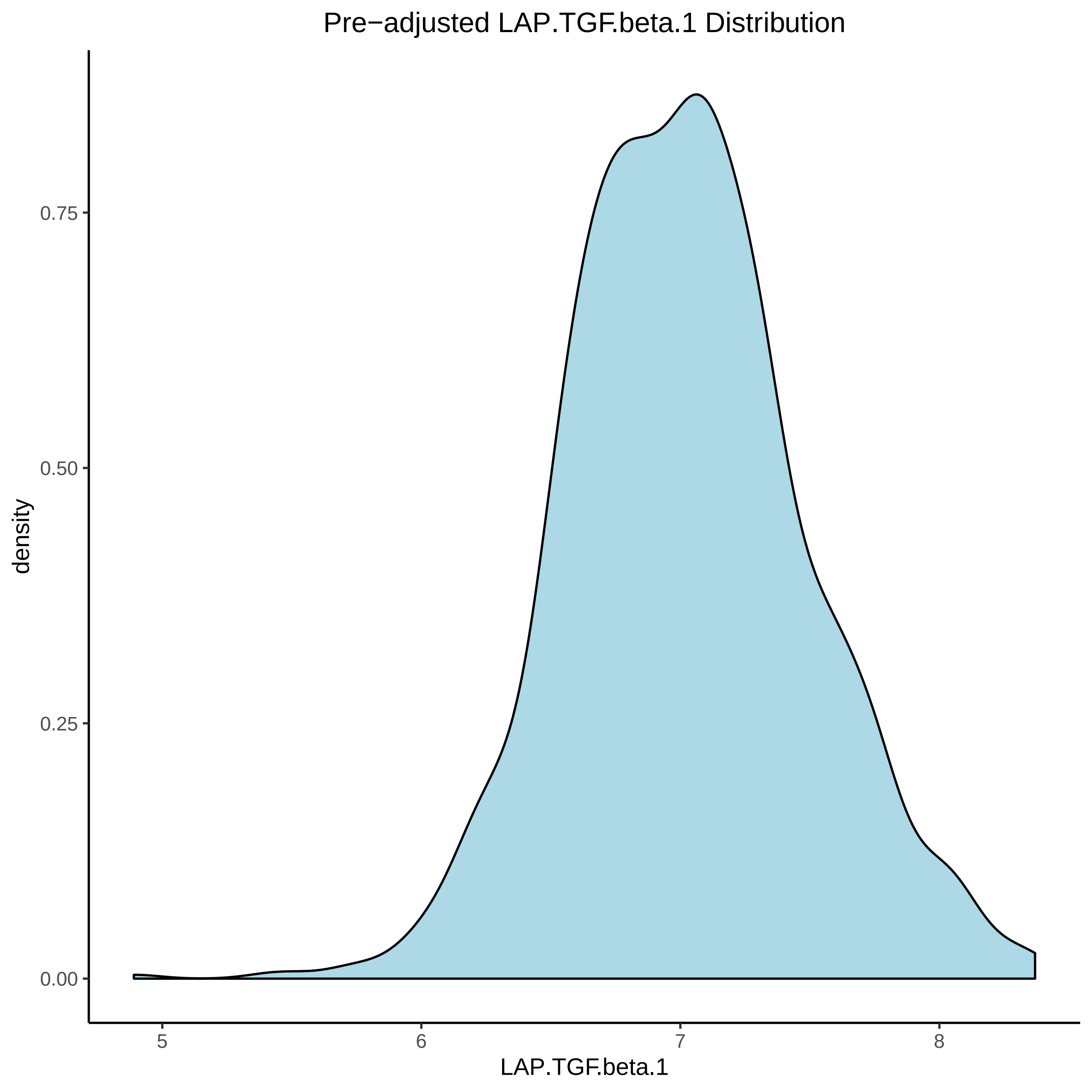
Pre−adjusted LAP.TGF.beta.1 Distribution
density
0.00
0.25
0.50
0.75
5
6
7
8
LAP.TGF.beta.1

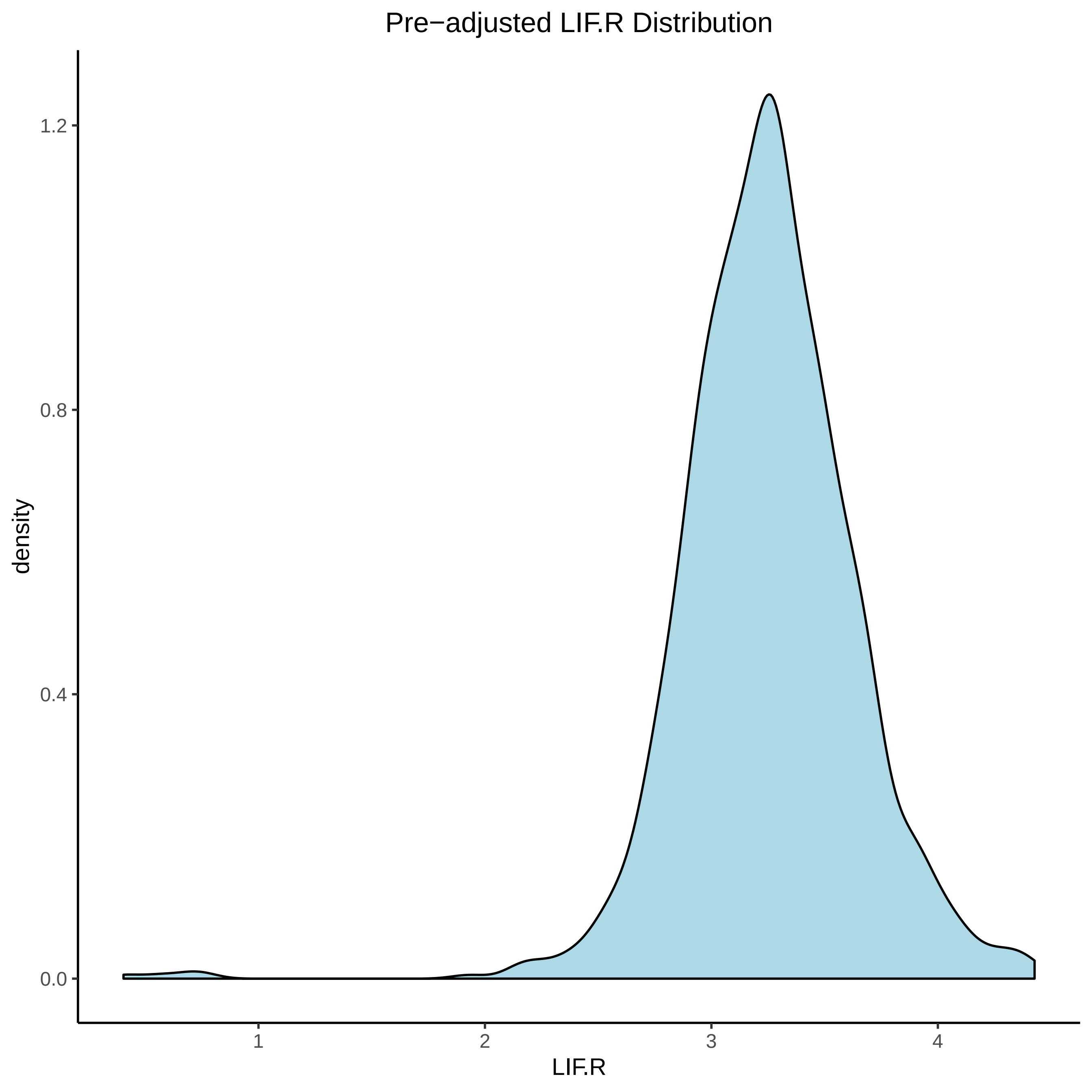
Pre−adjusted LIF.R Distribution
density
1.2
0.8
0.4
0.0
1
2
3
4
LIF.R

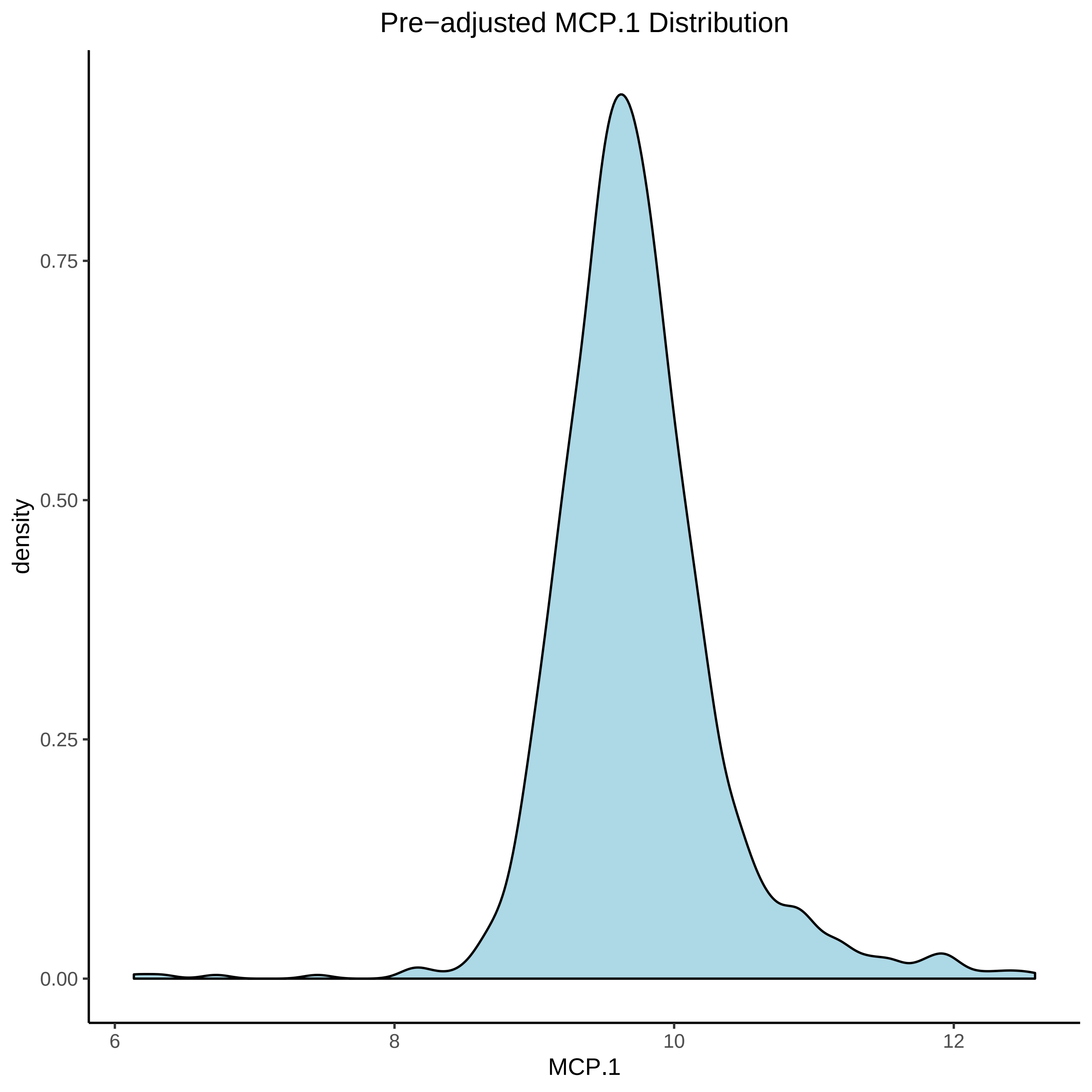

Pre−adjusted MCP.1 Distribution
density
0.00
0.25
0.50
0.75
6
8
10
12
MCP.1

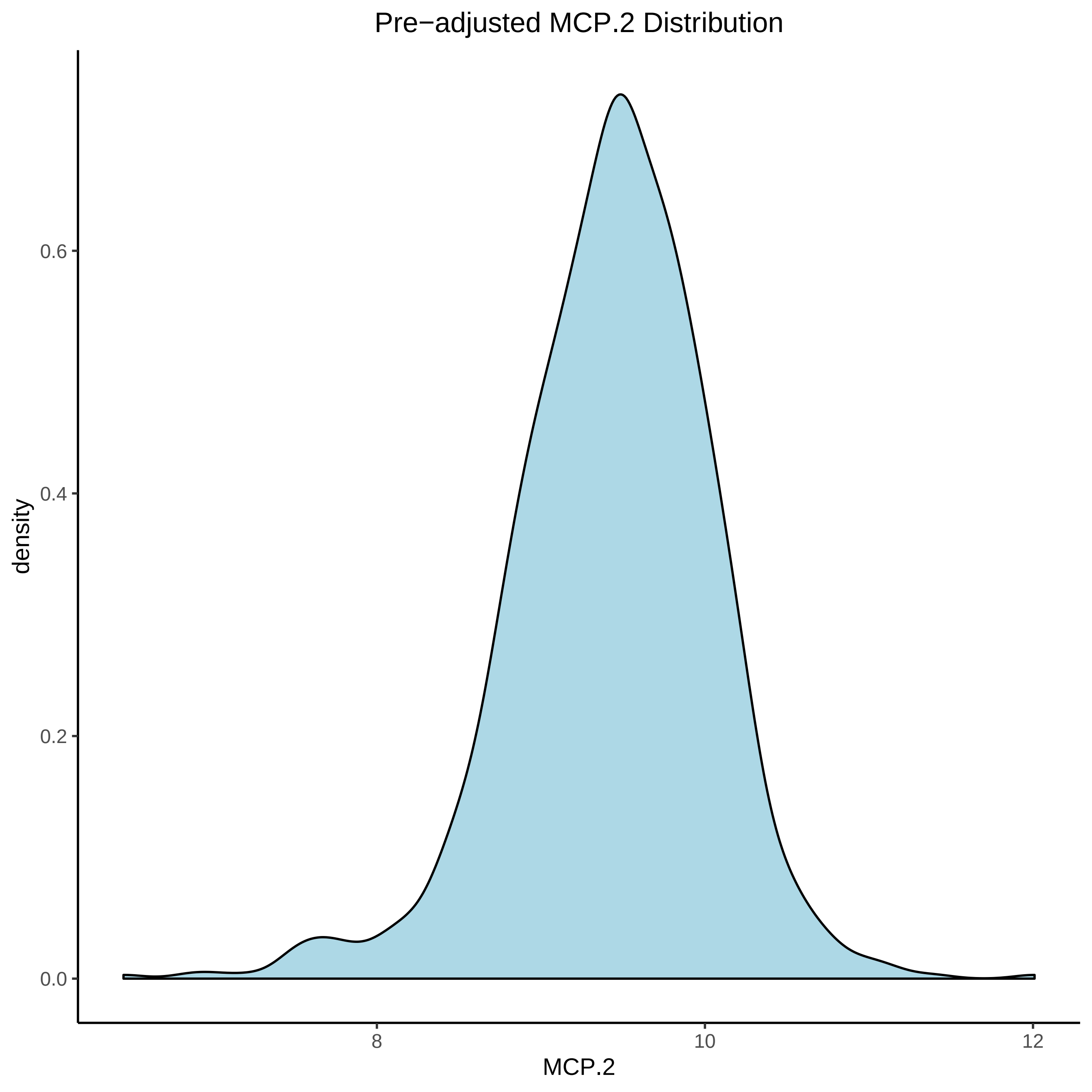

Pre−adjusted MCP.2 Distribution
density
0.0
0.2
0.4
0.6
8
10
12
MCP.2

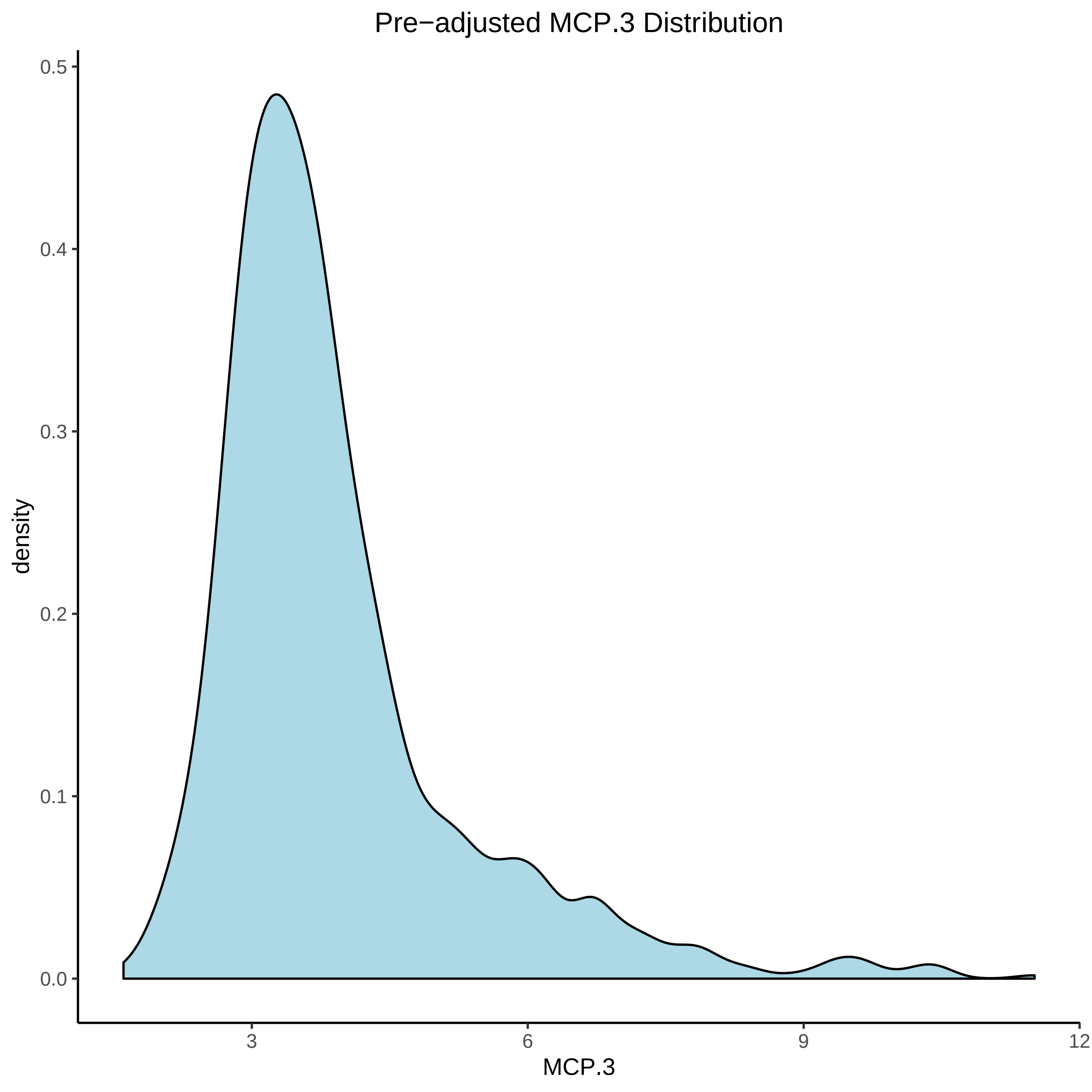

Pre−adjusted MCP.3 Distribution
density
0.5
0.4
0.3
0.2
0.1
0.0
3
6
9
12
MCP.3

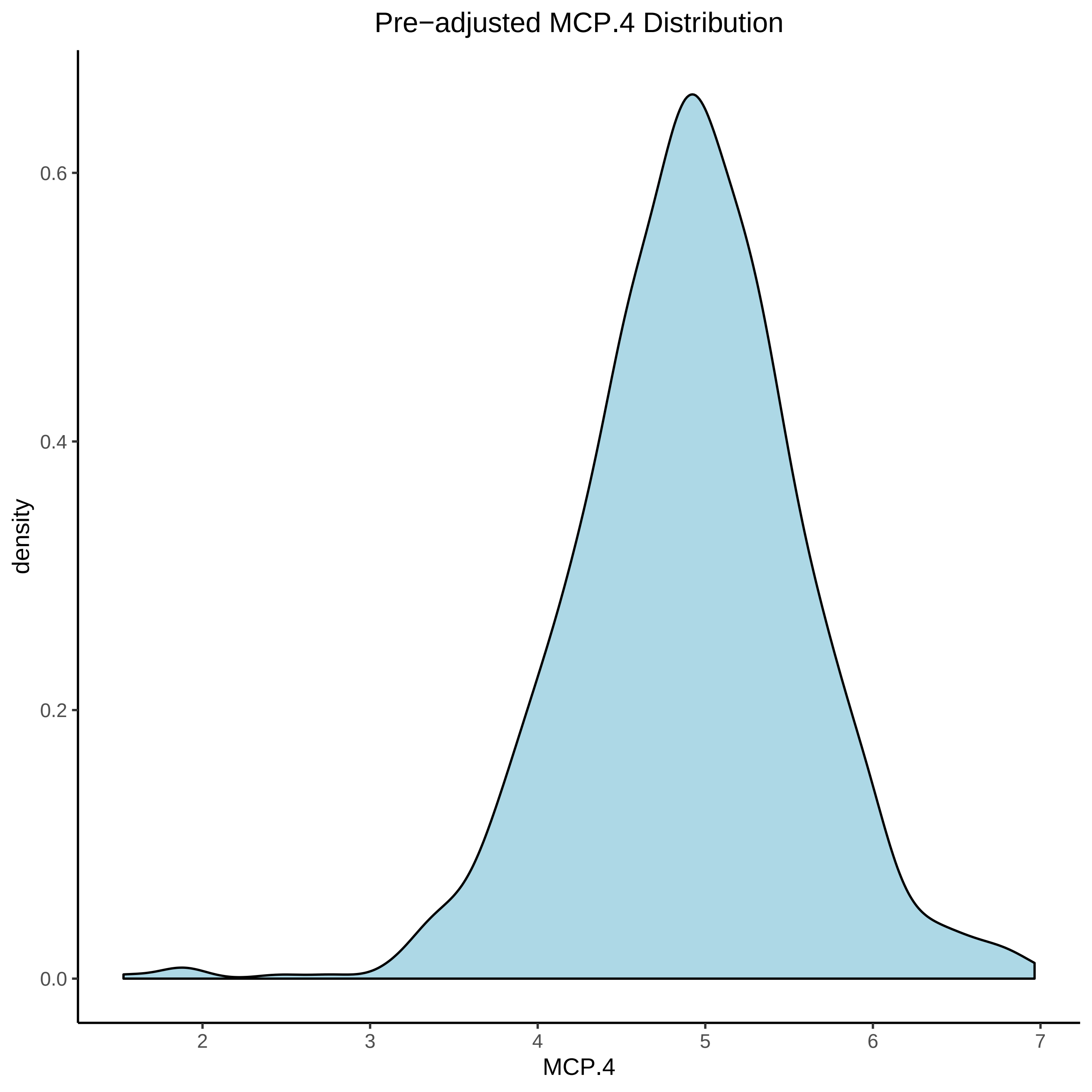

Pre−adjusted MCP.4 Distribution
density
0.0
0.2
0.4
0.6
2
3
4
5
6
7
MCP.4

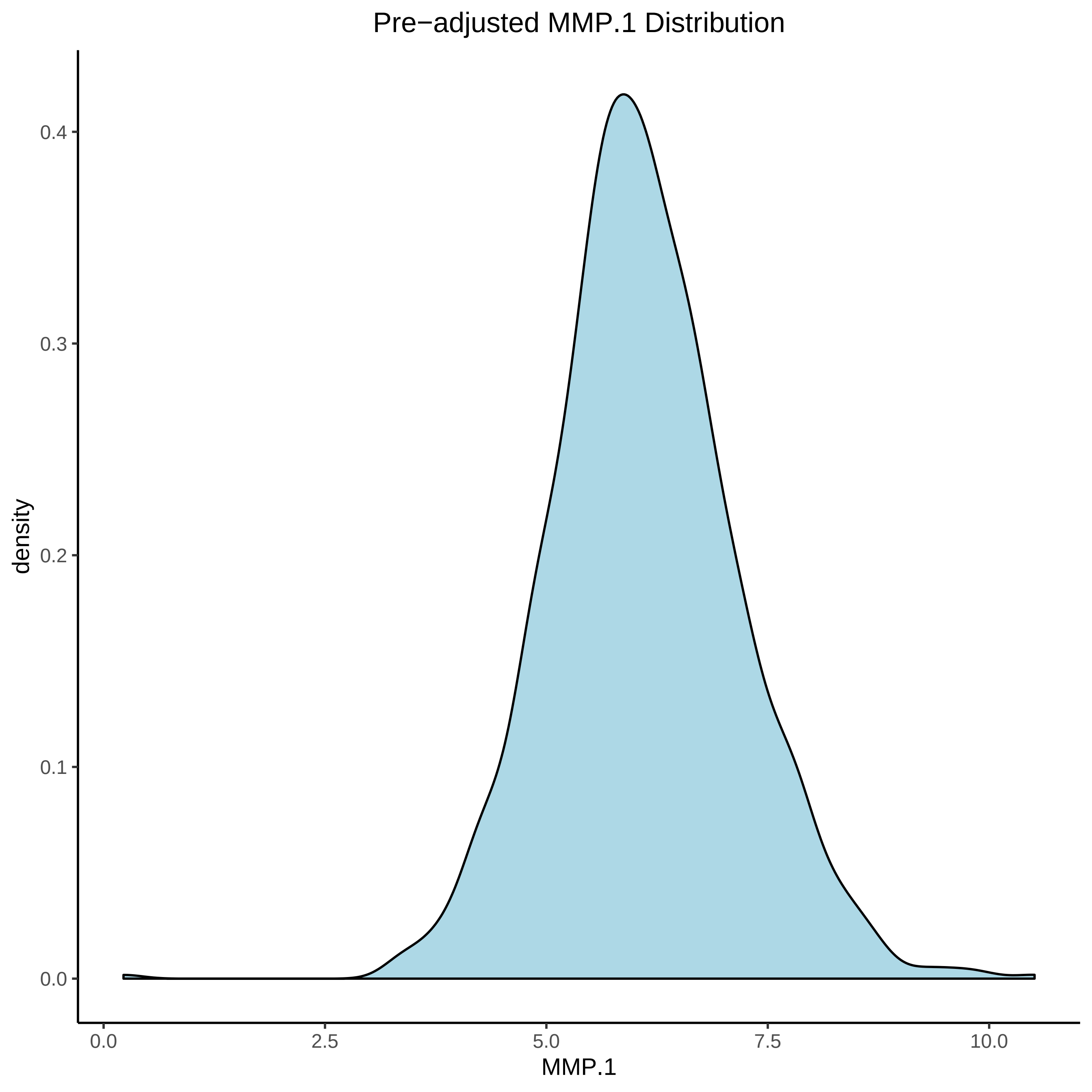
Pre−adjusted MMP.1 Distribution
density
0.0
0.1
0.2
0.3
0.4
0.0
2.5
5.0
7.5
10.0
MMP.1

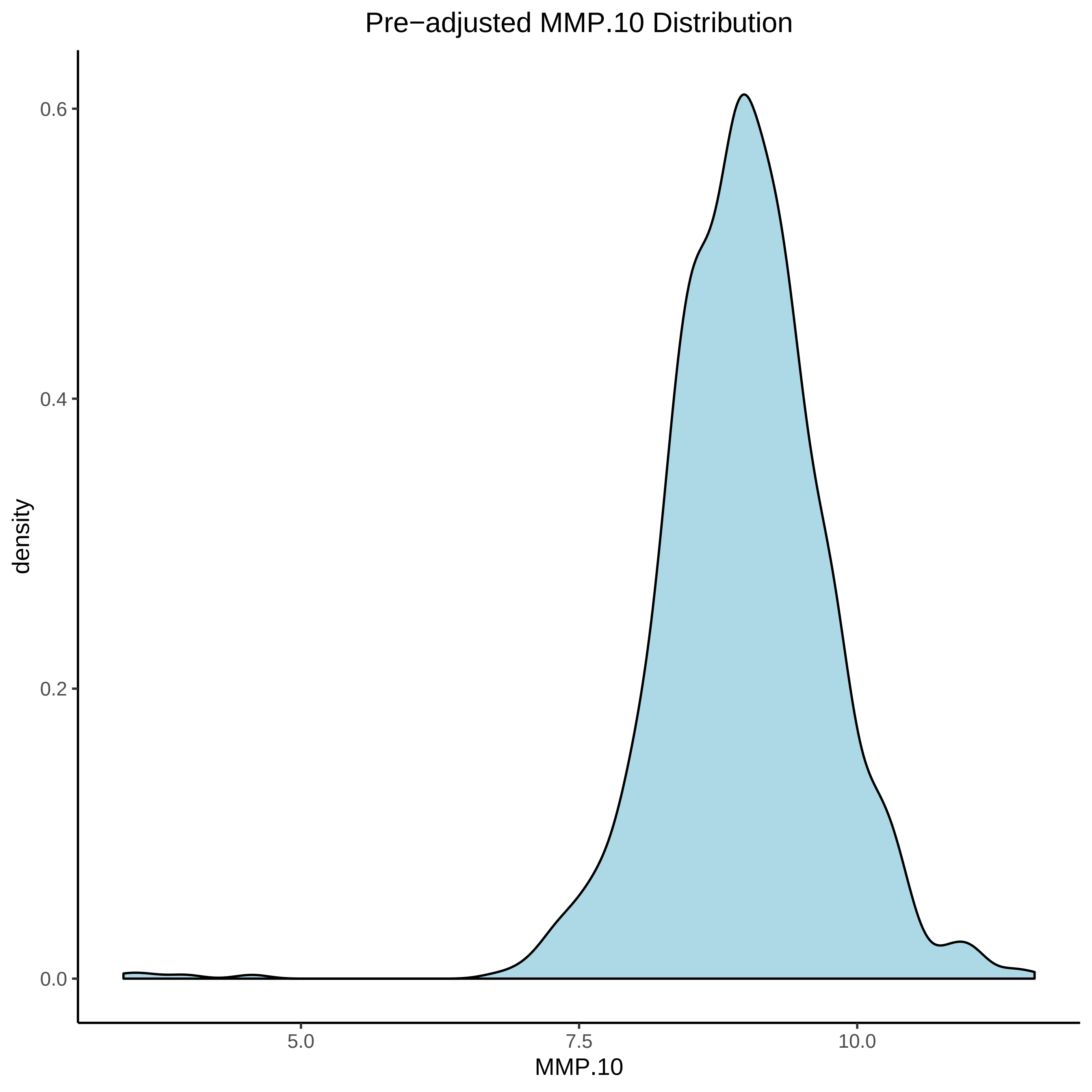

Pre−adjusted MMP.10 Distribution
0.6
0.4
0.2
0.0
density
5.0
7.5
10.0
MMP.10

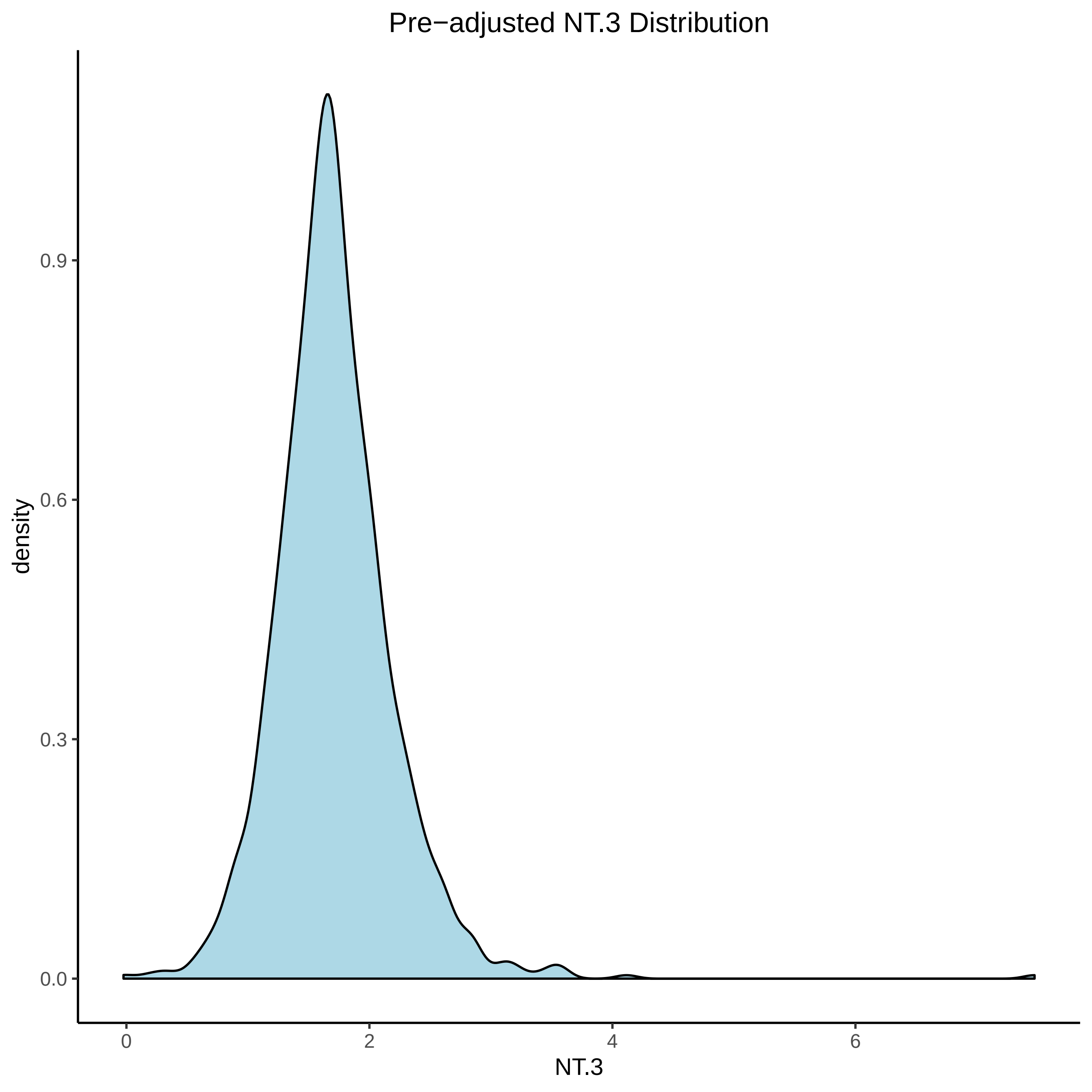
Pre−adjusted NT.3 Distribution
density
0.9
0.6
0.3
0.0
0
2
4
6
NT.3

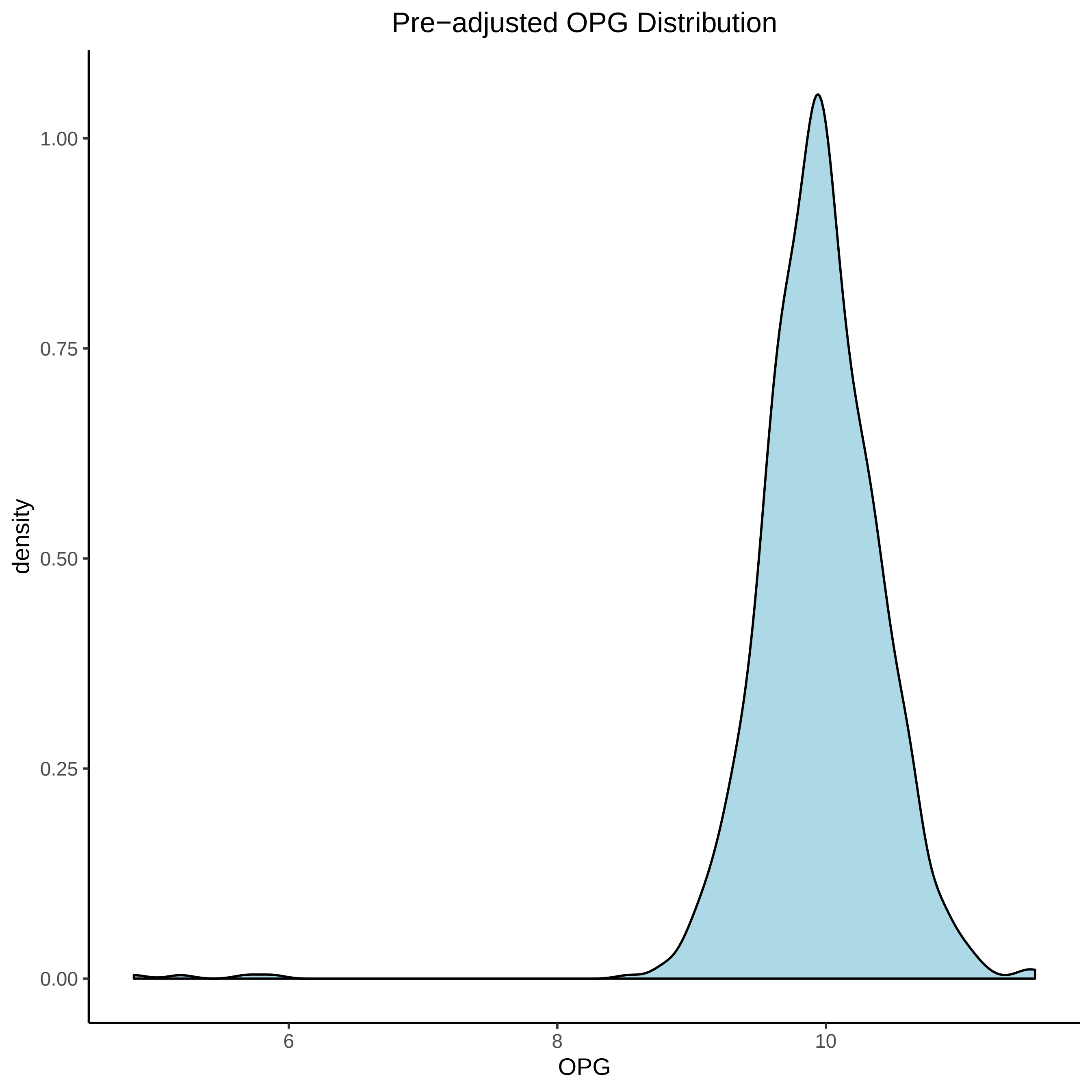
Pre−adjusted OPG Distribution
density
1.00
0.75
0.50
0.25
0.00
6
8
10
OPG

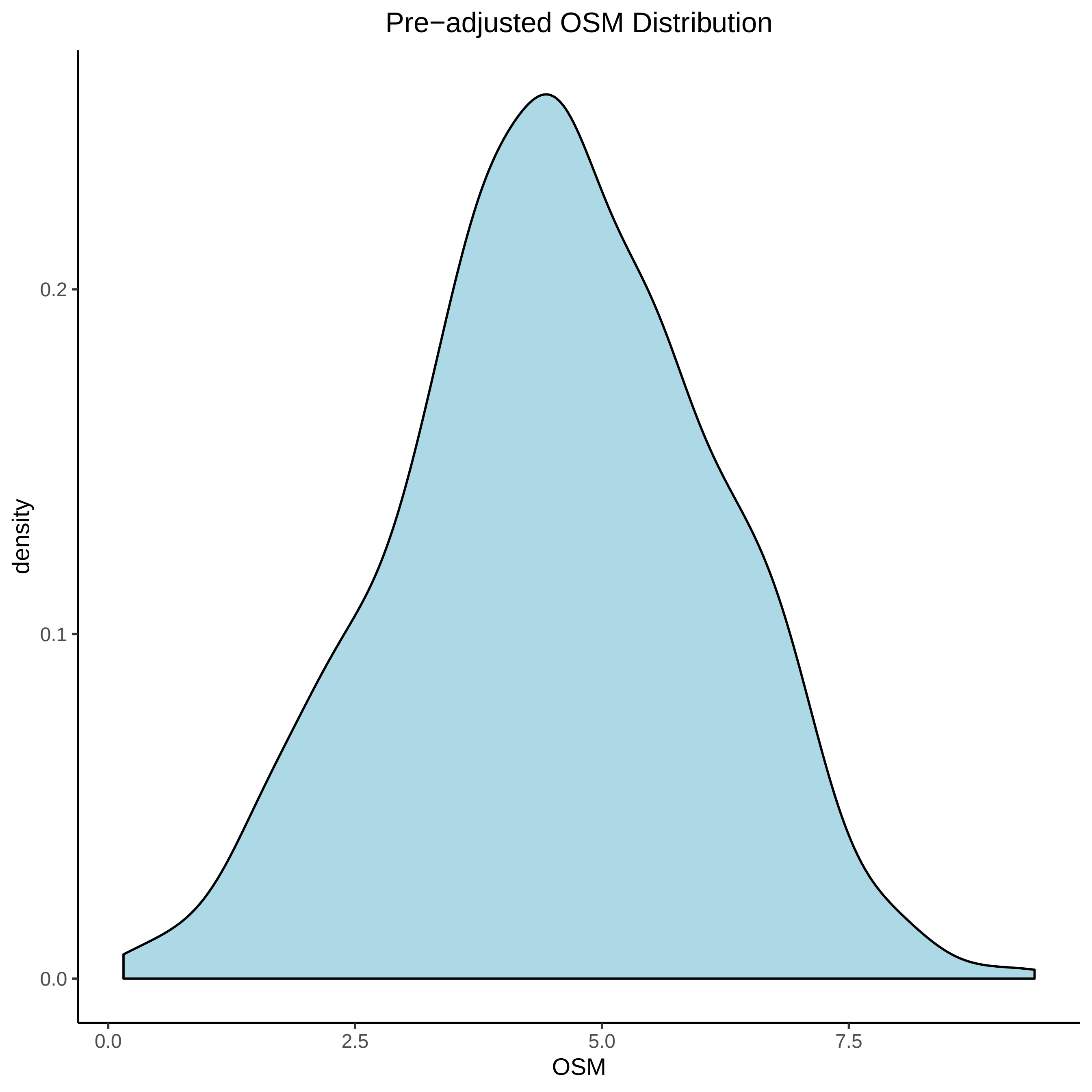

Pre−adjusted OSM Distribution
density
0.2
0.1
0.0
0.0
2.5
5.0
7.5
OSM

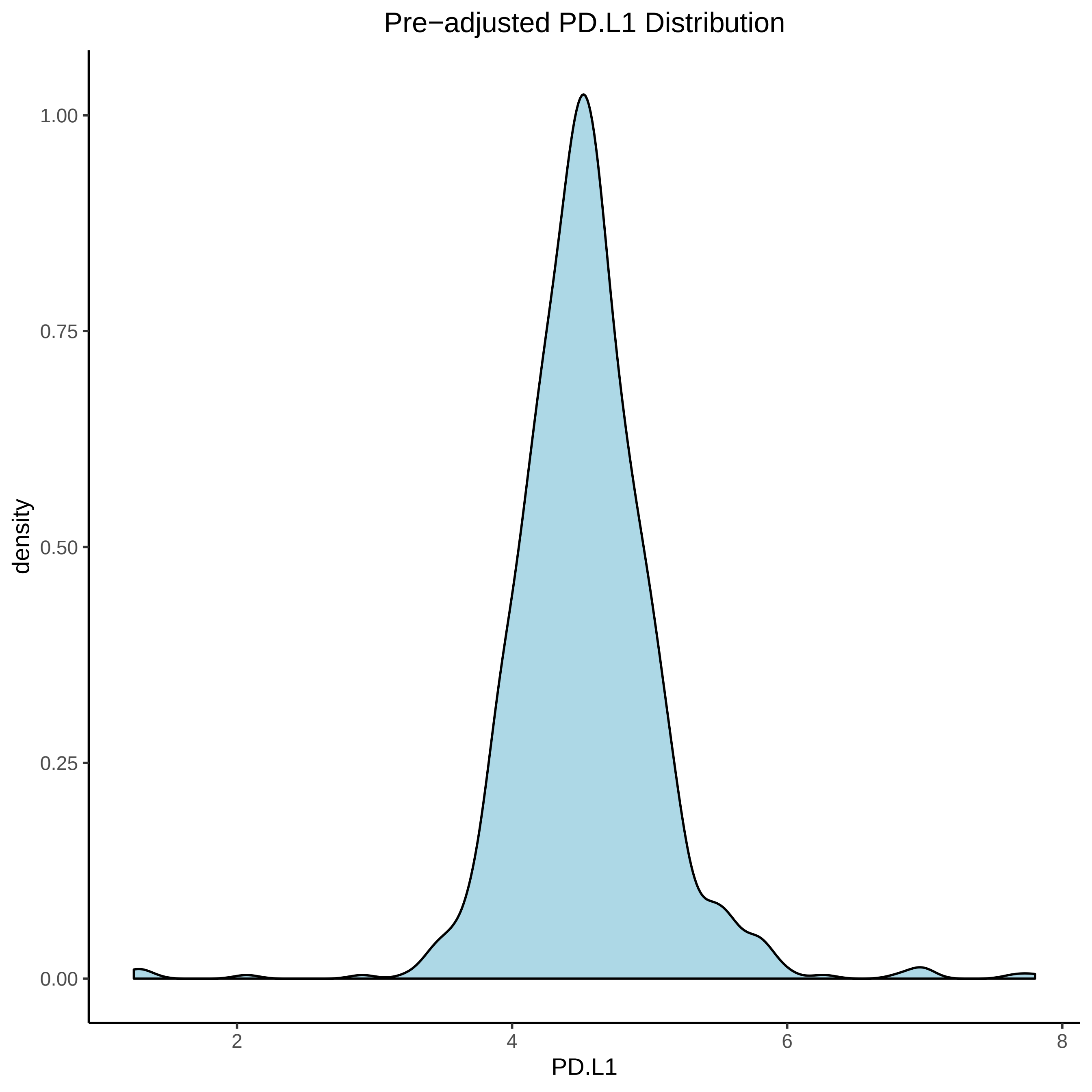

Pre−adjusted PD.L1 Distribution
1.00
0.75
0.50
0.25
0.00
density
2
4
6
8
PD.L1

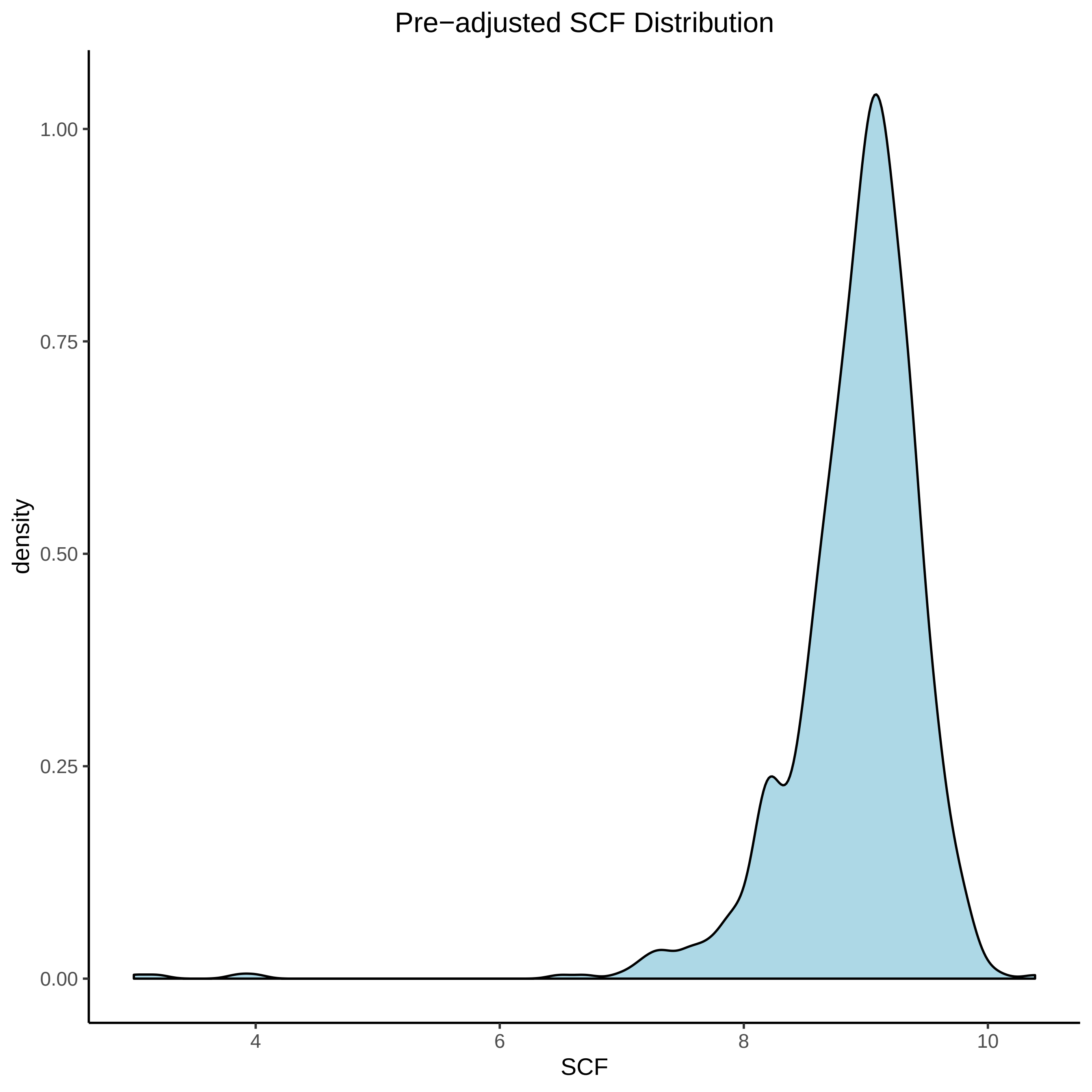

Pre−adjusted SCF Distribution
density
1.00
0.75
0.50
0.25
0.00
4
6
8
10
SCF

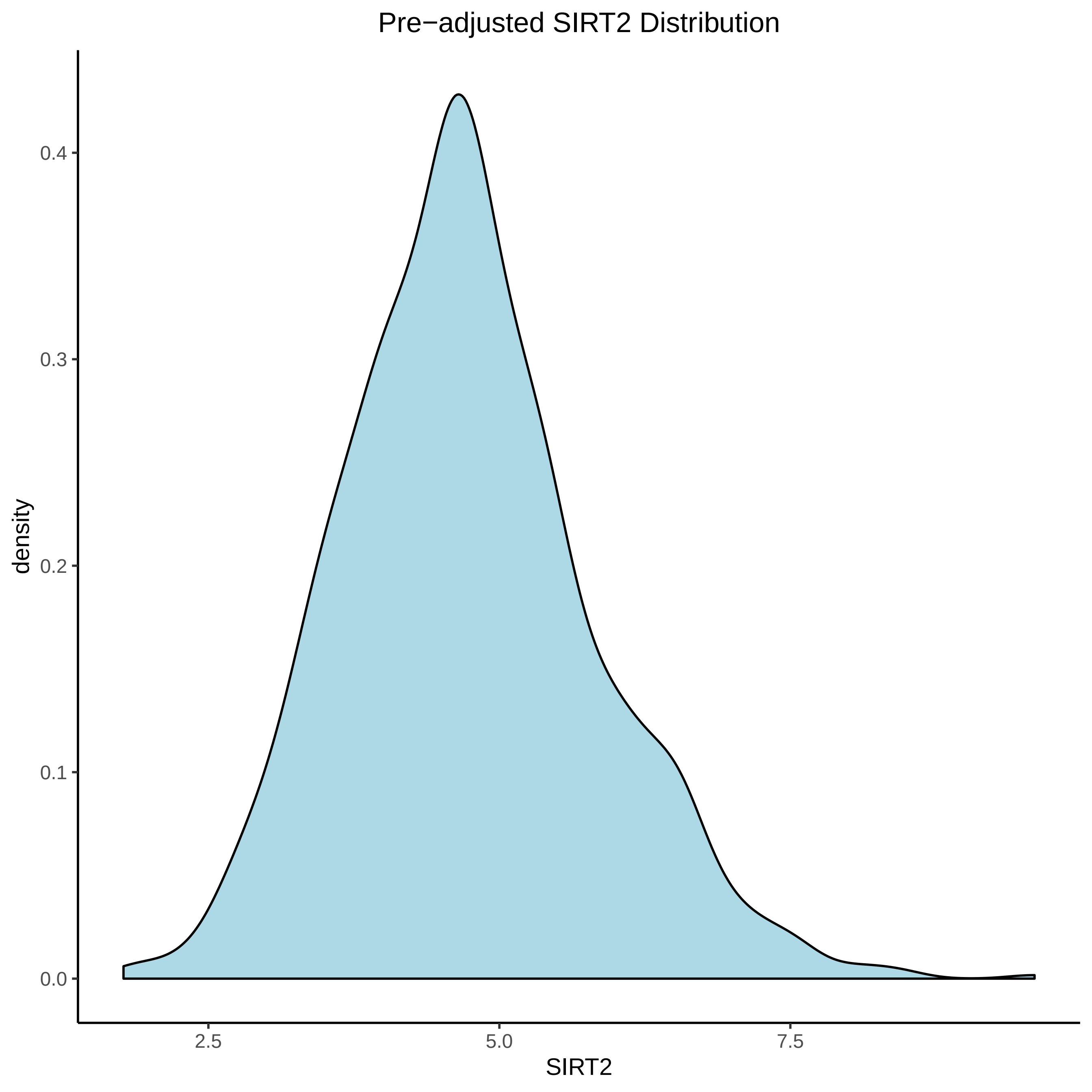
Pre−adjusted SIRT2 Distribution
density
0.4
0.3
0.2
0.1
0.0
2.5
5.0
7.5
SIRT2

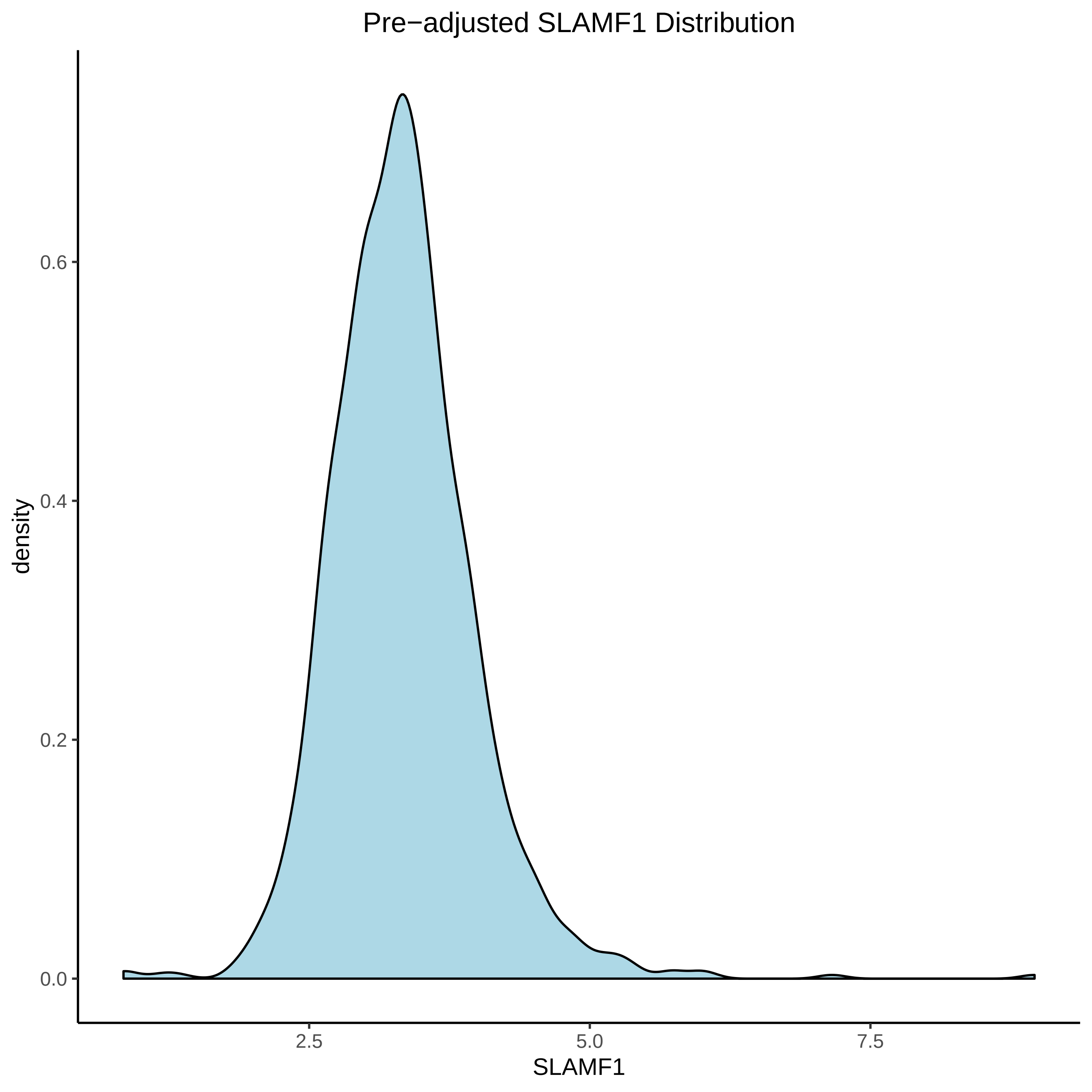
Pre−adjusted SLAMF1 Distribution
density
0.0
0.2
0.4
0.6
2.5
5.0
7.5
SLAMF1

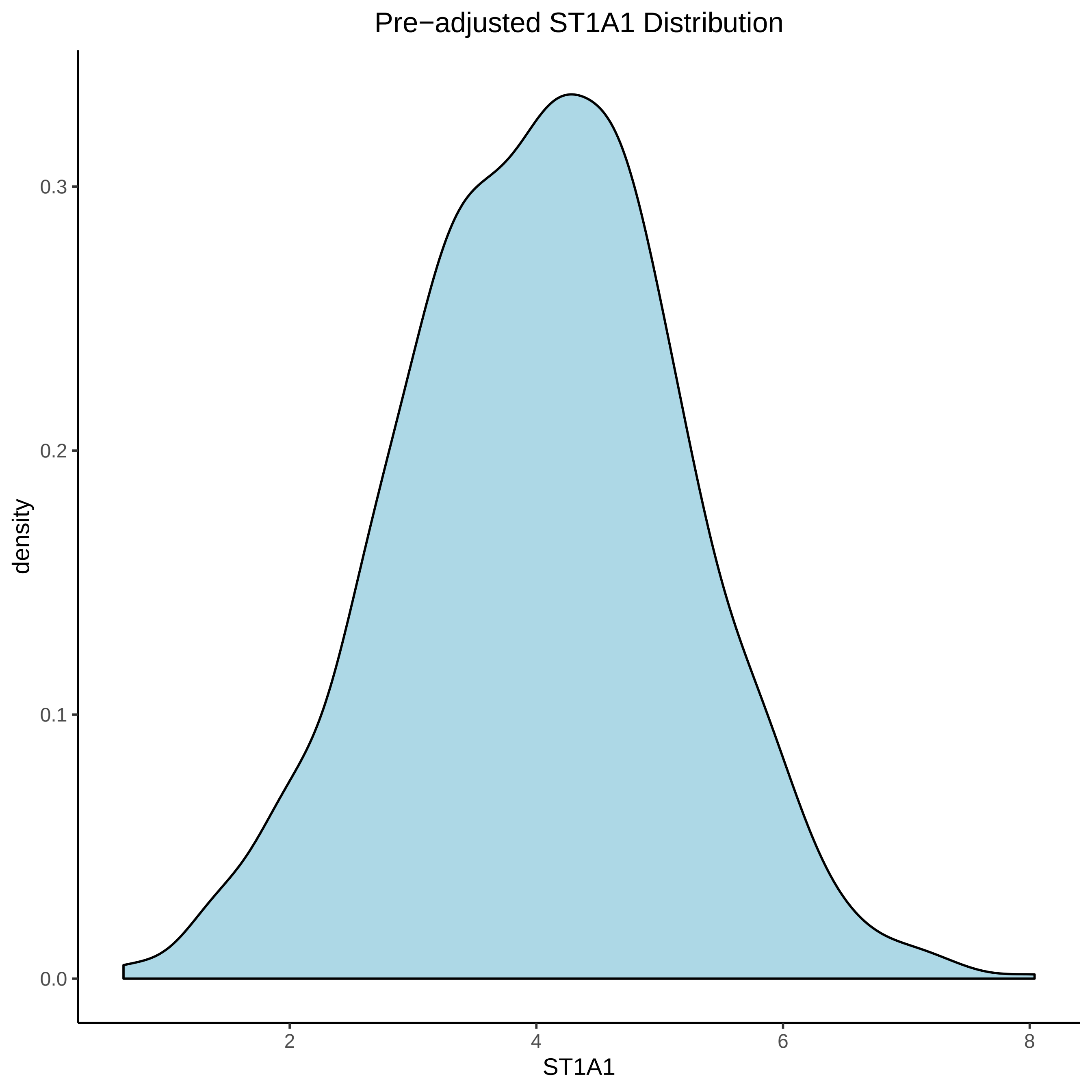
Pre−adjusted ST1A1 Distribution
density
0.3
0.2
0.1
0.0
2
4
6
8
ST1A1

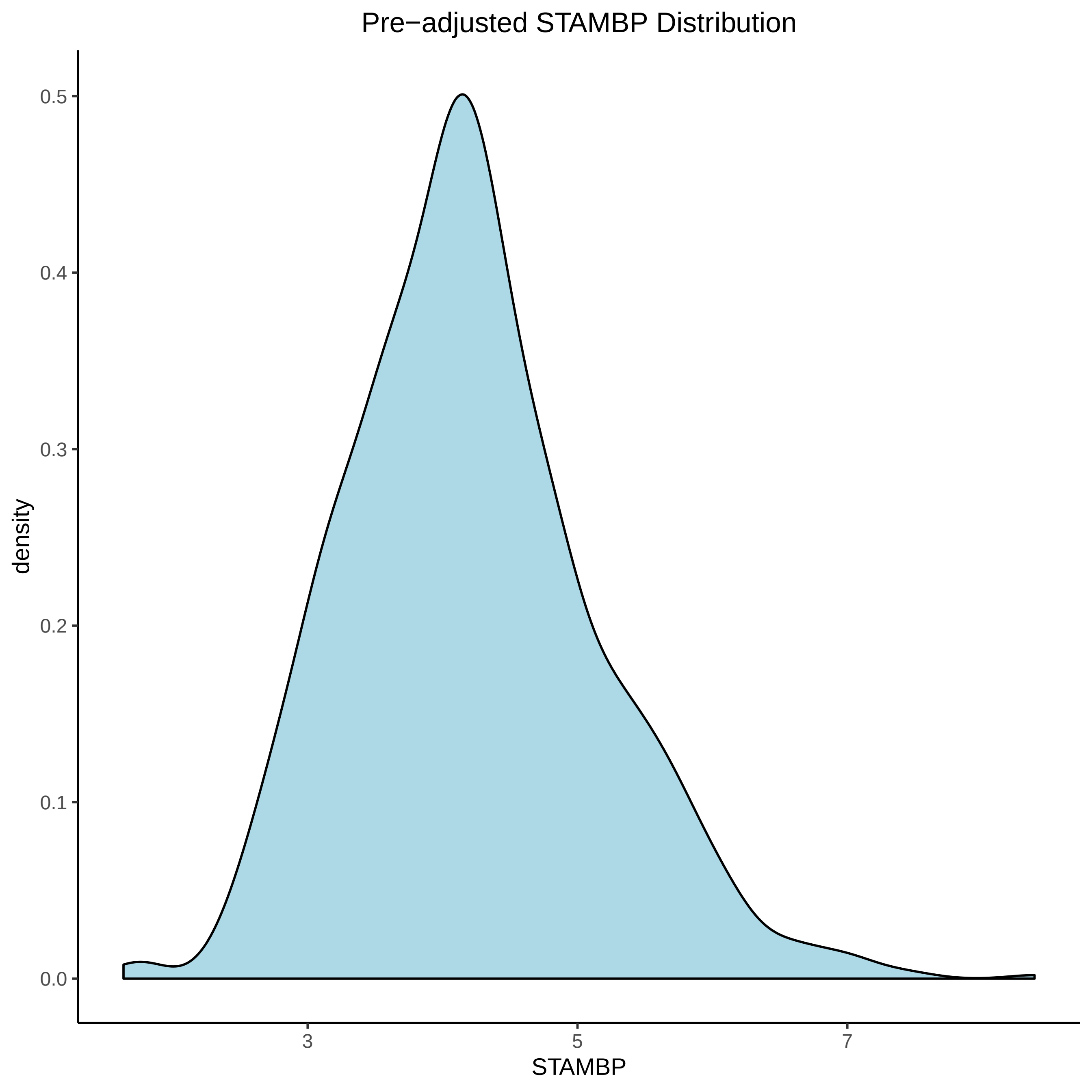

Pre−adjusted STAMBP Distribution
density
0.5
0.4
0.3
0.2
0.1
0.0
3
5
7
STAMBP

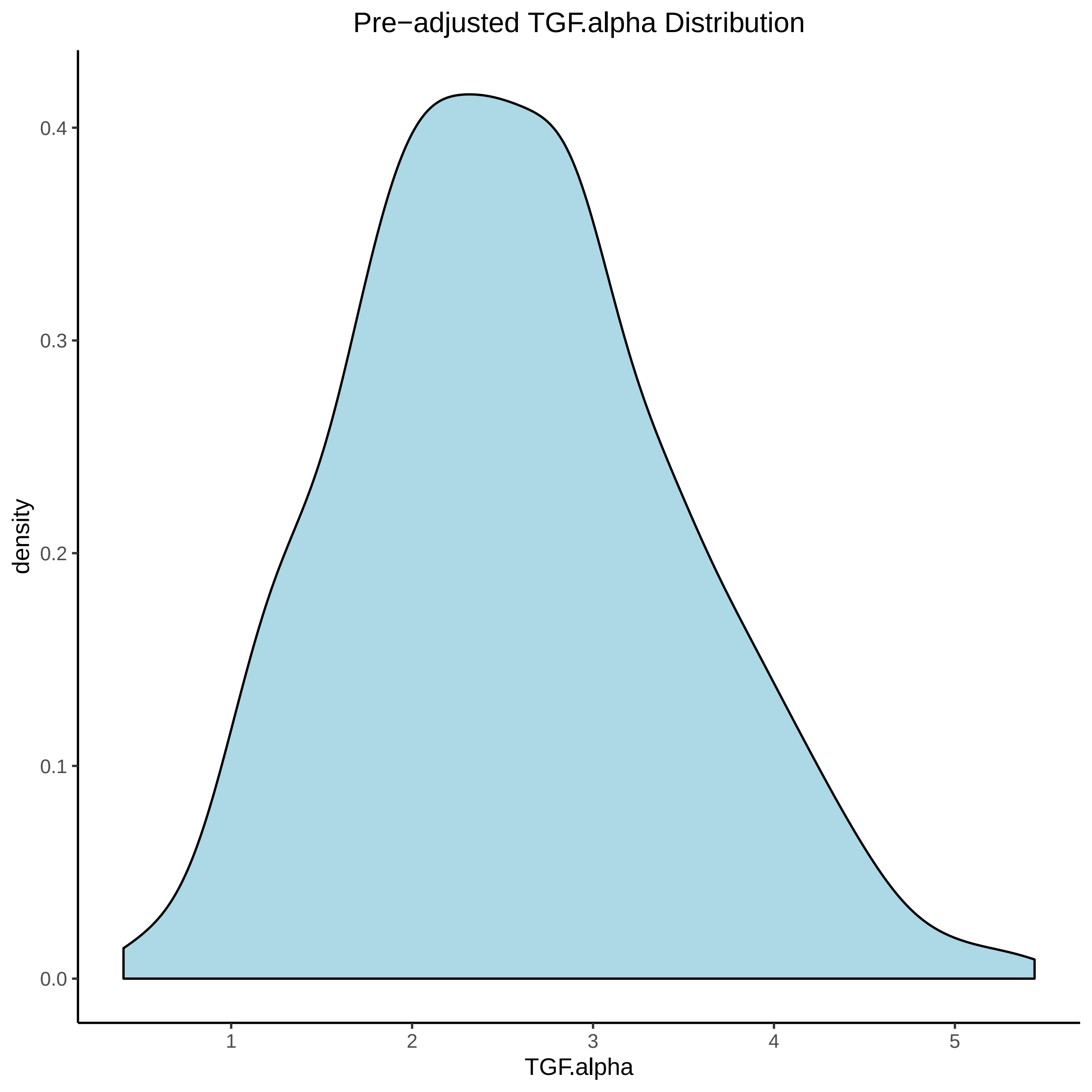

Pre−adjusted TGF.alpha Distribution
density
0.4
0.3
0.2
0.1
0.0
1
2
3
4
5
TGF.alpha

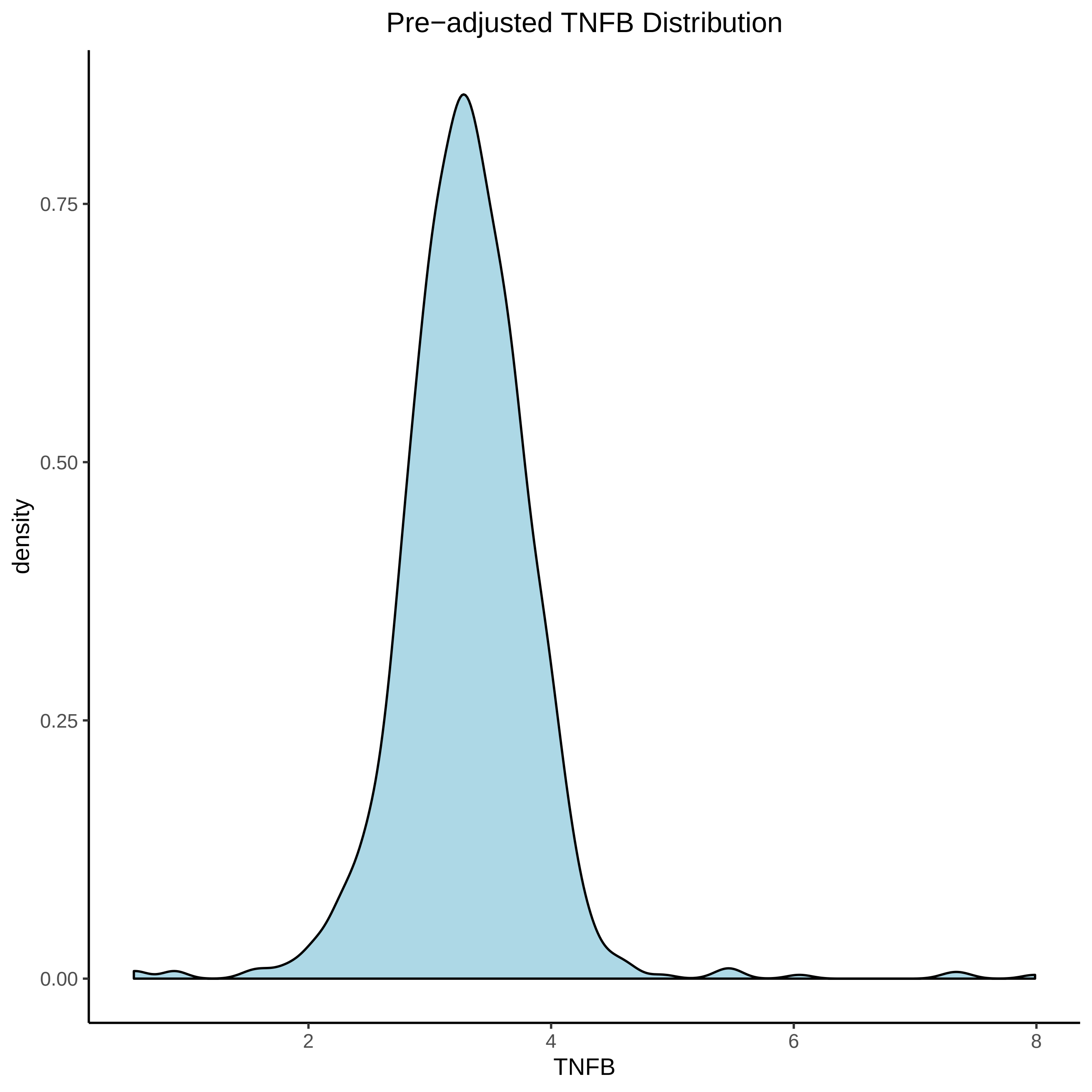
Pre−adjusted TNFB Distribution
0.75
0.50
0.25
0.00
density
2
4
6
8
TNFB

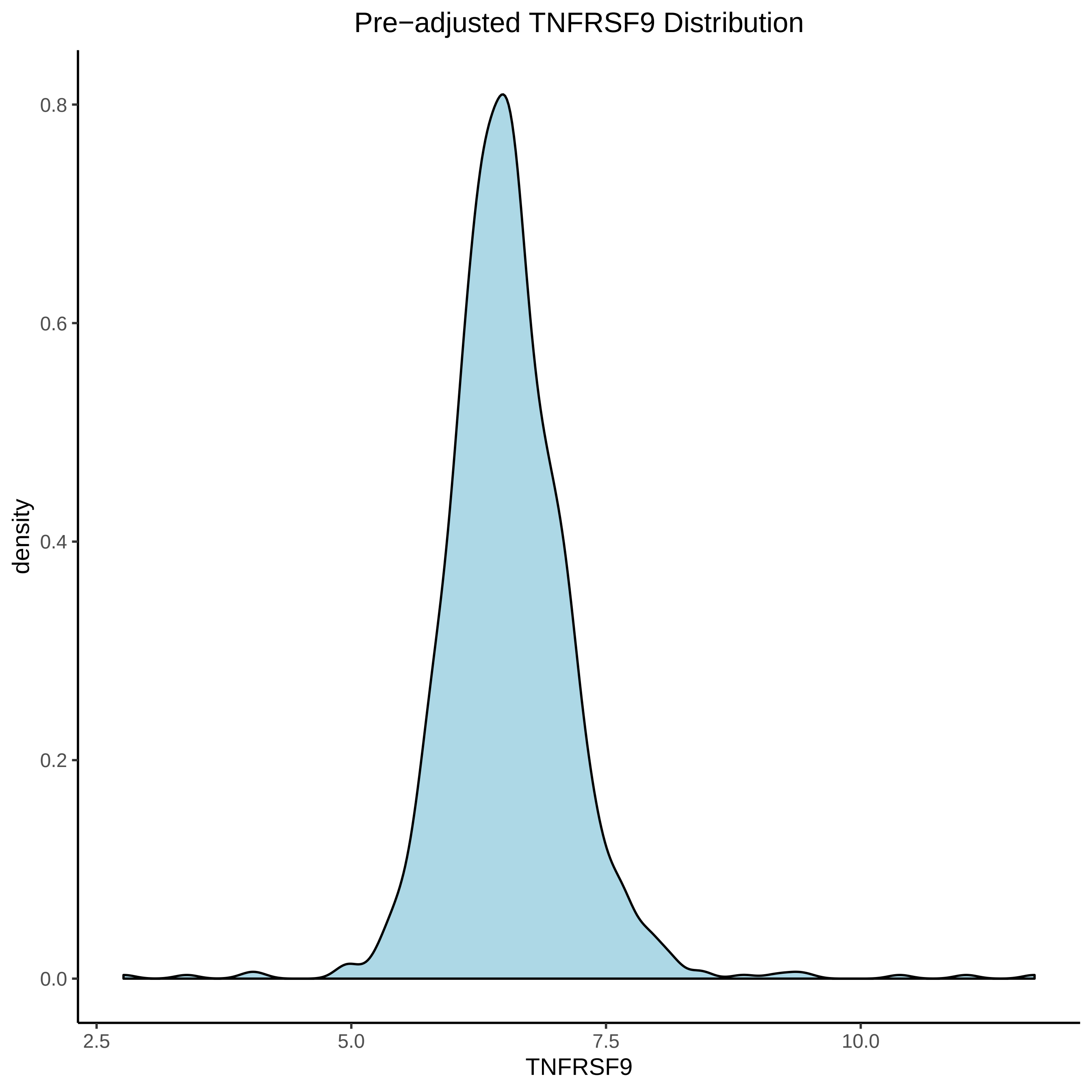

Pre−adjusted TNFRSF9 Distribution
density
0.8
0.6
0.4
0.2
0.0
2.5
5.0
7.5
10.0
TNFRSF9

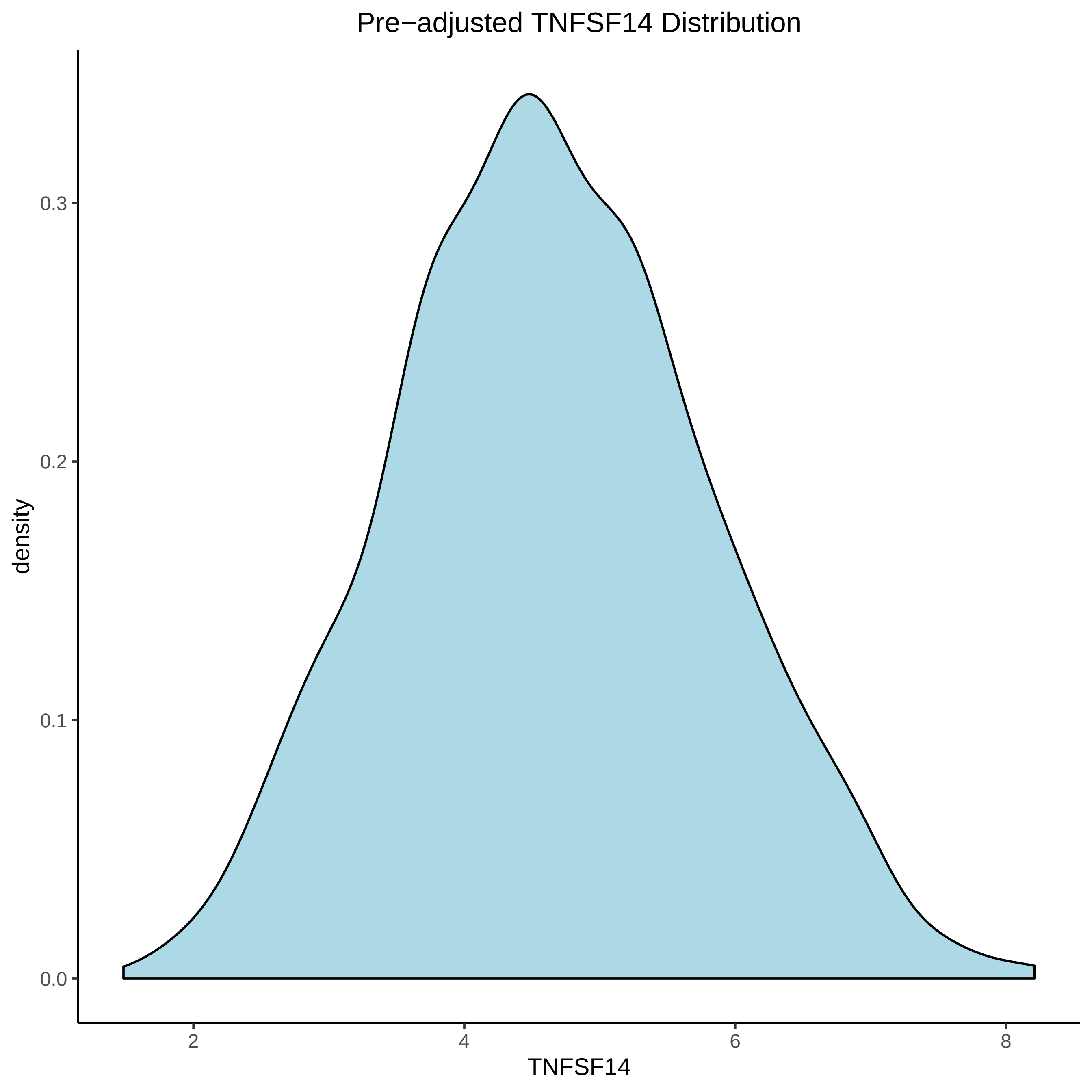

Pre−adjusted TNFSF14 Distribution
density
0.0
0.1
0.2
0.3
2
4
6
8
TNFSF14

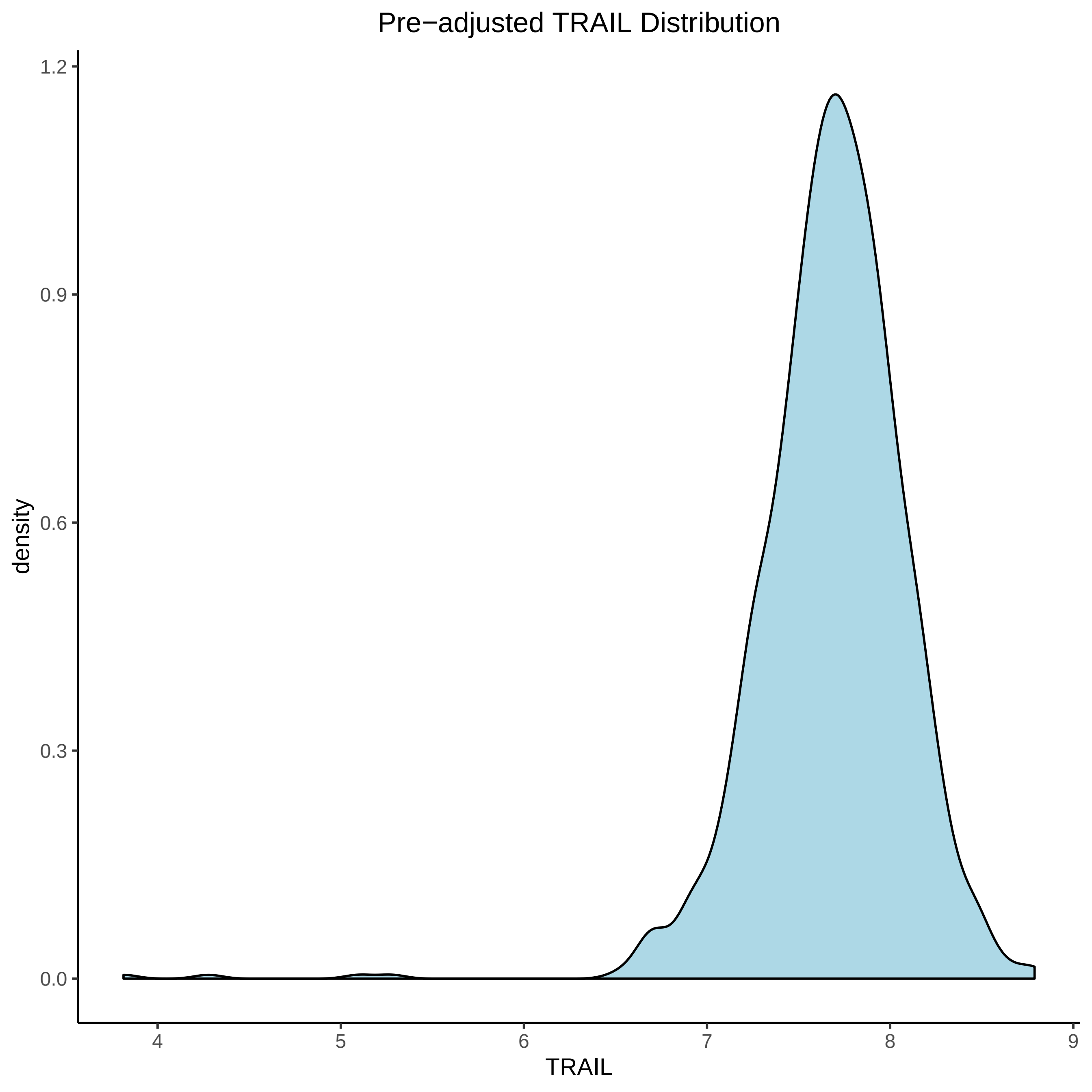

Pre−adjusted TRAIL Distribution
density
TRAIL
1.2
0.9
0.6
0.3
0.0
4
5
6
7
8
9

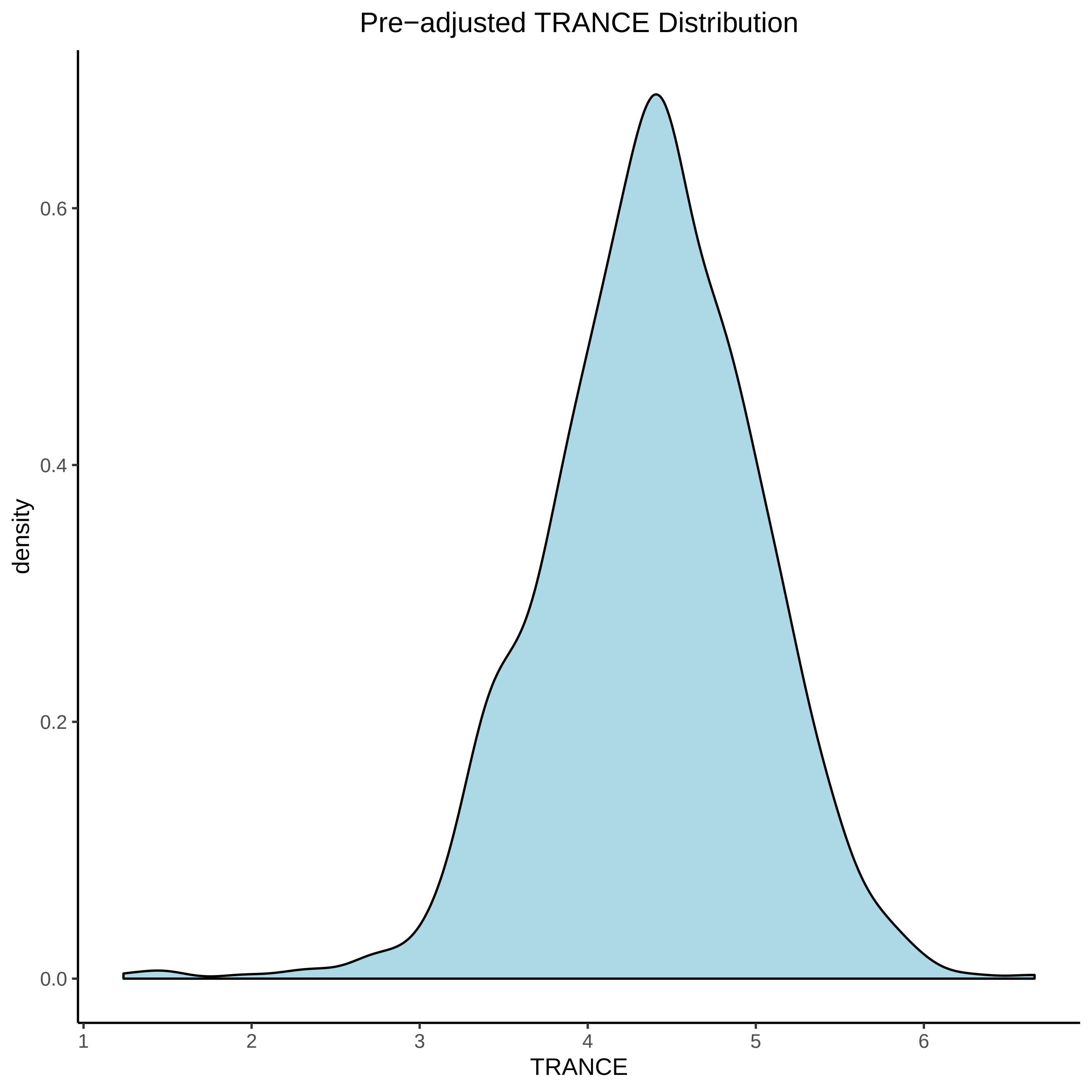
Pre−adjusted TRANCE Distribution
density
0.6
0.4
0.2
0.0
1
2
3
4
5
6
TRANCE

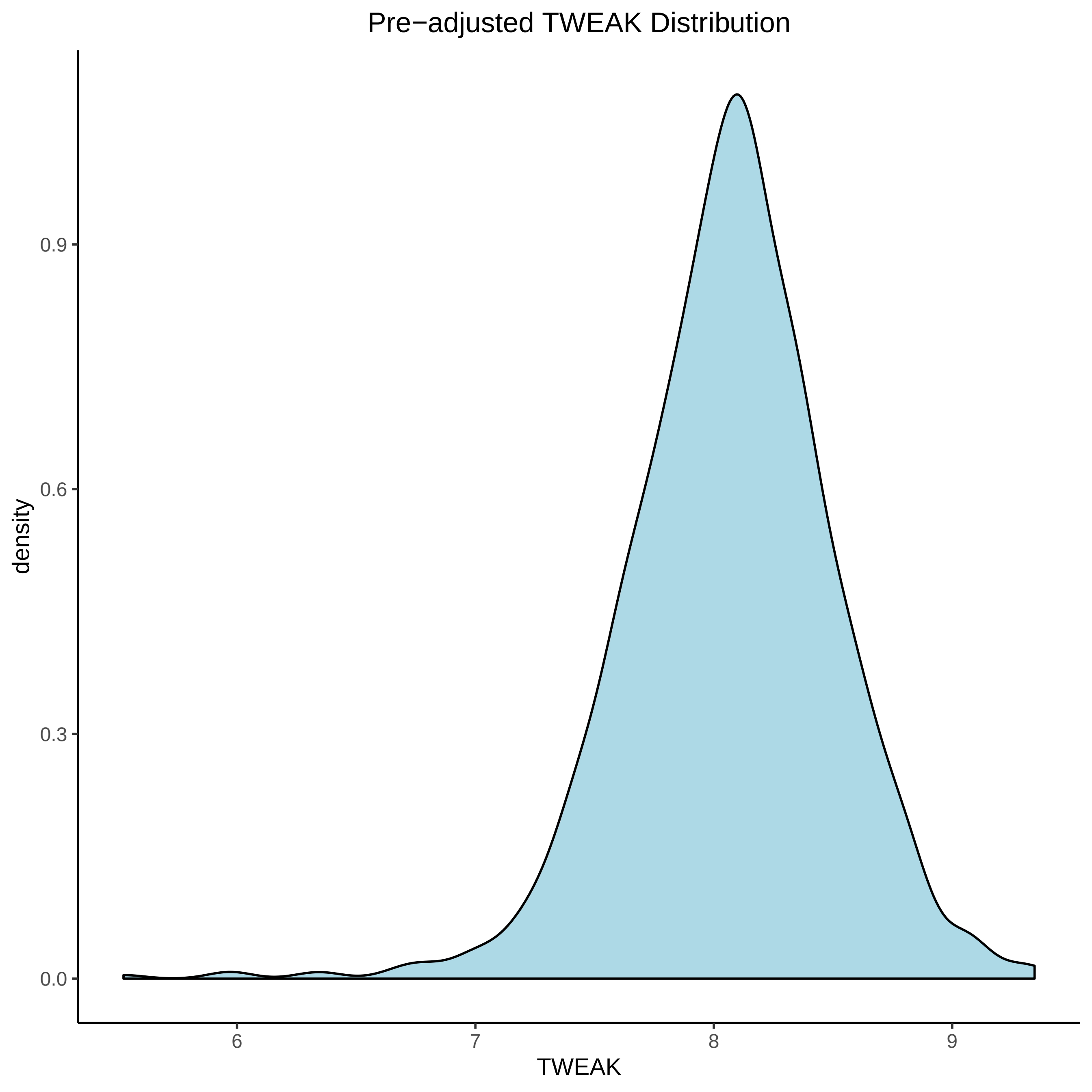
Pre−adjusted TWEAK Distribution
density
0.0
0.3
0.6
0.9
6
7
8
9
TWEAK

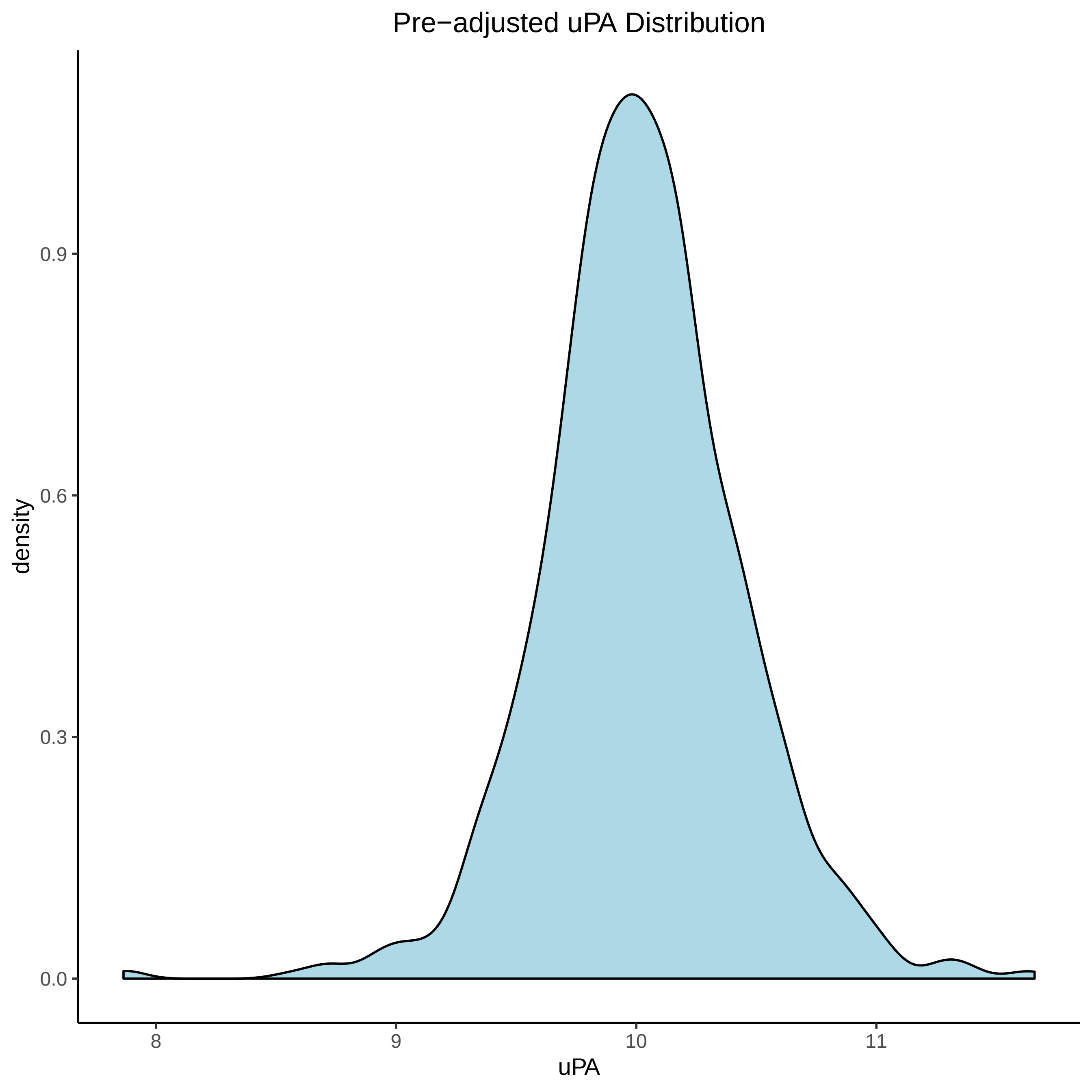
Pre−adjusted uPA Distribution
density
0.0
0.3
0.6
0.9
8
9
10
11
uPA

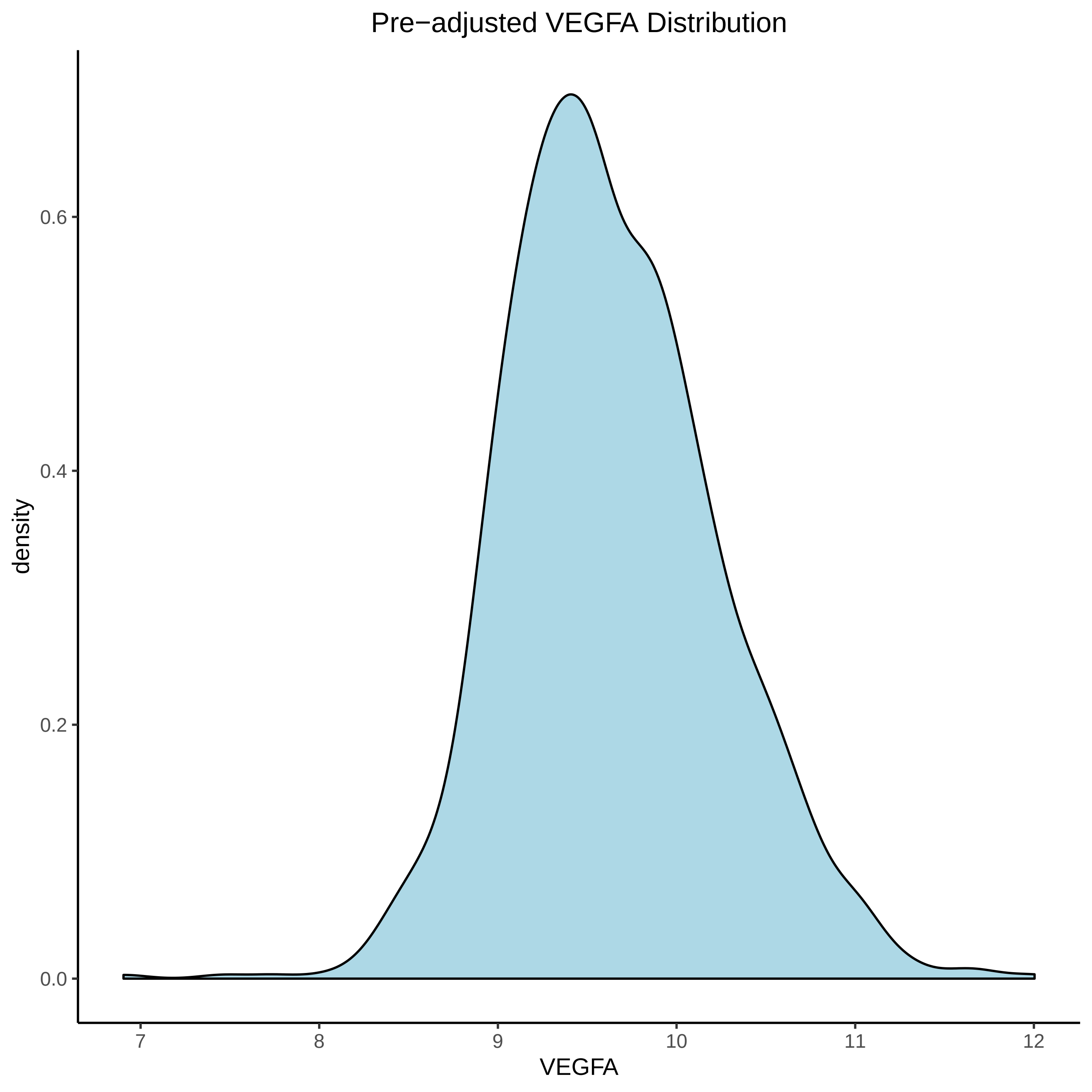

Pre−adjusted VEGFA Distribution
density
0.0
0.2
0.4
0.6
7
8
9
10
11
12
VEGFA

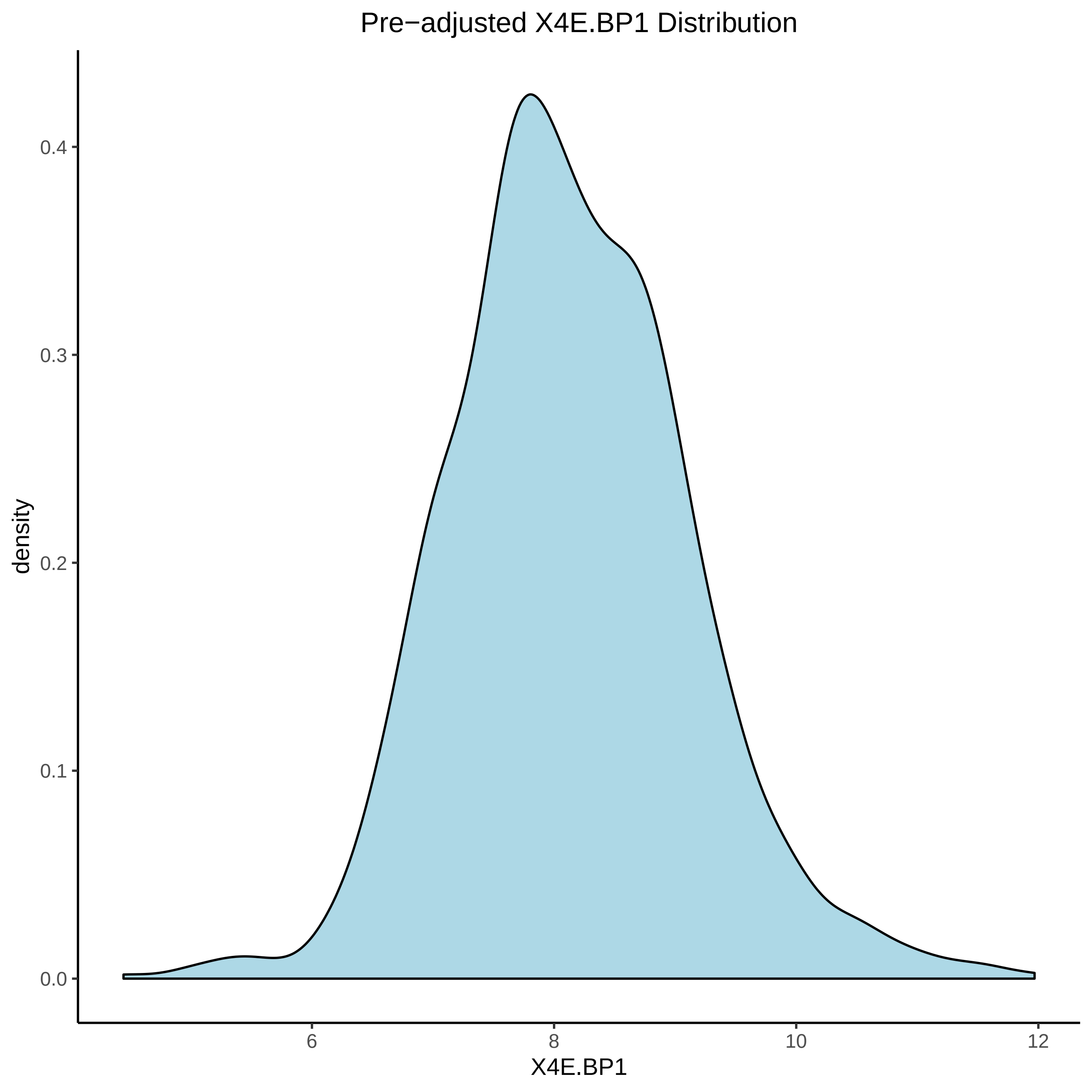
Pre−adjusted X4E.BP1 Distribution
density
0.0
0.1
0.2
0.3
0.4
6
8
10
12
X4E.BP1
