## Supplementary File 3 for "Integrative omics approach to identify the molecular architecture of inflammatory protein levels in healthy older adults"

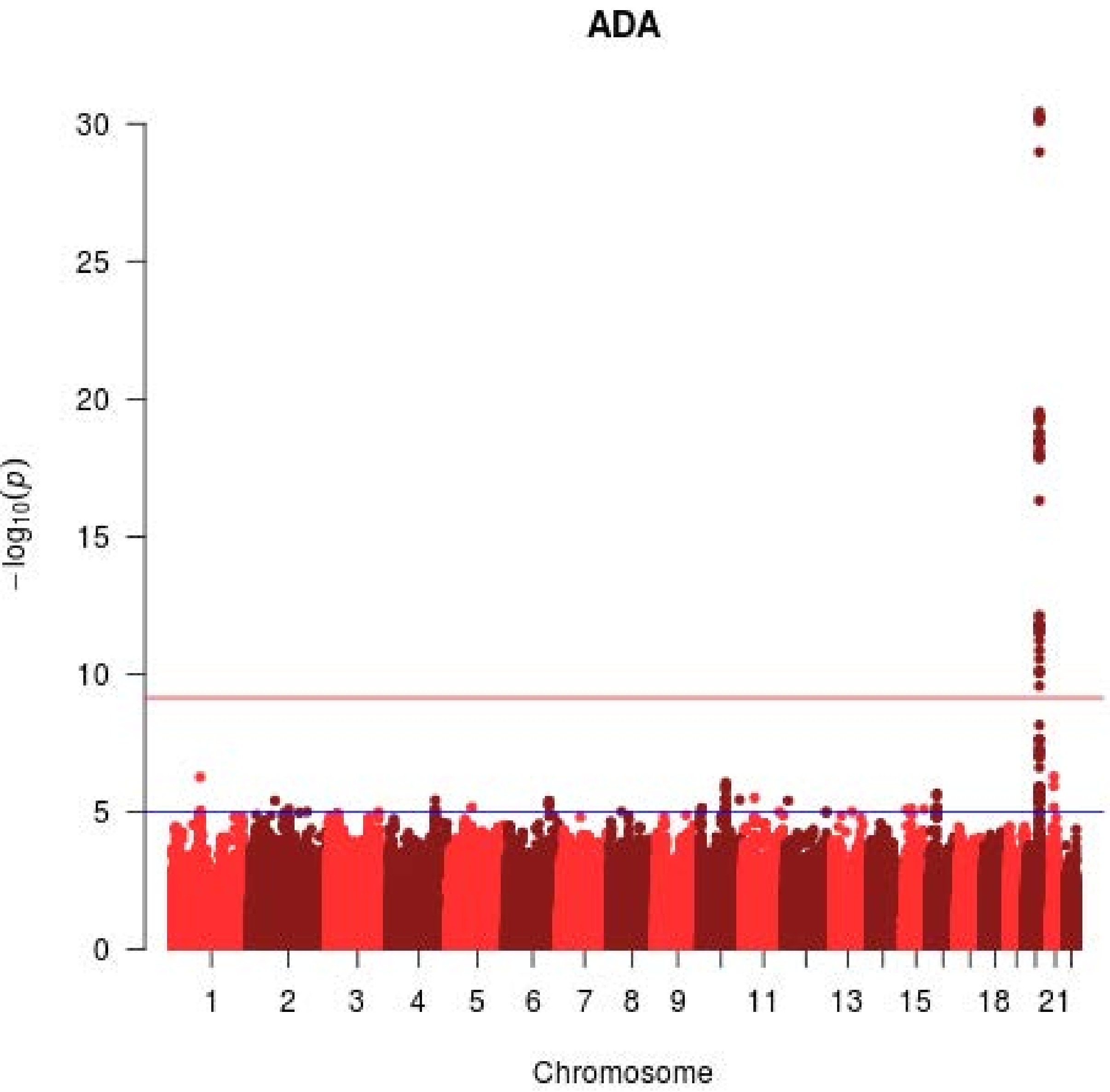
ADA
30
25
20
15
10
5
0
−log10(p)
1
2
3
4
5
6
7
8
9
11
13
15
18
21
Chromosome

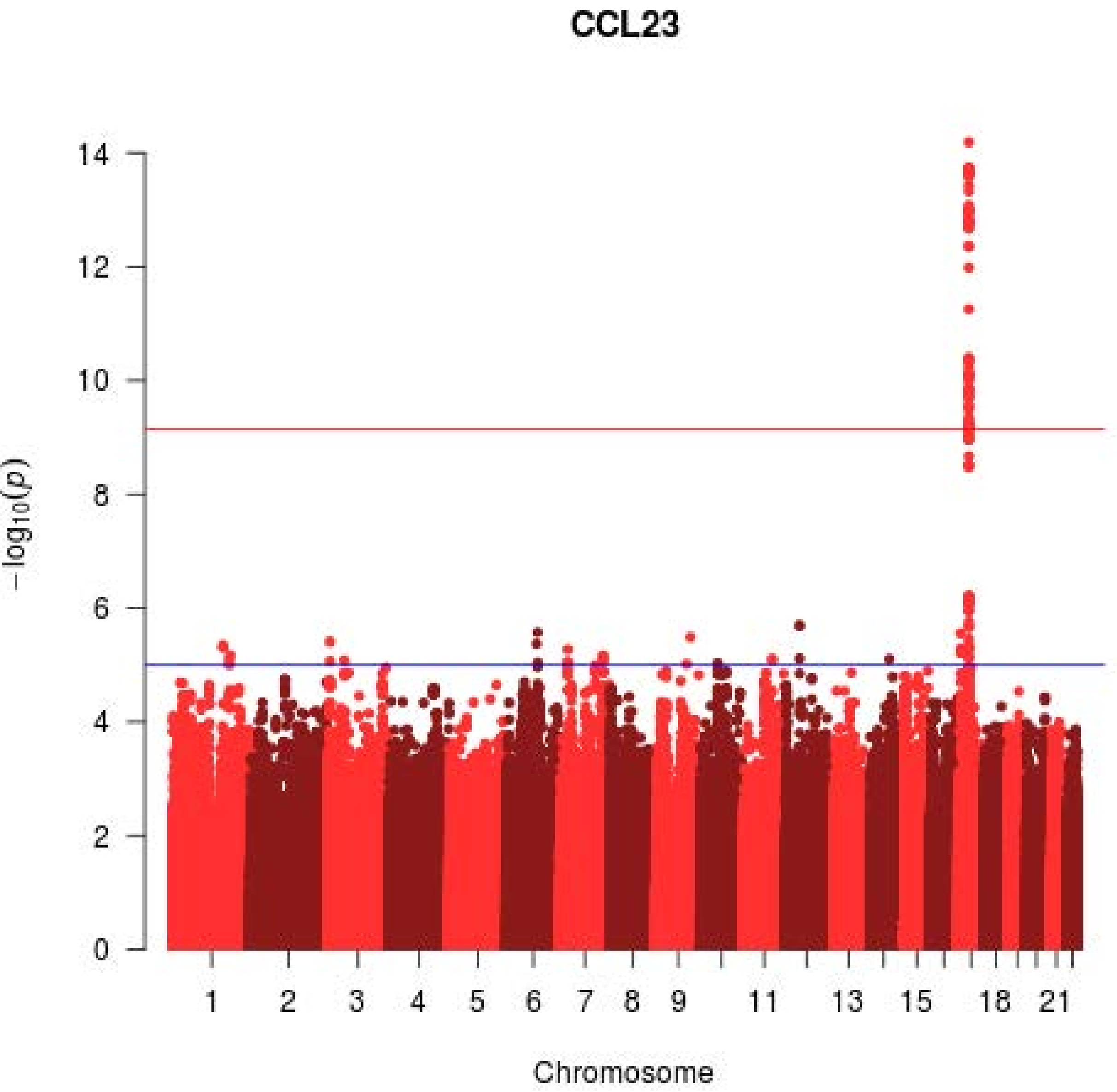
CCL23
14
12
10
8
6
4
2
0
$-\log_{10}(p)$
1
2
3
4
5
6
7
8
9
11
13
15
18
21
Chromosome

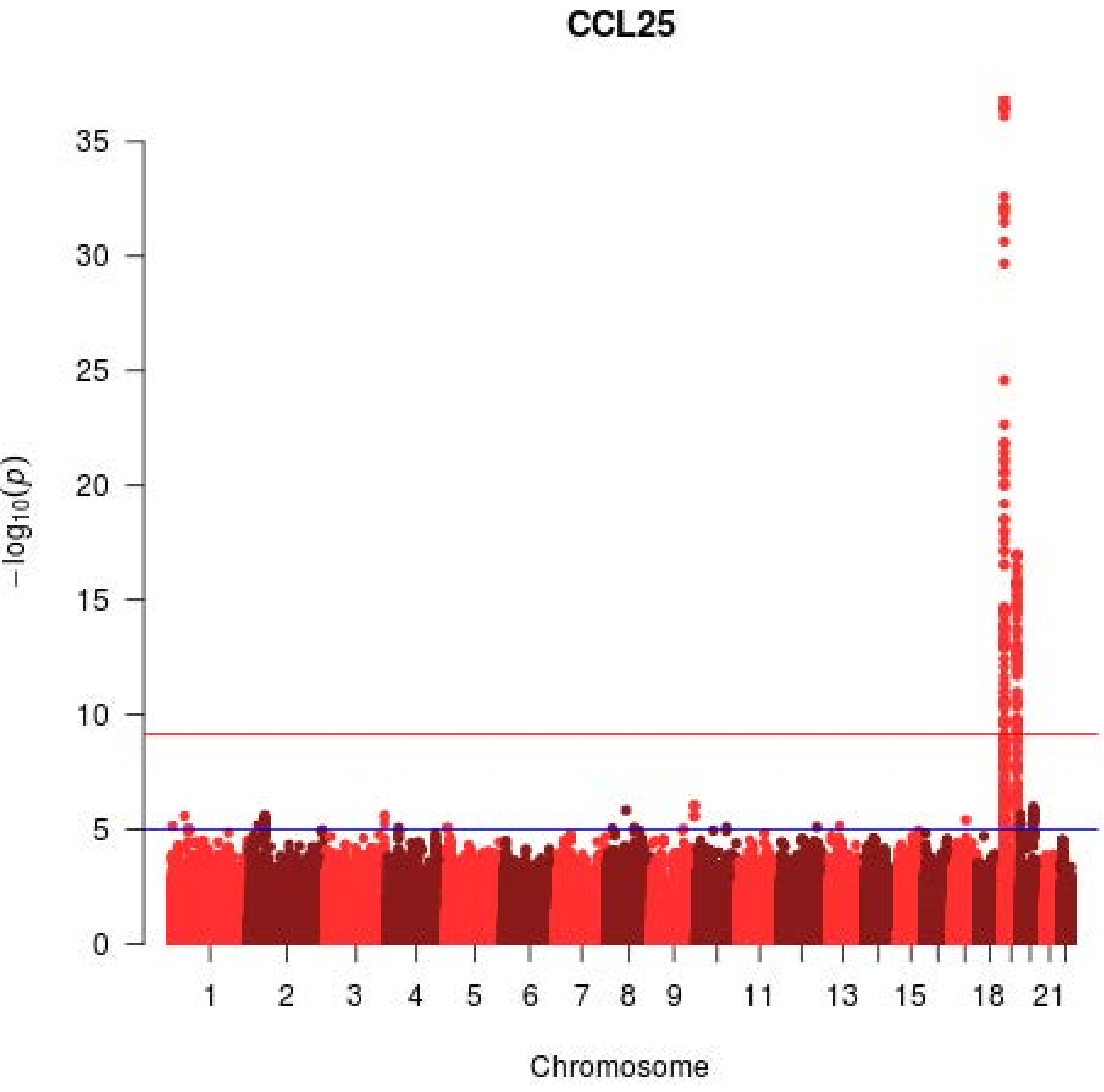

CCL25
35
30
25
20
15
10
5
0
$-\log_{10}(p)$
1
2
3
4
5
6
7
8
9
11
13
15
18
21
Chromosome

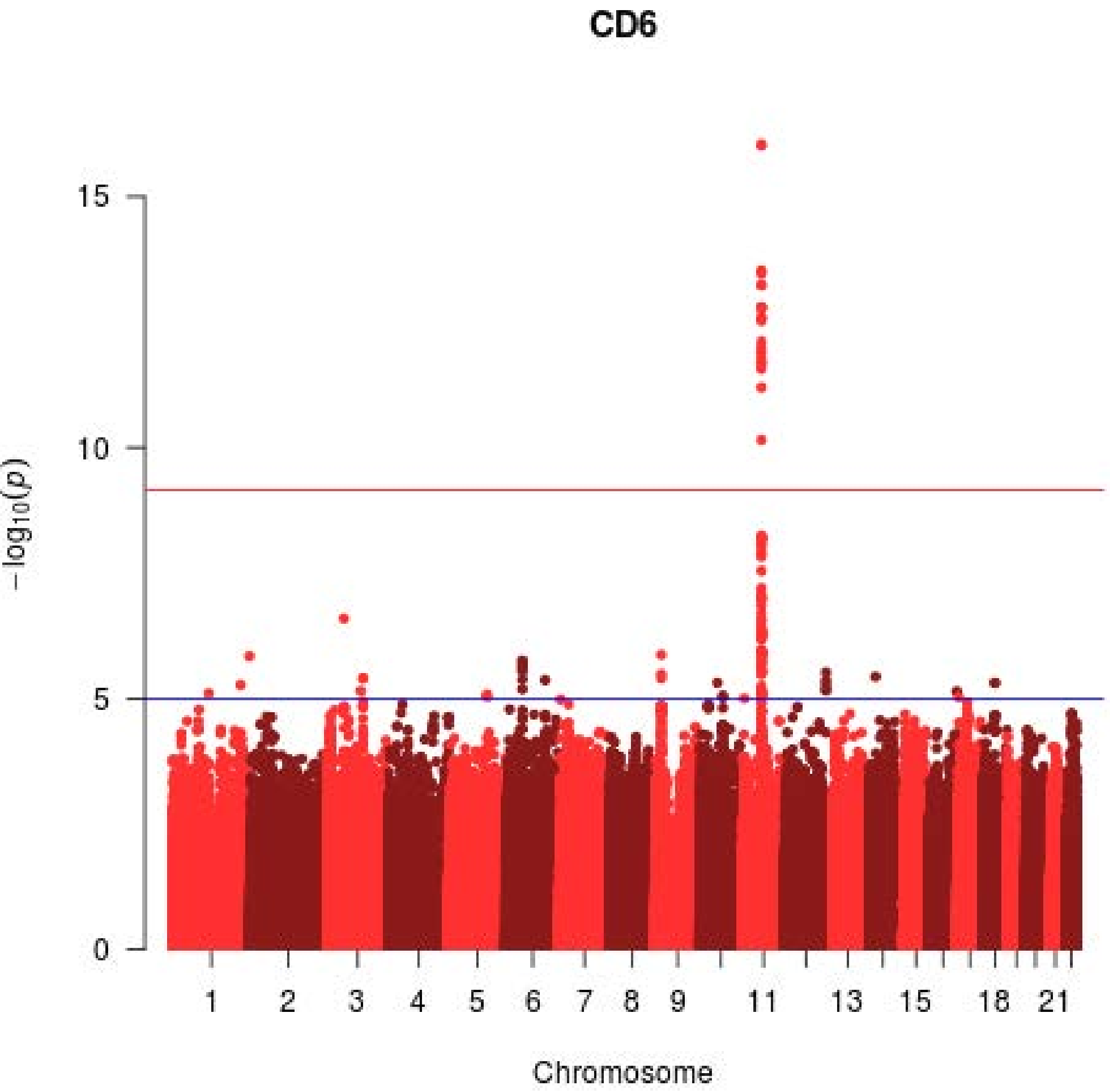
CD6
15
10
5
0
$-\log_{10}(p)$
1
2
3
4
5
6
7
8
9
11
13
15
18
21
Chromosome

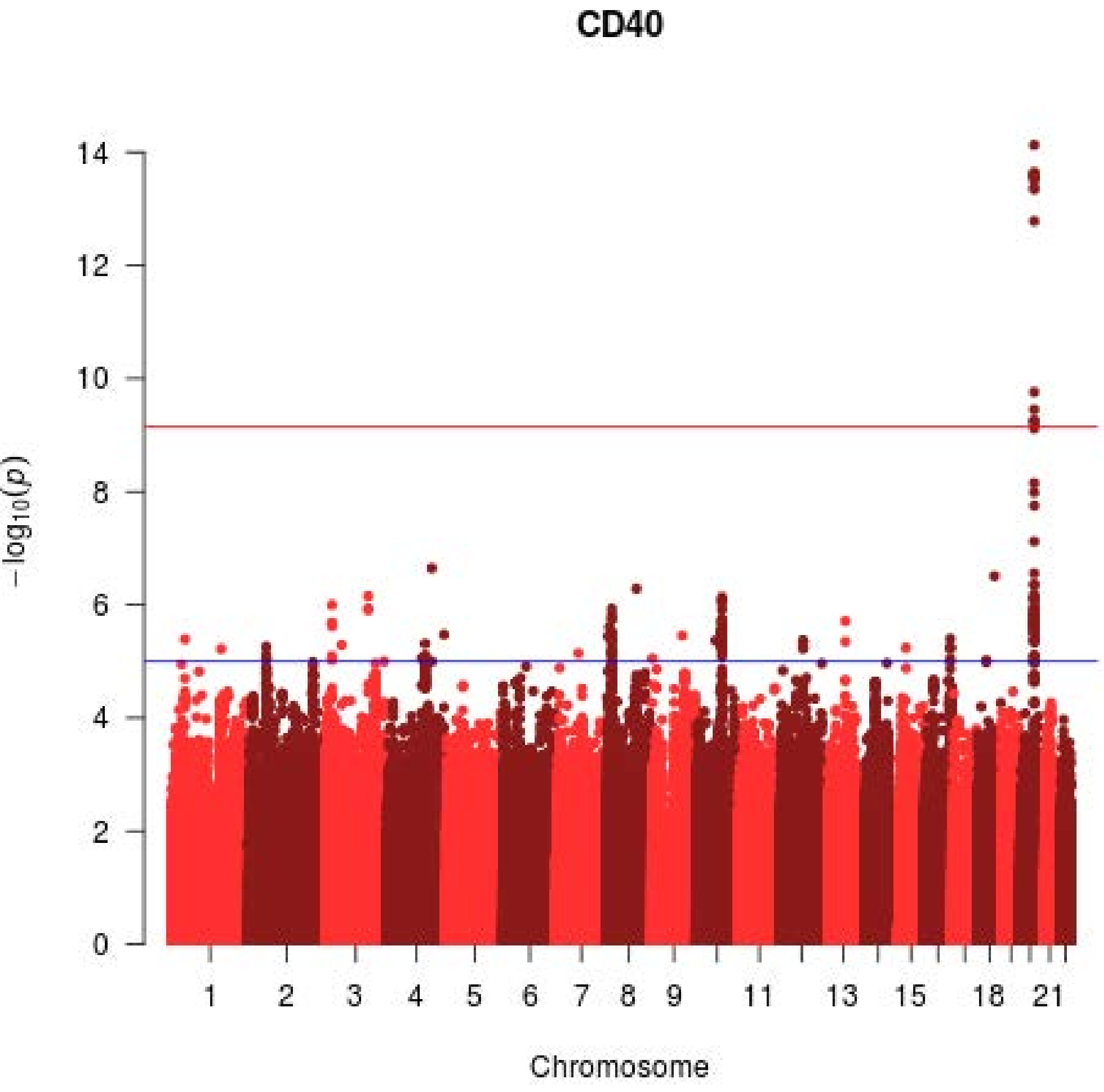

CD40
14
12
10
8
6
4
2
0
−log10(p)
1
2
3
4
5
6
7
8
9
11
13
15
18
21
Chromosome

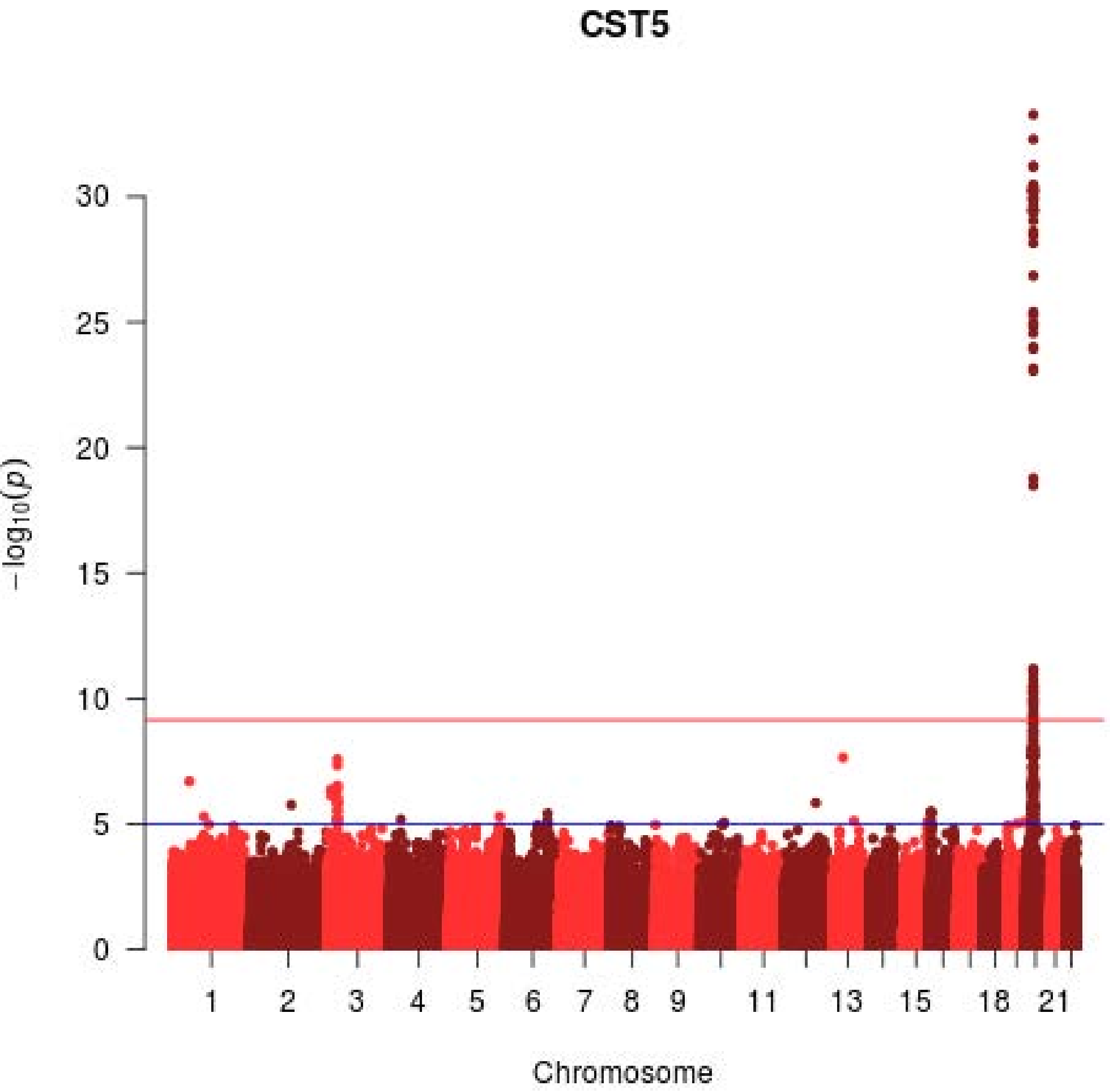

CST5
30
25
20
15
10
5
0
−log10(p)
1
2
3
4
5
6
7
8
9
11
13
15
18
21
Chromosome

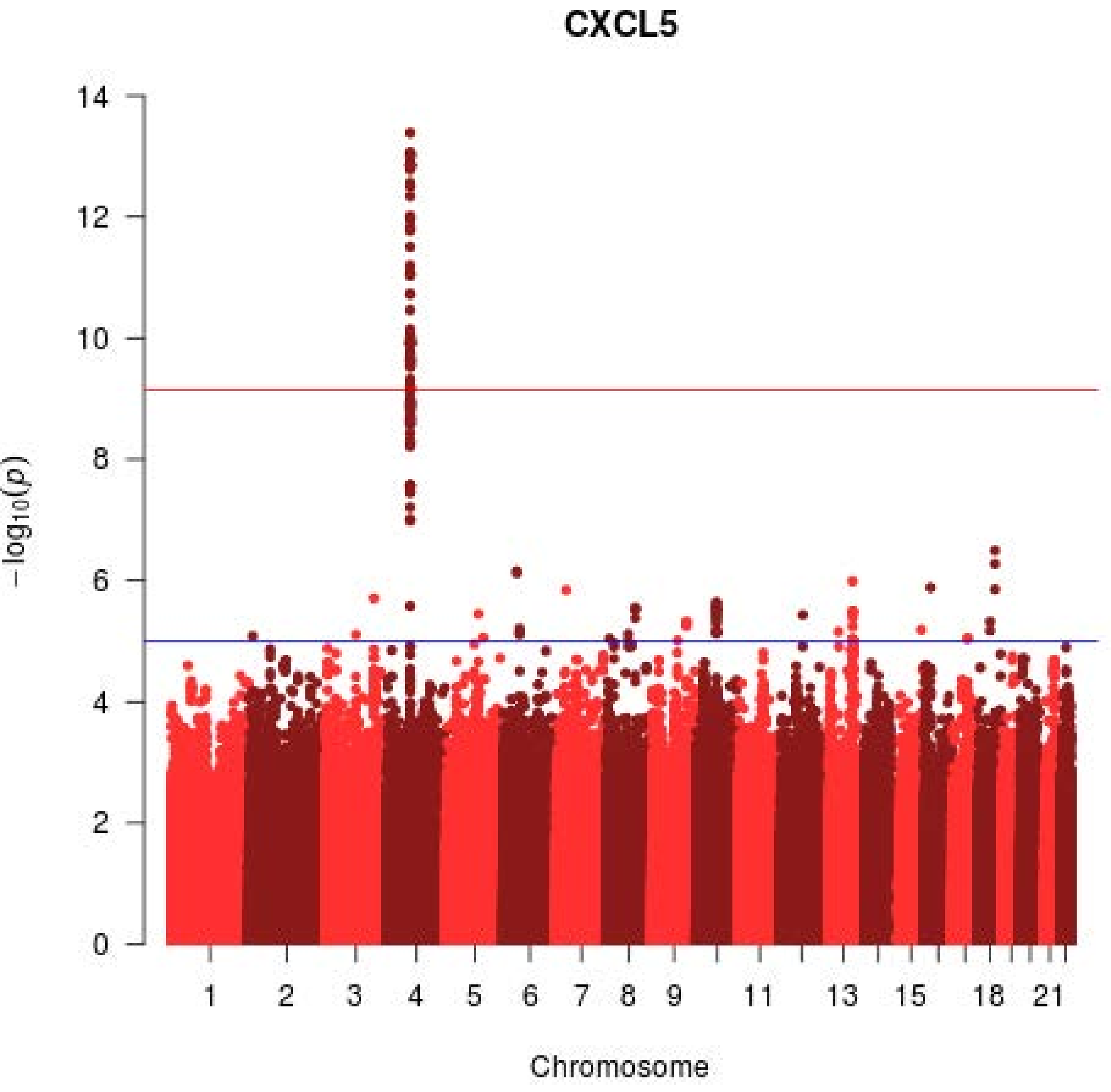

CXCL5
14
12
10
8
6
4
2
0
−log10(p)
1
2
3
4
5
6
7
8
9
11
13
15
18
21
Chromosome

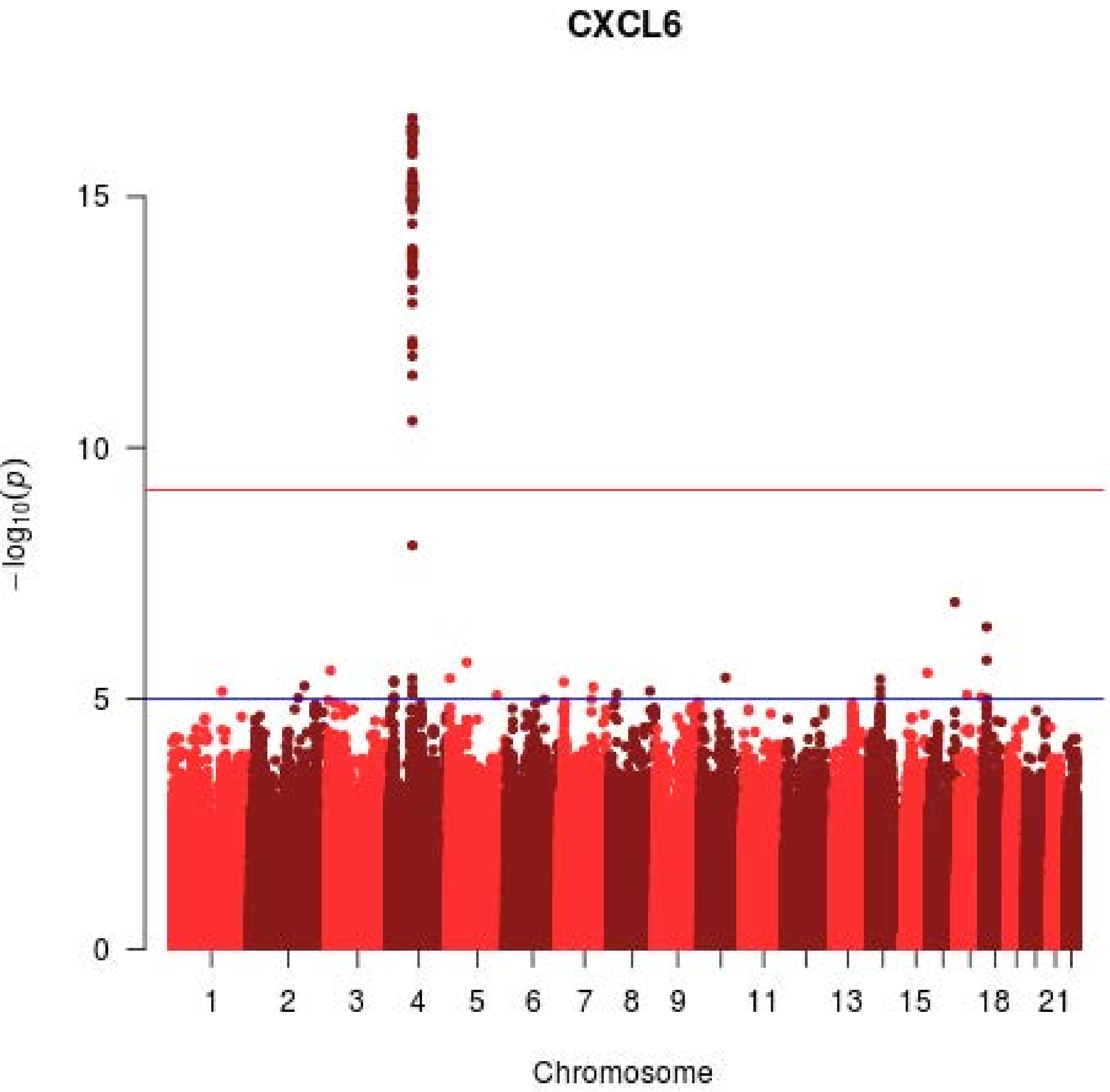
CXCL6
15
10
5
0
-log10(p)
1
2
3
4
5
6
7
8
9
11
13
15
18
21
Chromosome

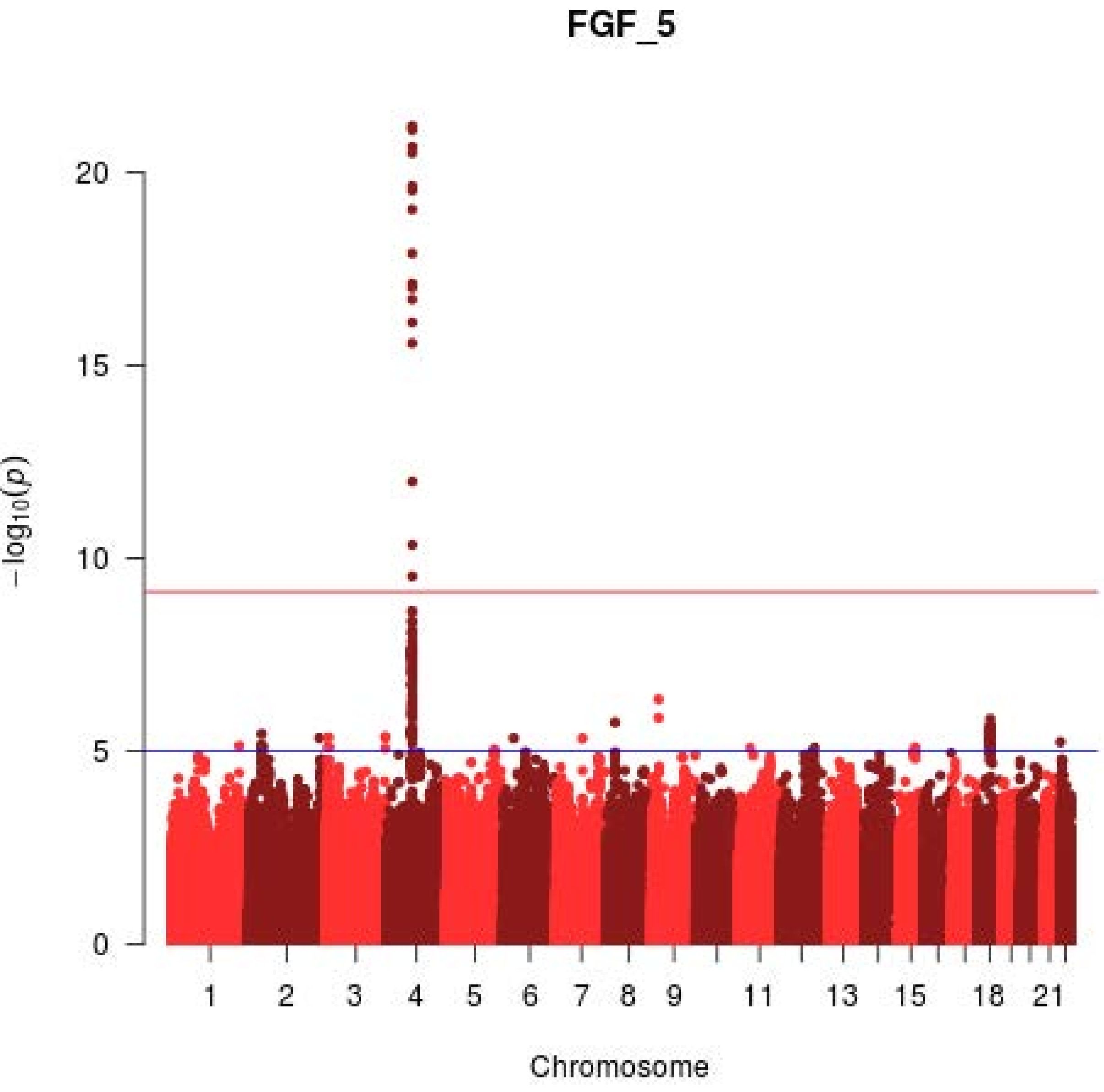
FGF_5
20
15
10
5
0
$-\log_{10}(p)$
1
2
3
4
5
6
7
8
9
11
13
15
18
21
Chromosome

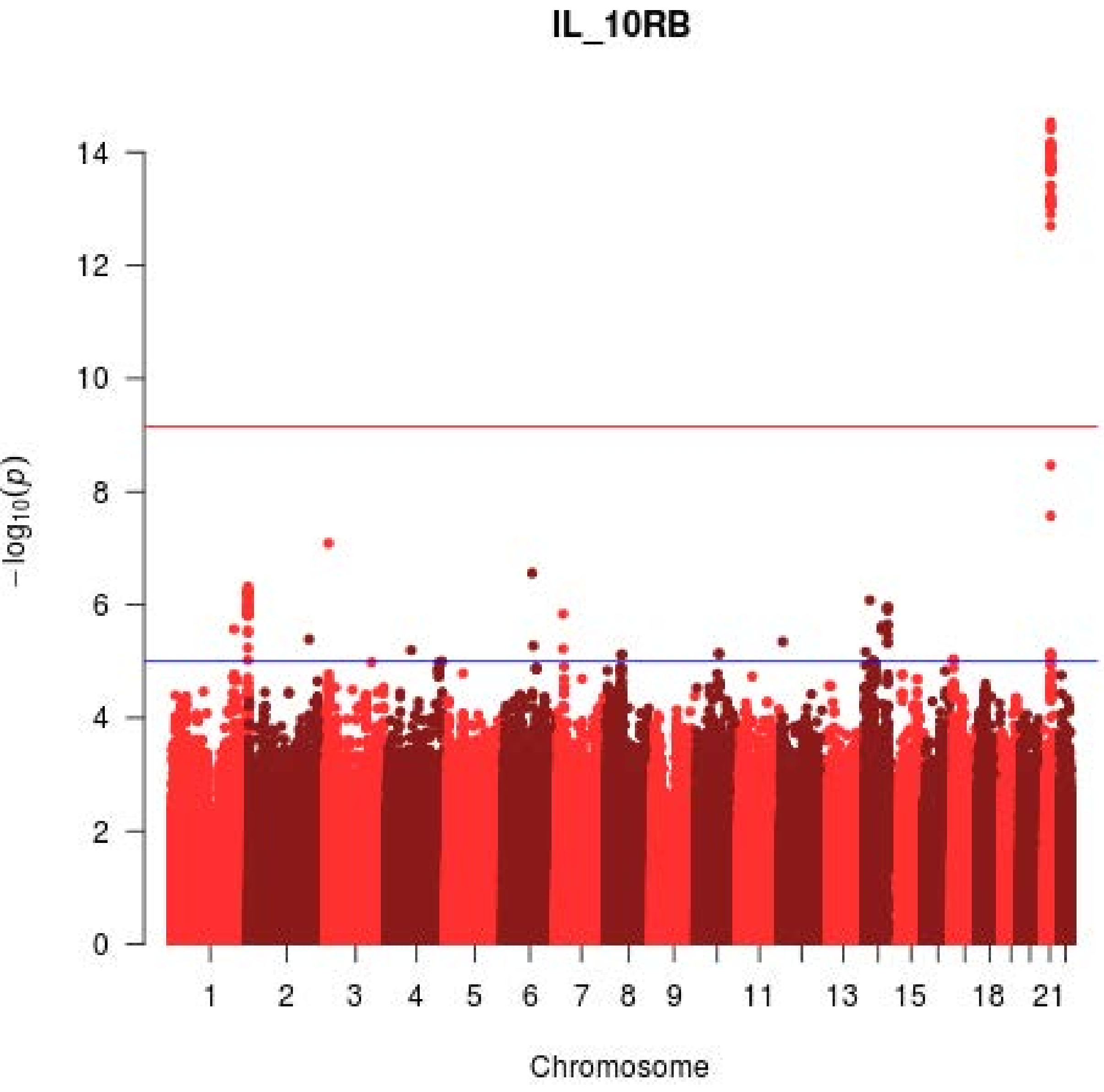
IL_10RB
−log10(p)
0
2
4
6
8
10
12
14
Chromosome
1
2
3
4
5
6
7
8
9
11
13
15
18
21

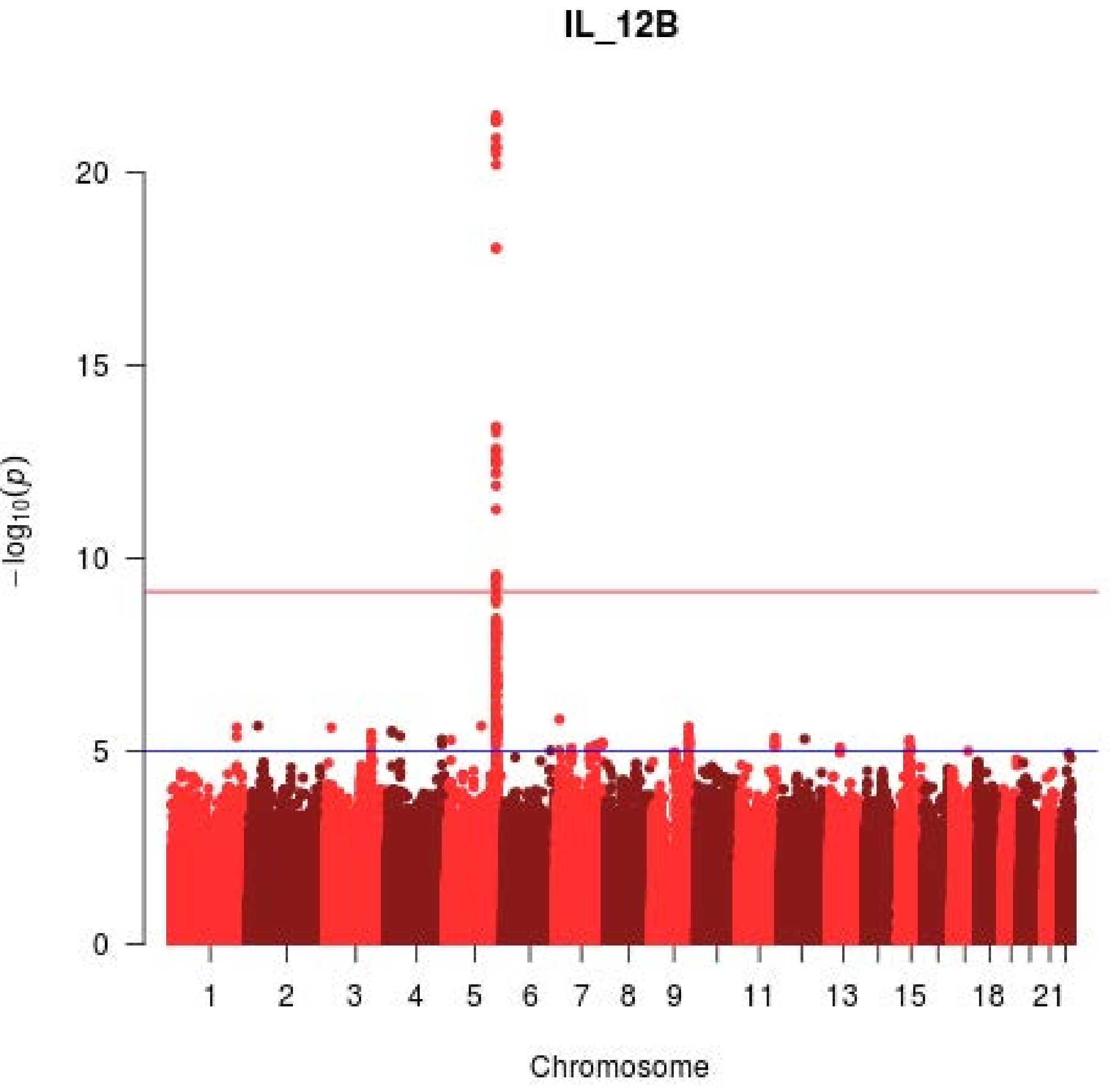
IL_12B
20
15
10
5
0
−log10(p)
1
2
3
4
5
6
7
8
9
11
13
15
18
21
Chromosome

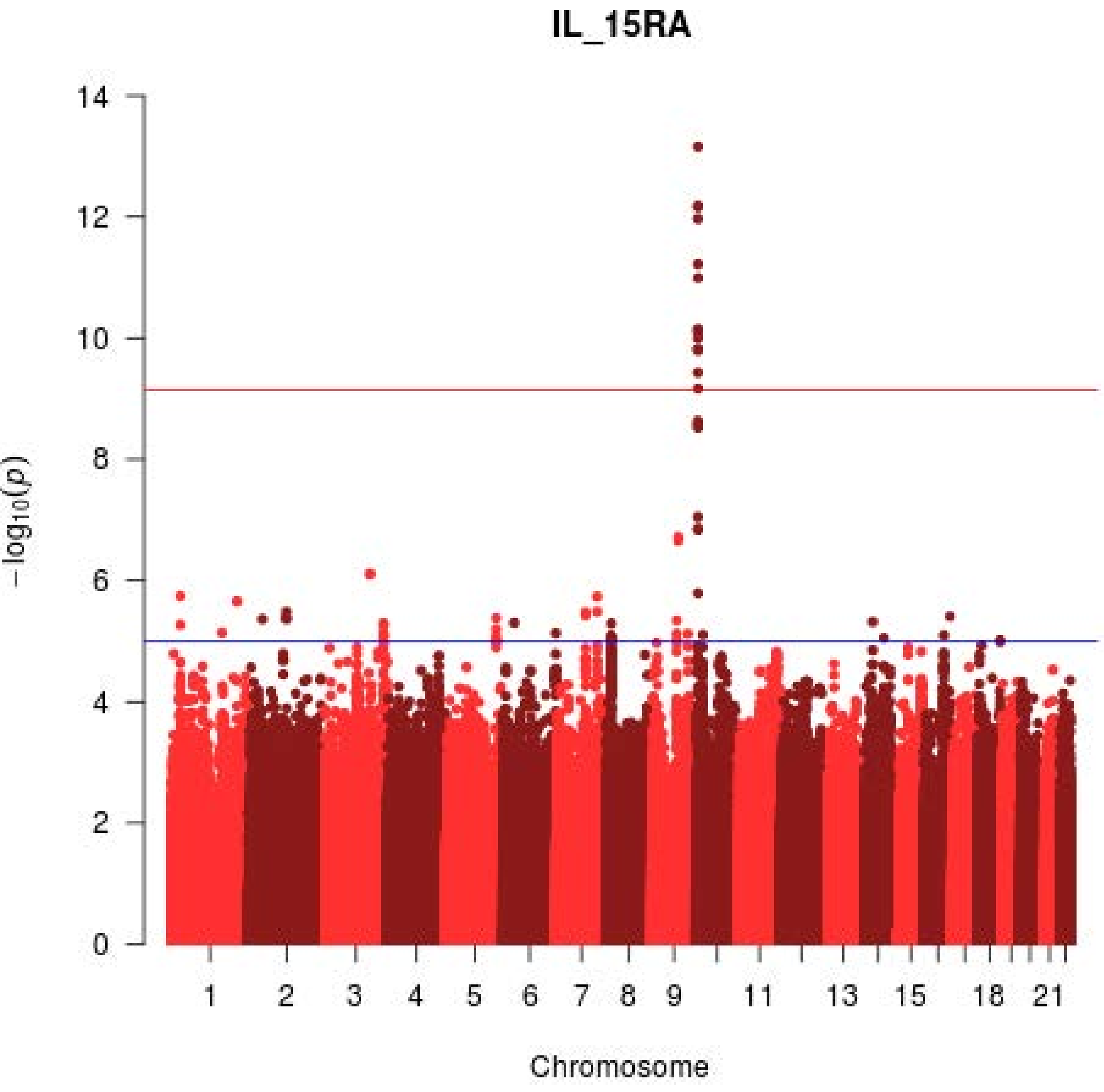
IL_15RA
14
12
10
8
6
4
2
0
$-\log_{10}(p)$
1
2
3
4
5
6
7
8
9
11
13
15
18
21
Chromosome

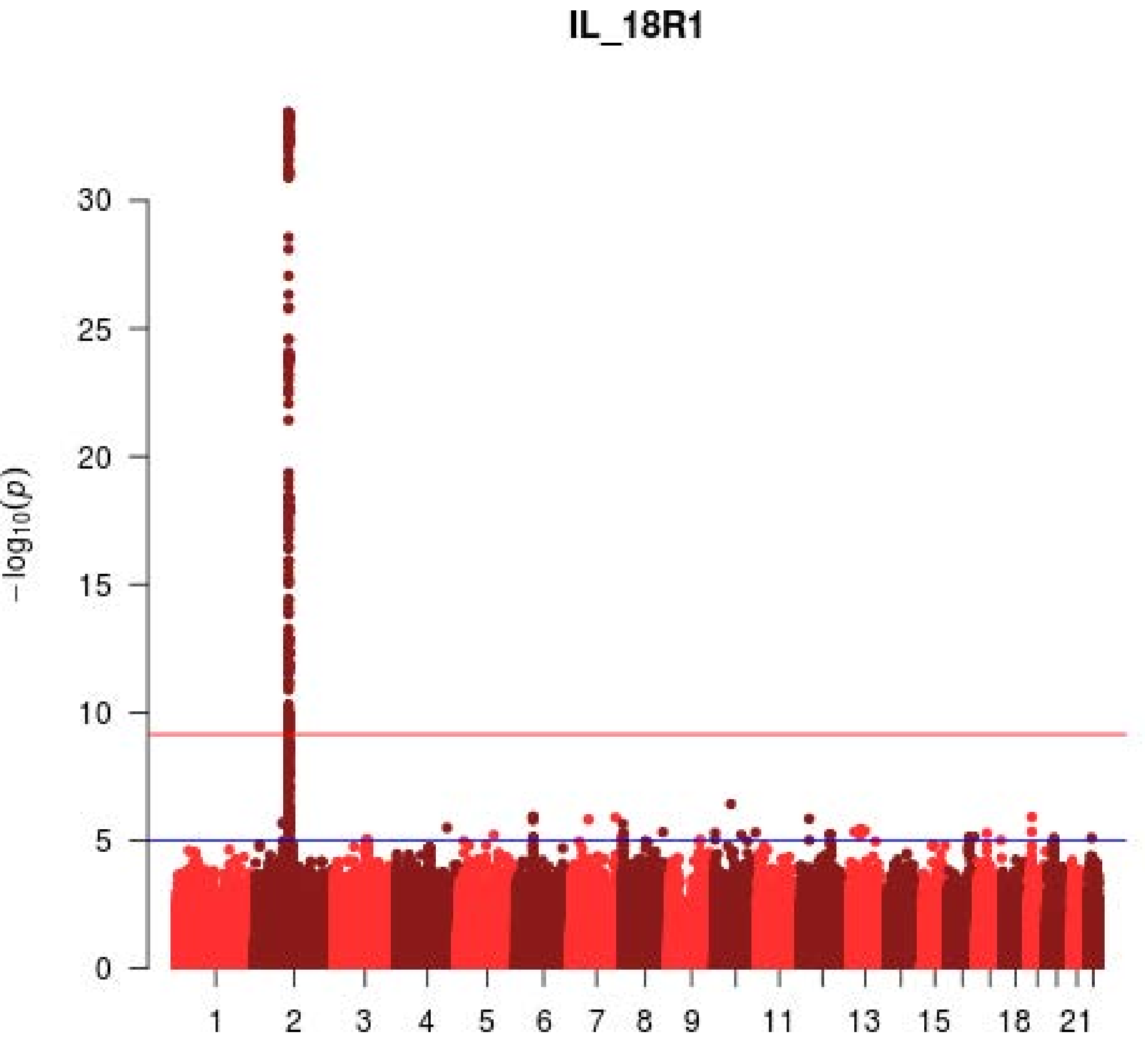
IL_18R1
30
25
20
15
10
5
0
−log10(p)
1
2
3
4
5
6
7
8
9
11
13
15
18
21
Chromosome

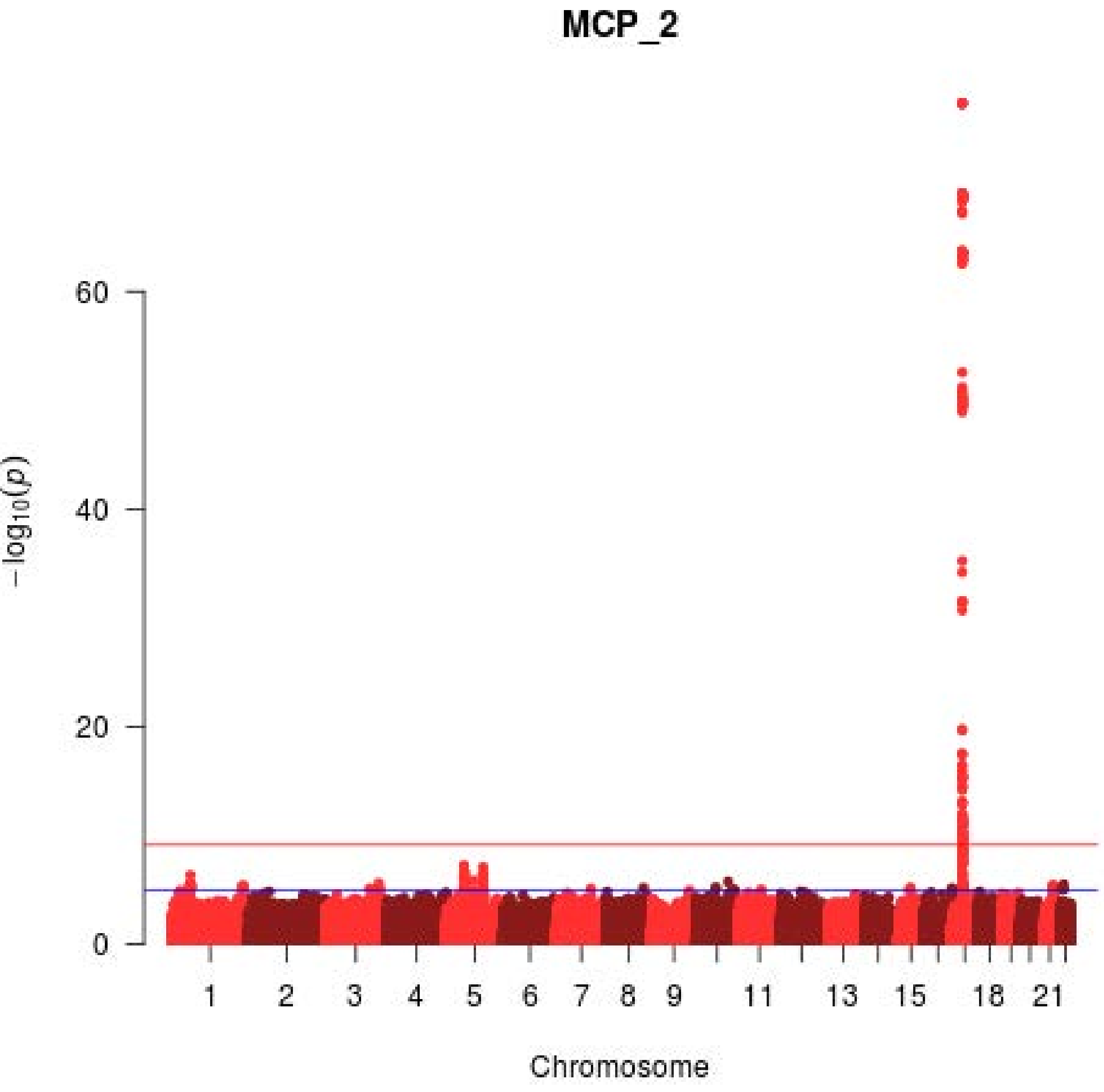

MCP_2
60
40
20
0
−log10(p)
1
2
3
4
5
6
7
8
9
11
13
15
18
21
Chromosome

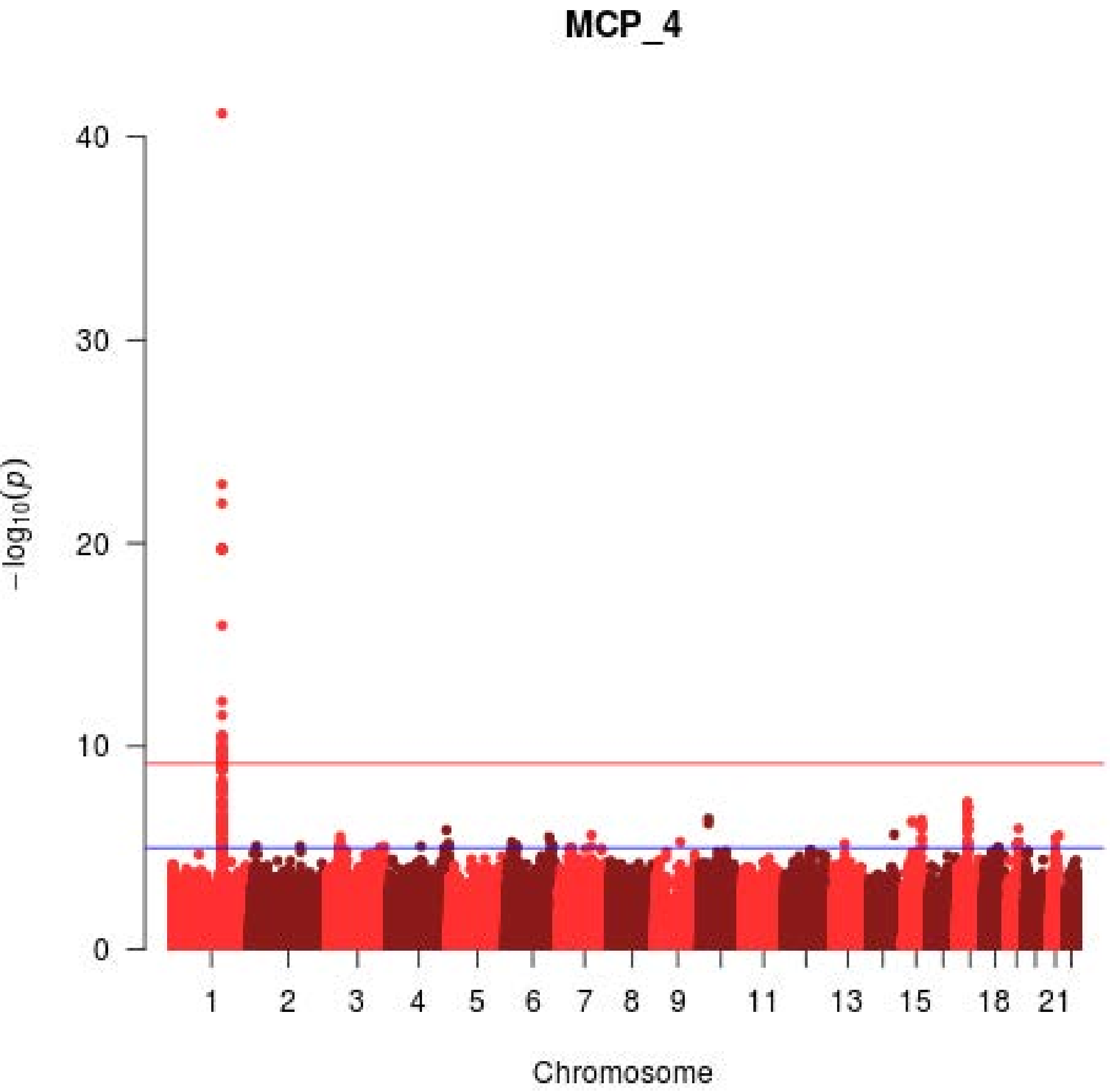

MCP_4
−log10(p)
0
10
20
30
40
1
2
3
4
5
6
7
8
9
11
13
15
18
21
Chromosome

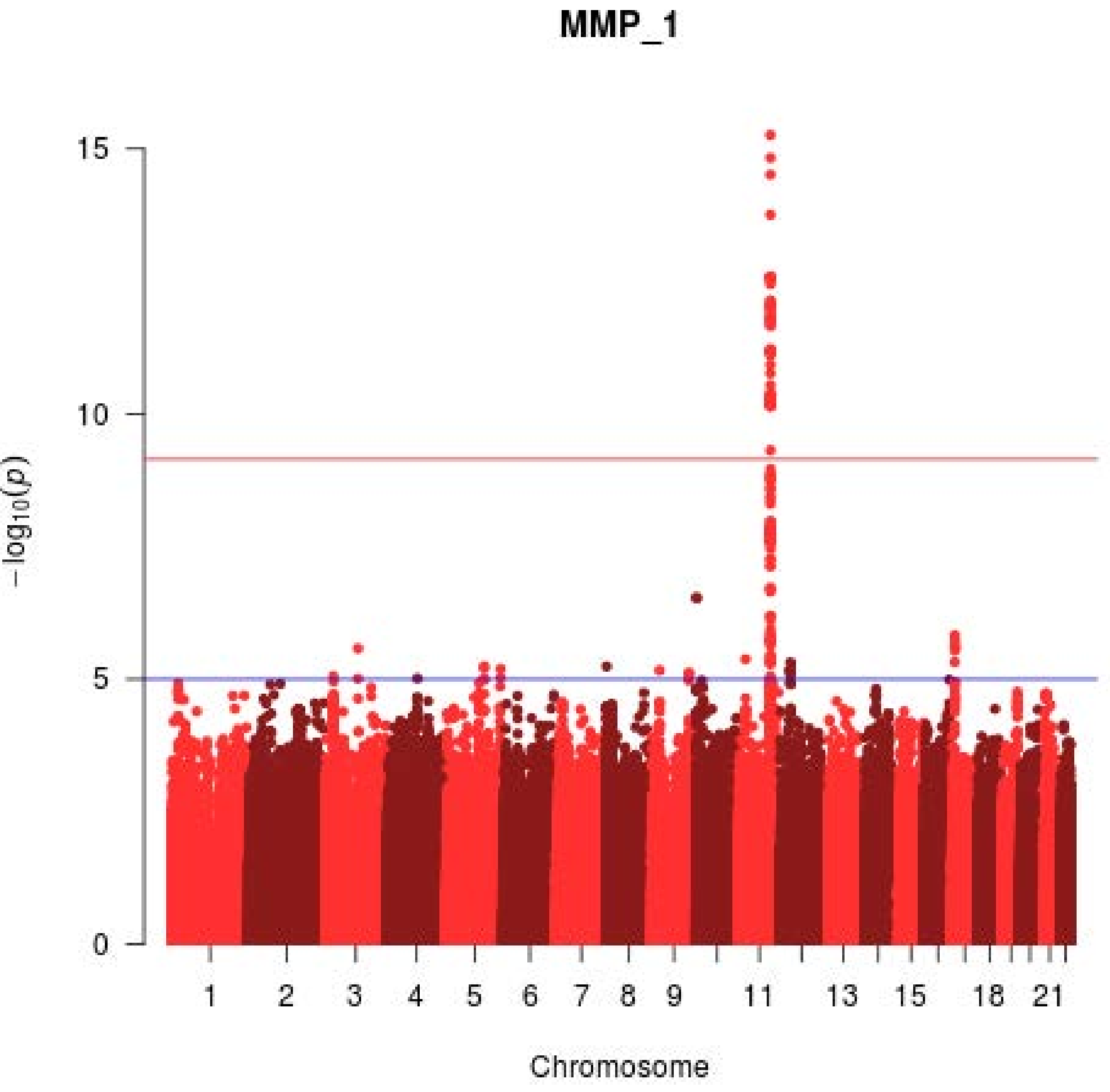
MMP_1
15
10
5
0
$-\log_{10}(p)$
1
2
3
4
5
6
7
8
9
11
13
15
18
21
Chromosome

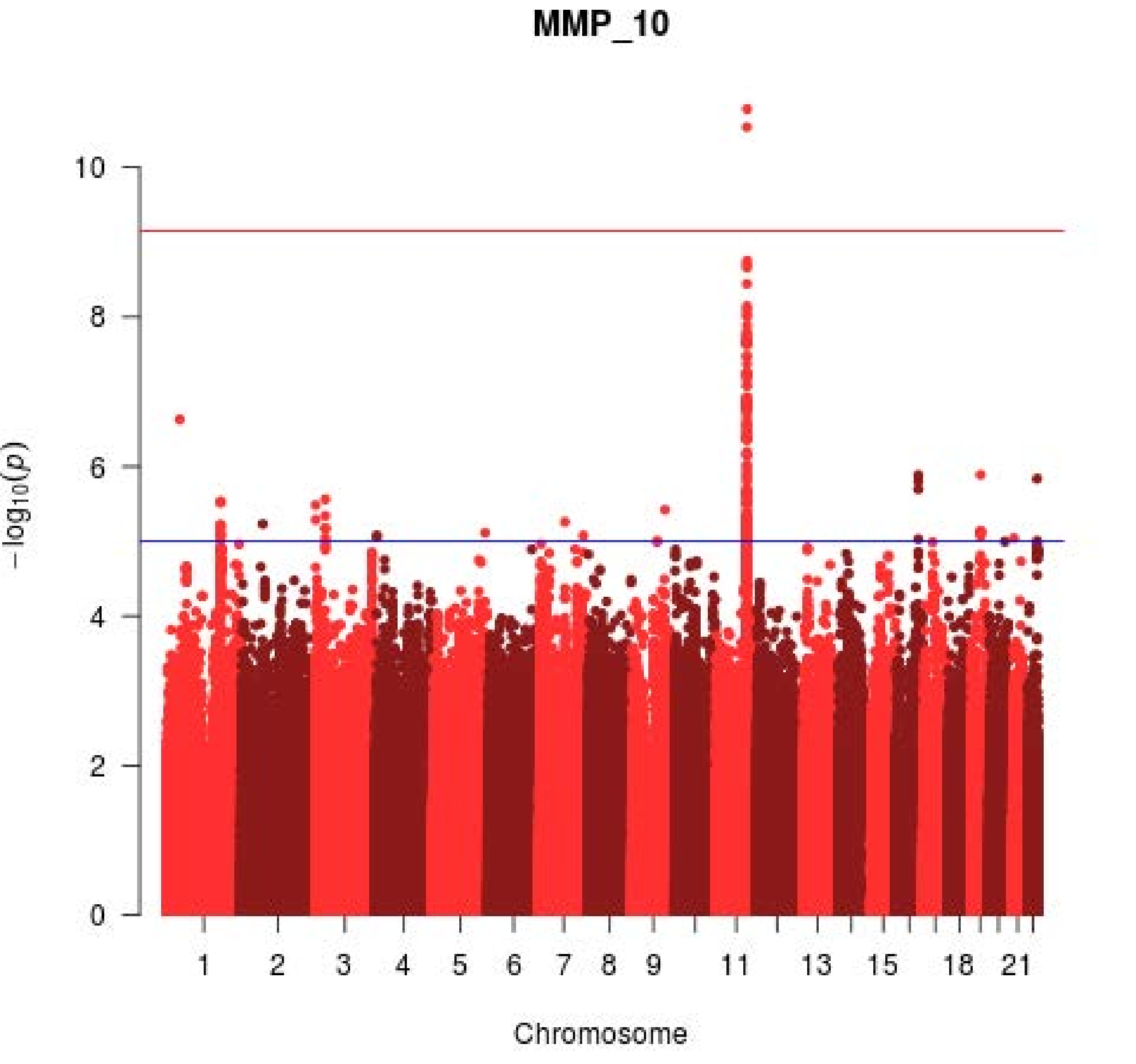
MMP_10
10
8
6
4
2
0
−log10(p)
1
2
3
4
5
6
7
8
9
11
13
15
18
21
Chromosome

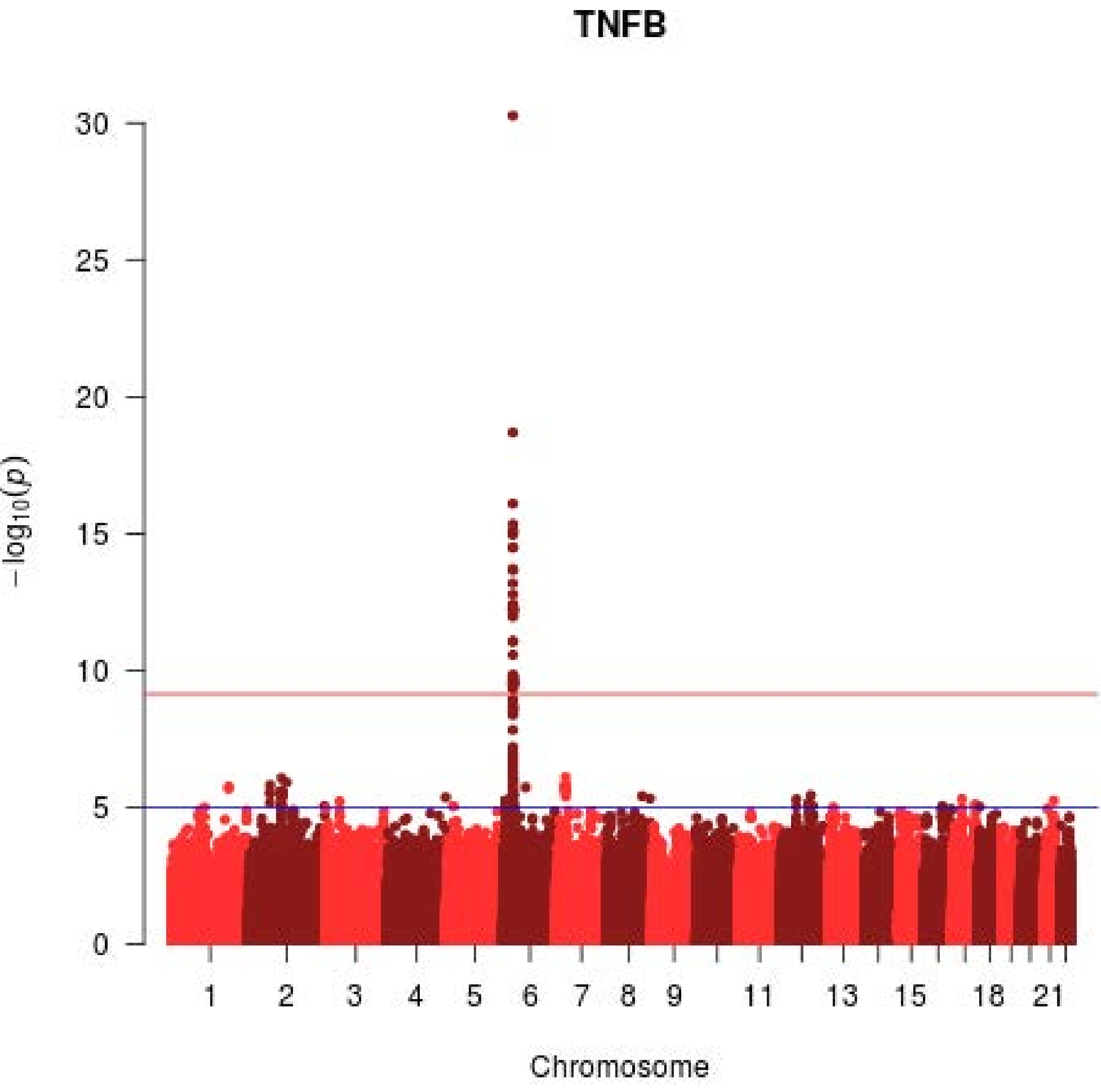

TNFB
30
25
20
15
10
5
0
−log10(p)
1
2
3
4
5
6
7
8
9
11
13
15
18
21
Chromosome

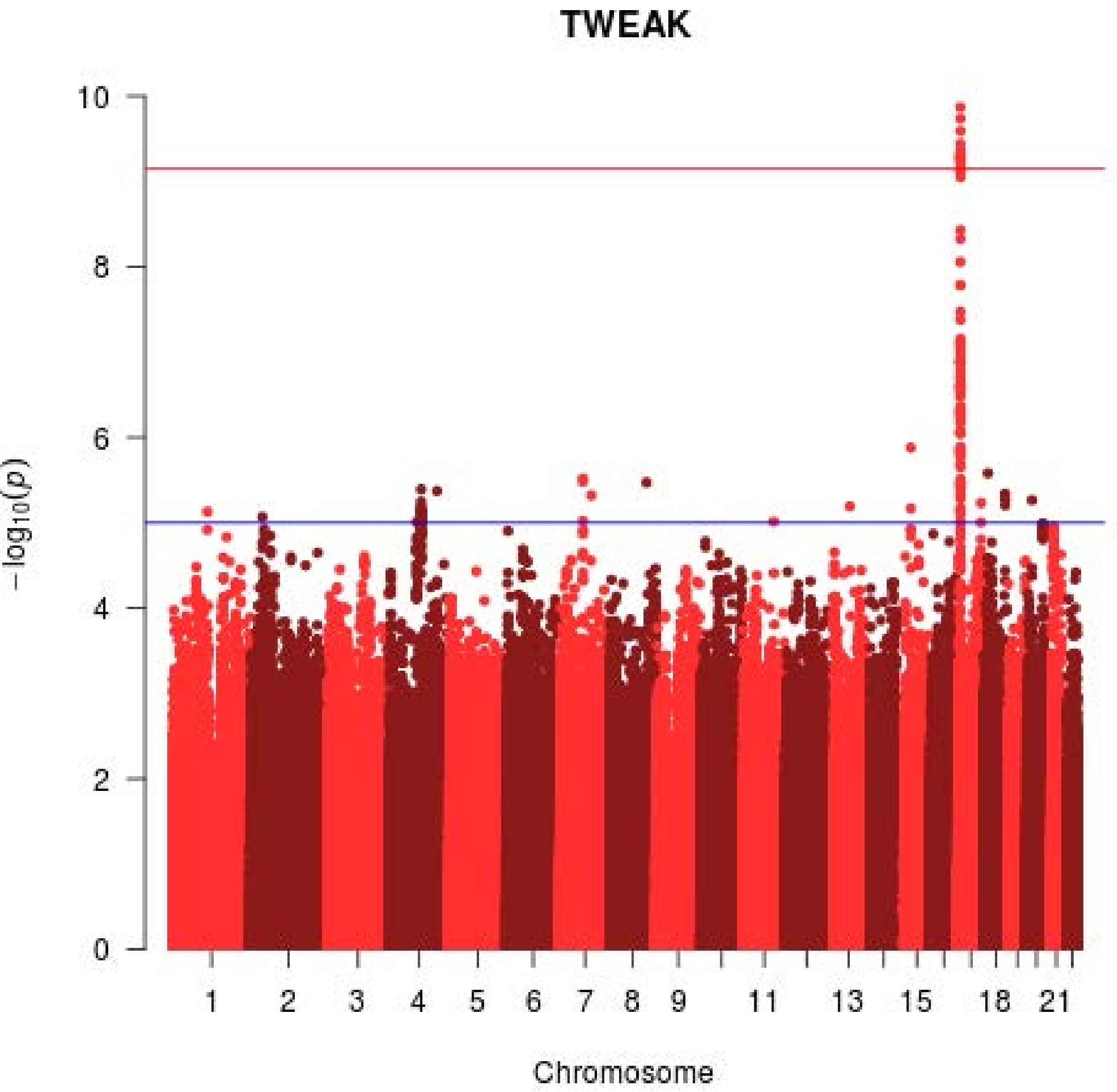
TWEAK
10
8
6
4
2
0
−log10(p)
1
2
3
4
5
6
7
8
9
11
13
15
18
21
Chromosome
