## Supplementary File 4 for "Integrative omics approach to identify the molecular architecture of inflammatory protein levels in healthy older adults"

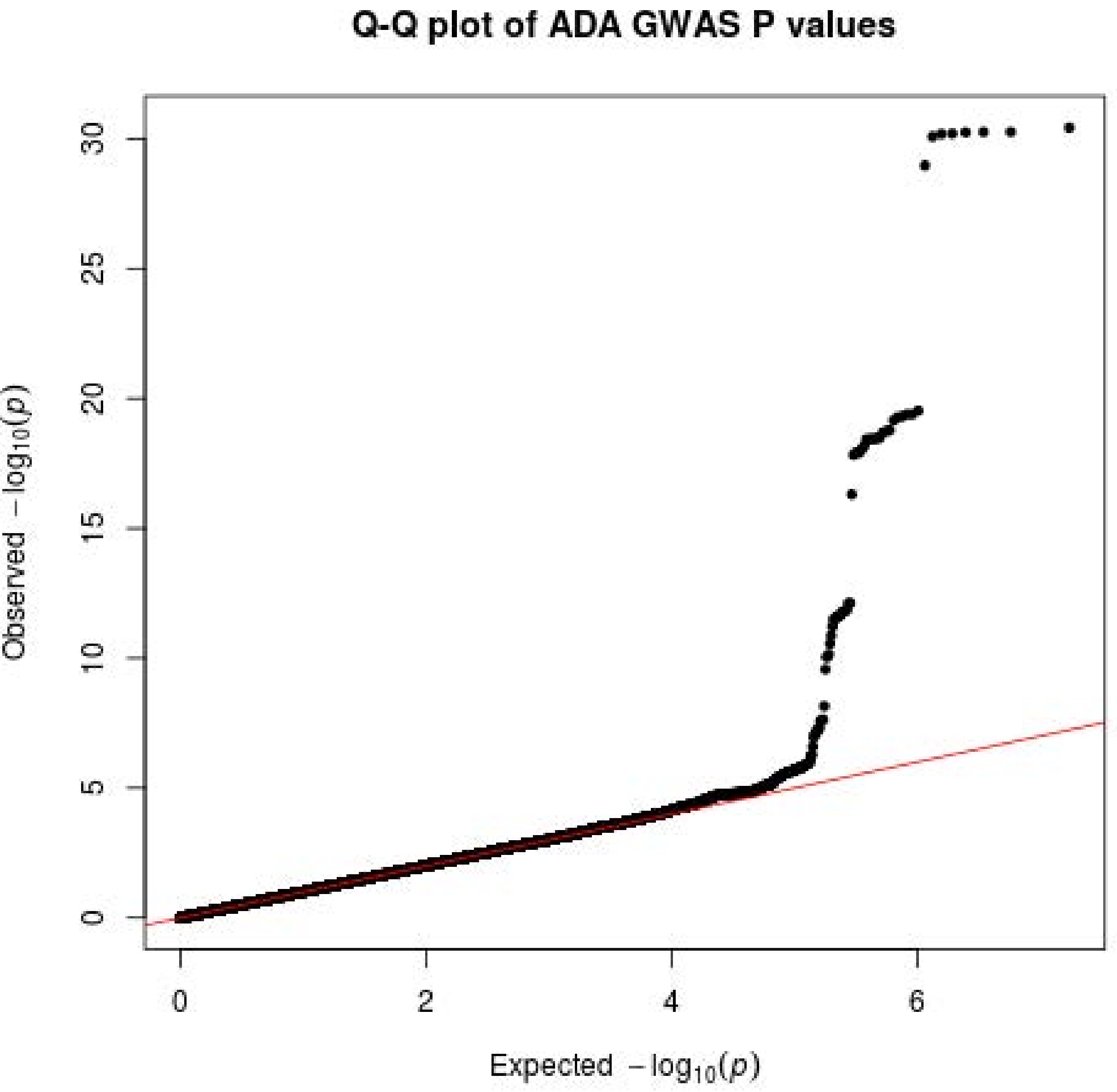

Q-Q plot of ADA GWAS P values
Observed $-\log_{10}(p)$
Expected $-\log_{10}(p)$
0
5
10
15
20
25
30
0
2
4
6

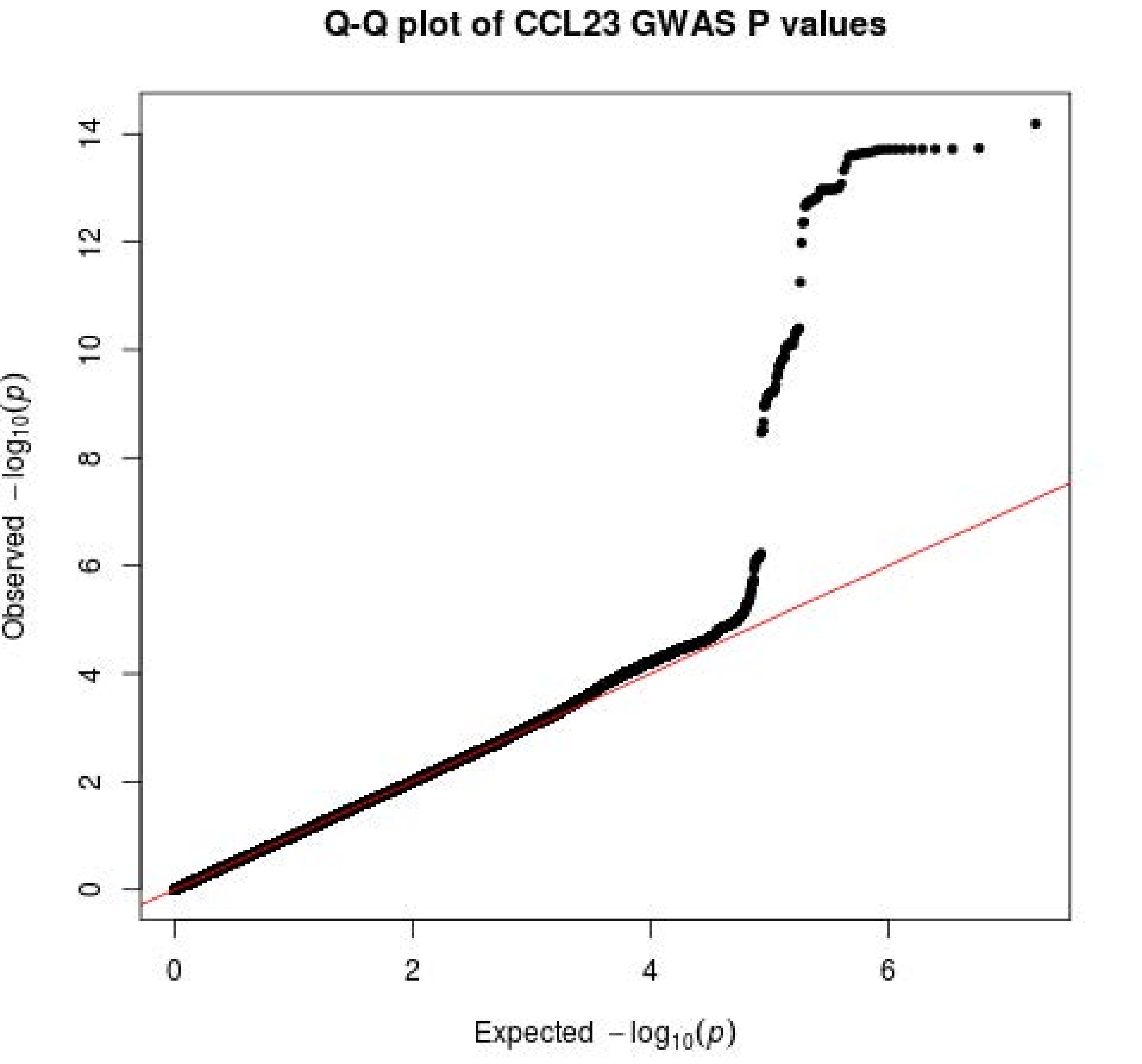

Q-Q plot of CCL23 GWAS P values
Observed −log10(p)
Expected −log10(p)
0
2
4
6
8
10
12
14
0
2
4
6

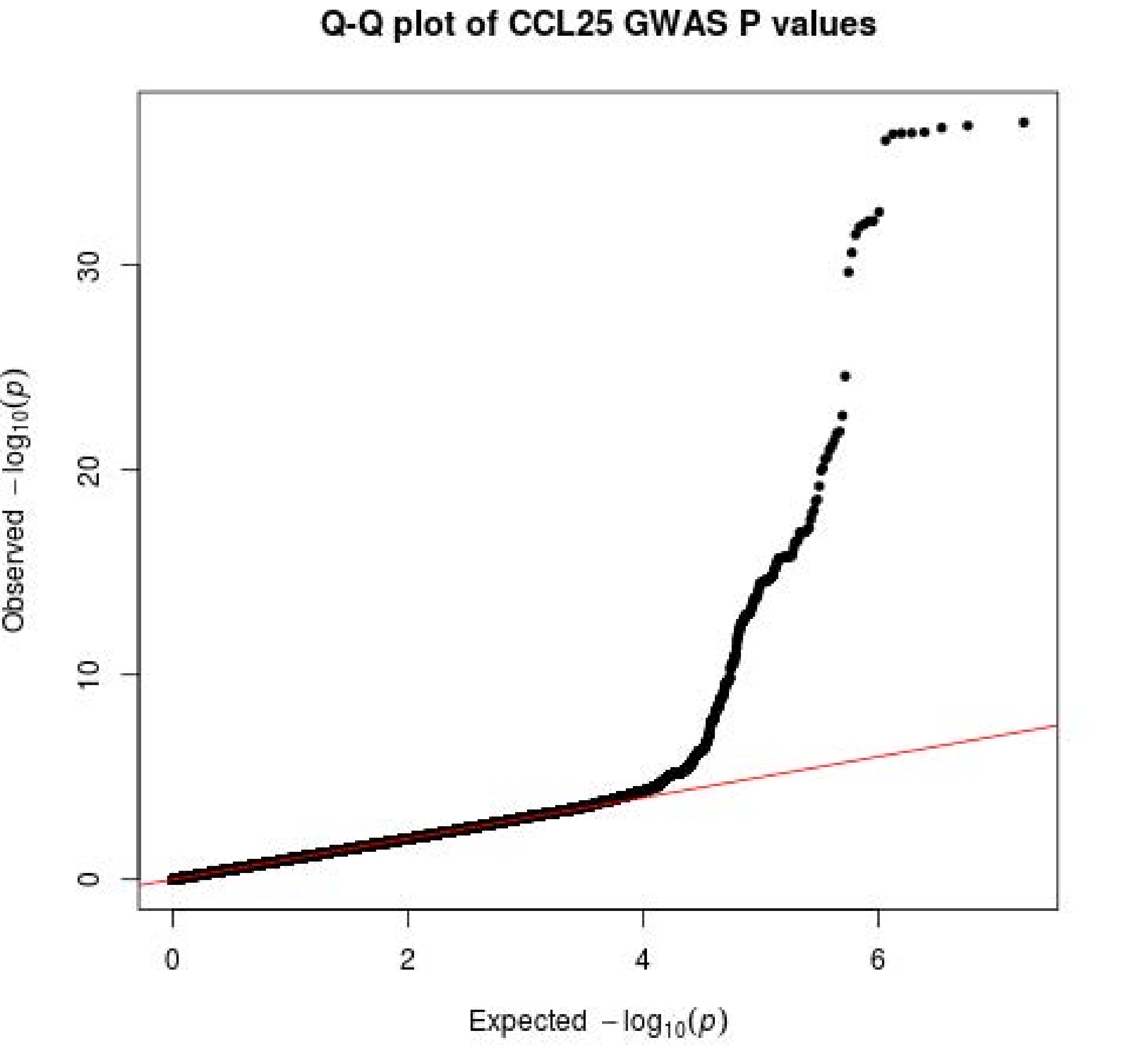
Q-Q plot of CCL25 GWAS P values
Observed $-\log_{10}(p)$
0
10
20
30
0
2
4
6
Expected $-\log_{10}(p)$

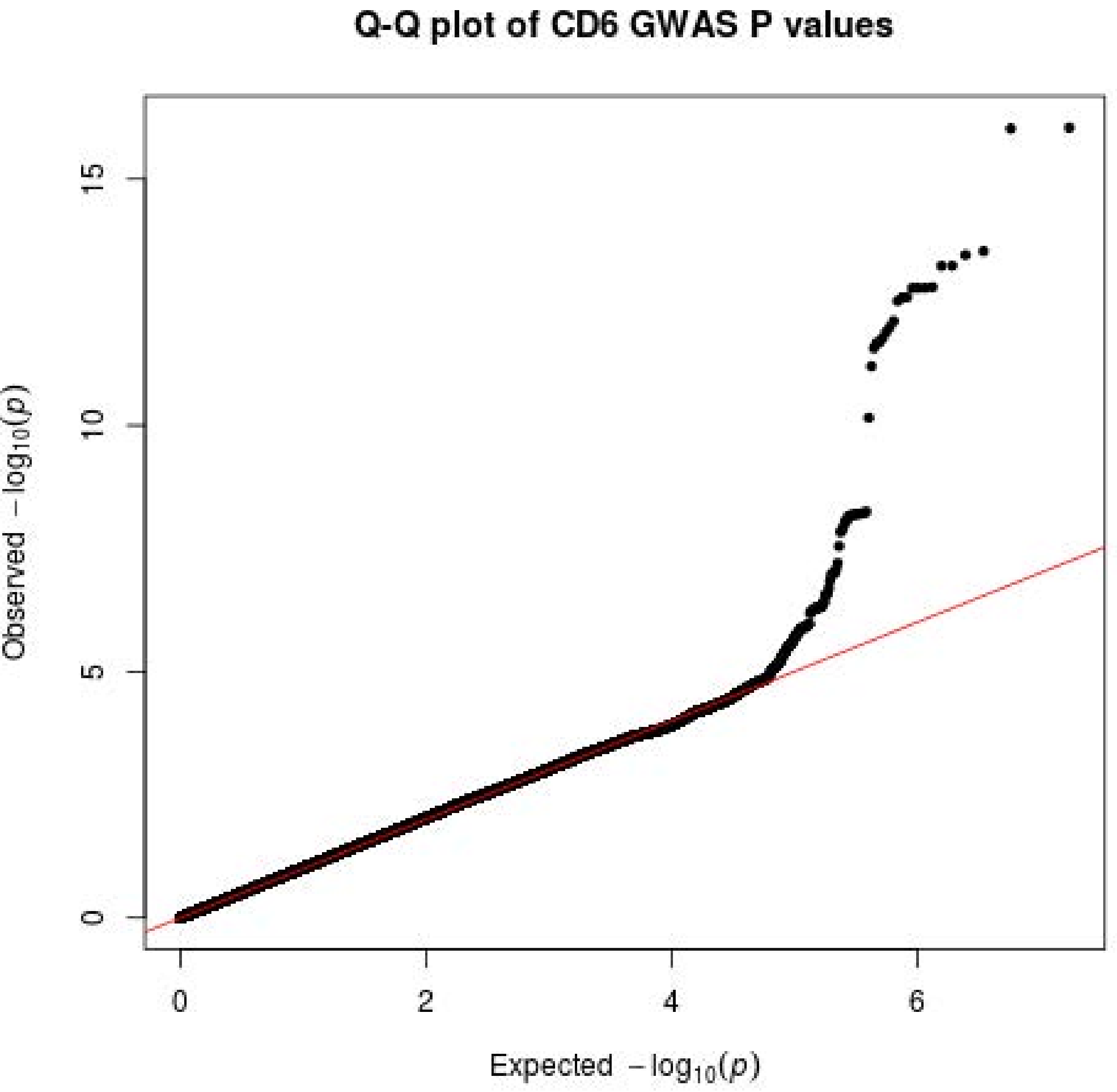

Q-Q plot of CD6 GWAS P values
Observed −log10(p)
Expected −log10(p)
0
5
10
15
0
2
4
6

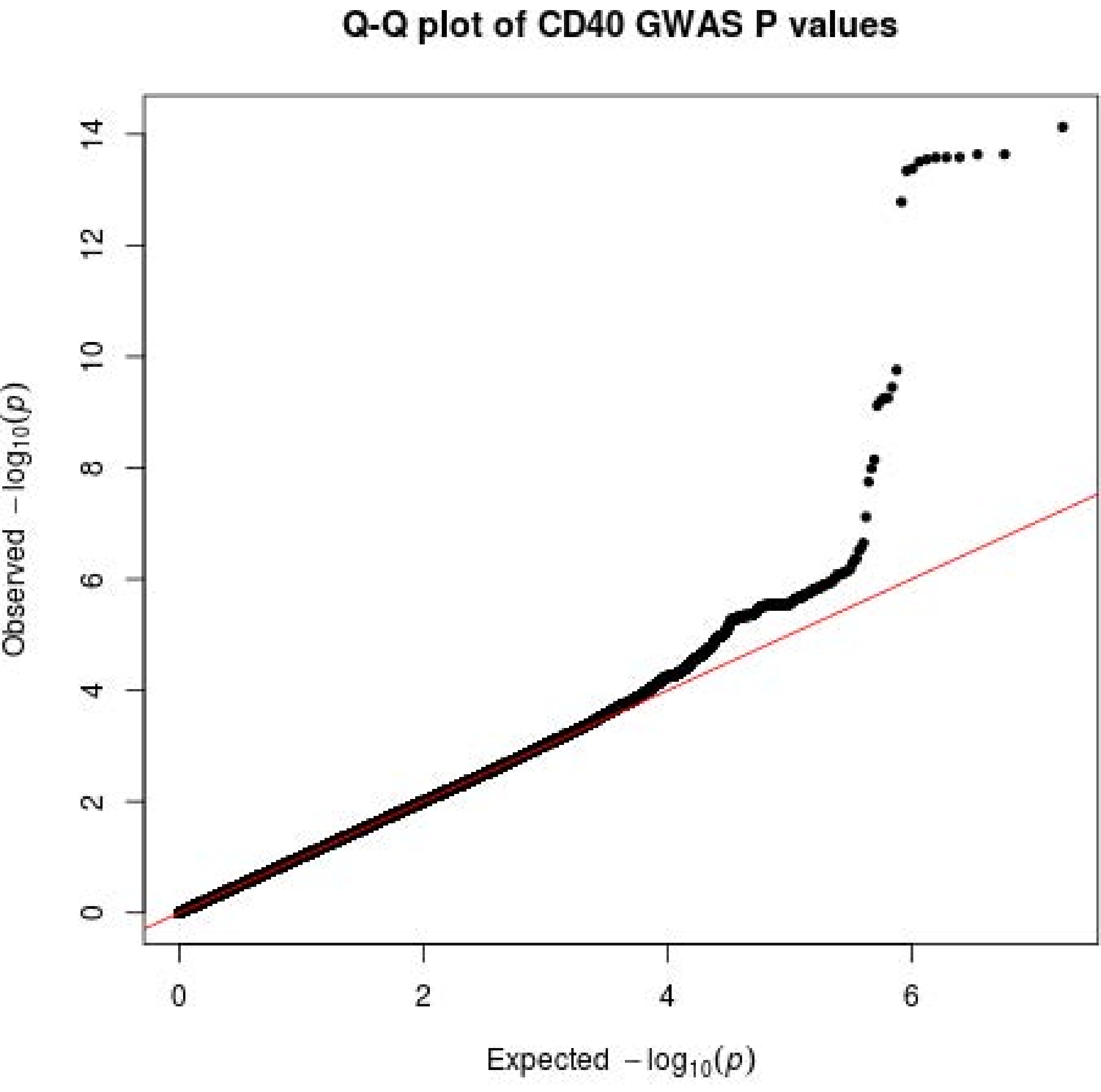
Q-Q plot of CD40 GWAS P values
Observed $-\log_{10}(p)$
Expected $-\log_{10}(p)$
0
2
4
6
8
10
12
14
0
2
4
6

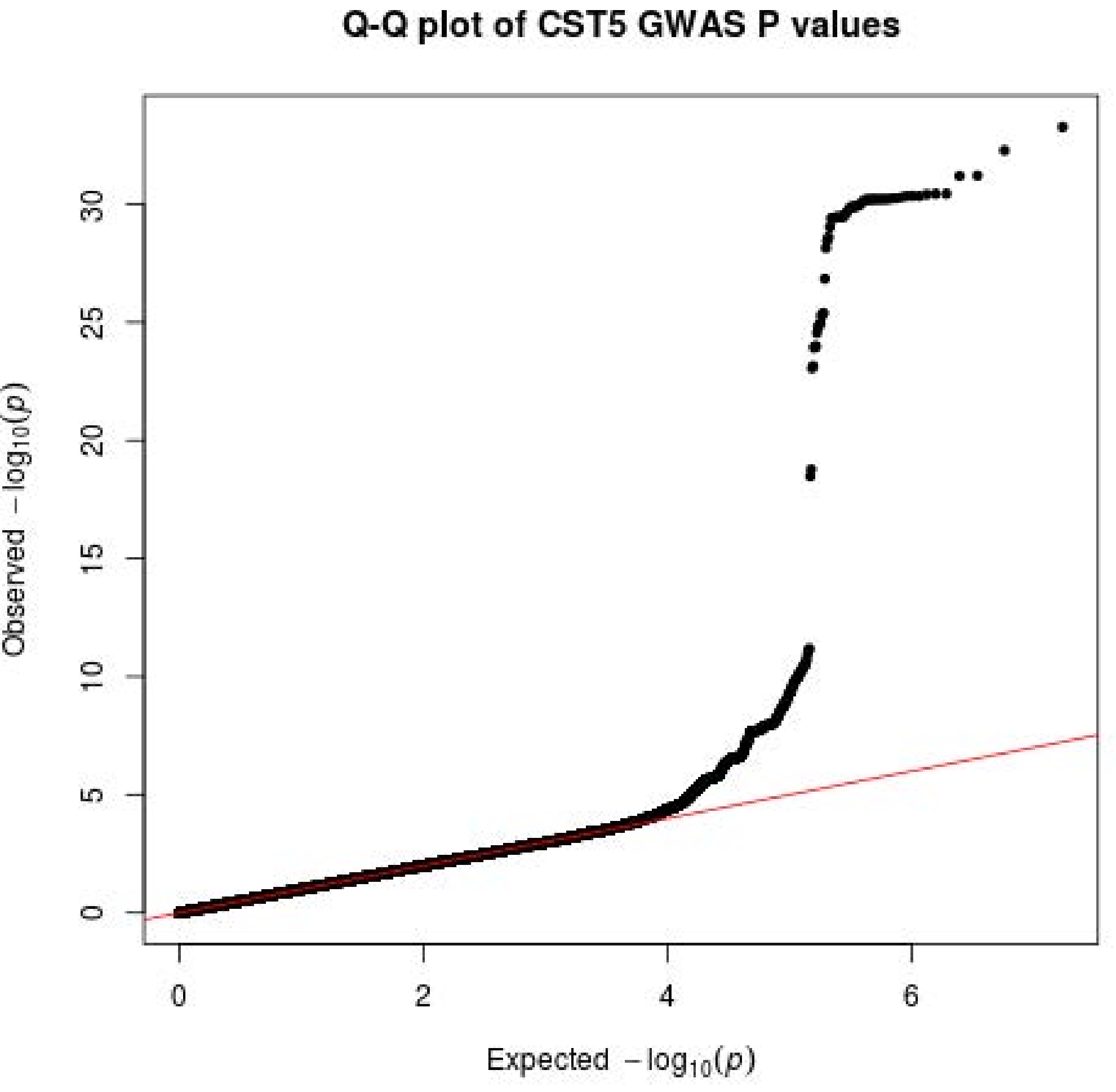
Q-Q plot of CST5 GWAS P values
Observed $-\log_{10}(p)$
Expected $-\log_{10}(p)$
0
5
10
15
20
25
30
0
2
4
6

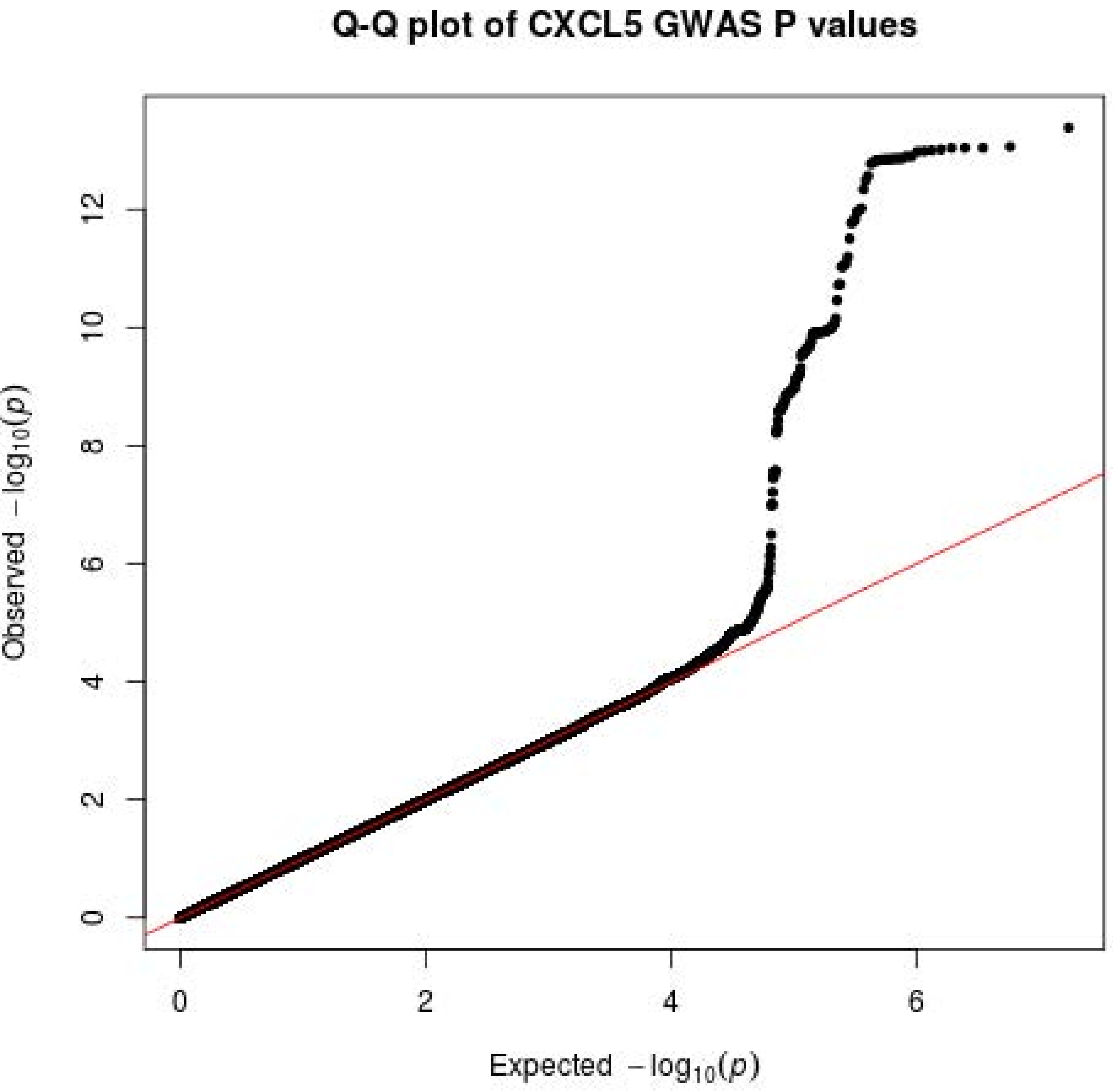
Q-Q plot of CXCL5 GWAS P values
Observed −log10(p)
Expected −log10(p)
0
2
4
6
8
10
12
0
2
4
6

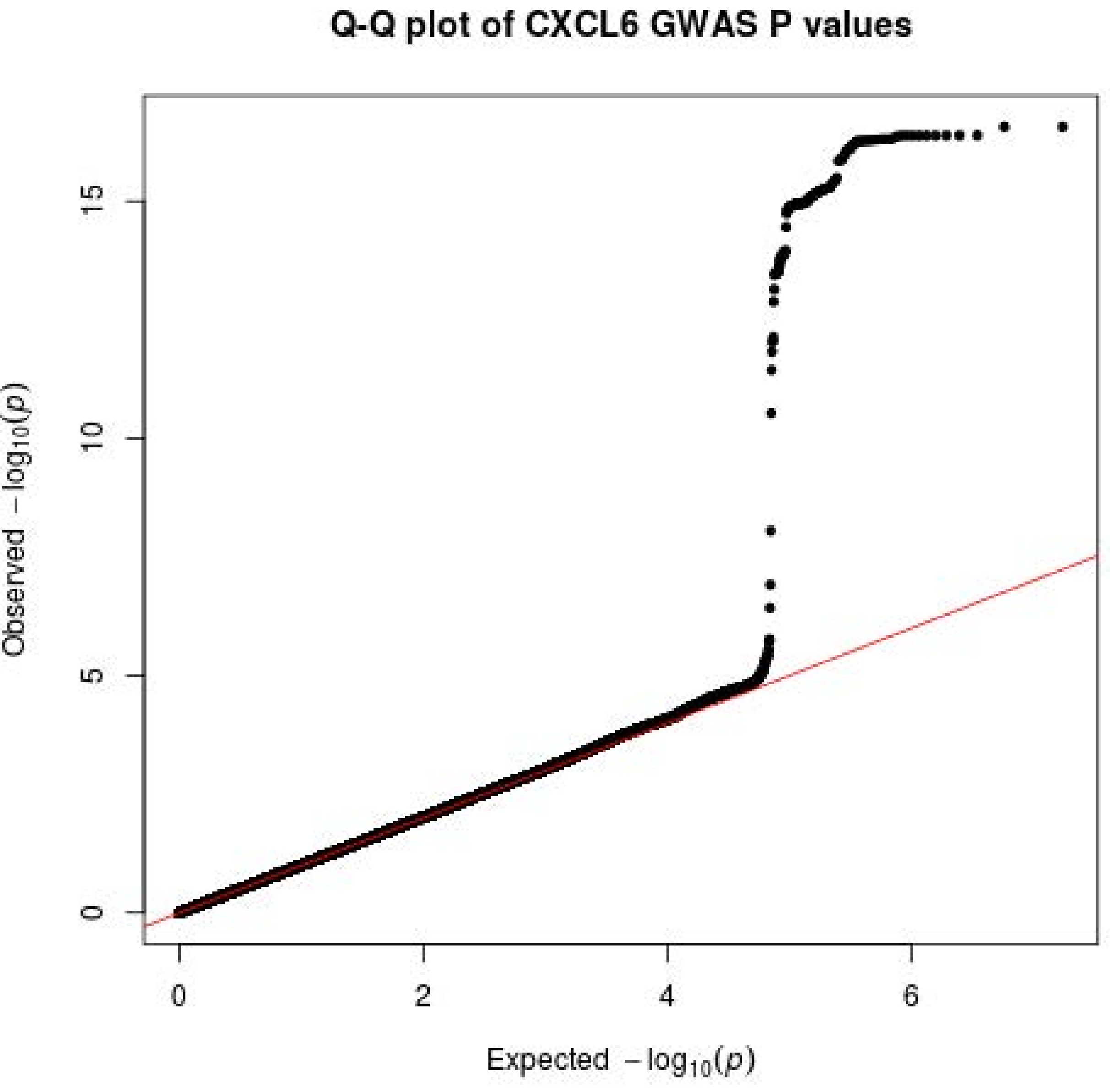
Q-Q plot of CXCL6 GWAS P values
Observed $-\log_{10}(p)$
Expected $-\log_{10}(p)$
0
5
10
15
0
2
4
6

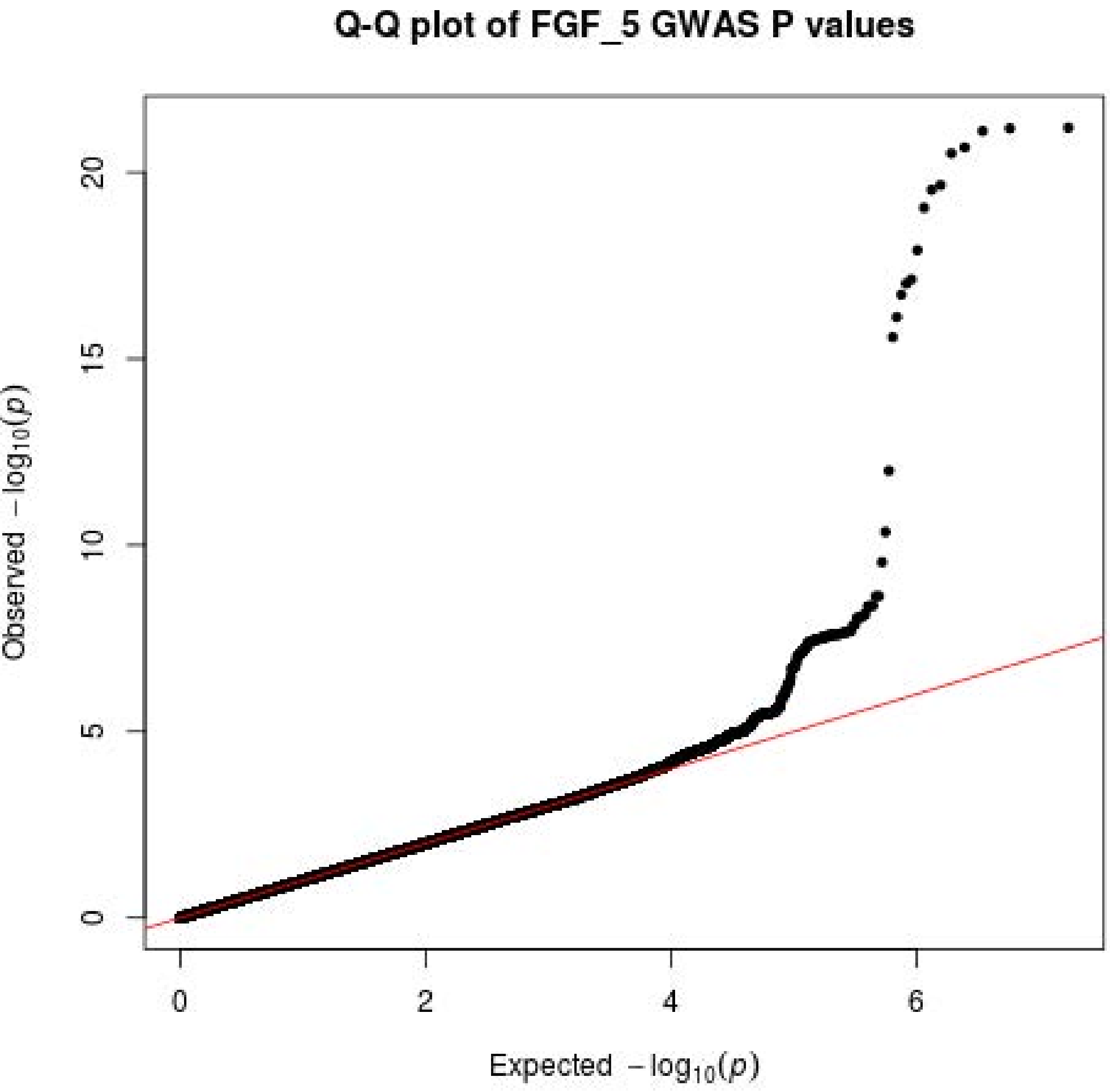
Q-Q plot of FGF_5 GWAS P values
Observed $-\log_{10}(p)$
Expected $-\log_{10}(p)$
0
5
10
15
20
0
2
4
6

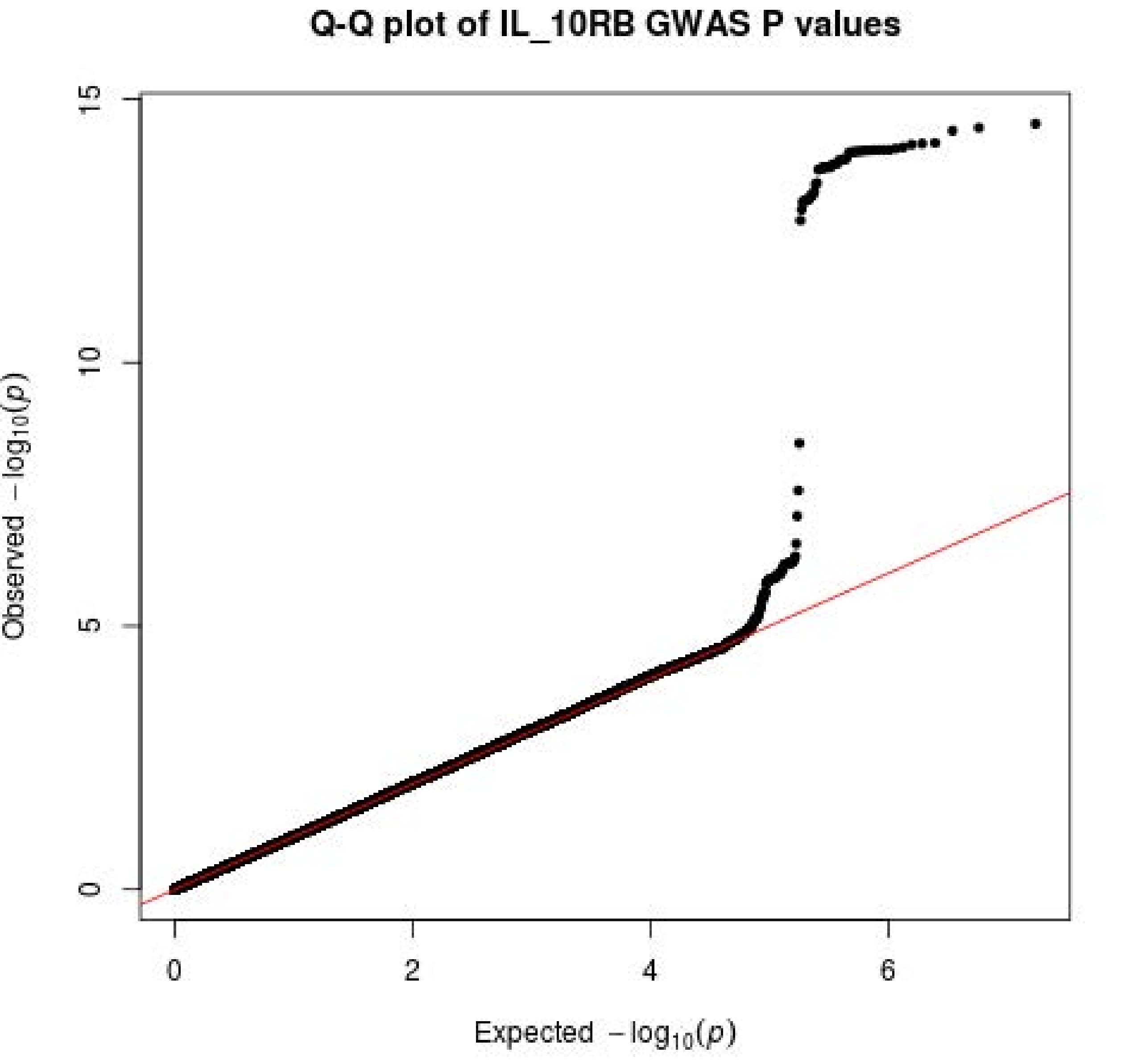
Q-Q plot of IL_10RB GWAS P values
Observed $-\log_{10}(p)$
Expected $-\log_{10}(p)$
0
5
10
15
0
2
4
6

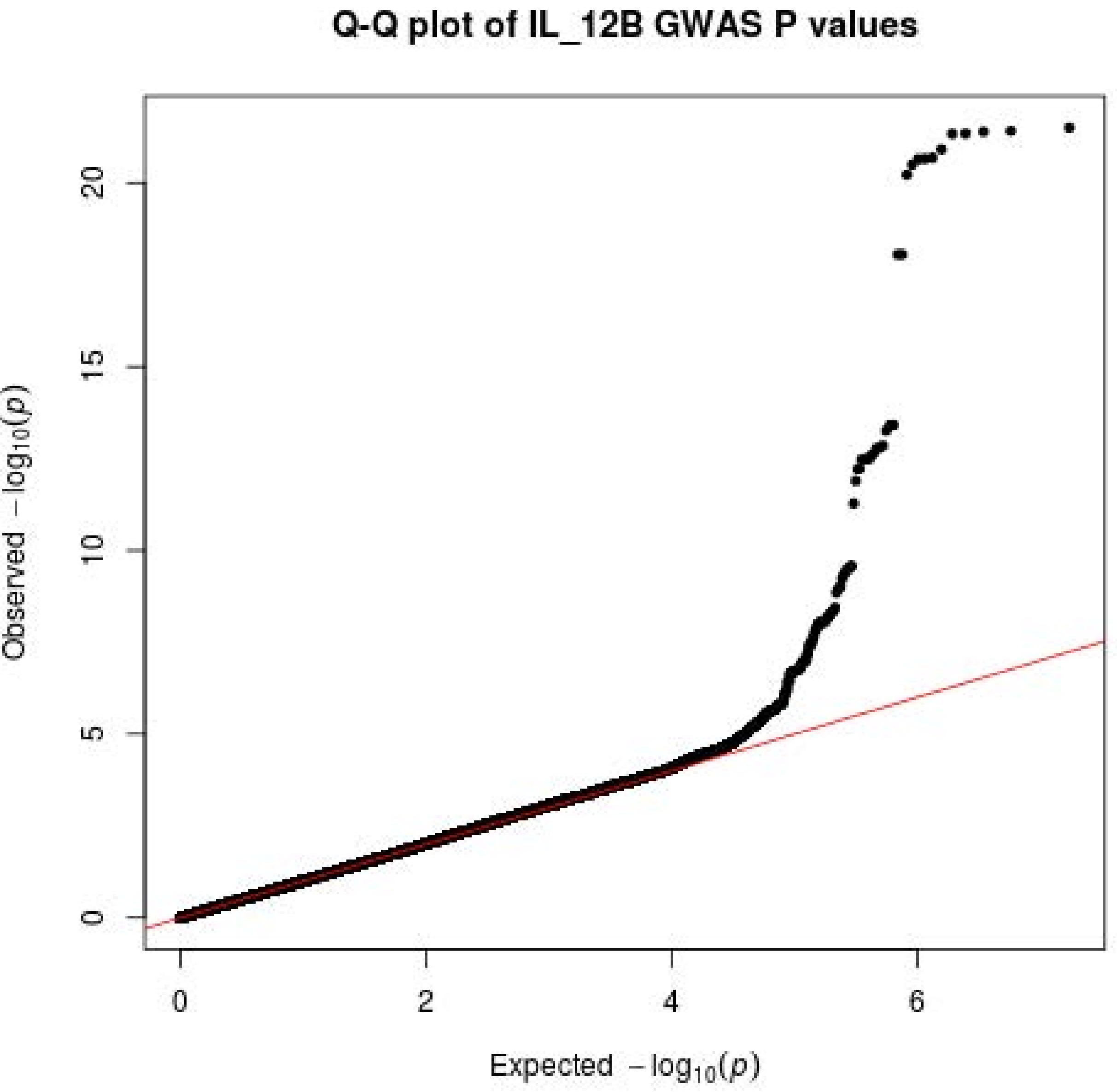

Q-Q plot of IL_12B GWAS P values
Observed $-\log_{10}(p)$
0
5
10
15
20
0
2
4
6
Expected $-\log_{10}(p)$

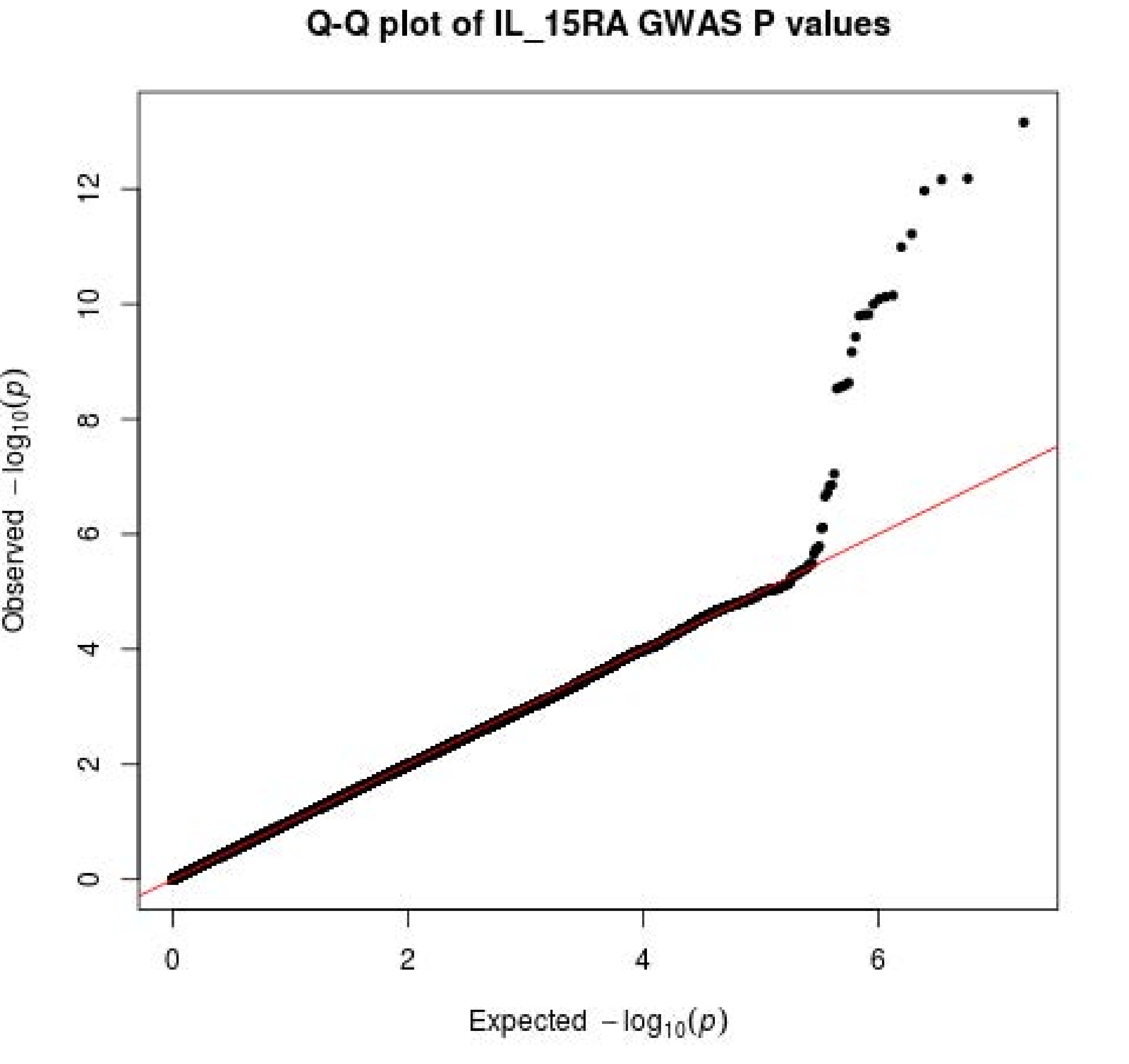
Q-Q plot of IL_15RA GWAS P values
Observed −log10(p)
Expected −log10(p)
0
2
4
6
8
10
12
0
2
4
6

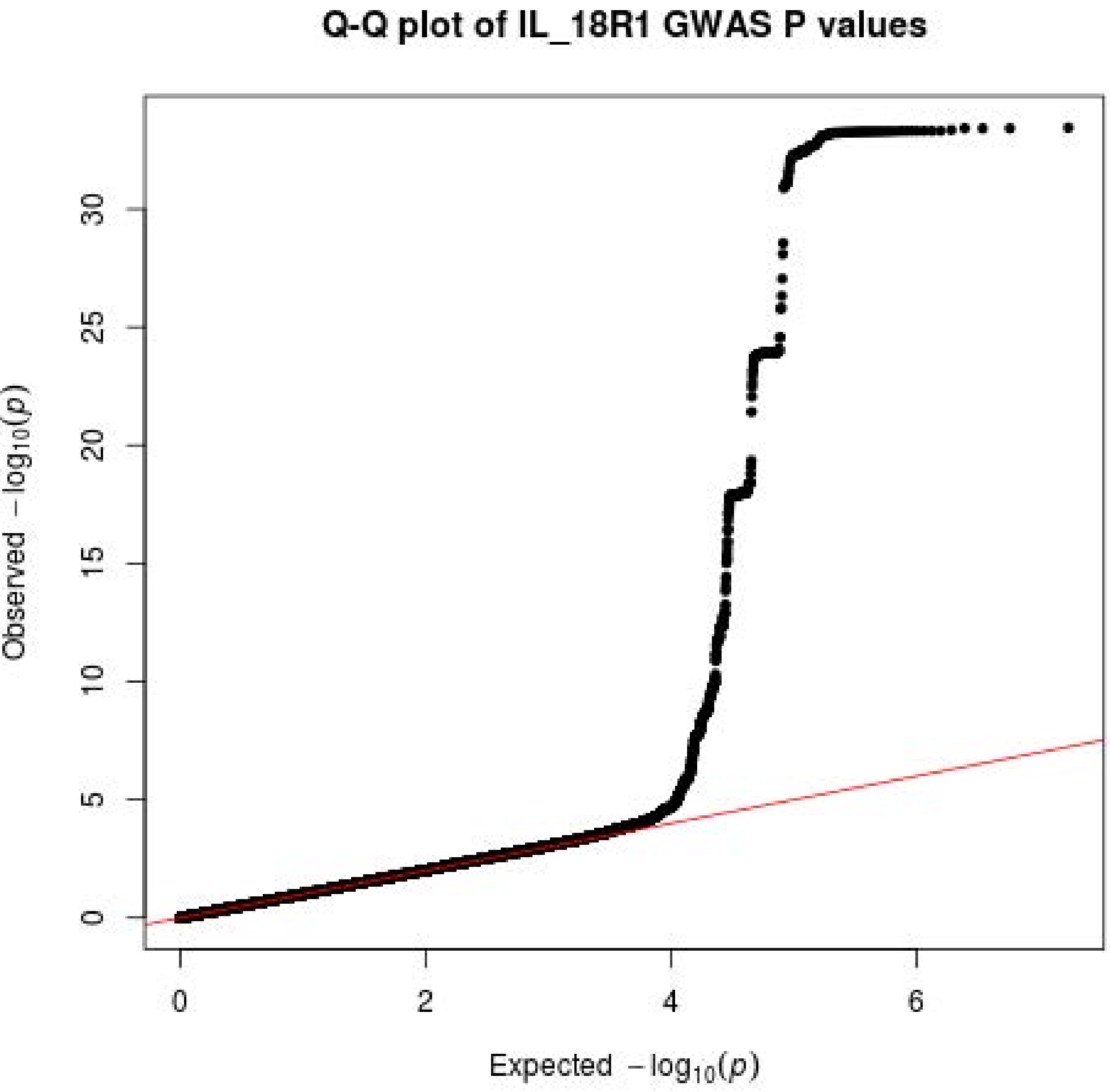

Q-Q plot of IL_18R1 GWAS P values
Observed $-\log_{10}(p)$
Expected $-\log_{10}(p)$
0
5
10
15
20
25
30
0
2
4
6

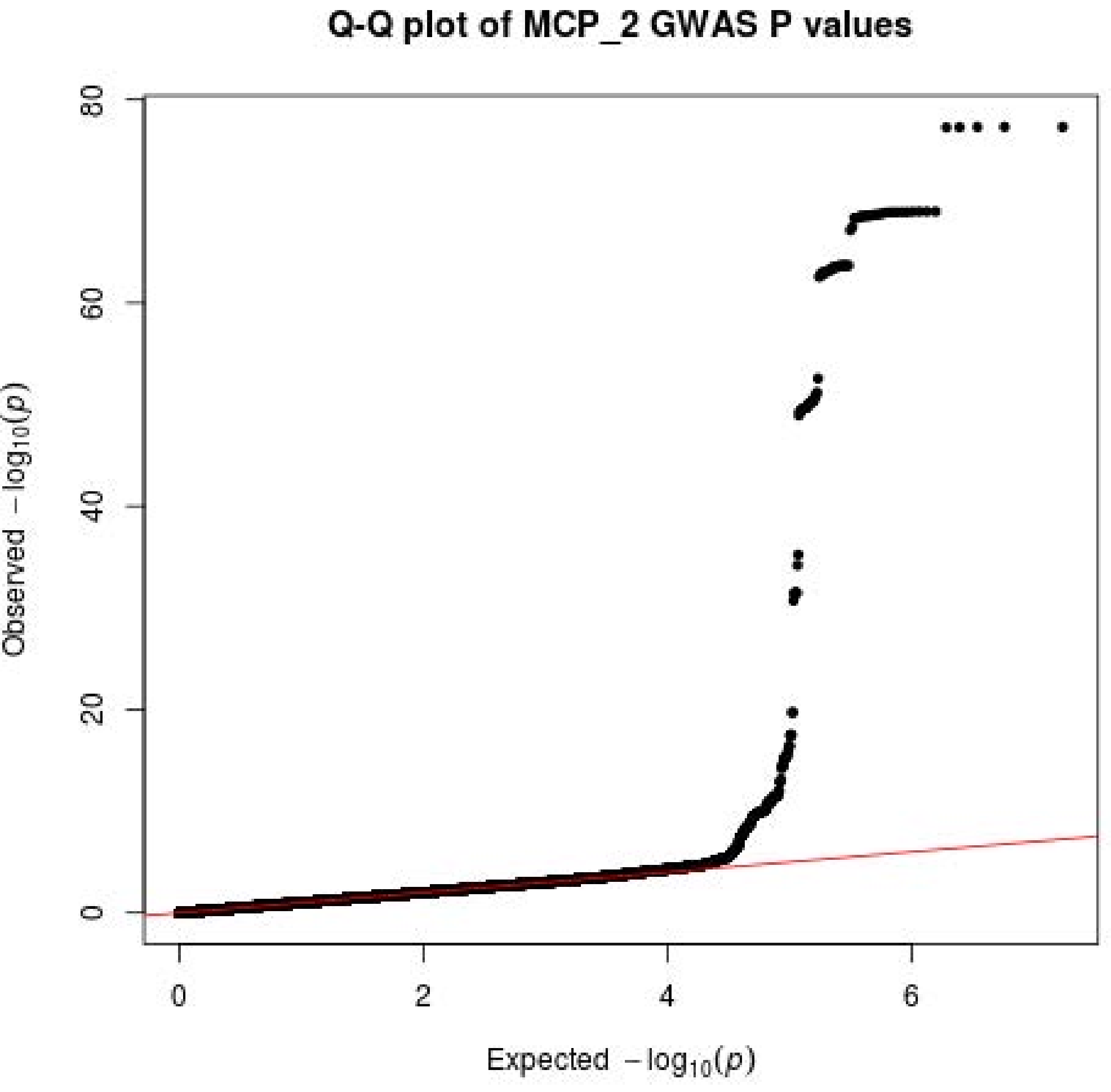
Q-Q plot of MCP_2 GWAS P values
Observed −log10(p)
Expected −log10(p)
0
20
40
60
80
0
2
4
6

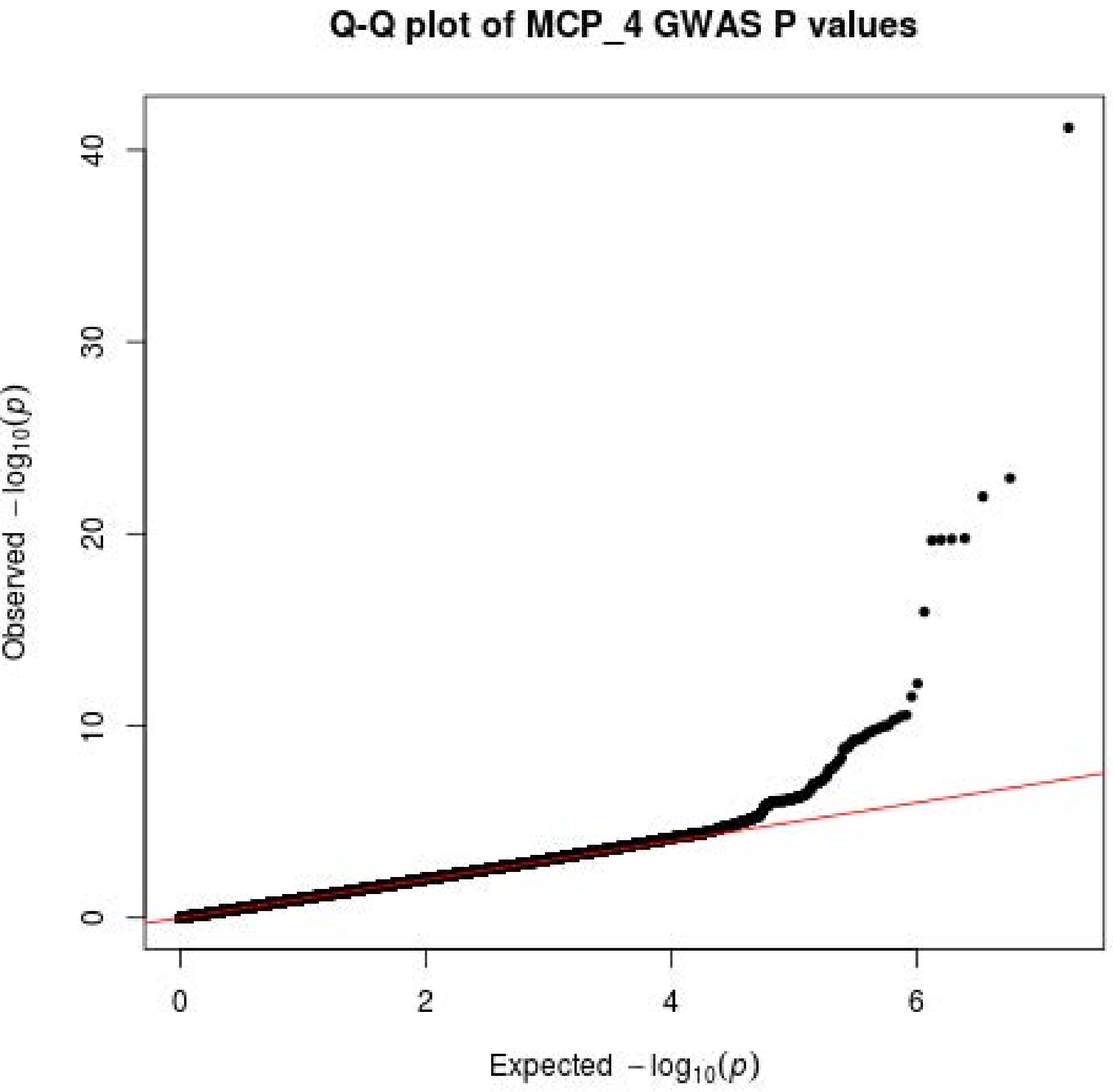

Q-Q plot of MCP_4 GWAS P values
Observed −log10(p)
Expected −log10(p)
0
10
20
30
40
0
2
4
6

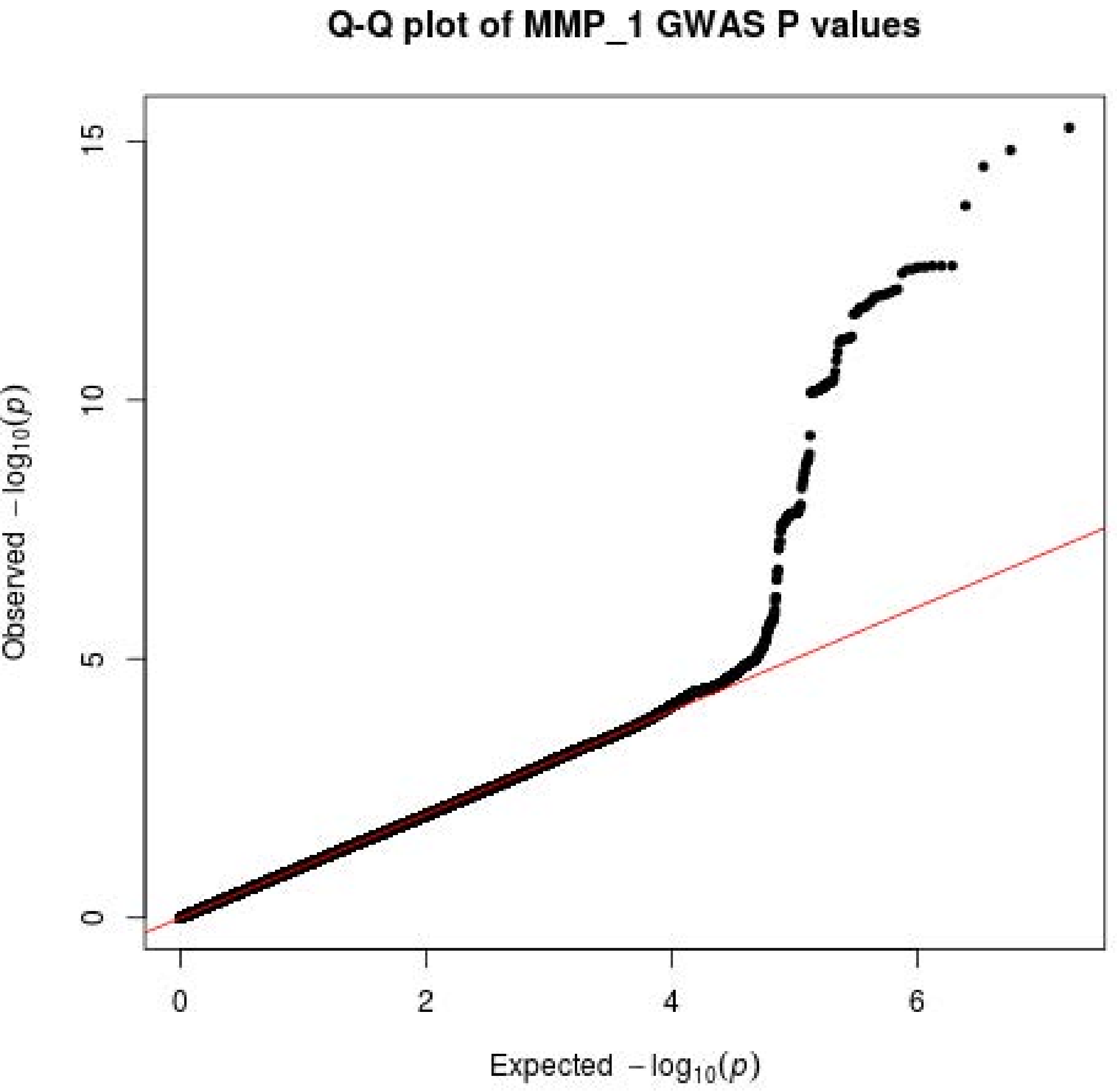
Q-Q plot of MMP_1 GWAS P values
Observed $-\log_{10}(p)$
Expected $-\log_{10}(p)$
0
5
10
15
0
2
4
6

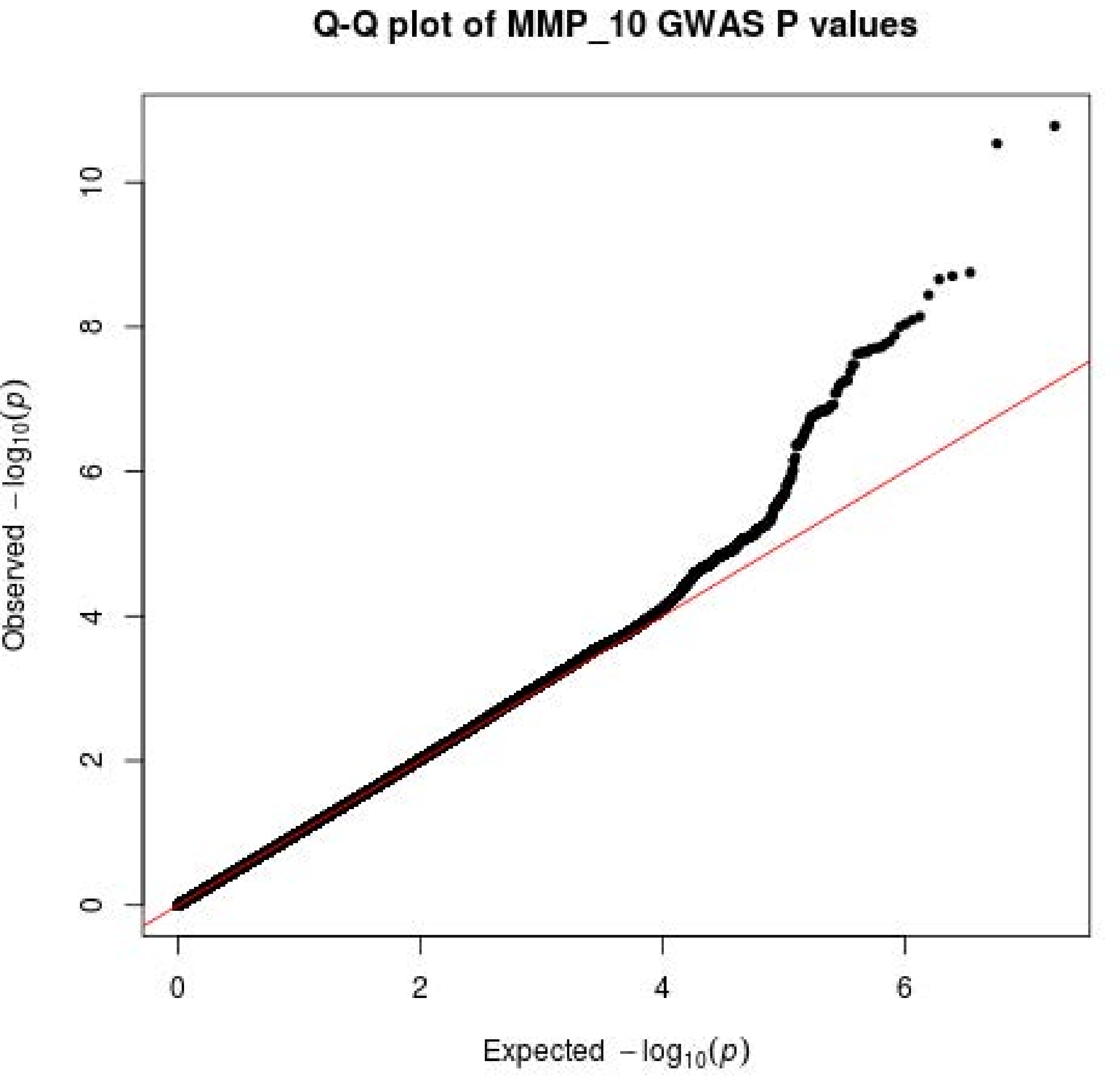

Q-Q plot of MMP_10 GWAS P values
Observed −log10(p)
Expected −log10(p)
0
2
4
6
8
10

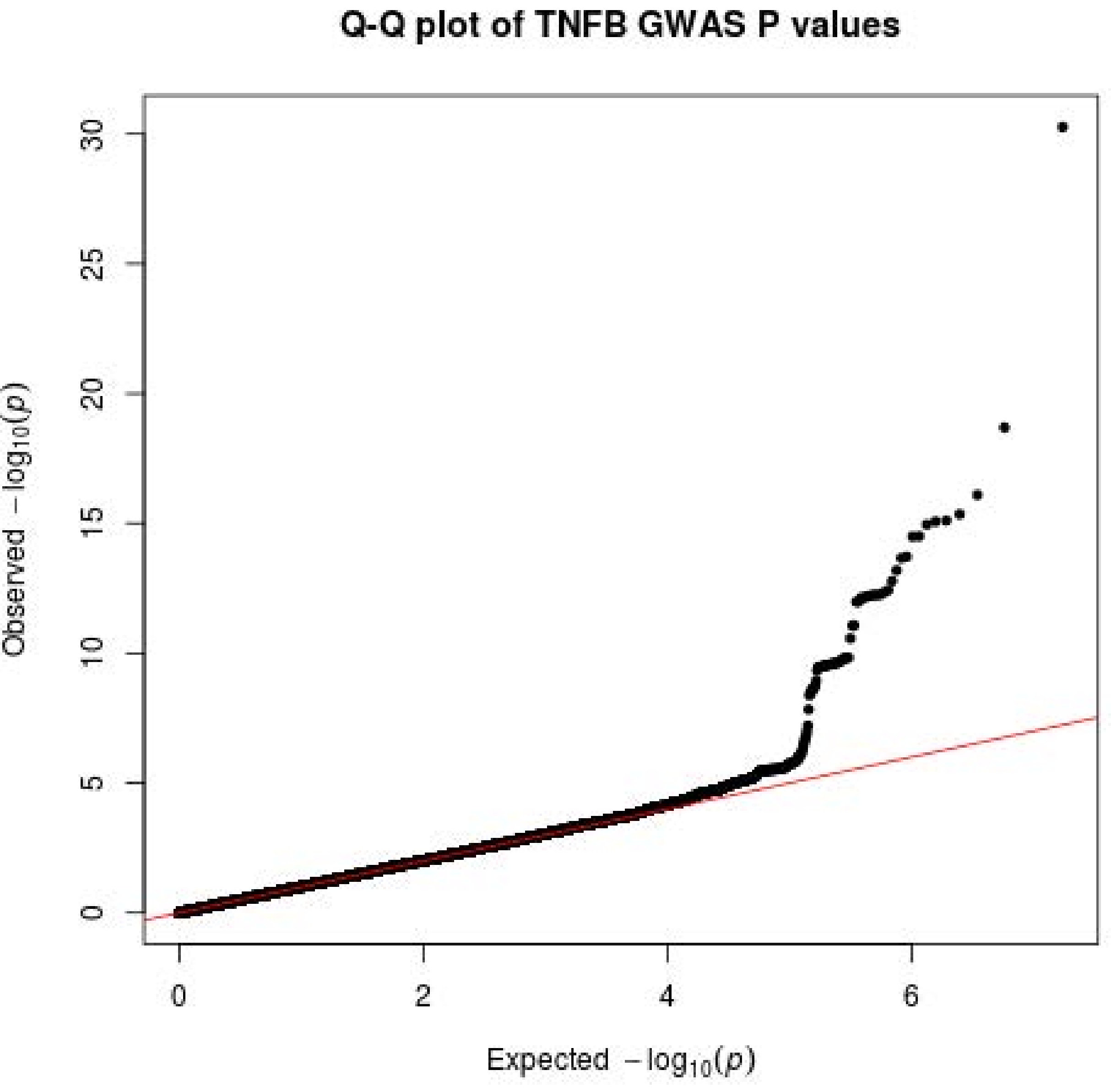

Q-Q plot of TNFB GWAS P values
Observed $-\log_{10}(p)$
Expected $-\log_{10}(p)$
0
5
10
15
20
25
30
0
2
4
6

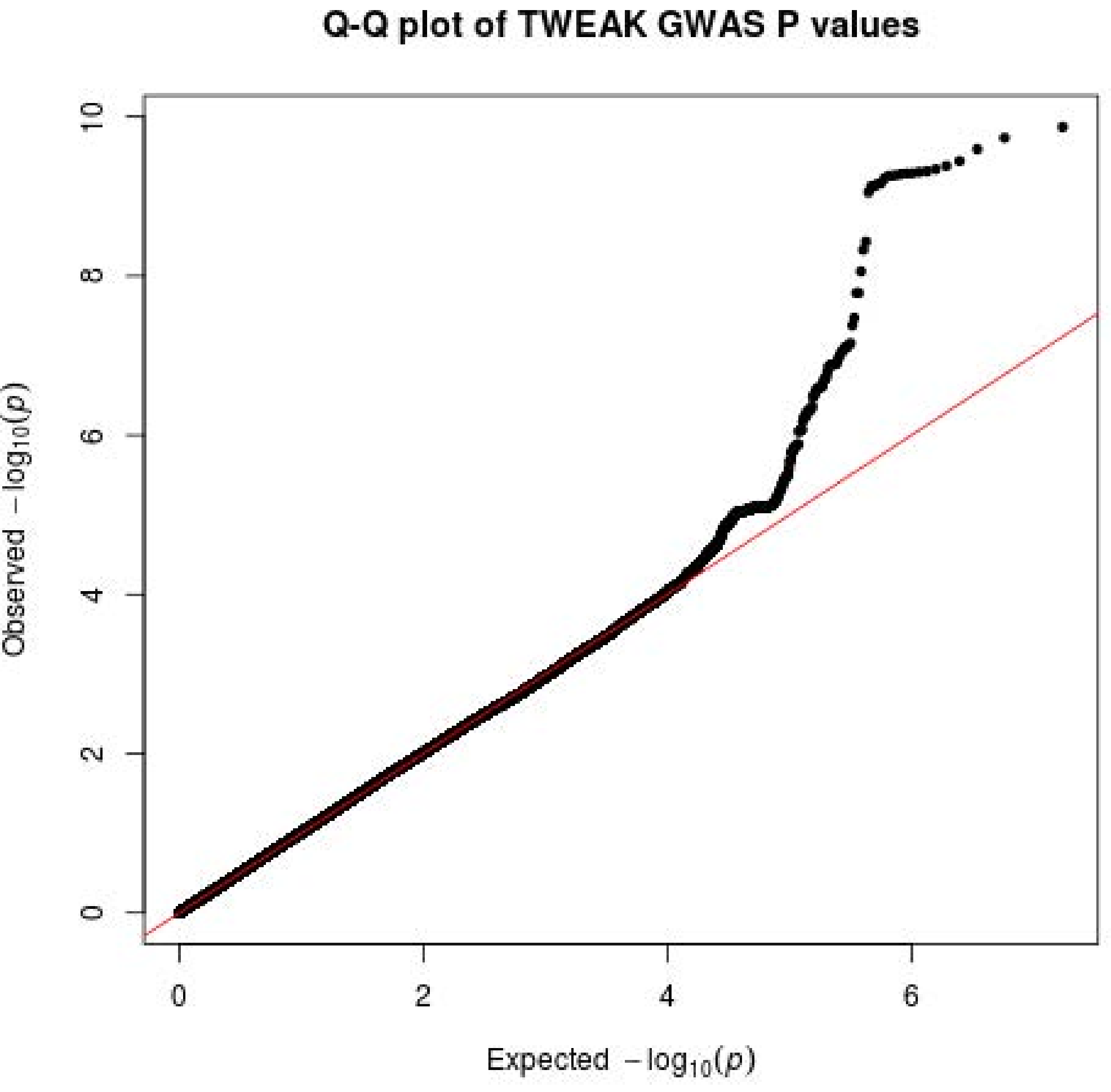

Q-Q plot of TWEAK GWAS P values
0
2
4
6
8
10
0
2
4
6
Observed −log10(p)
Expected −log10(p)
